# Fitness Functions Determine Optimal Parental Allocation: Redefining Trivers–Willard Theory

**DOI:** 10.1101/870881

**Authors:** Jibeom Choi, Hyungmin Roh, Sang-im Lee, Hee-Dae Kwon, Myungjoo Kang, Piotr G Jablonski

**Affiliations:** School of Biological Science, Seoul National University; Interdisciplinary Program of Computational Science and Technology, Seoul National University; School of Basic Science, DGIST; Department of Mathematics, Inha University; Department of Mathematical Sciences, Seoul National University

## Abstract

According to Trivers-Willard theory^1^, females in a good condition should carry more male offspring to maximize their fitness while should carry more females in a poor condition. Diverse theoretical and empirical studies has been performed to verify the validity of this claim^2,3^. Some portion of the empirical observations, however, exhibited contrary outcome to Trivers-Willard theory^4^. To resolve this problem, we computationally and mathematically show in here that reversed Trivers-Willard theory actually could be the outcome of the parental fitness optimization. In our models with identical fitness functions, we found that selective equitable care is optimal, and the number of the cared offspring should monotonically increase with maternal condition (or expendable parental investment). In some of our models with two distinguished male and female fitness functions, optimizations results were congruent with the conventional Trivers-Willard theory. In other models of ours, contrary to Trivers-Willard theory, it was optimal to invest in males when maternal condition was low. The results along with our hypothesis can explain the empirical observations that were previously thought to be the counterexample of Trivers-Willard theory. We propose that Trivers-Willard theory should be interpreted in multidimensional way, and more elaborate empirical data need to be collected to verify such propositions.

---

Trivers and Willard^1^ expected that the brood sex ratio is adjusted by female’s maternal condition in sexually dimorphic polygynous species. That is, the better the maternal condition is, the more bias goes toward the sex whose conditional variability highly affects the reproductive success. In a polygamous species, for instance, suppose that upper bound of good-conditioned male’s reproductive success is greater than that of a female. One, therefore, can expect female-biased offspring sex ratio when maternal condition is not favorable and male-biased sex ratio when maternal condition is advantageous. The empirical evidences that are for and against the Trivers-Willard theory are mutually present, and some of the empirical reports that are contrary to Trivers-Willard theory exhibited significant results^2,4^. There also have been analytical models which are based on the different fitness curves of offspring (e.g. Carranza 2001, Leimar 1996, Winckler 1987, Veller et al. 2016, Schindler et al. 2015).

To elucidate contribution of diverse factors, we built computational model that are composed of logistic functions, the shape of which would be appropriate in biological sense^5^. A brood in the computational model is composed of 10 individuals whose fitness (or reproductive value) functions are same or classified into type I and type II. Type I fitness function refers to the curve which exhibits certain extent when the given care is low but has a lower upper bound, and type II fitness function is a curve which requires considerable care to reach its plateau but has a higher upper bound. In many, but not all, species, type I is analogical to female fitness function, and type II to that of male’s.

We modified *S*, the index of maternal condition which would be correlated to the expendable care to the offspring. Let *v*_*i*_’s stand for the amount of care given to the *i*-th offspring, then 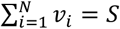 where *N* is the number of the offspring in a brood. The total offspring fitness (*F*) is therefore, 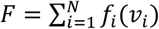 where *f*_*i*_ is the fitness function of *i*-th offspring. Then we computed the optimal care distribution toward each offspring by exhaustive search using fmincon function of MATLAB, which was not conducted in the aforementioned model papers.

In our models composed of identical fitness functions (denoted by model version H1 and H2), we explored which distribution of parental care among 10 identical offspring or embryos is optimal. (Henceforth, offspring refer to postnatal or prenatal offspring.) We identified that the number of the cared offspring monotonically increases and it can be represented by summation of step functions (Fig.1). Furthermore, picking some of the offspring and providing equitable care to them (selective equitable care) is proven to be optimal. These observation are verified by mathematical analysis as well (**Theorem 1**, **Supplementary Materials**). We infer this to be the theoretical basis of brood reduction as it is optimal not to invest in certain offspring in this model. The optimal *per capita* investment toward each cared offspring converged to the most efficient point of energy expense mentioned by Smith and Fretwell^6^ unless the expendable care is too abundant (**Theorem 2**). As an expansion of Smith and Fretwell’s (1974) model, we introduced the constraint of total expendable care. By considering the constraints, we showed the range of the total expendable care during which *per capita* investment lies near the most optimal point (Fig. 1d).

**Figure 1.**
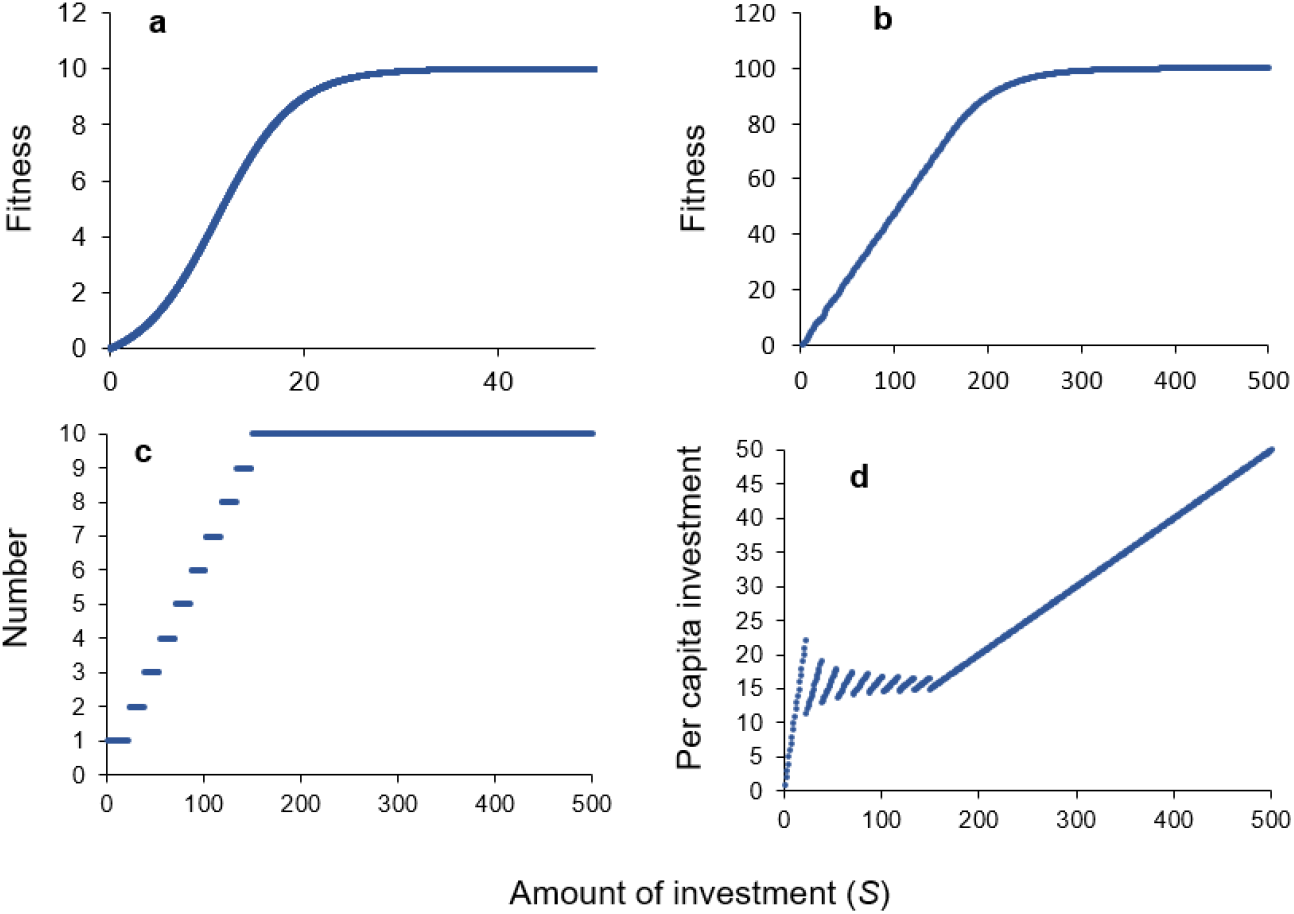
Optimization results of model version H1. (a) Fitness function plotted against cared investment. The curve is a logistic function whose inflection point lies on the first quadrant. (b) Optimized total offspring fitness plotted against total expendable care (*S*). (c) The number of the cared offspring plotted against total expendable care. (D) Optimal *per capita* investment to each cared offspring. At a certain interval of the *S*, *per capita* investment to the cared offspring lies around the most efficient point.

In subsequent models (denoted by model version 1 to 4, and model version 5-1 to 5-15), we divided offspring into two groups: type I and type II. Model version 1 and 2 exhibit classical cases of Trivers-Willard theory (Fig.2). In these models, it is optimal to raise more and feed more type I offspring rather than type II offspring when the maternal condition or total expendable care denoted as *S* is low. In an interval low in *S*, not a single type II offspring was cared. Type I offspring were dominant with respect to total care toward each type and the number of the cared offspring. BSR (brood sex ratio) and total investment should be biased toward type I to ensure the maximal parental fitness. In contrast, certain extent of *S* should be guaranteed for type II offspring to receive parental care. As *S* increases, more proportion of parental care was attributed to type II offspring. When *S* became plentiful enough, all offspring received care though optimal BSR and investment were skewed to type II. Calculation results of all versions are provided in the Fig. S3-S22 of **Supplementary Materials**.

**Figure 2.**
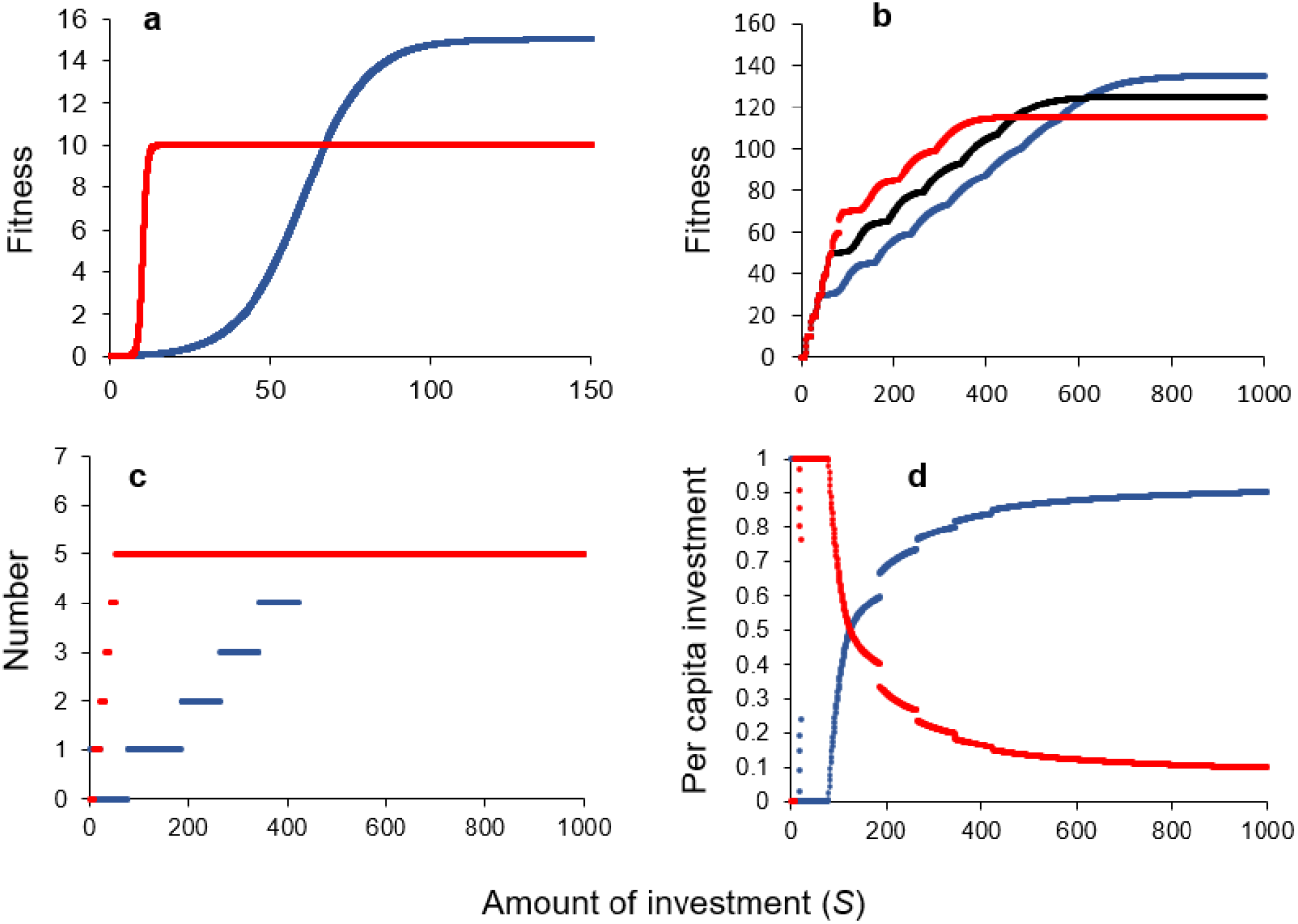
Optimization results of model version 2. (A) A type I fitness function (red) and a type II fitness (blue) function. Type I fitness function exhibits lower plateau but reaches most of its maximum with small amount of investment. Type II fitness function, on the other hand, has a greater upper bound but requires considerable care to reach most of its maximum. (B) Optimized total offspring fitness plotted against total expendable care. Each line indicates male to type I to type II ratio of 3:7 (red, type-I-biased), 5:5 (black), and 7:3 (blue, type-II-biased). (C) The number of the cared offspring plotted against total expendable care in 5:5 BSR. Only type I offspring receive the care when the condition is adverse (red line), while all offspring receive the care when condition become advantageous. (D) Optimal ratio of total type I and total type II investment. Optimal proportion to type II offspring increases as the maternal condition increases.

Results from model version 3 are contrary to Trivers-Willard theory (Fig.3). When *S* is low, but not too low, it is optimal to invest more in type II offspring and rear more of them. As *S* increases, it is optimal to invest in both types of offspring. We believe this to be the case of the reversed Trivers-Willard theory. Intuitively interpreting these results, we suspect these trends are related to the maximal efficiency. In version 1 and 2, the slope of maximal tangent that passes the origin is greater for type I than type II. Accordingly, the maximal efficiency of fitness against given care is greater for type I than type II. The opposite holds true for version 3: maximal tangent of type II is greater. Similar results were observed with same upper bound with different maximal slopes. (Model version 4: In this case, for simplicity, let the fitness function with higher fitness from 0 to cross point be type I function.) This standard is intuitive but not strict. The results of model version 5 reveals that this determinant of maximal efficiency is not general. Though type II fitness functions are endowed with greater maximal tangential line, strict biased care is not observed in some cases.

**Figure 3.**
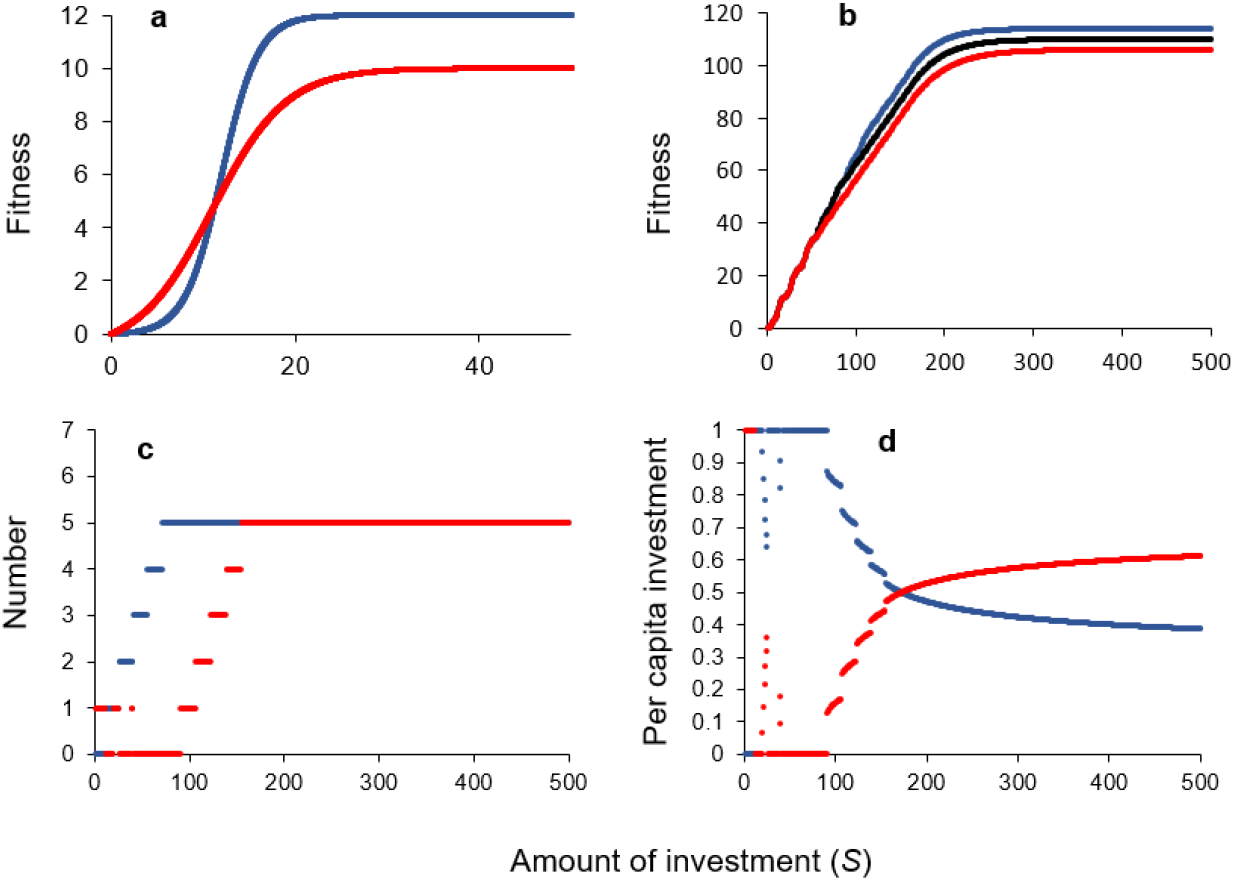
Optimization results of model version 3. (A) Same as Fig. 2A for model version 3. (B) Same as Fig. 2B for model version 3. (C) Same as Fig. 2C for model version 3. Contrary to Fig. 2B, type II offspring receive the majority of the care when condition is adverse (blue line), while all offspring receive the care when condition become advantageous. (D) Same as Fig. 2D for model version 3. Optimal proportion to type I increases as expendable care increases.

There have been indications of the reversed Trivers-Willard theory in both empirical^4^ and theoretical studies^7^. (A counterexample of Trivers-Willard theory or reversed Trivers-Willard theory in the present paper refers to the circumstances where male offspring are preferred in the harsh condition, while female offspring are preferred in mild condition.) Our results are not only in full support of Trivers-Willard theory but also can explain the empirical cases which were thought to be the counterexamples of the Trivers Willard theory. In some of the empirical studies, higher-ranking or better-conditioned females produced more daughters while lower-ranking or lower-conditioned females produced more sons^8,9^. We have two possible explanations for those cases. First, the fitness of sons could be rather prenatally determined presumably due to physical competence, while daughters could be relatively altricial. If the reproductive success of the sons are largely determined by innate factors rather than postnatal care, it suffices to provide sons with minimal care until they reach the maturity. For females, on the other hand, the success of gestation and breeding could be related to the nutritional status which can be augmented by postnatal parental care(Champagne & Meaney, 2006). Assume that fitness of males are less affected by the parental care, and the expected upper bound of the male fitness function is low. (The lower expected upper bound may occur in a broods of weak parents, due to the heritability of physical robustness.) Then the results that were considered to be the reversed Trivers-Willard theory can be explained by the model version 1 and 2. Under this circumstance, male fitness function is analogical to type I and that of female is analogical to type II. Hence, more male are cared for when expendable care is low.

Alternatively, the fitness functions of the offspring could follow the forms that resemble model version 3 and 4 (Fig.3). In those cases, it is optimal to invest in type II offspring when S is low. According to the model results where the number of type I and type II offspring are given, as *S* increases, more type I offspring receive care, and finally, all offspring receive care from the parents. As studies suggest that mammals can adaptively adjust the sex ratio at birth^11,12^, suppose that there are *N* male embryos and *N* females embryos. According to the model results, it is optimal to care for male embryos when condition is adverse, inducing male-skewed population at birth. As condition gets better, it is optimal to care for males and female embryos. On assumption that type II offspring has higher prenatal mortality (or other means such as lower conception probability) than type I offspring, model version 3 and 4 can explain female-skewed sex ratio at birth when maternal condition/rank is advantageous which was reported in diverse species^8,9^. As type I (females) and type II (males) prenatal offspring receive certain amount of care in decent maternal condition, mortality becomes the major factor to skew the sex ratio at birth.

With means of computational expectation and mathematical analysis, we provided possible explanations of the reversed Trivers-Willard theory. We suggest that the schematics of fitness functions could induce what was expected by Trivers-Willard theory and opposite cases which was not considered in the original model. The model and explanation are therefore not a denial of Trivers-Willard theory, but the expansion of such concept.

As pointed by Hewison and Gaillard^4^, and Veller et al.^13^, what is meant by prefer is ambiguous in the original text of Trivers and Willard^1^. The usage of the term ‘prefer’ has been equivocal in that in can be interpreted as bias on sex ratio at birth or bias on postnatal investment toward certain offspring^13^. Our results reveal that investment version and ratio version of Trivers-Willard theory^13^ may not coincide. For instance, preference of sex ratio toward type I ranges from *S* = 67 to 540 while that of investment ranges from *S* = 6 to *S* = 130 in model version 1. The usage of the term ‘variance’ is obscure as well because it may refer to extent of upper bound or change of fitness against given care. To avoid vagueness, we propose to use well-defined mathematical terms such as upper bound or slope when describing fitness curves.

Our models, however, do not take into account of cost of reproduction as in Leimar^3^, indicating that our model is fitted to a simple semelparous organisms. Unlike the model of Veller et al. (2016), we did not strictly distinguish maternal condition and expendable care. At cost of the simplifications, we were able to mathematically and computationally analyzed fitness functions to show that reversed Trivers–Willard effect is possible. To our knowledge, this is the first study to prove reversed Trivers–Willard effect in a system composed of multiple logistic fitness functions.

Though the model results reveal that ultimate bias is optimal in some circumstances, we do not expect animals to explicitly follow such tactics. This is due to the lack of the information about present and upcoming maternal conditions and the risk of extreme investment. Rather, we believe that the model results can explain the deviations from the 1:1 sex ratio or investment in natural circumstances.

To verify the validity of the ideas and to reconcile models with empirical findings, we encourage empirical scientists to measure the attributes of the fitness functions in diverse species. Not only the upper bound of the fitness function but the shape against the investment needs to be clarified. Though there have been attempts to measure and calculate the fitness functions^14,15^, it is questionable whether unambiguous definition of fitness can exist^16,17^. There also are confounding factors in measuring parental conditions^18^. Let alone the controversies of its existence and measurement, if such function can be measured, one can expect which tactics of investment is optimal by looking into the structure of each fitness function.

