## Supplemental Data for "Fitness Functions Determine Optimal Parental Allocation: Redefining Trivers–Willard Theory"

**Mathematical Propositions and Proofs**

**General Assumptions**

Fitness functions of individuals, $f_{i}(x)$ and $f_{j}(x)$ are logistic which are continuous and differentiable on $x\in\boldsymbol{R}$. The conditions of fitness functions include,

(1) The fitness functions have positive inflection points,

(2) The fitness functions are monotonically increasing,

(3) zero if $x=0$.

In our study, fitness functions are expressed as

$$f_{i}(x)= \frac{\alpha_{i}}{1+e^{\beta_{i}\left( x-\gamma_{i} \right)}}+ \delta_{i}$$

$X$ is defined as $X=(x_{1},x_{2}, x_{3},\ldots,x_{N})$ (where $0\leq x_{i}\leq S$ and $\sum x_{i}=S$) and if $X^{*}=(v_{1},v_{2}, v_{3},\ldots,v_{N})$ is optimal, then for any $Y\neq X^{*}$, $G(Y)\leq G(X^{*})$holds where $G(X)= \sum_{i=1}^{N} f_{i}(x_{i})$. As proven in subsequent propositions, $X^{*}$ may not be unique. Take $F^{*}$ as a function which satisfies $F^{*}\left( S \right)=G\left( X^{*} \right)=\sum_{i=1}^{N} f_{i}(v_{i})$.

**Theorem 1**

For any $v_{i}, v_{j}>0 (i\neq j)$ in optimized condition, ${f'}_{i}\left( v_{i} \right)={f'}_{j}(v_{j})$ holds.

**Proof**

Suppose ${f'}_{i}\left( v_{i} \right)\neq{f'}_{j}(v_{j})$ ($f_{i}\left( v_{i} \right), f_{j}(v_{j}) >0$; $v_{i}$, $v_{j}>0$) holds for an optimal $X^{*}$. For simplicity, set ${f'}_{i}\left( v_{i} \right)>{f'}_{j}(v_{j})$. Take $C\left( h \right)= \frac{f_{i}\left( v_{i}+h \right)-f_{i}\left( v_{i} \right)}{h}-\frac{f_{j}\left( v_{j} \right)-f_{i}\left( v_{j}-h \right)}{h}$. Then $\lim_{h\to+0} C(h)={f'}_{i}\left( v_{i} \right)-{f^{'}}_{j}\left( v_{j} \right)=K>0$. Therefore, for any $\varepsilon>0$, there exists some $\delta>0$ such that $0<\alpha<\delta$ indicates that $\left| C\left( \alpha\right)-K \right|< \varepsilon$. Taking $\left| \varepsilon\right|<K$ produces $K-\varepsilon<C\left( \alpha\right)<K+\varepsilon$ for some $0<\alpha<\delta$.

If we take $\alpha$ to satisfy the aforementioned conditions and $v_{j}-\alpha>0$, then $\frac{f_{i}\left( v_{i}+\alpha\right)-f_{i}\left( v_{i} \right)}{\alpha}-\frac{f_{j}\left( v_{j} \right)-f_{j}\left( v_{j}-\alpha\right)}{\alpha}=C\left( \alpha\right)>0$. Consequently, $f_{i}\left( v_{i}+\alpha\right)+f_{j}\left( v_{j}-\alpha\right)=f_{i}\left( v_{i} \right)+f_{j}\left( v_{j} \right)+C\left( \alpha\right)\alpha$. As both $C\left( \alpha\right)$ and $\alpha$ are positive, $f_{i}\left( v_{i}+\alpha\right)+f_{j}\left( v_{j}-\alpha\right)>f_{i}\left( v_{i} \right)+f_{j}\left( v_{j} \right)$ holds, indicating that $X^{*}$ is not optimal. Therefore, by reduction to absurdity, ${f'}_{i}\left( v_{i} \right)={f'}_{j}(v_{j})$.

**Definition 1**

Selective equitable distribution or SED is a strategy of parental care distribution by which select *n* offspring and provide equitable care to those *n* offspring.

**Definition 2**

$p^{*}$ is the inflection point of the fitness function. $N_{L} and$ $N_{H}$ are the number of elements in optimal points that are less or larger than $p^{*}$, respectively. $\beta and$ $\gamma$ are the *x*-values of the optimal point that are less or larger than $p^{*}$, respectively.

**Theorem 2**

SED is globally optimal for a brood of homogeneous fitness functions, but optimal strategy may not be unique. When a distribution strategy other than SED is optimal, $N_{L}=N_{H}=1$ and $\frac{\left( \beta+\gamma\right)}{2}=p^{*}$ should be satisfied.

**Proof**

Suppose that $v_{i}$’s are optimized. By **Theorem 1**, $f_{\alpha}^{'}(v_{i})$ should be the same for every $v_{i}>0$. For $v_{i}$ and $v_{j}$ to differ while $f_{\alpha}^{'}\left( v_{i} \right)=f_{\alpha}^{'}(v_{j})$, they should be symmetric to inflection point $p^{*}$ as $f_{\alpha}^{'}$ is a single-peaked function.

Suppose that optimized $v_{i}$’s are not homogeneous in a group of identical fitness functions. This indicates that $v_{i}$’s are divided into 2 sets: a set whose values are less than $p^{*}$, a set whose values are greater than $p^{*}$. By **Theorem 1**, differential values in each set should be the same. As it was assumed that optimized $v_{i}$’s are not homogeneous in a group of identical fitness functions, $N_{L}\geq1$ and $N_{H}\geq1$ hold.

(i) $N_{L}\geq2$

Let $\beta_{1}$ and $\beta_{2}$ denote the optimized care that are lower than $p^{*}$. As $f_{\alpha}^{'}$ is single-peaked, by **Theorem 1**, $\beta_{1}=\beta_{2}$. Take $\varepsilon$ as $0<\varepsilon<min(\beta_{1}, p^{*}-\beta_{1})$ so that ${0<\beta}_{1}-\varepsilon$ and $\beta_{1}+\varepsilon<p^{*}$ holds. Then $\int_{\beta_{1}}^{\beta_{1}+\varepsilon} f_{\alpha}^{'}\left( t \right)dt>\int_{\beta_{1}-\varepsilon}^{\beta_{1}} f_{\alpha}^{'}\left( t \right)dt$ because $f_{\alpha}^{'}\left( t_{1} \right)>f_{\alpha}^{'}(t_{2})$ for $t_{1}\in[\beta_{1},\beta_{1}+\varepsilon]$ and $t_{2}\in[\beta_{1}-\varepsilon,\beta_{1}]$. By adding $f_{\alpha}\left( \beta_{1} \right)+f_{\alpha}(\beta_{2})$ to both sides yields

$$\int_{\beta_{1}}^{\beta_{1}+\varepsilon} f_{\alpha}^{'}\left( t \right)dt+f_{\alpha}\left( \beta_{1} \right)+f_{\alpha}\left( \beta_{2} \right)>\int_{\beta_{1}-\varepsilon}^{\beta_{1}} f_{\alpha}^{'}\left( t \right)dt+f_{\alpha}\left( \beta_{1} \right)+f_{\alpha}(\beta_{2})$$

$$f_{\alpha}\left( \beta_{1} \right)+\int_{\beta_{1}}^{\beta_{1}+\varepsilon} f_{\alpha}^{'}\left( t \right)dt+f_{\alpha}\left( \beta_{2} \right)+\int_{\beta_{2}}^{\beta_{2}-\varepsilon} f_{\alpha}^{'}\left( t \right)dt>f_{\alpha}\left( \beta_{1} \right)+f_{\alpha}(\beta_{2})$$

$$f_{\alpha}\left( \beta_{1}+\varepsilon\right)+f_{\alpha}\left( \beta_{2}-\varepsilon\right)>f_{\alpha}\left( \beta_{1} \right)+f_{\alpha}(\beta_{2})$$

Thus, $\beta_{1}$ and $\beta_{2}$ are not optimal points which is contradictory to the assumption. By reduction to absurdity, $N_{L}<2$. As $N_{L}$ is non-negative integer, $N_{L}$ is either 0 or 1.

(ii) $N_{H}\geq2$

Let $\gamma_{1}$ and $\gamma_{2}$ denote the optimized care that are greater than $p^{*}$. By **Theorem 1**, $\gamma_{1}=\gamma_{2}$ holds. As heterogeneous optimization in a group was assumed, $N_{L}\neq0$. Take $\beta_{1}$ as the optimized care that is smaller than $p^{*}$. Due to the symmetry of the $f_{\alpha}^{'}$, $\frac{\beta_{1}+\gamma_{1}}{2}= p^{*}$.

Take $0<\varepsilon<{(p}^{*}-\beta_{1})/2$ so that $\beta_{1}+2\varepsilon<p^{*}$ holds, then $\int_{\beta_{1}+ \varepsilon}^{\beta_{1}+ 2\varepsilon} f_{\alpha}^{'}\left( t \right)dt>\int_{\gamma_{1}}^{\gamma_{1}+ \varepsilon} f_{\alpha}^{'}\left( t \right)dt$ due to the symmetry and single-peakedness of $p^{*}$. By the same grounds, $\int_{\beta_{1}}^{\beta_{1}+ \varepsilon} f_{\alpha}^{'}\left( t \right)dt>\int_{\gamma_{1}}^{\gamma_{1}+ \varepsilon} f_{\alpha}^{'}\left( t \right)dt$. Consequently,

$$\int_{\beta_{1}}^{\beta_{1}+ \varepsilon} f_{\alpha}^{'}\left( t \right)dt+\int_{\beta_{1}+ \varepsilon}^{\beta_{1}+ 2\varepsilon} f_{\alpha}^{'}\left( t \right)dt>2\int_{\gamma_{1}}^{\gamma_{1}+ \varepsilon} f_{\alpha}^{'}\left( t \right)dt$$

Adding $f_{\alpha}\left( \beta_{1} \right)+f_{\alpha}\left( \gamma_{1} \right)+f_{\alpha}\left( \gamma_{2} \right)$ on both side produces

$$f_{\alpha}\left( \beta_{1} \right)+f_{\alpha}\left( \gamma_{1} \right)+f_{\alpha}\left( \gamma_{2} \right)+\int_{\beta_{1}}^{\beta_{1}+ \varepsilon} f_{\alpha}^{'}\left( t \right)dt+\int_{\beta_{1}+ \varepsilon}^{\beta_{1}+ 2\varepsilon} f_{\alpha}^{'}\left( t \right)dt-2\int_{\gamma_{1}}^{\gamma_{1}+ \varepsilon} f_{\alpha}^{'}\left( t \right)dt >f_{\alpha}\left( \beta_{1} \right)+f_{\alpha}\left( \gamma_{1} \right)+f_{\alpha}\left( \gamma_{2} \right)$$

$$f_{\alpha}\left( \beta_{1}+2\varepsilon\right)+f_{\alpha}\left( \gamma_{1}-\varepsilon\right)+f_{\alpha}\left( \gamma_{2}-\varepsilon\right) >f_{\alpha}\left( \beta_{1} \right)+f_{\alpha}\left( \gamma_{1} \right)+f_{\alpha}\left( \gamma_{2} \right)$$

Therefore $\beta_{1}$, $\gamma_{1}$ and $\gamma_{2}$ are not optimized care which is contradictory to the assumption. By reduction to absurdity, $N_{H}<2$. As $N_{H}$ is non-negative integer, $N_{H}$ is either 0 or 1.

By (i) and (ii), if heterogeneous optimal cares exist in a group of identical fitness functions, $N_{L}=N_{H}=1$. Otherwise, it cannot be optimal.

Suppose heterogeneous optimal cares exist in a group of identical fitness functions with the condition of $N_{L}=N_{H}=1$. By symmetry, $f_{\alpha}^{'}\left( \beta\right)=f_{\alpha}^{'}\left( \gamma\right)$ and $\gamma-p^{*}=p^{*}-\beta$ where $\beta<\gamma$. Take $0<\varepsilon<min(\beta, {(p}^{*}-\beta))$ and $0<\omega<\beta$. Again by symmetry,

$$\int_{\beta}^{\beta+ \varepsilon} f_{\alpha}^{'}\left( t \right)dt=\int_{\gamma- \varepsilon}^{\gamma} f_{\alpha}^{'}\left( t \right)dt$$

and

$$\int_{\beta-\omega}^{\beta} f_{\alpha}^{'}\left( t \right)dt=\int_{\gamma}^{\gamma+\omega} f_{\alpha}^{'}\left( t \right)dt$$

This produces that whichever $\varepsilon$ and $\omega$ we choose,

$$f_{\alpha}\left( \beta+\varepsilon\right)+f_{\alpha}\left( \gamma-\varepsilon\right)=f_{\alpha}\left( \beta\right)+ \int_{\beta}^{\beta+\varepsilon} f_{\alpha}^{'}\left( t \right)dt+f_{\alpha}\left( \gamma\right)+\int_{\gamma}^{\gamma-\varepsilon} f_{\alpha}^{'}\left( t \right)dt =f_{\alpha}\left( \beta\right)+f_{\alpha}\left( \gamma\right)$$

and similarly,

$$f_{\alpha}\left( \beta-\omega\right)+f_{\alpha}\left( \gamma+\omega\right)=f_{\alpha}\left( \beta\right)+f_{\alpha}\left( \gamma\right)$$

Therefore, the optimal solution is not fixed but multiple optima are derived. However this condition is narrow because $N_{L}=N_{H}=1$ and $\frac{\left( \beta+\gamma\right)}{2}=p^{*}$ (or, $S=2p^{*})$ should be satisfied. Otherwise, $v_{i}$’s are the same ($v_{i}>0$) for the group of identical fitness functions.

**Theorem 3 (homogeneous fitness functions)**

For a brood composed of homogeneous logistic fitness functions, suppose SED is uniquely optimal (*i*.*e*. $S\neq2p^{*}$). Then the number of the cared offspring can be represented as summation of floor functions.

**Proof**

As SED is optimal by **Theorem 2**, one needs to find $n$ ($\in\{1,2,3,\ldots,N\}$) which maximizes $nf(\frac{S}{n})$.

$nf\left( \frac{S}{n} \right)=\frac{f\left( \frac{S}{n} \right)}{\frac{1}{n}}=S\left[ \frac{f\left( \frac{S}{n} \right)}{\frac{S}{n}} \right]$ holds where $S$ is fixed, and $\psi\left( x \right)=\frac{f\left( x \right)}{x}$ is a single-peaked function who has a peak at $a^{*}$ (the point of tangency). Therefore, $\psi\left( \frac{S}{n} \right)-\psi\left( \frac{S}{n+1} \right)>0$ if $\frac{S}{n}<a^{*}$ and $\psi\left( \frac{S}{n} \right)-\psi\left( \frac{S}{n+1} \right)<0$ if $\frac{S}{n+1}<a^{*}$. Then the problem is to find $n^{*}$ satisfying

$$n^{*}=\underset{n}{\mathrm{argmax}} nf\left( \frac{S}{n} \right)= \underset{n}{\mathrm{argmax}} \frac{f\left( \frac{S}{n} \right)}{\frac{1}{n}}$$

$$= \underset{n}{\mathrm{argmax}}S\left( \frac{f\left( \frac{S}{n} \right)}{\frac{S}{n}} \right)= \underset{n}{\mathrm{argmax}}\psi\left( \frac{S}{n} \right)$$

Take $n_{S}$ ($\in\{1,2,3,\ldots,N\}$) as the integer which satisfies the inequality condition $\frac{S}{n_{S}+1}<a^{*}<\frac{S}{n_{S}}$. The condition where $\frac{S}{N}>a^{*}$ or $S<a^{*}$ will be discussed later. For now, restrict $S$ so that $a^{*}<S<{Na}^{*}$. As $\frac{S}{a^{*}}-1<n_{S}<\frac{S}{a^{*}}$, $n_{S}=\left[ \frac{S}{a^{*}} \right]$ where [] is a floor function. $n^{*}$ is either $n_{S}$ or $n_{S}+1$ given that $a^{*}\leq S\leq{Na}^{*}$ because $\psi(x)$ has its peak when $x=a^{*}$. In case that there is $n_{S}$ such that $\frac{S}{n}=a^{*}$, the optimizing $n^{*}=n_{S}=\frac{S}{a^{*}}=\left[ \frac{S}{a^{*}} \right]$.

Suppose $S<a^{*}$, then $n^{*}=1$ because $f\left( S \right)>nf(\frac{S}{n})$ for $n\geq2$ as $\psi\left( S \right)>\psi\left( \frac{S}{n} \right)$. Suppose $S>Na^{*}$, then $Nf\left( S/N \right)>nf(\frac{S}{n})$ for $n<N$, again as $\psi\left( \frac{S}{N} \right)>\psi\left( \frac{S}{n} \right)$.

To sum up,

$n^{*}=1$	($S<a^{*}$)

$n^{*}=k$ or $n^{*}=k+1$	($ka^{*}\leq S\leq(k+1)a^{*}$, $k\in\{1,2,3,\ldots,(N-1)\}$)

$n^{*}=N$	($S>Na^{*}$)

For a fixed $k\in\{1,2,3,\ldots,(N-1)\}$, restrict $S$ so that $ka^{*}\leq S<(k+1)a^{*}$ holds. In this restricted domain, $n_{S}=k$. Let $\Psi$ be defined as $\Psi\left( S \right)=\psi\left( \frac{S}{n_{S}+1} \right)-\psi\left( \frac{S}{n_{S}} \right)$. Then $\Psi\left( ka^{*} \right)<0$ because $\psi(x)$ is maximized at $x=a^{*}$. On the other hand, $\lim_{h\to0^{+}} \psi\left( \frac{\left( k+1 \right)a^{*}-h}{n_{S}+1} \right)=\lim_{h\to0^{+}} \psi\left( a^{*}-\frac{h}{n_{S}+1} \right)=\psi(a^{*})$. Thus for some $\varepsilon>0$, $\Psi\left( \left( k+1 \right)a^{*}-\varepsilon\right)>0$ ($\left( k+1 \right)a^{*}-\varepsilon>ka^{*}$).

As $\psi$ is continuous and differentiable, $\Psi$ is accordingly continuous and differentiable in the restriction domain. In addition, $\Psi$ is (strictly) monotonically increasing because $\psi\left( \frac{S}{n_{S}+1} \right)$ is (strictly) monotonically increasing and $\psi\left( \frac{S}{n_{S}} \right)$ is (strictly) monotonically decreasing given that $ka^{*}\leq S<(k+1)a^{*}$. Due to **Mean Value Theorem**, there is $S^{d}\in(ka^{*},\left( k+1 \right)a^{*})$ such that $\Psi\left( S \right)>0$ if $S>S^{d}$ and $\Psi\left( S \right)<0$ if $S<S^{d}$. By the definition of $\Psi$,

$$n^{*}= \left\{ \begin{aligned} k if ka^{*}\leq S<S^{d} \\ k+1 if S^{d}<S\leq(k+1)a^{*} \end{aligned} \right.$$

where $k\in\{1,2,3,\ldots,(N-1)\}$.

As a consequence, let $A$ be the set of all $S^{d}$’s and let $\tau_{k}(x)$ be the step function such that

$$\tau_{c}\left( x \right) = \left\{ \begin{aligned} 0 if x<c \\ 1 if x>c \end{aligned} \right.$$

where $\tau_{k}$ is not defined at $x=c$. Let $\rho^{*}(S)$ represent the optimized number of cared offspring ($n^{*}$) by $S$. Then $\rho^{*}$ can be represented by the sum of those step functions.

$$\rho^{*}(x)=1+\sum_{c_{i}\in A} \tau_{c_{i}}\left( x \right)$$

where $x\in\boldsymbol{R}-A$. If $x\in A$, no unique solution exists, *i*.*e*. $\psi\left( \frac{S}{n_{S}+1} \right)=\psi\left( \frac{S}{n_{S}} \right)$ so that $n_{S}$ and $n_{S}+1$ exhibit same amount of the total offspring fitness.

**Definition**

Define $L_{i,k}$ and $H_{i,k}$ as $\frac{S}{k+1}$ and $\frac{S}{k}$, respectively, such that $\left( k+1 \right) f_{i}\left( \frac{S}{k+1} \right)=k f_{i}(\frac{S}{k})$ holds.

**Corollary of Theorem 3**

If $2f_{i}\left( \frac{S}{2} \right)=f_{i}\left( S \right)$, then $S={2p}^{*}$ (i.e., $L_{i,1}=p^{*}$ and $H_{i,1}=2p^{*}$)

**Proof**

Let us put $S={2p}^{*}$, then $2f_{i}\left( \frac{S}{2} \right)={2f}_{i}(p^{*})=f_{i}({2p}^{*})=f_{i}(S)$ holds by point symmetry of $f_{i}$.

To show that $S={2p}^{*}$ is the unique solution, suppose $2f_{i}\left( \frac{S}{2} \right)=f_{i}\left( S \right)$ holds for some $S\neq{2p}^{*}$. Then $\psi_{i}\left( \frac{S}{2} \right)={2f_{i}\left( \frac{S}{2} \right)}/S={f_{i}\left( S \right)}/S=\psi_{i}\left( S \right)$ should hold.

If $\psi_{i}\left( \frac{S}{2} \right)=\psi_{i}\left( S \right)$is smaller than $\psi_{i}\left( p^{*} \right){=\psi}_{i}\left( 2p^{*} \right)$, then $S$ should satisfy both $\frac{S}{2}<p^{*}$ and ${2p}^{*}<S$ because $\psi_{i}\left( \frac{S}{2} \right)<\psi_{i}\left( p^{*} \right)$ and $\psi_{i}\left( S \right)<\psi_{i}\left( {2p}^{*} \right)$ due to single-peakedness of $\psi_{i}$. Thus, $\psi_{i}\left( S \right)$ cannot be smaller than $\psi_{i}\left( {2p}^{*} \right)$.

On the other hands, if $\psi_{i}\left( \frac{S}{2} \right)=\psi_{i}\left( S \right)$ is larger than $\psi_{i}\left( p^{*} \right){=\psi}_{i}\left( 2p^{*} \right)$, then $S$ should satisfy both $p^{*}<\frac{S}{2}$ and ${S<2p}^{*}$ again by single-peakedness. Thus $\psi_{i}\left( S \right)$ cannot be larger than $\psi_{i}\left( 2p^{*} \right)$.

Now we have $\psi_{i}\left( S \right)=\psi_{i}\left( p^{*} \right)=\psi_{i}\left( 2p^{*} \right)$ which gives$S=p^{*}$ or $S={2p}^{*}$. But if we put $S=p^{*}$, then

$$2f_{i}\left( \frac{S}{2} \right)={2f}_{i}\left( \frac{p^{*}}{2} \right){=p^{*}\psi}_{i}\left( \frac{p^{*}}{2} \right)\neq{p^{*}\psi}_{i}\left( p^{*} \right)=f_{i}\left( p^{*} \right)=f_{i}\left( S \right)$$

which proves that $S={2p}^{*}$ is the unique solution.

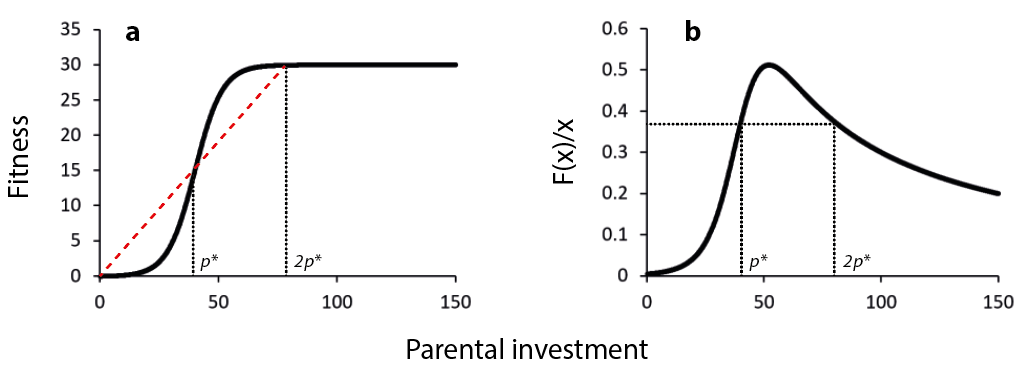

**Figure S1 | The graphical illustration of** $\boldsymbol{p}^{\boldsymbol{*}}$**.**

(a) For a logistic fitness function, $(p^{*}, f(p^{*}))$ is an inflection point. Due to symmetry of logistic functions, $\frac{f\left( p^{*} \right)}{p^{*}}=\frac{f\left( 2p^{*} \right)}{2p^{*}}$ holds.

(b) By the aforementioned symmetry, $\psi\left( p^{*} \right)=\psi\left( 2p^{*} \right)$ holds where $\psi\left( x \right)=f(x)/x$.

**Definition**

Suppose that the brood is composed of two groups, each with fitness function of $f_{1}$ and $f_{2}$. Then define $S_{1}^{*}$ and $S_{2}^{*}$ as parental investment to each group that maximizes total fitness with constraint that $S_{1}^{*}+S_{2}^{*}=S$. That is, $F_{1}\left( S_{1}^{*} \right)+F_{2}\left( S_{2}^{*} \right)=F^{*}(S)$ where $F_{i}(X)$ is the optimal offspring fitness of *i*-th group to which *X* amount of care is given. Define $F_{1}\left( S \right)=n_{1}^{*} f_{1}(\frac{S}{n_{1}^{*}})$, $F_{2}\left( S \right)=n_{2}^{*} f_{2}(\frac{S}{n_{2}^{*}})$ and $F\left( S_{1}, S_{2} \right)=F_{1}\left( S_{1} \right)+F_{2}(S_{2})$.

**Algorithm 1**: Finding optimal distribution in brood composed of two different fitness functions

$$F\left( S \right)=F_{1}\left( S_{1} \right)+F_{2}\left( S_{2} \right)=F_{1}\left( S_{1} \right)+F_{2}\left( S-S_{1} \right)$$

$$n_{i}^{*}=\underset{n_{i}\in\{n_{s}, n_{s}+1\}}{\mathrm{argmax}} n_{i}f_{i}(\frac{S_{i}}{n_{i}})$$

Therefore

$$F\left( S \right)=n_{1}^{*}f_{1}\left( \frac{S_{1}}{n_{1}^{*}} \right)+n_{2}^{*}f_{2}(\frac{S-S_{1}}{n_{2}^{*}})$$

As $F\left( S \right)$ is a univariate function of $S_{1}$,

$$S_{1}^{*}= \underset{S_{1}}{\mathrm{argmax}} \{n_{1}^{*}f_{1}\left( \frac{S_{1}}{n_{1}^{*}} \right)+n_{2}^{*}f_{2}\left( \frac{S-S_{1}}{n_{2}^{*}} \right)\}$$

and

$$S_{2}^{*}=S-S_{1}^{*}$$

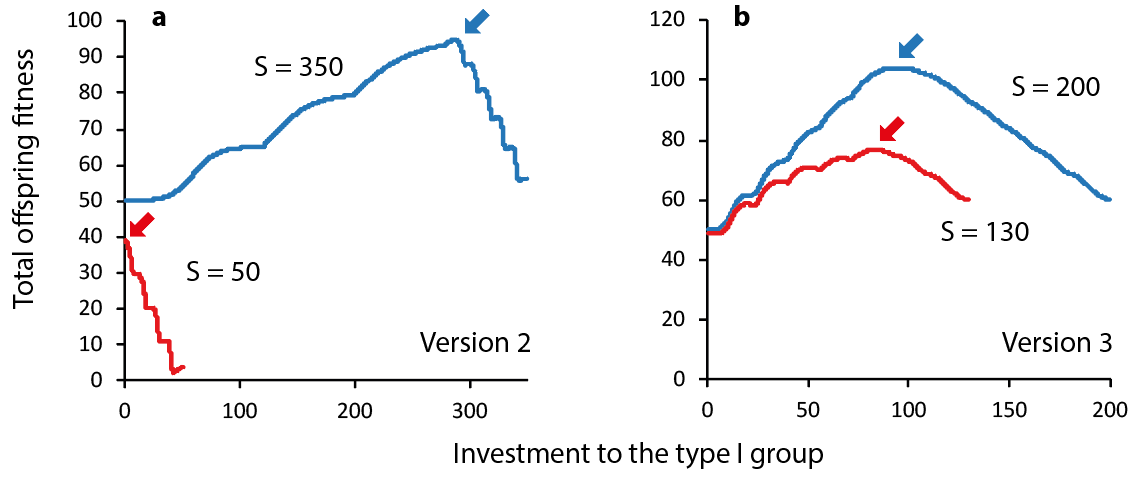

**Figure S2 | Comparison of the total offspring fitness in diverse conditions computed by Algorithm 1.**

(a) In model version 2, when S=50 which is correspondent to the adverse condition, it is optimal to invest all of resources to type II group as offspring fitness is maximized when there is no investment to type I offspring (S1=0). When S=350, offspring fitness is optimized if S1=285 (S1/S=0.8166). The peak of each graph is indicated by an arrow.

(b) In model version 3, when S=130, it is optimal if S1=82.9 (S1/S= 0.6377). When S=200, it is optimal if S1=94.1 (S1/S = 0.4705).

List of Variables and Functions in the Proof

| Name | Description |
| --- | --- |
| $f_{i}$ | *i*-th fitness function |
| $p_{i}^{*}$ | *x*-coordinate of *i*-th fitness function’s inflection point |
| $a_{i}^{*}$ | *x*-coordinate of *i*-th fitness function’s tangential point |
| $\psi(x)$ | $\psi\left( x \right)=f(x)/x$ |
| *S* | The amount of expendable care, or parental condition |
| $L_{i,j}$ | Lowest *x*-coordinates of the optimized care when SED was applied for (*j+1*) offspring with *i*-th fitness function. |
| $H_{i,j}$ | Highest *x*-coordinates of the optimized care when SED was applied for *j* offspring with *i*-th fitness function. |
| $F^{*}\left( S \right)$ | The optimized sum of offspring fitness given that total care is *S*. |
| $n^{*}$ | The number of offspring that maximizes sum of offspring fitness. |

**Computational Methods**

Total Expendable Care by Sire and Dam

Parents (a sire and a dam) in a decent condition can bear physically sound offspring who are robust at birth; additionally, parents in a decent condition can provide ample postnatal feeding and care during their immaturity. To reduce complexity, we assumed that the maternal condition is in proportion to the amount of care provided to the offspring. In other words, the concept of care in the present paper includes parental condition which affects prenatal offspring condition and provision ability which affects postnatal offspring condition. This concept is embodied by *S*, the total expendable care.

Constraints of Care Distribution

Let $x_{i}$ be the amount of provision toward the *i* th offspring. There are constraints on $x_{i}$’s such that

${0\leq x}_{i}\leq S$ for all *i*’s

$$\sum_{i=1}^{N} x_{i}=S$$

where $S (\geq0)$ is the amount of total parental provisioning given by parents and *N* is the number of the litters in the brood.

The aim of the model is to solve the optimizing solution of the problem. Let $V_{O}$ denote an ordered tuple such that $X_{O}= (x_{1}, x_{2}, x_{3}, \ldots, x_{N})$. For simplicity, let $F\left( X \right)$ be defined as $F\left( X \right)=\sum_{i=1}^{N} F_{i}(x_{i})$. Then one needs to find the tuple that maximizes $F\left( X \right)$, or $\underset{X\mathcal{\in H}}{arg max} F(X)$ where $\mathcal{H}$ is the set of tuples satisfying aforementioned constraints. As there are inequality constraints in domain, one would be tempted to utilize Karush–Kuhn–Tucker (KKT) conditions to find optimal *V*. To solve optimizing solution of the problem, however, we compared fitness of all possible distribution of *V* for simplicity rather than using KKT conditions. We used *fmincon* in MATLAB 2017a to find multiple local maxima. By comparing possible combinations of *X*’s, it is possible to find *X* that maximizes $F\left( X \right)$. Let this optimal *X* be donoted as *V*. By looking into the distribution of *V*, one can retrieve how many litters should be cared, how males and females should be differentiated in feeding in order to maximize total offspring fitness (the sum of all offspring fitness).

Optimized Care Distribution in Various Conditions

To analyze the dynamics of the feeding strategy, we modified the amount of the total expendable care (*S*). The total number of the offspring in the model was fixed to 10. In the models in which males and females have different fitness functions, we classified the brood into male-biased, equal, and female-biased condition. In male-biased brood, 7 males and 3 females are present and vice versa for the female-biased brood. For instance, to expect the optimal feeding strategy of male biased brood when the total expendable care is *S*, the objective of the maximization is

$$F\left( X \right)=\sum_{i=1}^{N} F_{i}\left( x_{i} \right)$$

$$=F_{m}\left( x_{1} \right)+F_{m}\left( x_{2} \right)+F_{m}\left( x_{3} \right)+ F_{f}\left( x_{4} \right)+F_{f}\left( x_{5} \right)+F_{f}\left( x_{6} \right)+F_{f}\left( x_{7} \right)+F_{f}\left( x_{8} \right)+F_{f}\left( x_{9} \right)+F_{f}\left( x_{10} \right)$$

with constraints of $\sum_{i=1}^{N} x_{i}=S$ where $F_{m}$ is a fitness function of a male and $F_{f}$ is a fitness function of a female.

We parametrically varied the value of S from 1 to 1000 (or to 500 if upper bound of functions are low) in steps of 1. In each S, the optimized X was calculated via MATLAB’s optimization toolbox. This allowed us to retrieved how much care should be given to each offspring in order to maximize total fitness. The same optimization calculation was performed to each of male-biased, equal, and female-biased brood to see which gender ratio is optimal given the same resources. We tested *N* number of models with different pairs of fitness functions.

**Computation Results**

**Model Version H1**

a

b

d

c

H1 Function

|  |  |
| --- | --- |
| 0 | -3.88578e-15 |
| 0.01 | 0.00150382 |
| 0.02 | 0.00301095 |
| 0.03 | 0.0045214 |
| 0.04 | 0.00603516 |
| 0.05 | 0.00755226 |
| 0.06 | 0.0090727 |
| 0.07 | 0.0105965 |
| 0.08 | 0.0121236 |
| 0.09 | 0.0136541 |
| 0.1 | 0.0151879 |
| 0.11 | 0.0167251 |
| 0.12 | 0.0182657 |
| 0.13 | 0.0198096 |
| 0.14 | 0.021357 |
| 0.15 | 0.0229077 |
| 0.16 | 0.0244618 |
| 0.17 | 0.0260194 |
| 0.18 | 0.0275803 |
| 0.19 | 0.0291447 |
| 0.2 | 0.0307125 |
| 0.21 | 0.0322837 |
| 0.22 | 0.0338583 |
| 0.23 | 0.0354364 |
| 0.24 | 0.037018 |
| 0.25 | 0.038603 |
| 0.26 | 0.0401914 |
| 0.27 | 0.0417833 |
| 0.28 | 0.0433787 |
| 0.29 | 0.0449776 |
| 0.3 | 0.04658 |
| 0.31 | 0.0481859 |
| 0.32 | 0.0497952 |
| 0.33 | 0.0514081 |
| 0.34 | 0.0530245 |
| 0.35 | 0.0546444 |
| 0.36 | 0.0562678 |
| 0.37 | 0.0578947 |
| 0.38 | 0.0595252 |
| 0.39 | 0.0611593 |
| 0.4 | 0.0627968 |
| 0.41 | 0.064438 |
| 0.42 | 0.0660827 |
| 0.43 | 0.067731 |
| 0.44 | 0.0693828 |
| 0.45 | 0.0710383 |
| 0.46 | 0.0726973 |
| 0.47 | 0.0743599 |
| 0.48 | 0.0760262 |
| 0.49 | 0.077696 |
| 0.5 | 0.0793695 |
| 0.51 | 0.0810465 |
| 0.52 | 0.0827272 |
| 0.53 | 0.0844116 |
| 0.54 | 0.0860996 |
| 0.55 | 0.0877912 |
| 0.56 | 0.0894865 |
| 0.57 | 0.0911854 |
| 0.58 | 0.092888 |
| 0.59 | 0.0945943 |
| 0.6 | 0.0963043 |
| 0.61 | 0.098018 |
| 0.62 | 0.0997353 |
| 0.63 | 0.101456 |
| 0.64 | 0.103181 |
| 0.65 | 0.10491 |
| 0.66 | 0.106642 |
| 0.67 | 0.108378 |
| 0.68 | 0.110117 |
| 0.69 | 0.111861 |
| 0.7 | 0.113608 |
| 0.71 | 0.115359 |
| 0.72 | 0.117113 |
| 0.73 | 0.118872 |
| 0.74 | 0.120634 |
| 0.75 | 0.1224 |
| 0.76 | 0.12417 |
| 0.77 | 0.125943 |
| 0.78 | 0.127721 |
| 0.79 | 0.129502 |
| 0.8 | 0.131287 |
| 0.81 | 0.133076 |
| 0.82 | 0.134868 |
| 0.83 | 0.136665 |
| 0.84 | 0.138465 |
| 0.85 | 0.140269 |
| 0.86 | 0.142077 |
| 0.87 | 0.143889 |
| 0.88 | 0.145705 |
| 0.89 | 0.147524 |
| 0.9 | 0.149348 |
| 0.91 | 0.151175 |
| 0.92 | 0.153007 |
| 0.93 | 0.154842 |
| 0.94 | 0.156681 |
| 0.95 | 0.158524 |
| 0.96 | 0.160371 |
| 0.97 | 0.162222 |
| 0.98 | 0.164076 |
| 0.99 | 0.165935 |
| 1 | 0.167798 |
| 1.01 | 0.169664 |
| 1.02 | 0.171535 |
| 1.03 | 0.173409 |
| 1.04 | 0.175288 |
| 1.05 | 0.17717 |
| 1.06 | 0.179057 |
| 1.07 | 0.180947 |
| 1.08 | 0.182842 |
| 1.09 | 0.18474 |
| 1.1 | 0.186643 |
| 1.11 | 0.188549 |
| 1.12 | 0.19046 |
| 1.13 | 0.192375 |
| 1.14 | 0.194293 |
| 1.15 | 0.196216 |
| 1.16 | 0.198143 |
| 1.17 | 0.200073 |
| 1.18 | 0.202008 |
| 1.19 | 0.203947 |
| 1.2 | 0.20589 |
| 1.21 | 0.207837 |
| 1.22 | 0.209789 |
| 1.23 | 0.211744 |
| 1.24 | 0.213703 |
| 1.25 | 0.215667 |
| 1.26 | 0.217635 |
| 1.27 | 0.219606 |
| 1.28 | 0.221582 |
| 1.29 | 0.223562 |
| 1.3 | 0.225547 |
| 1.31 | 0.227535 |
| 1.32 | 0.229528 |
| 1.33 | 0.231524 |
| 1.34 | 0.233525 |
| 1.35 | 0.23553 |
| 1.36 | 0.23754 |
| 1.37 | 0.239553 |
| 1.38 | 0.241571 |
| 1.39 | 0.243593 |
| 1.4 | 0.245619 |
| 1.41 | 0.247649 |
| 1.42 | 0.249684 |
| 1.43 | 0.251723 |
| 1.44 | 0.253766 |
| 1.45 | 0.255813 |
| 1.46 | 0.257865 |
| 1.47 | 0.259921 |
| 1.48 | 0.261981 |
| 1.49 | 0.264045 |
| 1.5 | 0.266114 |
| 1.51 | 0.268187 |
| 1.52 | 0.270264 |
| 1.53 | 0.272346 |
| 1.54 | 0.274432 |
| 1.55 | 0.276522 |
| 1.56 | 0.278617 |
| 1.57 | 0.280716 |
| 1.58 | 0.282819 |
| 1.59 | 0.284926 |
| 1.6 | 0.287038 |
| 1.61 | 0.289155 |
| 1.62 | 0.291275 |
| 1.63 | 0.293401 |
| 1.64 | 0.29553 |
| 1.65 | 0.297664 |
| 1.66 | 0.299802 |
| 1.67 | 0.301945 |
| 1.68 | 0.304092 |
| 1.69 | 0.306244 |
| 1.7 | 0.308399 |
| 1.71 | 0.31056 |
| 1.72 | 0.312725 |
| 1.73 | 0.314894 |
| 1.74 | 0.317068 |
| 1.75 | 0.319246 |
| 1.76 | 0.321429 |
| 1.77 | 0.323616 |
| 1.78 | 0.325807 |
| 1.79 | 0.328003 |
| 1.8 | 0.330204 |
| 1.81 | 0.332409 |
| 1.82 | 0.334619 |
| 1.83 | 0.336833 |
| 1.84 | 0.339051 |
| 1.85 | 0.341275 |
| 1.86 | 0.343502 |
| 1.87 | 0.345735 |
| 1.88 | 0.347971 |
| 1.89 | 0.350213 |
| 1.9 | 0.352459 |
| 1.91 | 0.354709 |
| 1.92 | 0.356964 |
| 1.93 | 0.359224 |
| 1.94 | 0.361488 |
| 1.95 | 0.363757 |
| 1.96 | 0.366031 |
| 1.97 | 0.368309 |
| 1.98 | 0.370592 |
| 1.99 | 0.372879 |

H1 Total fitness

|  |  |
| --- | --- |
| 1 | 0.167798 |
| 2 | 0.375171 |
| 3 | 0.628954 |
| 4 | 0.935837 |
| 5 | 1.3016 |
| 6 | 1.73009 |
| 7 | 2.22206 |
| 8 | 2.77403 |
| 9 | 3.37748 |
| 10 | 4.01885 |
| 11 | 4.68036 |
| 12 | 5.34187 |
| 13 | 5.98324 |
| 14 | 6.58669 |
| 15 | 7.13866 |
| 16 | 7.63063 |
| 17 | 8.05912 |
| 18 | 8.42488 |
| 19 | 8.73177 |
| 20 | 8.98555 |
| 21 | 9.19292 |
| 22 | 9.36072 |
| 23 | 10.0248 |
| 24 | 10.6837 |
| 25 | 11.3325 |
| 26 | 11.9665 |
| 27 | 12.5813 |
| 28 | 13.1734 |
| 29 | 13.7395 |
| 30 | 14.2773 |
| 31 | 14.7849 |
| 32 | 15.2613 |
| 33 | 15.7057 |
| 34 | 16.1182 |
| 35 | 16.4993 |
| 36 | 16.8498 |
| 37 | 17.1707 |
| 38 | 17.4635 |
| 39 | 17.9497 |
| 40 | 18.5681 |
| 41 | 19.1721 |
| 42 | 19.7601 |
| 43 | 20.3308 |
| 44 | 20.8831 |
| 45 | 21.416 |
| 46 | 21.9288 |
| 47 | 22.4209 |
| 48 | 22.8919 |
| 49 | 23.3417 |
| 50 | 23.7701 |
| 51 | 24.1774 |
| 52 | 24.5636 |
| 53 | 24.9292 |
| 54 | 25.2747 |
| 55 | 25.7608 |
| 56 | 26.3468 |
| 57 | 26.9197 |
| 58 | 27.4791 |
| 59 | 28.0242 |
| 60 | 28.5546 |
| 61 | 29.07 |
| 62 | 29.5699 |
| 63 | 30.0541 |
| 64 | 30.5225 |
| 65 | 30.975 |
| 66 | 31.4114 |
| 67 | 31.8319 |
| 68 | 32.2365 |
| 69 | 32.6254 |
| 70 | 32.9987 |
| 71 | 33.5078 |
| 72 | 34.0713 |
| 73 | 34.6236 |
| 74 | 35.1643 |
| 75 | 35.6933 |
| 76 | 36.2102 |
| 77 | 36.7147 |
| 78 | 37.2068 |
| 79 | 37.6864 |
| 80 | 38.1532 |
| 81 | 38.6072 |
| 82 | 39.0484 |
| 83 | 39.4769 |
| 84 | 39.8926 |
| 85 | 40.2956 |
| 86 | 40.686 |
| 87 | 41.2186 |
| 88 | 41.7661 |
| 89 | 42.304 |
| 90 | 42.8319 |
| 91 | 43.3498 |
| 92 | 43.8575 |
| 93 | 44.3548 |
| 94 | 44.8417 |
| 95 | 45.318 |
| 96 | 45.7838 |
| 97 | 46.2389 |
| 98 | 46.6833 |
| 99 | 47.1171 |
| 100 | 47.5403 |
| 101 | 47.9528 |
| 102 | 48.3635 |
| 103 | 48.9076 |
| 104 | 49.4434 |
| 105 | 49.9706 |
| 106 | 50.4892 |
| 107 | 50.9991 |
| 108 | 51.5001 |
| 109 | 51.9922 |
| 110 | 52.4754 |
| 111 | 52.9494 |
| 112 | 53.4144 |
| 113 | 53.8703 |
| 114 | 54.317 |
| 115 | 54.7546 |
| 116 | 55.1831 |
| 117 | 55.6024 |
| 118 | 56.0484 |
| 119 | 56.5825 |
| 120 | 57.1093 |
| 121 | 57.6284 |
| 122 | 58.1399 |
| 123 | 58.6438 |
| 124 | 59.1398 |
| 125 | 59.628 |
| 126 | 60.1083 |
| 127 | 60.5806 |
| 128 | 61.045 |
| 129 | 61.5015 |
| 130 | 61.9499 |
| 131 | 62.3904 |
| 132 | 62.8228 |
| 133 | 63.2473 |
| 134 | 63.7216 |
| 135 | 64.2479 |
| 136 | 64.7675 |
| 137 | 65.2803 |
| 138 | 65.7863 |
| 139 | 66.2853 |
| 140 | 66.7774 |
| 141 | 67.2626 |
| 142 | 67.7407 |
| 143 | 68.2117 |
| 144 | 68.6757 |
| 145 | 69.1326 |
| 146 | 69.5823 |
| 147 | 70.025 |
| 148 | 70.4606 |
| 149 | 70.889 |
| 150 | 71.3866 |
| 151 | 71.9065 |
| 152 | 72.4203 |
| 153 | 72.928 |
| 154 | 73.4295 |
| 155 | 73.9247 |
| 156 | 74.4137 |
| 157 | 74.8964 |
| 158 | 75.3727 |
| 159 | 75.8427 |
| 160 | 76.3063 |
| 161 | 76.7635 |
| 162 | 77.2144 |
| 163 | 77.6588 |
| 164 | 78.0969 |
| 165 | 78.5285 |
| 166 | 78.9538 |
| 167 | 79.3727 |
| 168 | 79.7852 |
| 169 | 80.1914 |
| 170 | 80.5912 |
| 171 | 80.9848 |
| 172 | 81.3721 |
| 173 | 81.7531 |
| 174 | 82.128 |
| 175 | 82.4966 |
| 176 | 82.8592 |
| 177 | 83.2156 |
| 178 | 83.566 |
| 179 | 83.9104 |
| 180 | 84.2488 |
| 181 | 84.5814 |
| 182 | 84.9081 |
| 183 | 85.229 |
| 184 | 85.5441 |
| 185 | 85.8536 |
| 186 | 86.1574 |
| 187 | 86.4557 |
| 188 | 86.7484 |
| 189 | 87.0357 |
| 190 | 87.3177 |
| 191 | 87.5943 |
| 192 | 87.8656 |
| 193 | 88.1318 |
| 194 | 88.3929 |
| 195 | 88.6489 |
| 196 | 88.8999 |
| 197 | 89.146 |
| 198 | 89.3873 |
| 199 | 89.6237 |
| 200 | 89.8555 |

H1 Numbers

|  |  |
| --- | --- |
| 1 | 1 |
| 2 | 1 |
| 3 | 1 |
| 4 | 1 |
| 5 | 1 |
| 6 | 1 |
| 7 | 1 |
| 8 | 1 |
| 9 | 1 |
| 10 | 1 |
| 11 | 1 |
| 12 | 1 |
| 13 | 1 |
| 14 | 1 |
| 15 | 1 |
| 16 | 1 |
| 17 | 1 |
| 18 | 1 |
| 19 | 1 |
| 20 | 1 |
| 21 | 1 |
| 22 | 1 |
| 23 | 2 |
| 24 | 2 |
| 25 | 2 |
| 26 | 2 |
| 27 | 2 |
| 28 | 2 |
| 29 | 2 |
| 30 | 2 |
| 31 | 2 |
| 32 | 2 |
| 33 | 2 |
| 34 | 2 |
| 35 | 2 |
| 36 | 2 |
| 37 | 2 |
| 38 | 2 |
| 39 | 3 |
| 40 | 3 |
| 41 | 3 |
| 42 | 3 |
| 43 | 3 |
| 44 | 3 |
| 45 | 3 |
| 46 | 3 |
| 47 | 3 |
| 48 | 3 |
| 49 | 3 |
| 50 | 3 |
| 51 | 3 |
| 52 | 3 |
| 53 | 3 |
| 54 | 3 |
| 55 | 4 |
| 56 | 4 |
| 57 | 4 |
| 58 | 4 |
| 59 | 4 |
| 60 | 4 |
| 61 | 4 |
| 62 | 4 |
| 63 | 4 |
| 64 | 4 |
| 65 | 4 |
| 66 | 4 |
| 67 | 4 |
| 68 | 4 |
| 69 | 4 |
| 70 | 4 |
| 71 | 5 |
| 72 | 5 |
| 73 | 5 |
| 74 | 5 |
| 75 | 5 |
| 76 | 5 |
| 77 | 5 |
| 78 | 5 |
| 79 | 5 |
| 80 | 5 |
| 81 | 5 |
| 82 | 5 |
| 83 | 5 |
| 84 | 5 |
| 85 | 5 |
| 86 | 5 |
| 87 | 6 |
| 88 | 6 |
| 89 | 6 |
| 90 | 6 |
| 91 | 6 |
| 92 | 6 |
| 93 | 6 |
| 94 | 6 |
| 95 | 6 |
| 96 | 6 |
| 97 | 6 |
| 98 | 6 |
| 99 | 6 |
| 100 | 6 |
| 101 | 6 |
| 102 | 7 |
| 103 | 7 |
| 104 | 7 |
| 105 | 7 |
| 106 | 7 |
| 107 | 7 |
| 108 | 7 |
| 109 | 7 |
| 110 | 7 |
| 111 | 7 |
| 112 | 7 |
| 113 | 7 |
| 114 | 7 |
| 115 | 7 |
| 116 | 7 |
| 117 | 7 |
| 118 | 8 |
| 119 | 8 |
| 120 | 8 |
| 121 | 8 |
| 122 | 8 |
| 123 | 8 |
| 124 | 8 |
| 125 | 8 |
| 126 | 8 |
| 127 | 8 |
| 128 | 8 |
| 129 | 8 |
| 130 | 8 |
| 131 | 8 |
| 132 | 8 |
| 133 | 8 |
| 134 | 9 |
| 135 | 9 |
| 136 | 9 |
| 137 | 9 |
| 138 | 9 |
| 139 | 9 |
| 140 | 9 |
| 141 | 9 |
| 142 | 9 |
| 143 | 9 |
| 144 | 9 |
| 145 | 9 |
| 146 | 9 |
| 147 | 9 |
| 148 | 9 |
| 149 | 9 |
| 150 | 10 |
| 151 | 10 |
| 152 | 10 |
| 153 | 10 |
| 154 | 10 |
| 155 | 10 |
| 156 | 10 |
| 157 | 10 |
| 158 | 10 |
| 159 | 10 |
| 160 | 10 |
| 161 | 10 |
| 162 | 10 |
| 163 | 10 |
| 164 | 10 |
| 165 | 10 |
| 166 | 10 |
| 167 | 10 |
| 168 | 10 |
| 169 | 10 |
| 170 | 10 |
| 171 | 10 |
| 172 | 10 |
| 173 | 10 |
| 174 | 10 |
| 175 | 10 |
| 176 | 10 |
| 177 | 10 |
| 178 | 10 |
| 179 | 10 |
| 180 | 10 |
| 181 | 10 |
| 182 | 10 |
| 183 | 10 |
| 184 | 10 |
| 185 | 10 |
| 186 | 10 |
| 187 | 10 |
| 188 | 10 |
| 189 | 10 |
| 190 | 10 |
| 191 | 10 |
| 192 | 10 |
| 193 | 10 |
| 194 | 10 |
| 195 | 10 |
| 196 | 10 |
| 197 | 10 |
| 198 | 10 |
| 199 | 10 |
| 200 | 10 |

H1 Per capita

|  |  |
| --- | --- |
| 1 | 1 |
| 2 | 2 |
| 3 | 3 |
| 4 | 4 |
| 5 | 5 |
| 6 | 6 |
| 7 | 7 |
| 8 | 8 |
| 9 | 9 |
| 10 | 10 |
| 11 | 11 |
| 12 | 12 |
| 13 | 13 |
| 14 | 14 |
| 15 | 15 |
| 16 | 16 |
| 17 | 17 |
| 18 | 18 |
| 19 | 19 |
| 20 | 20 |
| 21 | 21 |
| 22 | 22 |
| 23 | 11.5 |
| 24 | 12 |
| 25 | 12.5 |
| 26 | 13 |
| 27 | 13.5 |
| 28 | 14 |
| 29 | 14.5 |
| 30 | 15 |
| 31 | 15.5 |
| 32 | 16 |
| 33 | 16.5 |
| 34 | 17 |
| 35 | 17.5 |
| 36 | 18 |
| 37 | 18.5 |
| 38 | 19 |
| 39 | 13 |
| 40 | 13.3333 |
| 41 | 13.6667 |
| 42 | 14 |
| 43 | 14.3333 |
| 44 | 14.6667 |
| 45 | 15 |
| 46 | 15.3333 |
| 47 | 15.6667 |
| 48 | 16 |
| 49 | 16.3333 |
| 50 | 16.6667 |
| 51 | 17 |
| 52 | 17.3333 |
| 53 | 17.6667 |
| 54 | 18 |
| 55 | 13.75 |
| 56 | 14 |
| 57 | 14.25 |
| 58 | 14.5 |
| 59 | 14.75 |
| 60 | 15 |
| 61 | 15.25 |
| 62 | 15.5 |
| 63 | 15.75 |
| 64 | 16 |
| 65 | 16.25 |
| 66 | 16.5 |
| 67 | 16.75 |
| 68 | 17 |
| 69 | 17.25 |
| 70 | 17.5 |
| 71 | 14.2 |
| 72 | 14.4 |
| 73 | 14.6 |
| 74 | 14.8 |
| 75 | 15 |
| 76 | 15.2 |
| 77 | 15.4 |
| 78 | 15.6 |
| 79 | 15.8 |
| 80 | 16 |
| 81 | 16.2 |
| 82 | 16.4 |
| 83 | 16.6 |
| 84 | 16.8 |
| 85 | 17 |
| 86 | 17.2 |
| 87 | 14.5 |
| 88 | 14.6667 |
| 89 | 14.8333 |
| 90 | 15 |
| 91 | 15.1667 |
| 92 | 15.3333 |
| 93 | 15.5 |
| 94 | 15.6667 |
| 95 | 15.8333 |
| 96 | 16 |
| 97 | 16.1667 |
| 98 | 16.3333 |
| 99 | 16.5 |
| 100 | 16.6667 |
| 101 | 16.8333 |
| 102 | 14.5714 |
| 103 | 14.7143 |
| 104 | 14.8571 |
| 105 | 15 |
| 106 | 15.1429 |
| 107 | 15.2857 |
| 108 | 15.4286 |
| 109 | 15.5714 |
| 110 | 15.7143 |
| 111 | 15.8571 |
| 112 | 16 |
| 113 | 16.1429 |
| 114 | 16.2857 |
| 115 | 16.4286 |
| 116 | 16.5714 |
| 117 | 16.7143 |
| 118 | 14.75 |
| 119 | 14.875 |
| 120 | 15 |
| 121 | 15.125 |
| 122 | 15.25 |
| 123 | 15.375 |
| 124 | 15.5 |
| 125 | 15.625 |
| 126 | 15.75 |
| 127 | 15.875 |
| 128 | 16 |
| 129 | 16.125 |
| 130 | 16.25 |
| 131 | 16.375 |
| 132 | 16.5 |
| 133 | 16.625 |
| 134 | 14.8889 |
| 135 | 15 |
| 136 | 15.1111 |
| 137 | 15.2222 |
| 138 | 15.3333 |
| 139 | 15.4444 |
| 140 | 15.5556 |
| 141 | 15.6667 |
| 142 | 15.7778 |
| 143 | 15.8889 |
| 144 | 16 |
| 145 | 16.1111 |
| 146 | 16.2222 |
| 147 | 16.3333 |
| 148 | 16.4444 |
| 149 | 16.5556 |
| 150 | 15 |
| 151 | 15.1 |
| 152 | 15.2 |
| 153 | 15.3 |
| 154 | 15.4 |
| 155 | 15.5 |
| 156 | 15.6 |
| 157 | 15.7 |
| 158 | 15.8 |
| 159 | 15.9 |
| 160 | 16 |
| 161 | 16.1 |
| 162 | 16.2 |
| 163 | 16.3 |
| 164 | 16.4 |
| 165 | 16.5 |
| 166 | 16.6 |
| 167 | 16.7 |
| 168 | 16.8 |
| 169 | 16.9 |
| 170 | 17 |
| 171 | 17.1 |
| 172 | 17.2 |
| 173 | 17.3 |
| 174 | 17.4 |
| 175 | 17.5 |
| 176 | 17.6 |
| 177 | 17.7 |
| 178 | 17.8 |
| 179 | 17.9 |
| 180 | 18 |
| 181 | 18.1 |
| 182 | 18.2 |
| 183 | 18.3 |
| 184 | 18.4 |
| 185 | 18.5 |
| 186 | 18.6 |
| 187 | 18.7 |
| 188 | 18.8 |
| 189 | 18.9 |
| 190 | 19 |
| 191 | 19.1 |
| 192 | 19.2 |
| 193 | 19.3 |
| 194 | 19.4 |
| 195 | 19.5 |
| 196 | 19.6 |
| 197 | 19.7 |
| 198 | 19.8 |
| 199 | 19.9 |
| 200 | 20 |

Figure S3.

(a) The logistic fitness function of model H1. In this model, every offspring has an identical fitness function.

(b) The optimized sum of total offspring fitness.

(c) The optimized number of the cared offspring.

(d) The Per Capita investment of each cared offspring.

**Model Version H2**

H2 Function

|  |  |
| --- | --- |
| 0 | -1.6263e-18 |
| 0.01 | 4.56256e-06 |
| 0.02 | 9.17097e-06 |
| 0.03 | 1.38257e-05 |
| 0.04 | 1.85272e-05 |
| 0.05 | 2.32759e-05 |
| 0.06 | 2.80724e-05 |
| 0.07 | 3.29171e-05 |
| 0.08 | 3.78104e-05 |
| 0.09 | 4.27529e-05 |
| 0.1 | 4.77451e-05 |
| 0.11 | 5.27875e-05 |
| 0.12 | 5.78805e-05 |
| 0.13 | 6.30247e-05 |
| 0.14 | 6.82206e-05 |
| 0.15 | 7.34688e-05 |
| 0.16 | 7.87696e-05 |
| 0.17 | 8.41237e-05 |
| 0.18 | 8.95317e-05 |
| 0.19 | 9.4994e-05 |
| 0.2 | 0.000100511 |
| 0.21 | 0.000106084 |
| 0.22 | 0.000111712 |
| 0.23 | 0.000117398 |
| 0.24 | 0.00012314 |
| 0.25 | 0.00012894 |
| 0.26 | 0.000134798 |
| 0.27 | 0.000140715 |
| 0.28 | 0.000146692 |
| 0.29 | 0.000152729 |
| 0.3 | 0.000158826 |
| 0.31 | 0.000164985 |
| 0.32 | 0.000171205 |
| 0.33 | 0.000177488 |
| 0.34 | 0.000183834 |
| 0.35 | 0.000190244 |
| 0.36 | 0.000196718 |
| 0.37 | 0.000203258 |
| 0.38 | 0.000209863 |
| 0.39 | 0.000216534 |
| 0.4 | 0.000223273 |
| 0.41 | 0.000230079 |
| 0.42 | 0.000236954 |
| 0.43 | 0.000243898 |
| 0.44 | 0.000250911 |
| 0.45 | 0.000257995 |
| 0.46 | 0.00026515 |
| 0.47 | 0.000272377 |
| 0.48 | 0.000279677 |
| 0.49 | 0.00028705 |
| 0.5 | 0.000294497 |
| 0.51 | 0.000302019 |
| 0.52 | 0.000309616 |
| 0.53 | 0.00031729 |
| 0.54 | 0.000325041 |
| 0.55 | 0.00033287 |
| 0.56 | 0.000340778 |
| 0.57 | 0.000348765 |
| 0.58 | 0.000356832 |
| 0.59 | 0.00036498 |
| 0.6 | 0.00037321 |
| 0.61 | 0.000381523 |
| 0.62 | 0.00038992 |
| 0.63 | 0.000398401 |
| 0.64 | 0.000406967 |
| 0.65 | 0.000415619 |
| 0.66 | 0.000424358 |
| 0.67 | 0.000433185 |
| 0.68 | 0.0004421 |
| 0.69 | 0.000451105 |
| 0.7 | 0.000460201 |
| 0.71 | 0.000469388 |
| 0.72 | 0.000478667 |
| 0.73 | 0.00048804 |
| 0.74 | 0.000497507 |
| 0.75 | 0.000507068 |
| 0.76 | 0.000516726 |
| 0.77 | 0.000526481 |
| 0.78 | 0.000536334 |
| 0.79 | 0.000546286 |
| 0.8 | 0.000556339 |
| 0.81 | 0.000566492 |
| 0.82 | 0.000576747 |
| 0.83 | 0.000587105 |
| 0.84 | 0.000597567 |
| 0.85 | 0.000608134 |
| 0.86 | 0.000618808 |
| 0.87 | 0.000629588 |
| 0.88 | 0.000640477 |
| 0.89 | 0.000651476 |
| 0.9 | 0.000662585 |
| 0.91 | 0.000673805 |
| 0.92 | 0.000685139 |
| 0.93 | 0.000696586 |
| 0.94 | 0.000708148 |
| 0.95 | 0.000719827 |
| 0.96 | 0.000731622 |
| 0.97 | 0.000743537 |
| 0.98 | 0.000755571 |
| 0.99 | 0.000767725 |
| 1 | 0.000780002 |
| 1.01 | 0.000792403 |
| 1.02 | 0.000804928 |
| 1.03 | 0.000817579 |
| 1.04 | 0.000830357 |
| 1.05 | 0.000843263 |
| 1.06 | 0.000856299 |
| 1.07 | 0.000869466 |
| 1.08 | 0.000882765 |
| 1.09 | 0.000896198 |
| 1.1 | 0.000909766 |
| 1.11 | 0.00092347 |
| 1.12 | 0.000937312 |
| 1.13 | 0.000951293 |
| 1.14 | 0.000965414 |
| 1.15 | 0.000979678 |
| 1.16 | 0.000994084 |
| 1.17 | 0.00100864 |
| 1.18 | 0.00102333 |
| 1.19 | 0.00103818 |
| 1.2 | 0.00105317 |
| 1.21 | 0.00106832 |
| 1.22 | 0.00108361 |
| 1.23 | 0.00109907 |
| 1.24 | 0.00111467 |
| 1.25 | 0.00113043 |
| 1.26 | 0.00114636 |
| 1.27 | 0.00116244 |
| 1.28 | 0.00117868 |
| 1.29 | 0.00119509 |
| 1.3 | 0.00121166 |
| 1.31 | 0.00122839 |
| 1.32 | 0.0012453 |
| 1.33 | 0.00126238 |
| 1.34 | 0.00127962 |
| 1.35 | 0.00129704 |
| 1.36 | 0.00131464 |
| 1.37 | 0.00133241 |
| 1.38 | 0.00135036 |
| 1.39 | 0.00136849 |
| 1.4 | 0.0013868 |
| 1.41 | 0.0014053 |
| 1.42 | 0.00142398 |
| 1.43 | 0.00144285 |
| 1.44 | 0.00146191 |
| 1.45 | 0.00148117 |
| 1.46 | 0.00150061 |
| 1.47 | 0.00152025 |
| 1.48 | 0.00154009 |
| 1.49 | 0.00156012 |
| 1.5 | 0.00158036 |
| 1.51 | 0.0016008 |
| 1.52 | 0.00162145 |
| 1.53 | 0.00164231 |
| 1.54 | 0.00166337 |
| 1.55 | 0.00168464 |
| 1.56 | 0.00170613 |
| 1.57 | 0.00172784 |
| 1.58 | 0.00174976 |
| 1.59 | 0.0017719 |
| 1.6 | 0.00179427 |
| 1.61 | 0.00181686 |
| 1.62 | 0.00183968 |
| 1.63 | 0.00186272 |
| 1.64 | 0.001886 |
| 1.65 | 0.00190951 |
| 1.66 | 0.00193326 |
| 1.67 | 0.00195725 |
| 1.68 | 0.00198148 |
| 1.69 | 0.00200595 |
| 1.7 | 0.00203066 |
| 1.71 | 0.00205563 |
| 1.72 | 0.00208085 |
| 1.73 | 0.00210631 |
| 1.74 | 0.00213204 |
| 1.75 | 0.00215802 |
| 1.76 | 0.00218427 |
| 1.77 | 0.00221077 |
| 1.78 | 0.00223755 |
| 1.79 | 0.00226459 |
| 1.8 | 0.00229191 |
| 1.81 | 0.0023195 |
| 1.82 | 0.00234736 |
| 1.83 | 0.00237551 |
| 1.84 | 0.00240394 |
| 1.85 | 0.00243265 |
| 1.86 | 0.00246165 |
| 1.87 | 0.00249095 |
| 1.88 | 0.00252054 |
| 1.89 | 0.00255042 |
| 1.9 | 0.00258061 |
| 1.91 | 0.0026111 |
| 1.92 | 0.00264189 |
| 1.93 | 0.002673 |
| 1.94 | 0.00270441 |
| 1.95 | 0.00273615 |
| 1.96 | 0.0027682 |
| 1.97 | 0.00280057 |
| 1.98 | 0.00283327 |
| 1.99 | 0.0028663 |

H2 Total fitness

|  |  |
| --- | --- |
| 1 | 0.000780002 |
| 2 | 0.00289965 |
| 3 | 0.00865693 |
| 4 | 0.0242734 |
| 5 | 0.0664775 |
| 6 | 0.179416 |
| 7 | 0.473826 |
| 8 | 1.19163 |
| 9 | 2.68908 |
| 10 | 4.99977 |
| 11 | 7.31046 |
| 12 | 8.80792 |
| 13 | 9.52572 |
| 14 | 9.82013 |
| 15 | 9.93307 |
| 16 | 9.97527 |
| 17 | 9.99089 |
| 18 | 9.99665 |
| 19 | 9.99877 |
| 20 | 9.99955 |
| 21 | 12.4488 |
| 22 | 14.6209 |
| 23 | 16.3513 |
| 24 | 17.6158 |
| 25 | 18.4828 |
| 26 | 19.0514 |
| 27 | 19.4137 |
| 28 | 19.6403 |
| 29 | 19.7803 |
| 30 | 19.8661 |
| 31 | 19.9186 |
| 32 | 19.9505 |
| 33 | 21.9314 |
| 34 | 23.7415 |
| 35 | 25.2337 |
| 36 | 26.4237 |
| 37 | 27.3479 |
| 38 | 28.0508 |
| 39 | 28.5772 |
| 40 | 28.9666 |
| 41 | 29.2522 |
| 42 | 29.4604 |
| 43 | 29.6114 |
| 44 | 29.7205 |
| 45 | 31.0916 |
| 46 | 32.7026 |
| 47 | 34.0778 |
| 48 | 35.2317 |
| 49 | 36.1858 |
| 50 | 36.9655 |
| 51 | 37.5964 |
| 52 | 38.1029 |
| 53 | 38.5069 |
| 54 | 38.8275 |
| 55 | 39.0809 |
| 56 | 39.2805 |
| 57 | 40.1087 |
| 58 | 41.6005 |
| 59 | 42.9071 |
| 60 | 44.0396 |
| 61 | 45.0122 |
| 62 | 45.8412 |
| 63 | 46.5429 |
| 64 | 47.1337 |
| 65 | 47.6286 |
| 66 | 48.0416 |
| 67 | 48.3852 |
| 68 | 48.6701 |
| 69 | 49.054 |
| 70 | 50.4674 |
| 71 | 51.7291 |
| 72 | 52.8475 |
| 73 | 53.8327 |
| 74 | 54.6958 |
| 75 | 55.4483 |
| 76 | 56.1017 |
| 77 | 56.6669 |
| 78 | 57.1543 |
| 79 | 57.5735 |
| 80 | 57.9332 |
| 81 | 58.2412 |
| 82 | 59.3169 |
| 83 | 60.547 |
| 84 | 61.6554 |
| 85 | 62.6496 |
| 86 | 63.5378 |
| 87 | 64.3284 |
| 88 | 65.0298 |
| 89 | 65.6503 |
| 90 | 66.1979 |
| 91 | 66.68 |
| 92 | 67.1037 |
| 93 | 67.4754 |
| 94 | 68.1557 |
| 95 | 69.3624 |
| 96 | 70.4633 |
| 97 | 71.4644 |
| 98 | 72.3717 |
| 99 | 73.1918 |
| 100 | 73.9311 |
| 101 | 74.596 |
| 102 | 75.1928 |
| 103 | 75.7275 |
| 104 | 76.2058 |
| 105 | 76.6328 |
| 106 | 77.0137 |
| 107 | 78.1761 |
| 108 | 79.2712 |
| 109 | 80.2776 |
| 110 | 81.2001 |
| 111 | 82.0437 |
| 112 | 82.8136 |
| 113 | 83.5148 |
| 114 | 84.1525 |
| 115 | 84.7314 |
| 116 | 85.2563 |
| 117 | 85.7315 |
| 118 | 86.1612 |
| 119 | 86.9886 |
| 120 | 88.0792 |
| 121 | 89.0898 |
| 122 | 90.0245 |
| 123 | 90.8873 |
| 124 | 91.6824 |
| 125 | 92.4138 |
| 126 | 93.0858 |
| 127 | 93.7024 |
| 128 | 94.2673 |
| 129 | 94.7844 |
| 130 | 95.2572 |
| 131 | 95.6891 |
| 132 | 96.0832 |
| 133 | 96.4427 |
| 134 | 96.7703 |
| 135 | 97.0686 |
| 136 | 97.3402 |
| 137 | 97.5872 |
| 138 | 97.8118 |
| 139 | 98.0159 |
| 140 | 98.2013 |
| 141 | 98.3697 |
| 142 | 98.5225 |
| 143 | 98.6612 |
| 144 | 98.7871 |
| 145 | 98.9013 |
| 146 | 99.0048 |
| 147 | 99.0986 |
| 148 | 99.1837 |
| 149 | 99.2608 |
| 150 | 99.3307 |
| 151 | 99.394 |
| 152 | 99.4513 |
| 153 | 99.5033 |
| 154 | 99.5504 |
| 155 | 99.593 |
| 156 | 99.6316 |
| 157 | 99.6665 |
| 158 | 99.6981 |
| 159 | 99.7268 |
| 160 | 99.7527 |
| 161 | 99.7762 |
| 162 | 99.7975 |
| 163 | 99.8167 |
| 164 | 99.8341 |
| 165 | 99.8499 |
| 166 | 99.8641 |
| 167 | 99.8771 |
| 168 | 99.8887 |
| 169 | 99.8993 |
| 170 | 99.9089 |
| 171 | 99.9176 |
| 172 | 99.9254 |
| 173 | 99.9325 |
| 174 | 99.9389 |
| 175 | 99.9447 |
| 176 | 99.95 |
| 177 | 99.9547 |
| 178 | 99.959 |
| 179 | 99.9629 |
| 180 | 99.9665 |
| 181 | 99.9697 |
| 182 | 99.9725 |
| 183 | 99.9752 |
| 184 | 99.9775 |
| 185 | 99.9797 |
| 186 | 99.9816 |
| 187 | 99.9833 |
| 188 | 99.9849 |
| 189 | 99.9864 |
| 190 | 99.9877 |
| 191 | 99.9888 |
| 192 | 99.9899 |
| 193 | 99.9909 |
| 194 | 99.9917 |
| 195 | 99.9925 |
| 196 | 99.9932 |
| 197 | 99.9939 |
| 198 | 99.9945 |
| 199 | 99.995 |
| 200 | 99.9955 |

H2 Numbers

|  |  |
| --- | --- |
| 1 | 1 |
| 2 | 1 |
| 3 | 1 |
| 4 | 1 |
| 5 | 1 |
| 6 | 1 |
| 7 | 1 |
| 8 | 1 |
| 9 | 1 |
| 10 | 1 |
| 11 | 1 |
| 12 | 1 |
| 13 | 1 |
| 14 | 1 |
| 15 | 1 |
| 16 | 1 |
| 17 | 1 |
| 18 | 1 |
| 19 | 1 |
| 20 | 1 |
| 21 | 2 |
| 22 | 2 |
| 23 | 2 |
| 24 | 2 |
| 25 | 2 |
| 26 | 2 |
| 27 | 2 |
| 28 | 2 |
| 29 | 2 |
| 30 | 2 |
| 31 | 2 |
| 32 | 2 |
| 33 | 3 |
| 34 | 3 |
| 35 | 3 |
| 36 | 3 |
| 37 | 3 |
| 38 | 3 |
| 39 | 3 |
| 40 | 3 |
| 41 | 3 |
| 42 | 3 |
| 43 | 3 |
| 44 | 3 |
| 45 | 4 |
| 46 | 4 |
| 47 | 4 |
| 48 | 4 |
| 49 | 4 |
| 50 | 4 |
| 51 | 4 |
| 52 | 4 |
| 53 | 4 |
| 54 | 4 |
| 55 | 4 |
| 56 | 4 |
| 57 | 5 |
| 58 | 5 |
| 59 | 5 |
| 60 | 5 |
| 61 | 5 |
| 62 | 5 |
| 63 | 5 |
| 64 | 5 |
| 65 | 5 |
| 66 | 5 |
| 67 | 5 |
| 68 | 5 |
| 69 | 6 |
| 70 | 6 |
| 71 | 6 |
| 72 | 6 |
| 73 | 6 |
| 74 | 6 |
| 75 | 6 |
| 76 | 6 |
| 77 | 6 |
| 78 | 6 |
| 79 | 6 |
| 80 | 6 |
| 81 | 6 |
| 82 | 7 |
| 83 | 7 |
| 84 | 7 |
| 85 | 7 |
| 86 | 7 |
| 87 | 7 |
| 88 | 7 |
| 89 | 7 |
| 90 | 7 |
| 91 | 7 |
| 92 | 7 |
| 93 | 7 |
| 94 | 8 |
| 95 | 8 |
| 96 | 8 |
| 97 | 8 |
| 98 | 8 |
| 99 | 8 |
| 100 | 8 |
| 101 | 8 |
| 102 | 8 |
| 103 | 8 |
| 104 | 8 |
| 105 | 8 |
| 106 | 8 |
| 107 | 9 |
| 108 | 9 |
| 109 | 9 |
| 110 | 9 |
| 111 | 9 |
| 112 | 9 |
| 113 | 9 |
| 114 | 9 |
| 115 | 9 |
| 116 | 9 |
| 117 | 9 |
| 118 | 9 |
| 119 | 10 |
| 120 | 10 |
| 121 | 10 |
| 122 | 10 |
| 123 | 10 |
| 124 | 10 |
| 125 | 10 |
| 126 | 10 |
| 127 | 10 |
| 128 | 10 |
| 129 | 10 |
| 130 | 10 |
| 131 | 10 |
| 132 | 10 |
| 133 | 10 |
| 134 | 10 |
| 135 | 10 |
| 136 | 10 |
| 137 | 10 |
| 138 | 10 |
| 139 | 10 |
| 140 | 10 |
| 141 | 10 |
| 142 | 10 |
| 143 | 10 |
| 144 | 10 |
| 145 | 10 |
| 146 | 10 |
| 147 | 10 |
| 148 | 10 |
| 149 | 10 |
| 150 | 10 |
| 151 | 10 |
| 152 | 10 |
| 153 | 10 |
| 154 | 10 |
| 155 | 10 |
| 156 | 10 |
| 157 | 10 |
| 158 | 10 |
| 159 | 10 |
| 160 | 10 |
| 161 | 10 |
| 162 | 10 |
| 163 | 10 |
| 164 | 10 |
| 165 | 10 |
| 166 | 10 |
| 167 | 10 |
| 168 | 10 |
| 169 | 10 |
| 170 | 10 |
| 171 | 10 |
| 172 | 10 |
| 173 | 10 |
| 174 | 10 |
| 175 | 10 |
| 176 | 10 |
| 177 | 10 |
| 178 | 10 |
| 179 | 10 |
| 180 | 10 |
| 181 | 10 |
| 182 | 10 |
| 183 | 10 |
| 184 | 10 |
| 185 | 10 |
| 186 | 10 |
| 187 | 10 |
| 188 | 10 |
| 189 | 10 |
| 190 | 10 |
| 191 | 10 |
| 192 | 10 |
| 193 | 10 |
| 194 | 10 |
| 195 | 10 |
| 196 | 10 |
| 197 | 10 |
| 198 | 10 |
| 199 | 10 |
| 200 | 10 |

H2 Per capita

|  |  |
| --- | --- |
| 1 | 1 |
| 2 | 2 |
| 3 | 3 |
| 4 | 4 |
| 5 | 5 |
| 6 | 6 |
| 7 | 7 |
| 8 | 8 |
| 9 | 9 |
| 10 | 10 |
| 11 | 11 |
| 12 | 12 |
| 13 | 13 |
| 14 | 14 |
| 15 | 15 |
| 16 | 16 |
| 17 | 17 |
| 18 | 18 |
| 19 | 19 |
| 20 | 19.9999 |
| 21 | 10.5 |
| 22 | 11 |
| 23 | 11.5 |
| 24 | 12 |
| 25 | 12.5 |
| 26 | 13 |
| 27 | 13.5 |
| 28 | 14 |
| 29 | 14.5 |
| 30 | 15 |
| 31 | 15.5 |
| 32 | 16 |
| 33 | 11 |
| 34 | 11.3333 |
| 35 | 11.6667 |
| 36 | 12 |
| 37 | 12.3333 |
| 38 | 12.6667 |
| 39 | 13 |
| 40 | 13.3333 |
| 41 | 13.6667 |
| 42 | 14 |
| 43 | 14.3333 |
| 44 | 14.6667 |
| 45 | 11.25 |
| 46 | 11.5 |
| 47 | 11.75 |
| 48 | 12 |
| 49 | 12.25 |
| 50 | 12.5 |
| 51 | 12.75 |
| 52 | 13 |
| 53 | 13.25 |
| 54 | 13.5 |
| 55 | 13.75 |
| 56 | 14 |
| 57 | 11.4 |
| 58 | 11.6 |
| 59 | 11.8 |
| 60 | 12 |
| 61 | 12.2 |
| 62 | 12.4 |
| 63 | 12.6 |
| 64 | 12.8 |
| 65 | 13 |
| 66 | 13.2 |
| 67 | 13.4 |
| 68 | 13.6 |
| 69 | 11.5 |
| 70 | 11.6667 |
| 71 | 11.8333 |
| 72 | 12 |
| 73 | 12.1667 |
| 74 | 12.3333 |
| 75 | 12.5 |
| 76 | 12.6667 |
| 77 | 12.8333 |
| 78 | 13 |
| 79 | 13.1667 |
| 80 | 13.3333 |
| 81 | 13.5 |
| 82 | 11.7143 |
| 83 | 11.8571 |
| 84 | 12 |
| 85 | 12.1429 |
| 86 | 12.2857 |
| 87 | 12.4286 |
| 88 | 12.5714 |
| 89 | 12.7143 |
| 90 | 12.8571 |
| 91 | 13 |
| 92 | 13.1429 |
| 93 | 13.2857 |
| 94 | 11.75 |
| 95 | 11.875 |
| 96 | 12 |
| 97 | 12.125 |
| 98 | 12.25 |
| 99 | 12.375 |
| 100 | 12.5 |
| 101 | 12.625 |
| 102 | 12.75 |
| 103 | 12.875 |
| 104 | 13 |
| 105 | 13.125 |
| 106 | 13.25 |
| 107 | 11.8889 |
| 108 | 12 |
| 109 | 12.1111 |
| 110 | 12.2222 |
| 111 | 12.3333 |
| 112 | 12.4444 |
| 113 | 12.5556 |
| 114 | 12.6667 |
| 115 | 12.7778 |
| 116 | 12.8889 |
| 117 | 13 |
| 118 | 13.1111 |
| 119 | 11.9 |
| 120 | 12 |
| 121 | 12.1 |
| 122 | 12.2 |
| 123 | 12.3 |
| 124 | 12.4 |
| 125 | 12.5 |
| 126 | 12.6 |
| 127 | 12.7 |
| 128 | 12.8 |
| 129 | 12.9 |
| 130 | 13 |
| 131 | 13.1 |
| 132 | 13.2 |
| 133 | 13.3 |
| 134 | 13.4 |
| 135 | 13.5 |
| 136 | 13.6 |
| 137 | 13.7 |
| 138 | 13.8 |
| 139 | 13.9 |
| 140 | 14 |
| 141 | 14.1 |
| 142 | 14.2 |
| 143 | 14.3 |
| 144 | 14.4 |
| 145 | 14.5 |
| 146 | 14.6 |
| 147 | 14.7 |
| 148 | 14.8 |
| 149 | 14.9 |
| 150 | 15 |
| 151 | 15.1 |
| 152 | 15.2 |
| 153 | 15.3 |
| 154 | 15.4 |
| 155 | 15.5 |
| 156 | 15.6 |
| 157 | 15.7 |
| 158 | 15.8 |
| 159 | 15.9 |
| 160 | 16 |
| 161 | 16.1 |
| 162 | 16.2 |
| 163 | 16.3 |
| 164 | 16.4 |
| 165 | 16.5 |
| 166 | 16.6 |
| 167 | 16.7 |
| 168 | 16.8 |
| 169 | 16.9 |
| 170 | 17 |
| 171 | 17.1 |
| 172 | 17.2 |
| 173 | 17.3 |
| 174 | 17.4 |
| 175 | 17.5 |
| 176 | 17.6 |
| 177 | 17.7 |
| 178 | 17.8 |
| 179 | 17.9 |
| 180 | 18 |
| 181 | 18.1 |
| 182 | 18.2 |
| 183 | 18.3 |
| 184 | 18.4 |
| 185 | 18.5 |
| 186 | 18.6 |
| 187 | 18.7 |
| 188 | 18.8 |
| 189 | 18.9 |
| 190 | 19 |
| 191 | 19.1 |
| 192 | 19.2 |
| 193 | 19.3 |
| 194 | 19.4 |
| 195 | 19.5 |
| 196 | 19.6 |
| 197 | 19.7 |
| 198 | 19.8 |
| 199 | 19.9 |
| 200 | 20 |

Same as Figure S3 for model version H2.

**Model Version 1**

a

b

c

d

e

f

g

h

V1 Functions

|  |  |  |
| --- | --- | --- |
| 0 | -9.81853e-16 | 1.82578e-17 |
| 0.01 | 2.24651e-06 | 4.62312e-08 |
| 0.02 | 4.49527e-06 | 9.31611e-08 |
| 0.03 | 6.74627e-06 | 1.408e-07 |
| 0.04 | 8.99953e-06 | 1.89159e-07 |
| 0.05 | 1.1255e-05 | 2.38249e-07 |
| 0.06 | 1.35128e-05 | 2.88081e-07 |
| 0.07 | 1.57728e-05 | 3.38666e-07 |
| 0.08 | 1.80351e-05 | 3.90016e-07 |
| 0.09 | 2.02997e-05 | 4.42141e-07 |
| 0.1 | 2.25665e-05 | 4.95055e-07 |
| 0.11 | 2.48356e-05 | 5.48768e-07 |
| 0.12 | 2.71069e-05 | 6.03292e-07 |
| 0.13 | 2.93806e-05 | 6.58641e-07 |
| 0.14 | 3.16565e-05 | 7.14826e-07 |
| 0.15 | 3.39346e-05 | 7.71861e-07 |
| 0.16 | 3.62151e-05 | 8.29757e-07 |
| 0.17 | 3.84978e-05 | 8.88528e-07 |
| 0.18 | 4.07829e-05 | 9.48188e-07 |
| 0.19 | 4.30702e-05 | 1.00875e-06 |
| 0.2 | 4.53598e-05 | 1.07023e-06 |
| 0.21 | 4.76516e-05 | 1.13263e-06 |
| 0.22 | 4.99458e-05 | 1.19598e-06 |
| 0.23 | 5.22423e-05 | 1.26029e-06 |
| 0.24 | 5.45411e-05 | 1.32556e-06 |
| 0.25 | 5.68421e-05 | 1.39183e-06 |
| 0.26 | 5.91455e-05 | 1.45909e-06 |
| 0.27 | 6.14512e-05 | 1.52738e-06 |
| 0.28 | 6.37592e-05 | 1.59669e-06 |
| 0.29 | 6.60695e-05 | 1.66705e-06 |
| 0.3 | 6.83821e-05 | 1.73848e-06 |
| 0.31 | 7.0697e-05 | 1.81098e-06 |
| 0.32 | 7.30142e-05 | 1.88458e-06 |
| 0.33 | 7.53337e-05 | 1.9593e-06 |
| 0.34 | 7.76556e-05 | 2.03514e-06 |
| 0.35 | 7.99798e-05 | 2.11213e-06 |
| 0.36 | 8.23063e-05 | 2.19028e-06 |
| 0.37 | 8.46352e-05 | 2.26961e-06 |
| 0.38 | 8.69663e-05 | 2.35015e-06 |
| 0.39 | 8.92999e-05 | 2.43189e-06 |
| 0.4 | 9.16357e-05 | 2.51488e-06 |
| 0.41 | 9.39739e-05 | 2.59912e-06 |
| 0.42 | 9.63144e-05 | 2.68463e-06 |
| 0.43 | 9.86572e-05 | 2.77143e-06 |
| 0.44 | 0.000101002 | 2.85955e-06 |
| 0.45 | 0.00010335 | 2.949e-06 |
| 0.46 | 0.0001057 | 3.0398e-06 |
| 0.47 | 0.000108052 | 3.13197e-06 |
| 0.48 | 0.000110407 | 3.22553e-06 |
| 0.49 | 0.000112764 | 3.32051e-06 |
| 0.5 | 0.000115123 | 3.41693e-06 |
| 0.51 | 0.000117485 | 3.5148e-06 |
| 0.52 | 0.000119849 | 3.61415e-06 |
| 0.53 | 0.000122215 | 3.715e-06 |
| 0.54 | 0.000124584 | 3.81738e-06 |
| 0.55 | 0.000126955 | 3.9213e-06 |
| 0.56 | 0.000129329 | 4.02679e-06 |
| 0.57 | 0.000131704 | 4.13388e-06 |
| 0.58 | 0.000134083 | 4.24259e-06 |
| 0.59 | 0.000136463 | 4.35294e-06 |
| 0.6 | 0.000138846 | 4.46496e-06 |
| 0.61 | 0.000141232 | 4.57867e-06 |
| 0.62 | 0.00014362 | 4.6941e-06 |
| 0.63 | 0.00014601 | 4.81127e-06 |
| 0.64 | 0.000148402 | 4.93021e-06 |
| 0.65 | 0.000150797 | 5.05095e-06 |
| 0.66 | 0.000153195 | 5.17352e-06 |
| 0.67 | 0.000155594 | 5.29794e-06 |
| 0.68 | 0.000157997 | 5.42424e-06 |
| 0.69 | 0.000160401 | 5.55245e-06 |
| 0.7 | 0.000162808 | 5.68259e-06 |
| 0.71 | 0.000165217 | 5.81471e-06 |
| 0.72 | 0.000167629 | 5.94881e-06 |
| 0.73 | 0.000170043 | 6.08495e-06 |
| 0.74 | 0.00017246 | 6.22314e-06 |
| 0.75 | 0.000174879 | 6.36343e-06 |
| 0.76 | 0.0001773 | 6.50583e-06 |
| 0.77 | 0.000179724 | 6.65038e-06 |
| 0.78 | 0.000182151 | 6.79712e-06 |
| 0.79 | 0.000184579 | 6.94608e-06 |
| 0.8 | 0.000187011 | 7.09728e-06 |
| 0.81 | 0.000189444 | 7.25078e-06 |
| 0.82 | 0.00019188 | 7.40659e-06 |
| 0.83 | 0.000194319 | 7.56476e-06 |
| 0.84 | 0.000196759 | 7.72531e-06 |
| 0.85 | 0.000199203 | 7.8883e-06 |
| 0.86 | 0.000201649 | 8.05375e-06 |
| 0.87 | 0.000204097 | 8.22169e-06 |
| 0.88 | 0.000206547 | 8.39218e-06 |
| 0.89 | 0.000209001 | 8.56524e-06 |
| 0.9 | 0.000211456 | 8.74092e-06 |
| 0.91 | 0.000213914 | 8.91925e-06 |
| 0.92 | 0.000216375 | 9.10028e-06 |
| 0.93 | 0.000218838 | 9.28405e-06 |
| 0.94 | 0.000221303 | 9.47059e-06 |
| 0.95 | 0.000223771 | 9.65995e-06 |
| 0.96 | 0.000226241 | 9.85217e-06 |
| 0.97 | 0.000228714 | 1.00473e-05 |
| 0.98 | 0.000231189 | 1.02454e-05 |
| 0.99 | 0.000233667 | 1.04464e-05 |
| 1 | 0.000236148 | 1.06506e-05 |
| 1.01 | 0.00023863 | 1.08577e-05 |
| 1.02 | 0.000241115 | 1.10681e-05 |
| 1.03 | 0.000243603 | 1.12816e-05 |
| 1.04 | 0.000246093 | 1.14983e-05 |
| 1.05 | 0.000248586 | 1.17183e-05 |
| 1.06 | 0.000251081 | 1.19416e-05 |
| 1.07 | 0.000253579 | 1.21683e-05 |
| 1.08 | 0.000256079 | 1.23985e-05 |
| 1.09 | 0.000258582 | 1.26321e-05 |
| 1.1 | 0.000261087 | 1.28692e-05 |
| 1.11 | 0.000263595 | 1.311e-05 |
| 1.12 | 0.000266105 | 1.33543e-05 |
| 1.13 | 0.000268618 | 1.36024e-05 |
| 1.14 | 0.000271133 | 1.38542e-05 |
| 1.15 | 0.000273651 | 1.41098e-05 |
| 1.16 | 0.000276171 | 1.43693e-05 |
| 1.17 | 0.000278694 | 1.46327e-05 |
| 1.18 | 0.000281219 | 1.49e-05 |
| 1.19 | 0.000283747 | 1.51714e-05 |
| 1.2 | 0.000286277 | 1.5447e-05 |
| 1.21 | 0.00028881 | 1.57266e-05 |
| 1.22 | 0.000291345 | 1.60106e-05 |
| 1.23 | 0.000293883 | 1.62987e-05 |
| 1.24 | 0.000296424 | 1.65913e-05 |
| 1.25 | 0.000298967 | 1.68883e-05 |
| 1.26 | 0.000301512 | 1.71897e-05 |
| 1.27 | 0.000304061 | 1.74958e-05 |
| 1.28 | 0.000306611 | 1.78064e-05 |
| 1.29 | 0.000309164 | 1.81217e-05 |
| 1.3 | 0.00031172 | 1.84419e-05 |
| 1.31 | 0.000314279 | 1.87668e-05 |
| 1.32 | 0.000316839 | 1.90967e-05 |
| 1.33 | 0.000319403 | 1.94315e-05 |
| 1.34 | 0.000321969 | 1.97714e-05 |
| 1.35 | 0.000324537 | 2.01164e-05 |
| 1.36 | 0.000327109 | 2.04667e-05 |
| 1.37 | 0.000329682 | 2.08222e-05 |
| 1.38 | 0.000332259 | 2.11831e-05 |
| 1.39 | 0.000334838 | 2.15495e-05 |
| 1.4 | 0.000337419 | 2.19214e-05 |
| 1.41 | 0.000340003 | 2.2299e-05 |
| 1.42 | 0.00034259 | 2.26822e-05 |
| 1.43 | 0.000345179 | 2.30712e-05 |
| 1.44 | 0.000347771 | 2.34661e-05 |
| 1.45 | 0.000350365 | 2.3867e-05 |
| 1.46 | 0.000352962 | 2.42739e-05 |
| 1.47 | 0.000355562 | 2.4687e-05 |
| 1.48 | 0.000358164 | 2.51063e-05 |
| 1.49 | 0.000360769 | 2.5532e-05 |
| 1.5 | 0.000363376 | 2.59641e-05 |
| 1.51 | 0.000365986 | 2.64027e-05 |
| 1.52 | 0.000368599 | 2.6848e-05 |
| 1.53 | 0.000371214 | 2.73e-05 |
| 1.54 | 0.000373832 | 2.77588e-05 |
| 1.55 | 0.000376452 | 2.82245e-05 |
| 1.56 | 0.000379075 | 2.86973e-05 |
| 1.57 | 0.000381701 | 2.91773e-05 |
| 1.58 | 0.000384329 | 2.96645e-05 |
| 1.59 | 0.00038696 | 3.0159e-05 |
| 1.6 | 0.000389594 | 3.0661e-05 |
| 1.61 | 0.00039223 | 3.11706e-05 |
| 1.62 | 0.000394869 | 3.1688e-05 |
| 1.63 | 0.000397511 | 3.22131e-05 |
| 1.64 | 0.000400155 | 3.27461e-05 |
| 1.65 | 0.000402802 | 3.32873e-05 |
| 1.66 | 0.000405451 | 3.38366e-05 |
| 1.67 | 0.000408103 | 3.43942e-05 |
| 1.68 | 0.000410758 | 3.49602e-05 |
| 1.69 | 0.000413415 | 3.55348e-05 |
| 1.7 | 0.000416075 | 3.61181e-05 |
| 1.71 | 0.000418738 | 3.67101e-05 |
| 1.72 | 0.000421403 | 3.73112e-05 |
| 1.73 | 0.000424071 | 3.79213e-05 |
| 1.74 | 0.000426742 | 3.85406e-05 |
| 1.75 | 0.000429416 | 3.91693e-05 |
| 1.76 | 0.000432092 | 3.98075e-05 |
| 1.77 | 0.00043477 | 4.04554e-05 |
| 1.78 | 0.000437452 | 4.1113e-05 |
| 1.79 | 0.000440136 | 4.17806e-05 |
| 1.8 | 0.000442823 | 4.24582e-05 |
| 1.81 | 0.000445512 | 4.31461e-05 |
| 1.82 | 0.000448204 | 4.38444e-05 |
| 1.83 | 0.000450899 | 4.45533e-05 |
| 1.84 | 0.000453597 | 4.52728e-05 |
| 1.85 | 0.000456297 | 4.60033e-05 |
| 1.86 | 0.000459 | 4.67448e-05 |
| 1.87 | 0.000461706 | 4.74974e-05 |
| 1.88 | 0.000464414 | 4.82615e-05 |
| 1.89 | 0.000467125 | 4.90371e-05 |
| 1.9 | 0.000469839 | 4.98244e-05 |
| 1.91 | 0.000472555 | 5.06237e-05 |
| 1.92 | 0.000475275 | 5.1435e-05 |
| 1.93 | 0.000477997 | 5.22585e-05 |
| 1.94 | 0.000480721 | 5.30946e-05 |
| 1.95 | 0.000483449 | 5.39432e-05 |
| 1.96 | 0.000486179 | 5.48047e-05 |
| 1.97 | 0.000488912 | 5.56792e-05 |
| 1.98 | 0.000491647 | 5.65669e-05 |
| 1.99 | 0.000494386 | 5.7468e-05 |

V1 Total fitness

|  |  |  |  |
| --- | --- | --- | --- |
| 1 | 0.000236148 | 0.000236148 | 0.000236148 |
| 2 | 0.000497127 | 0.000497127 | 0.000497127 |
| 3 | 0.000785548 | 0.000785548 | 0.000785548 |
| 4 | 0.0011043 | 0.0011043 | 0.0011043 |
| 5 | 0.00145656 | 0.00145656 | 0.00145656 |
| 6 | 0.00184586 | 0.0247231 | 0.0247232 |
| 7 | 0.00227609 | 0.109866 | 0.109866 |
| 8 | 0.00275156 | 0.474256 | 0.474256 |
| 9 | 1.82425 | 1.82425 | 1.82425 |
| 10 | 5 | 5 | 5 |
| 11 | 8.17574 | 8.17574 | 8.17574 |
| 12 | 9.52574 | 9.52574 | 9.52574 |
| 13 | 9.89013 | 9.89013 | 9.89013 |
| 14 | 9.97527 | 9.97527 | 9.97527 |
| 15 | 9.99447 | 9.99447 | 9.99447 |
| 16 | 9.99877 | 9.99877 | 9.99877 |
| 17 | 9.99972 | 9.99972 | 9.99972 |
| 18 | 9.99999 | 9.99999 | 9.99999 |
| 19 | 10.0002 | 10.0002 | 10.0002 |
| 20 | 10.0005 | 10.0005 | 10.0005 |
| 21 | 13.5836 | 13.5836 | 13.5836 |
| 22 | 16.3515 | 16.3515 | 16.3515 |
| 23 | 18.093 | 18.093 | 18.093 |
| 24 | 19.0515 | 19.0515 | 19.0515 |
| 25 | 19.5405 | 19.5405 | 19.5405 |
| 26 | 19.7803 | 19.7803 | 19.7803 |
| 27 | 19.8956 | 19.8956 | 19.8956 |
| 28 | 19.9505 | 19.9505 | 19.9505 |
| 29 | 19.9766 | 19.9766 | 19.9766 |
| 30 | 19.9889 | 19.9889 | 19.9889 |
| 31 | 19.9948 | 19.9948 | 19.9948 |
| 32 | 21.9318 | 21.9318 | 21.9318 |
| 33 | 24.5272 | 24.5272 | 24.5272 |
| 34 | 26.4239 | 26.4239 | 26.4239 |
| 35 | 27.7243 | 27.7243 | 27.7243 |
| 36 | 28.5772 | 28.5772 | 28.5772 |
| 37 | 29.1206 | 29.1206 | 29.1206 |
| 38 | 29.4604 | 29.4604 | 29.4604 |
| 39 | 29.6704 | 29.6704 | 29.6704 |
| 40 | 29.7992 | 29.7992 | 29.7992 |
| 41 | 29.8779 | 29.8779 | 29.8779 |
| 42 | 29.9258 | 29.9258 | 29.9258 |
| 43 | 29.955 | 30.1966 | 30.1966 |
| 44 | 29.9727 | 32.703 | 32.703 |
| 45 | 29.9834 | 34.6814 | 34.6814 |
| 46 | 29.9899 | 36.186 | 36.186 |
| 47 | 29.9939 | 37.2981 | 37.2981 |
| 48 | 29.9963 | 38.103 | 38.103 |
| 49 | 29.9978 | 38.6766 | 38.6766 |
| 50 | 29.9986 | 39.0809 | 39.0809 |
| 51 | 29.9992 | 39.3637 | 39.3637 |
| 52 | 29.9995 | 39.5605 | 39.5605 |
| 53 | 29.9997 | 39.6969 | 39.6969 |
| 54 | 30 | 39.7912 | 39.7912 |
| 55 | 30.0003 | 40.8787 | 40.8787 |
| 56 | 30.0006 | 42.9074 | 42.9074 |
| 57 | 30.001 | 44.5452 | 44.5452 |
| 58 | 30.0015 | 45.8414 | 45.8414 |
| 59 | 30.0019 | 46.8513 | 46.8513 |
| 60 | 30.0025 | 47.6287 | 47.6287 |
| 61 | 30.0031 | 48.2214 | 48.2214 |
| 62 | 30.0039 | 48.6701 | 48.6701 |
| 63 | 30.0047 | 49.008 | 49.008 |
| 64 | 30.0056 | 49.2613 | 49.2613 |
| 65 | 30.0066 | 49.4507 | 49.4507 |
| 66 | 30.0078 | 49.5919 | 49.5919 |
| 67 | 30.0091 | 49.697 | 51.1172 |
| 68 | 30.0107 | 49.7752 | 52.8478 |
| 69 | 30.0124 | 49.8333 | 54.279 |
| 70 | 30.0143 | 49.8764 | 55.4485 |
| 71 | 30.0165 | 49.9084 | 56.3948 |
| 72 | 30.019 | 49.9321 | 57.1544 |
| 73 | 30.0219 | 49.9497 | 57.7604 |
| 74 | 30.0251 | 49.9627 | 58.2413 |
| 75 | 30.0287 | 49.9724 | 58.6214 |
| 76 | 30.0328 | 49.9795 | 58.9208 |
| 77 | 30.0375 | 49.9848 | 59.1562 |
| 78 | 30.0427 | 49.9888 | 59.3408 |
| 79 | 30.0487 | 49.9917 | 59.4854 |
| 80 | 30.0555 | 49.9938 | 62.65 |
| 81 | 30.0631 | 49.9954 | 63.945 |
| 82 | 30.0718 | 49.9966 | 65.03 |
| 83 | 30.0816 | 49.9975 | 65.9329 |
| 84 | 30.0927 | 49.9981 | 66.6802 |
| 85 | 30.1053 | 49.9986 | 67.2958 |
| 86 | 30.1196 | 49.999 | 67.801 |
| 87 | 30.1357 | 49.9992 | 68.2143 |
| 88 | 30.1539 | 49.9995 | 68.5516 |
| 89 | 30.1745 | 49.9998 | 68.8263 |
| 90 | 30.1978 | 50.0001 | 69.0496 |
| 91 | 30.2241 | 50.0005 | 69.2309 |
| 92 | 30.2539 | 50.0009 | 69.3779 |
| 93 | 30.2875 | 50.0014 | 69.4971 |
| 94 | 30.3254 | 50.002 | 69.5935 |
| 95 | 30.3683 | 50.0027 | 69.6715 |
| 96 | 30.4166 | 50.0035 | 69.7347 |
| 97 | 30.471 | 50.0044 | 69.7857 |
| 98 | 30.5324 | 50.0055 | 69.8269 |
| 99 | 30.6014 | 50.0068 | 69.8602 |
| 100 | 30.6791 | 50.0082 | 69.8871 |
| 101 | 30.7664 | 50.0099 | 69.9089 |
| 102 | 30.8643 | 50.0119 | 69.9264 |
| 103 | 30.9741 | 50.0142 | 69.9406 |
| 104 | 31.0971 | 50.0168 | 69.9521 |
| 105 | 31.2346 | 50.0199 | 69.9613 |
| 106 | 31.388 | 50.0235 | 69.9688 |
| 107 | 31.559 | 50.0276 | 69.9748 |
| 108 | 31.7491 | 50.0324 | 69.9796 |
| 109 | 31.9599 | 50.038 | 69.9836 |
| 110 | 32.1934 | 50.0445 | 69.9867 |
| 111 | 32.451 | 50.0521 | 69.9893 |
| 112 | 32.7346 | 50.0608 | 69.9914 |
| 113 | 33.0457 | 50.071 | 69.993 |
| 114 | 33.3858 | 50.0828 | 69.9944 |
| 115 | 33.7563 | 50.0964 | 69.9955 |
| 116 | 34.1584 | 50.1123 | 69.9963 |
| 117 | 34.5928 | 50.1306 | 69.997 |
| 118 | 35.0601 | 50.1519 | 69.9976 |
| 119 | 35.5603 | 50.1766 | 69.9981 |
| 120 | 36.0933 | 50.2051 | 69.9984 |
| 121 | 36.6583 | 50.2381 | 69.9987 |
| 122 | 37.254 | 50.2762 | 69.999 |
| 123 | 37.8788 | 50.3203 | 69.9992 |
| 124 | 38.5304 | 50.3711 | 69.9995 |
| 125 | 39.2062 | 50.4297 | 69.9999 |
| 126 | 39.9033 | 50.4973 | 70.0004 |
| 127 | 40.6184 | 50.5749 | 70.0009 |
| 128 | 41.3478 | 50.664 | 70.0016 |
| 129 | 42.0878 | 50.7662 | 70.0023 |
| 130 | 42.8346 | 50.883 | 70.0033 |
| 131 | 43.5844 | 51.0164 | 70.0044 |
| 132 | 44.3333 | 51.1683 | 70.0058 |
| 133 | 45.0778 | 51.3406 | 70.0074 |
| 134 | 45.8143 | 51.5357 | 70.0094 |
| 135 | 46.5398 | 51.7556 | 70.0118 |
| 136 | 47.2513 | 52.0026 | 70.0147 |
| 137 | 47.9462 | 52.2789 | 70.0182 |
| 138 | 48.6223 | 52.5864 | 70.0225 |
| 139 | 49.2777 | 52.927 | 70.0275 |
| 140 | 49.9108 | 53.3023 | 70.0337 |
| 141 | 50.5203 | 53.7133 | 70.041 |
| 142 | 51.1053 | 54.1608 | 70.0499 |
| 143 | 51.6651 | 54.645 | 70.0606 |
| 144 | 52.1994 | 55.1655 | 70.0734 |
| 145 | 52.7079 | 55.7214 | 70.0889 |
| 146 | 53.1909 | 56.3109 | 70.1075 |
| 147 | 53.6484 | 56.932 | 70.1298 |
| 148 | 54.0809 | 57.5821 | 70.1566 |
| 149 | 54.4891 | 58.2578 | 70.1887 |
| 150 | 54.8734 | 58.9557 | 70.2271 |
| 151 | 55.2348 | 59.6721 | 70.2731 |
| 152 | 55.574 | 60.4029 | 70.3279 |
| 153 | 55.892 | 61.144 | 70.3932 |
| 154 | 56.1896 | 61.8915 | 70.4708 |
| 155 | 56.4677 | 62.6414 | 70.5627 |
| 156 | 56.7274 | 63.3899 | 70.6712 |
| 157 | 56.9697 | 64.1334 | 70.7989 |
| 158 | 57.1953 | 64.8688 | 70.9485 |
| 159 | 57.4053 | 65.5931 | 71.1229 |
| 160 | 57.6006 | 66.3035 | 71.3251 |
| 161 | 57.7821 | 66.9978 | 71.558 |
| 162 | 57.9505 | 67.674 | 71.8245 |
| 163 | 58.1069 | 68.3304 | 72.1273 |
| 164 | 58.2518 | 68.9656 | 72.4685 |
| 165 | 58.3861 | 69.5787 | 72.8499 |
| 166 | 58.5104 | 70.1687 | 73.2726 |
| 167 | 58.6256 | 70.7351 | 73.7367 |
| 168 | 58.7321 | 71.2777 | 74.2417 |
| 169 | 58.8306 | 71.7963 | 74.7865 |
| 170 | 58.9216 | 72.2909 | 75.3687 |
| 171 | 59.0058 | 72.7617 | 75.9856 |
| 172 | 59.0835 | 73.2092 | 76.6338 |
| 173 | 59.1553 | 73.6337 | 77.3094 |
| 174 | 59.2216 | 74.0359 | 78.0082 |
| 175 | 59.2827 | 74.4163 | 78.726 |
| 176 | 59.3392 | 74.7756 | 79.4581 |
| 177 | 59.3912 | 75.1146 | 80.2004 |
| 178 | 59.4392 | 75.4341 | 80.9484 |
| 179 | 59.4835 | 75.7348 | 81.6983 |
| 180 | 59.5243 | 76.0176 | 82.4463 |
| 181 | 59.562 | 76.2833 | 83.189 |
| 182 | 59.5966 | 76.5327 | 83.9232 |
| 183 | 59.6286 | 76.7666 | 84.6462 |
| 184 | 59.6581 | 76.9858 | 85.3556 |
| 185 | 59.6852 | 77.1911 | 86.0493 |
| 186 | 59.7102 | 77.3832 | 86.7256 |
| 187 | 59.7332 | 77.5628 | 87.3829 |
| 188 | 59.7544 | 77.7308 | 88.0201 |
| 189 | 59.7739 | 77.8877 | 88.6363 |
| 190 | 59.7919 | 78.0343 | 89.2308 |
| 191 | 59.8085 | 78.171 | 89.8031 |
| 192 | 59.8237 | 78.2986 | 90.353 |
| 193 | 59.8378 | 78.4176 | 90.8803 |
| 194 | 59.8507 | 78.5285 | 91.3851 |
| 195 | 59.8626 | 78.6319 | 91.8677 |
| 196 | 59.8735 | 78.7282 | 92.3282 |
| 197 | 59.8836 | 78.8179 | 92.767 |
| 198 | 59.8929 | 78.9015 | 93.1847 |
| 199 | 59.9015 | 78.9792 | 93.5818 |
| 200 | 59.9093 | 79.0515 | 93.9588 |

V1 T-II-biased

|  | Male | Female |
| --- | --- | --- |
| 1 | 1 | 1.59796e-08 |
| 2 | 1 | 9.1248e-09 |
| 3 | 1 | 5.54976e-09 |
| 4 | 1 | 3.79266e-09 |
| 5 | 1 | 2.76178e-09 |
| 6 | 1 | 2.09319e-09 |
| 7 | 1 | 1.63063e-09 |
| 8 | 1 | 1.03681e-09 |
| 9 | 3.0906e-10 | 1 |
| 10 | 8.29681e-12 | 1 |
| 11 | 2.52868e-10 | 1 |
| 12 | 7.65431e-10 | 1 |
| 13 | 1.47039e-10 | 1 |
| 14 | 4.83515e-10 | 1 |
| 15 | 3.29496e-10 | 1 |
| 16 | 3.83848e-10 | 1 |
| 17 | 2.52227e-09 | 1 |
| 18 | 0.0353345 | 0.964665 |
| 19 | 0.0898686 | 0.910131 |
| 20 | 0.138946 | 0.861054 |
| 21 | 4.07972e-12 | 1 |
| 22 | 5.68951e-12 | 1 |
| 23 | 9.41055e-12 | 1 |
| 24 | 3.44444e-12 | 1 |
| 25 | 1.33129e-10 | 1 |
| 26 | 2.6465e-10 | 1 |
| 27 | 2.13633e-11 | 1 |
| 28 | 1.74039e-10 | 1 |
| 29 | 7.14486e-11 | 1 |
| 30 | 1.48181e-10 | 1 |
| 31 | 7.82785e-11 | 1 |
| 32 | 2.96715e-12 | 1 |
| 33 | 3.79301e-12 | 1 |
| 34 | 5.22983e-12 | 1 |
| 35 | 2.43501e-11 | 1 |
| 36 | 2.29628e-12 | 1 |
| 37 | 7.09618e-11 | 1 |
| 38 | 2.22681e-11 | 1 |
| 39 | 1.41147e-10 | 1 |
| 40 | 1.40709e-11 | 1 |
| 41 | 9.01866e-11 | 1 |
| 42 | 1.45034e-10 | 1 |
| 43 | 2.34033e-10 | 1 |
| 44 | 7.58224e-11 | 1 |
| 45 | 2.4695e-11 | 1 |
| 46 | 2.02754e-10 | 1 |
| 47 | 6.74547e-11 | 1 |
| 48 | 1.14777e-10 | 1 |
| 49 | 3.25697e-09 | 1 |
| 50 | 6.28314e-09 | 1 |
| 51 | 2.9799e-09 | 1 |
| 52 | 1.06273e-07 | 1 |
| 53 | 0.0184193 | 0.981581 |
| 54 | 0.0412232 | 0.958777 |
| 55 | 0.0632021 | 0.936798 |
| 56 | 0.0843949 | 0.915605 |
| 57 | 0.104843 | 0.895157 |
| 58 | 0.124586 | 0.875414 |
| 59 | 0.143658 | 0.856342 |
| 60 | 0.162096 | 0.837904 |
| 61 | 0.179929 | 0.820071 |
| 62 | 0.197187 | 0.802813 |
| 63 | 0.213896 | 0.786104 |
| 64 | 0.230083 | 0.769917 |
| 65 | 0.245771 | 0.754229 |
| 66 | 0.260984 | 0.739016 |
| 67 | 0.275742 | 0.724258 |
| 68 | 0.290065 | 0.709935 |
| 69 | 0.303973 | 0.696027 |
| 70 | 0.317483 | 0.682517 |
| 71 | 0.330612 | 0.669388 |
| 72 | 0.343376 | 0.656624 |
| 73 | 0.355789 | 0.644211 |
| 74 | 0.367865 | 0.632135 |
| 75 | 0.379619 | 0.620381 |
| 76 | 0.391062 | 0.608938 |
| 77 | 0.402207 | 0.597793 |
| 78 | 0.413065 | 0.586935 |
| 79 | 0.423647 | 0.576353 |
| 80 | 0.433962 | 0.566038 |
| 81 | 0.444021 | 0.555979 |
| 82 | 0.453832 | 0.546168 |
| 83 | 0.463405 | 0.536595 |
| 84 | 0.472747 | 0.527253 |
| 85 | 0.481866 | 0.518134 |
| 86 | 0.49077 | 0.50923 |
| 87 | 0.499466 | 0.500534 |
| 88 | 0.50796 | 0.49204 |
| 89 | 0.516259 | 0.483741 |
| 90 | 0.524368 | 0.475632 |
| 91 | 0.532293 | 0.467707 |
| 92 | 0.540039 | 0.459961 |
| 93 | 0.547612 | 0.452388 |
| 94 | 0.555016 | 0.444984 |
| 95 | 0.562255 | 0.437745 |
| 96 | 0.569334 | 0.430666 |
| 97 | 0.576255 | 0.423745 |
| 98 | 0.583023 | 0.416977 |
| 99 | 0.589641 | 0.410359 |
| 100 | 0.596111 | 0.403889 |
| 101 | 0.602436 | 0.397564 |
| 102 | 0.608618 | 0.391382 |
| 103 | 0.614659 | 0.385341 |
| 104 | 0.62056 | 0.37944 |
| 105 | 0.626324 | 0.373676 |
| 106 | 0.63195 | 0.36805 |
| 107 | 0.637439 | 0.362561 |
| 108 | 0.642793 | 0.357207 |
| 109 | 0.64801 | 0.35199 |
| 110 | 0.65309 | 0.34691 |
| 111 | 0.658034 | 0.341966 |
| 112 | 0.662839 | 0.337161 |
| 113 | 0.667505 | 0.332495 |
| 114 | 0.672032 | 0.327968 |
| 115 | 0.676416 | 0.323584 |
| 116 | 0.680658 | 0.319342 |
| 117 | 0.684755 | 0.315245 |
| 118 | 0.688707 | 0.311293 |
| 119 | 0.692512 | 0.307488 |
| 120 | 0.696169 | 0.303831 |
| 121 | 0.699679 | 0.300321 |
| 122 | 0.70304 | 0.29696 |
| 123 | 0.706253 | 0.293747 |
| 124 | 0.70932 | 0.29068 |
| 125 | 0.712243 | 0.287757 |
| 126 | 0.715022 | 0.284978 |
| 127 | 0.717663 | 0.282337 |
| 128 | 0.720167 | 0.279833 |
| 129 | 0.72254 | 0.27746 |
| 130 | 0.724786 | 0.275214 |
| 131 | 0.726911 | 0.273089 |
| 132 | 0.728919 | 0.271081 |
| 133 | 0.730817 | 0.269183 |
| 134 | 0.73261 | 0.26739 |
| 135 | 0.734305 | 0.265695 |
| 136 | 0.735906 | 0.264094 |
| 137 | 0.737421 | 0.262579 |
| 138 | 0.738854 | 0.261146 |
| 139 | 0.740211 | 0.259789 |
| 140 | 0.741498 | 0.258502 |
| 141 | 0.742718 | 0.257282 |
| 142 | 0.743878 | 0.256122 |
| 143 | 0.74498 | 0.25502 |
| 144 | 0.746031 | 0.253969 |
| 145 | 0.747032 | 0.252968 |
| 146 | 0.747988 | 0.252012 |
| 147 | 0.748902 | 0.251098 |
| 148 | 0.749778 | 0.250222 |
| 149 | 0.750617 | 0.249383 |
| 150 | 0.751424 | 0.248576 |
| 151 | 0.752199 | 0.247801 |
| 152 | 0.752945 | 0.247055 |
| 153 | 0.753665 | 0.246335 |
| 154 | 0.75436 | 0.24564 |
| 155 | 0.755031 | 0.244969 |
| 156 | 0.755681 | 0.244319 |
| 157 | 0.756311 | 0.243689 |
| 158 | 0.756923 | 0.243077 |
| 159 | 0.757516 | 0.242484 |
| 160 | 0.758093 | 0.241907 |
| 161 | 0.758655 | 0.241345 |
| 162 | 0.759202 | 0.240798 |
| 163 | 0.759736 | 0.240264 |
| 164 | 0.760257 | 0.239743 |
| 165 | 0.760766 | 0.239234 |
| 166 | 0.761263 | 0.238737 |
| 167 | 0.76175 | 0.23825 |
| 168 | 0.762226 | 0.237774 |
| 169 | 0.762693 | 0.237307 |
| 170 | 0.76315 | 0.23685 |
| 171 | 0.763599 | 0.236401 |
| 172 | 0.76404 | 0.23596 |
| 173 | 0.764473 | 0.235527 |
| 174 | 0.764898 | 0.235102 |
| 175 | 0.765316 | 0.234684 |
| 176 | 0.765727 | 0.234273 |
| 177 | 0.766131 | 0.233869 |
| 178 | 0.766529 | 0.233471 |
| 179 | 0.766922 | 0.233078 |
| 180 | 0.767308 | 0.232692 |
| 181 | 0.767688 | 0.232312 |
| 182 | 0.768063 | 0.231937 |
| 183 | 0.768433 | 0.231567 |
| 184 | 0.768798 | 0.231202 |
| 185 | 0.769158 | 0.230842 |
| 186 | 0.769513 | 0.230487 |
| 187 | 0.769864 | 0.230136 |
| 188 | 0.77021 | 0.22979 |
| 189 | 0.770551 | 0.229449 |
| 190 | 0.770889 | 0.229111 |
| 191 | 0.771222 | 0.228778 |
| 192 | 0.771552 | 0.228448 |
| 193 | 0.771877 | 0.228123 |
| 194 | 0.772199 | 0.227801 |
| 195 | 0.772517 | 0.227483 |
| 196 | 0.772832 | 0.227168 |
| 197 | 0.773143 | 0.226857 |
| 198 | 0.77345 | 0.22655 |
| 199 | 0.773754 | 0.226246 |
| 200 | 0.774055 | 0.225945 |

V1 T-II-biased

|  | Male | Female |
| --- | --- | --- |
| 1 | 1 | 0 |
| 2 | 1 | 0 |
| 3 | 1 | 0 |
| 4 | 1 | 0 |
| 5 | 1 | 0 |
| 6 | 1 | 0 |
| 7 | 1 | 0 |
| 8 | 1 | 0 |
| 9 | 0 | 1 |
| 10 | 0 | 1 |
| 11 | 0 | 1 |
| 12 | 0 | 1 |
| 13 | 0 | 1 |
| 14 | 0 | 1 |
| 15 | 0 | 1 |
| 16 | 0 | 1 |
| 17 | 0 | 1 |
| 18 | 1 | 1 |
| 19 | 1 | 1 |
| 20 | 1 | 1 |
| 21 | 0 | 2 |
| 22 | 0 | 2 |
| 23 | 0 | 2 |
| 24 | 0 | 2 |
| 25 | 0 | 2 |
| 26 | 0 | 2 |
| 27 | 0 | 2 |
| 28 | 0 | 2 |
| 29 | 0 | 2 |
| 30 | 0 | 2 |
| 31 | 0 | 2 |
| 32 | 0 | 3 |
| 33 | 0 | 3 |
| 34 | 0 | 3 |
| 35 | 0 | 3 |
| 36 | 0 | 3 |
| 37 | 0 | 3 |
| 38 | 0 | 3 |
| 39 | 0 | 3 |
| 40 | 0 | 3 |
| 41 | 0 | 3 |
| 42 | 0 | 3 |
| 43 | 0 | 3 |
| 44 | 0 | 3 |
| 45 | 0 | 3 |
| 46 | 0 | 3 |
| 47 | 0 | 3 |
| 48 | 0 | 3 |
| 49 | 0 | 3 |
| 50 | 0 | 3 |
| 51 | 0 | 3 |
| 52 | 0 | 3 |
| 53 | 1 | 3 |
| 54 | 1 | 3 |
| 55 | 1 | 3 |
| 56 | 1 | 3 |
| 57 | 1 | 3 |
| 58 | 1 | 3 |
| 59 | 1 | 3 |
| 60 | 1 | 3 |
| 61 | 1 | 3 |
| 62 | 1 | 3 |
| 63 | 1 | 3 |
| 64 | 1 | 3 |
| 65 | 1 | 3 |
| 66 | 1 | 3 |
| 67 | 1 | 3 |
| 68 | 1 | 3 |
| 69 | 1 | 3 |
| 70 | 1 | 3 |
| 71 | 1 | 3 |
| 72 | 1 | 3 |
| 73 | 1 | 3 |
| 74 | 1 | 3 |
| 75 | 1 | 3 |
| 76 | 1 | 3 |
| 77 | 1 | 3 |
| 78 | 1 | 3 |
| 79 | 1 | 3 |
| 80 | 1 | 3 |
| 81 | 1 | 3 |
| 82 | 1 | 3 |
| 83 | 1 | 3 |
| 84 | 1 | 3 |
| 85 | 1 | 3 |
| 86 | 1 | 3 |
| 87 | 1 | 3 |
| 88 | 1 | 3 |
| 89 | 1 | 3 |
| 90 | 1 | 3 |
| 91 | 1 | 3 |
| 92 | 1 | 3 |
| 93 | 1 | 3 |
| 94 | 1 | 3 |
| 95 | 1 | 3 |
| 96 | 1 | 3 |
| 97 | 1 | 3 |
| 98 | 1 | 3 |
| 99 | 1 | 3 |
| 100 | 1 | 3 |
| 101 | 1 | 3 |
| 102 | 1 | 3 |
| 103 | 1 | 3 |
| 104 | 1 | 3 |
| 105 | 1 | 3 |
| 106 | 1 | 3 |
| 107 | 1 | 3 |
| 108 | 1 | 3 |
| 109 | 1 | 3 |
| 110 | 1 | 3 |
| 111 | 1 | 3 |
| 112 | 1 | 3 |
| 113 | 1 | 3 |
| 114 | 1 | 3 |
| 115 | 1 | 3 |
| 116 | 1 | 3 |
| 117 | 1 | 3 |
| 118 | 1 | 3 |
| 119 | 1 | 3 |
| 120 | 1 | 3 |
| 121 | 1 | 3 |
| 122 | 1 | 3 |
| 123 | 1 | 3 |
| 124 | 1 | 3 |
| 125 | 1 | 3 |
| 126 | 1 | 3 |
| 127 | 1 | 3 |
| 128 | 1 | 3 |
| 129 | 1 | 3 |
| 130 | 1 | 3 |
| 131 | 1 | 3 |
| 132 | 1 | 3 |
| 133 | 1 | 3 |
| 134 | 1 | 3 |
| 135 | 1 | 3 |
| 136 | 1 | 3 |
| 137 | 1 | 3 |
| 138 | 1 | 3 |
| 139 | 1 | 3 |
| 140 | 1 | 3 |
| 141 | 1 | 3 |
| 142 | 1 | 3 |
| 143 | 1 | 3 |
| 144 | 1 | 3 |
| 145 | 1 | 3 |
| 146 | 1 | 3 |
| 147 | 1 | 3 |
| 148 | 1 | 3 |
| 149 | 1 | 3 |
| 150 | 1 | 3 |
| 151 | 1 | 3 |
| 152 | 1 | 3 |
| 153 | 1 | 3 |
| 154 | 1 | 3 |
| 155 | 1 | 3 |
| 156 | 1 | 3 |
| 157 | 1 | 3 |
| 158 | 1 | 3 |
| 159 | 1 | 3 |
| 160 | 1 | 3 |
| 161 | 1 | 3 |
| 162 | 1 | 3 |
| 163 | 1 | 3 |
| 164 | 1 | 3 |
| 165 | 1 | 3 |
| 166 | 1 | 3 |
| 167 | 1 | 3 |
| 168 | 1 | 3 |
| 169 | 1 | 3 |
| 170 | 1 | 3 |
| 171 | 1 | 3 |
| 172 | 1 | 3 |
| 173 | 1 | 3 |
| 174 | 1 | 3 |
| 175 | 1 | 3 |
| 176 | 1 | 3 |
| 177 | 1 | 3 |
| 178 | 1 | 3 |
| 179 | 1 | 3 |
| 180 | 1 | 3 |
| 181 | 1 | 3 |
| 182 | 1 | 3 |
| 183 | 1 | 3 |
| 184 | 1 | 3 |
| 185 | 1 | 3 |
| 186 | 1 | 3 |
| 187 | 1 | 3 |
| 188 | 1 | 3 |
| 189 | 1 | 3 |
| 190 | 1 | 3 |
| 191 | 1 | 3 |
| 192 | 1 | 3 |
| 193 | 1 | 3 |
| 194 | 1 | 3 |
| 195 | 1 | 3 |
| 196 | 1 | 3 |
| 197 | 1 | 3 |
| 198 | 1 | 3 |
| 199 | 1 | 3 |
| 200 | 1 | 3 |

V1 Equal

|  | Male | Female |
| --- | --- | --- |
| 1 | 1 | 2.67569e-08 |
| 2 | 1 | 1.53218e-08 |
| 3 | 1 | 9.33319e-09 |
| 4 | 1 | 5.10622e-09 |
| 5 | 1 | 3.71903e-09 |
| 6 | 1.00747e-07 | 1 |
| 7 | 1.9505e-08 | 1 |
| 8 | 4.10056e-09 | 1 |
| 9 | 2.20758e-10 | 1 |
| 10 | 1.18526e-12 | 1 |
| 11 | 1.8062e-10 | 1 |
| 12 | 5.46741e-10 | 1 |
| 13 | 1.05031e-10 | 1 |
| 14 | 3.45419e-10 | 1 |
| 15 | 2.35511e-10 | 1 |
| 16 | 3.4377e-10 | 1 |
| 17 | 9.17721e-09 | 1 |
| 18 | 0.0353341 | 0.964666 |
| 19 | 0.0898681 | 0.910132 |
| 20 | 0.138946 | 0.861054 |
| 21 | 2.91408e-12 | 1 |
| 22 | 4.06393e-12 | 1 |
| 23 | 6.72182e-12 | 1 |
| 24 | 4.92059e-11 | 1 |
| 25 | 4.7546e-12 | 1 |
| 26 | 1.89036e-12 | 1 |
| 27 | 3.05191e-12 | 1 |
| 28 | 6.21574e-10 | 1 |
| 29 | 6.37921e-11 | 1 |
| 30 | 5.29184e-10 | 1 |
| 31 | 1.1183e-09 | 1 |
| 32 | 2.11939e-12 | 1 |
| 33 | 2.70929e-12 | 1 |
| 34 | 3.73559e-12 | 1 |
| 35 | 1.08706e-12 | 1 |
| 36 | 1.64021e-12 | 1 |
| 37 | 5.06871e-11 | 1 |
| 38 | 7.95291e-11 | 1 |
| 39 | 6.30122e-12 | 1 |
| 40 | 4.02025e-11 | 1 |
| 41 | 1.28838e-11 | 1 |
| 42 | 4.14382e-10 | 1 |
| 43 | 6.70426e-12 | 1 |
| 44 | 1.01598e-11 | 1 |
| 45 | 1.28523e-11 | 1 |
| 46 | 1.68046e-11 | 1 |
| 47 | 9.01037e-11 | 1 |
| 48 | 6.15074e-12 | 1 |
| 49 | 8.50971e-12 | 1 |
| 50 | 1.90184e-10 | 1 |
| 51 | 6.68707e-13 | 1 |
| 52 | 3.78074e-10 | 1 |
| 53 | 5.36379e-10 | 1 |
| 54 | 1.90755e-10 | 1 |
| 55 | 1.62557e-10 | 1 |
| 56 | 3.91237e-11 | 1 |
| 57 | 4.81413e-09 | 1 |
| 58 | 3.01528e-10 | 1 |
| 59 | 7.66158e-09 | 1 |
| 60 | 7.87294e-09 | 1 |
| 61 | 6.37497e-10 | 1 |
| 62 | 1.66233e-10 | 1 |
| 63 | 2.1783e-10 | 1 |
| 64 | 2.86566e-10 | 1 |
| 65 | 3.02478e-10 | 1 |
| 66 | 3.99995e-10 | 1 |
| 67 | 5.29961e-10 | 1 |
| 68 | 7.03249e-10 | 1 |
| 69 | 9.34628e-10 | 1 |
| 70 | 1.24316e-09 | 1 |
| 71 | 1.65581e-09 | 1 |
| 72 | 4.41701e-10 | 1 |
| 73 | 2.94946e-09 | 1 |
| 74 | 7.89288e-10 | 1 |
| 75 | 2.64599e-10 | 1 |
| 76 | 1.13865e-09 | 1 |
| 77 | 2.40175e-09 | 1 |
| 78 | 2.08565e-09 | 1 |
| 79 | 3.56362e-09 | 1 |
| 80 | 4.91977e-09 | 1 |
| 81 | 5.51094e-09 | 1 |
| 82 | 7.89166e-09 | 1 |
| 83 | 1.16843e-08 | 1 |
| 84 | 1.46518e-08 | 1 |
| 85 | 3.19315e-08 | 1 |
| 86 | 5.81253e-08 | 1 |
| 87 | 1.89855e-06 | 0.999998 |
| 88 | 0.016487 | 0.983513 |
| 89 | 0.0331688 | 0.966831 |
| 90 | 0.0494731 | 0.950527 |
| 91 | 0.0654073 | 0.934593 |
| 92 | 0.0809996 | 0.919 |
| 93 | 0.0962553 | 0.903745 |
| 94 | 0.111188 | 0.888812 |
| 95 | 0.125802 | 0.874198 |
| 96 | 0.140113 | 0.859887 |
| 97 | 0.154129 | 0.845871 |
| 98 | 0.167858 | 0.832142 |
| 99 | 0.181309 | 0.818691 |
| 100 | 0.19449 | 0.80551 |
| 101 | 0.207409 | 0.792591 |
| 102 | 0.220074 | 0.779926 |
| 103 | 0.232492 | 0.767508 |
| 104 | 0.244669 | 0.755331 |
| 105 | 0.256613 | 0.743387 |
| 106 | 0.268329 | 0.731671 |
| 107 | 0.279825 | 0.720175 |
| 108 | 0.291105 | 0.708895 |
| 109 | 0.302175 | 0.697825 |
| 110 | 0.31304 | 0.68696 |
| 111 | 0.323707 | 0.676293 |
| 112 | 0.334178 | 0.665822 |
| 113 | 0.344459 | 0.655541 |
| 114 | 0.354554 | 0.645446 |
| 115 | 0.364466 | 0.635534 |
| 116 | 0.374201 | 0.625799 |
| 117 | 0.38376 | 0.61624 |
| 118 | 0.393147 | 0.606853 |
| 119 | 0.402365 | 0.597635 |
| 120 | 0.411416 | 0.588584 |
| 121 | 0.420302 | 0.579698 |
| 122 | 0.429026 | 0.570974 |
| 123 | 0.437588 | 0.562412 |
| 124 | 0.44599 | 0.55401 |
| 125 | 0.454231 | 0.545769 |
| 126 | 0.462313 | 0.537687 |
| 127 | 0.470234 | 0.529766 |
| 128 | 0.477993 | 0.522007 |
| 129 | 0.48559 | 0.51441 |
| 130 | 0.493021 | 0.506979 |
| 131 | 0.500285 | 0.499715 |
| 132 | 0.507379 | 0.492621 |
| 133 | 0.514297 | 0.485703 |
| 134 | 0.521037 | 0.478963 |
| 135 | 0.527593 | 0.472407 |
| 136 | 0.533961 | 0.466039 |
| 137 | 0.540135 | 0.459865 |
| 138 | 0.54611 | 0.45389 |
| 139 | 0.55188 | 0.44812 |
| 140 | 0.557441 | 0.442559 |
| 141 | 0.562789 | 0.437211 |
| 142 | 0.567919 | 0.432081 |
| 143 | 0.572829 | 0.427171 |
| 144 | 0.577517 | 0.422483 |
| 145 | 0.581983 | 0.418017 |
| 146 | 0.586229 | 0.413771 |
| 147 | 0.590255 | 0.409745 |
| 148 | 0.594068 | 0.405932 |
| 149 | 0.597671 | 0.402329 |
| 150 | 0.601071 | 0.398929 |
| 151 | 0.604277 | 0.395723 |
| 152 | 0.607295 | 0.392705 |
| 153 | 0.610136 | 0.389864 |
| 154 | 0.612808 | 0.387192 |
| 155 | 0.615321 | 0.384679 |
| 156 | 0.617685 | 0.382315 |
| 157 | 0.61991 | 0.38009 |
| 158 | 0.622004 | 0.377996 |
| 159 | 0.623977 | 0.376023 |
| 160 | 0.625838 | 0.374162 |
| 161 | 0.627594 | 0.372406 |
| 162 | 0.629254 | 0.370746 |
| 163 | 0.630825 | 0.369175 |
| 164 | 0.632313 | 0.367687 |
| 165 | 0.633725 | 0.366275 |
| 166 | 0.635066 | 0.364934 |
| 167 | 0.636343 | 0.363657 |
| 168 | 0.637559 | 0.362441 |
| 169 | 0.63872 | 0.36128 |
| 170 | 0.639829 | 0.360171 |
| 171 | 0.640891 | 0.359109 |
| 172 | 0.641908 | 0.358092 |
| 173 | 0.642885 | 0.357115 |
| 174 | 0.643824 | 0.356176 |
| 175 | 0.644728 | 0.355272 |
| 176 | 0.6456 | 0.3544 |
| 177 | 0.646441 | 0.353559 |
| 178 | 0.647253 | 0.352747 |
| 179 | 0.64804 | 0.35196 |
| 180 | 0.648802 | 0.351198 |
| 181 | 0.649541 | 0.350459 |
| 182 | 0.650258 | 0.349742 |
| 183 | 0.650955 | 0.349045 |
| 184 | 0.651634 | 0.348366 |
| 185 | 0.652295 | 0.347705 |
| 186 | 0.652939 | 0.347061 |
| 187 | 0.653567 | 0.346433 |
| 188 | 0.654181 | 0.345819 |
| 189 | 0.654781 | 0.345219 |
| 190 | 0.655368 | 0.344632 |
| 191 | 0.655942 | 0.344058 |
| 192 | 0.656505 | 0.343495 |
| 193 | 0.657056 | 0.342944 |
| 194 | 0.657597 | 0.342403 |
| 195 | 0.658128 | 0.341872 |
| 196 | 0.658649 | 0.341351 |
| 197 | 0.659161 | 0.340839 |
| 198 | 0.659665 | 0.340335 |
| 199 | 0.66016 | 0.33984 |
| 200 | 0.660647 | 0.339353 |

V1 Equal

|  | Male | Female |
| --- | --- | --- |
| 1 | 1 | 0 |
| 2 | 1 | 0 |
| 3 | 1 | 0 |
| 4 | 1 | 0 |
| 5 | 1 | 0 |
| 6 | 0 | 1 |
| 7 | 0 | 1 |
| 8 | 0 | 1 |
| 9 | 0 | 1 |
| 10 | 0 | 1 |
| 11 | 0 | 1 |
| 12 | 0 | 1 |
| 13 | 0 | 1 |
| 14 | 0 | 1 |
| 15 | 0 | 1 |
| 16 | 0 | 1 |
| 17 | 0 | 1 |
| 18 | 1 | 1 |
| 19 | 1 | 1 |
| 20 | 1 | 1 |
| 21 | 0 | 2 |
| 22 | 0 | 2 |
| 23 | 0 | 2 |
| 24 | 0 | 2 |
| 25 | 0 | 2 |
| 26 | 0 | 2 |
| 27 | 0 | 2 |
| 28 | 0 | 2 |
| 29 | 0 | 2 |
| 30 | 0 | 2 |
| 31 | 0 | 2 |
| 32 | 0 | 3 |
| 33 | 0 | 3 |
| 34 | 0 | 3 |
| 35 | 0 | 3 |
| 36 | 0 | 3 |
| 37 | 0 | 3 |
| 38 | 0 | 3 |
| 39 | 0 | 3 |
| 40 | 0 | 3 |
| 41 | 0 | 3 |
| 42 | 0 | 3 |
| 43 | 0 | 4 |
| 44 | 0 | 4 |
| 45 | 0 | 4 |
| 46 | 0 | 4 |
| 47 | 0 | 4 |
| 48 | 0 | 4 |
| 49 | 0 | 4 |
| 50 | 0 | 4 |
| 51 | 0 | 4 |
| 52 | 0 | 4 |
| 53 | 0 | 4 |
| 54 | 0 | 4 |
| 55 | 0 | 5 |
| 56 | 0 | 5 |
| 57 | 0 | 5 |
| 58 | 0 | 5 |
| 59 | 0 | 5 |
| 60 | 0 | 5 |
| 61 | 0 | 5 |
| 62 | 0 | 5 |
| 63 | 0 | 5 |
| 64 | 0 | 5 |
| 65 | 0 | 5 |
| 66 | 0 | 5 |
| 67 | 0 | 5 |
| 68 | 0 | 5 |
| 69 | 0 | 5 |
| 70 | 0 | 5 |
| 71 | 0 | 5 |
| 72 | 0 | 5 |
| 73 | 0 | 5 |
| 74 | 0 | 5 |
| 75 | 0 | 5 |
| 76 | 0 | 5 |
| 77 | 0 | 5 |
| 78 | 0 | 5 |
| 79 | 0 | 5 |
| 80 | 0 | 5 |
| 81 | 0 | 5 |
| 82 | 0 | 5 |
| 83 | 0 | 5 |
| 84 | 0 | 5 |
| 85 | 0 | 5 |
| 86 | 0 | 5 |
| 87 | 0 | 5 |
| 88 | 1 | 5 |
| 89 | 1 | 5 |
| 90 | 1 | 5 |
| 91 | 1 | 5 |
| 92 | 1 | 5 |
| 93 | 1 | 5 |
| 94 | 1 | 5 |
| 95 | 1 | 5 |
| 96 | 1 | 5 |
| 97 | 1 | 5 |
| 98 | 1 | 5 |
| 99 | 1 | 5 |
| 100 | 1 | 5 |
| 101 | 1 | 5 |
| 102 | 1 | 5 |
| 103 | 1 | 5 |
| 104 | 1 | 5 |
| 105 | 1 | 5 |
| 106 | 1 | 5 |
| 107 | 1 | 5 |
| 108 | 1 | 5 |
| 109 | 1 | 5 |
| 110 | 1 | 5 |
| 111 | 1 | 5 |
| 112 | 1 | 5 |
| 113 | 1 | 5 |
| 114 | 1 | 5 |
| 115 | 1 | 5 |
| 116 | 1 | 5 |
| 117 | 1 | 5 |
| 118 | 1 | 5 |
| 119 | 1 | 5 |
| 120 | 1 | 5 |
| 121 | 1 | 5 |
| 122 | 1 | 5 |
| 123 | 1 | 5 |
| 124 | 1 | 5 |
| 125 | 1 | 5 |
| 126 | 1 | 5 |
| 127 | 1 | 5 |
| 128 | 1 | 5 |
| 129 | 1 | 5 |
| 130 | 1 | 5 |
| 131 | 1 | 5 |
| 132 | 1 | 5 |
| 133 | 1 | 5 |
| 134 | 1 | 5 |
| 135 | 1 | 5 |
| 136 | 1 | 5 |
| 137 | 1 | 5 |
| 138 | 1 | 5 |
| 139 | 1 | 5 |
| 140 | 1 | 5 |
| 141 | 1 | 5 |
| 142 | 1 | 5 |
| 143 | 1 | 5 |
| 144 | 1 | 5 |
| 145 | 1 | 5 |
| 146 | 1 | 5 |
| 147 | 1 | 5 |
| 148 | 1 | 5 |
| 149 | 1 | 5 |
| 150 | 1 | 5 |
| 151 | 1 | 5 |
| 152 | 1 | 5 |
| 153 | 1 | 5 |
| 154 | 1 | 5 |
| 155 | 1 | 5 |
| 156 | 1 | 5 |
| 157 | 1 | 5 |
| 158 | 1 | 5 |
| 159 | 1 | 5 |
| 160 | 1 | 5 |
| 161 | 1 | 5 |
| 162 | 1 | 5 |
| 163 | 1 | 5 |
| 164 | 1 | 5 |
| 165 | 1 | 5 |
| 166 | 1 | 5 |
| 167 | 1 | 5 |
| 168 | 1 | 5 |
| 169 | 1 | 5 |
| 170 | 1 | 5 |
| 171 | 1 | 5 |
| 172 | 1 | 5 |
| 173 | 1 | 5 |
| 174 | 1 | 5 |
| 175 | 1 | 5 |
| 176 | 1 | 5 |
| 177 | 1 | 5 |
| 178 | 1 | 5 |
| 179 | 1 | 5 |
| 180 | 1 | 5 |
| 181 | 1 | 5 |
| 182 | 1 | 5 |
| 183 | 1 | 5 |
| 184 | 1 | 5 |
| 185 | 1 | 5 |
| 186 | 1 | 5 |
| 187 | 1 | 5 |
| 188 | 1 | 5 |
| 189 | 1 | 5 |
| 190 | 1 | 5 |
| 191 | 1 | 5 |
| 192 | 1 | 5 |
| 193 | 1 | 5 |
| 194 | 1 | 5 |
| 195 | 1 | 5 |
| 196 | 1 | 5 |
| 197 | 1 | 5 |
| 198 | 1 | 5 |
| 199 | 1 | 5 |
| 200 | 1 | 5 |

V1 T-I-biased

|  | Male | Female |
| --- | --- | --- |
| 1 | 1 | 3.78359e-08 |
| 2 | 1 | 2.17687e-08 |
| 3 | 1 | 1.32798e-08 |
| 4 | 1 | 9.0819e-09 |
| 5 | 1 | 6.60982e-09 |
| 6 | 6.04574e-10 | 1 |
| 7 | 1.17034e-10 | 1 |
| 8 | 4.92071e-10 | 1 |
| 9 | 1.32455e-10 | 1 |
| 10 | 3.55579e-12 | 1 |
| 11 | 5.41862e-12 | 1 |
| 12 | 3.28047e-10 | 1 |
| 13 | 5.04162e-11 | 1 |
| 14 | 1.65844e-10 | 1 |
| 15 | 1.41414e-10 | 1 |
| 16 | 1.32446e-10 | 1 |
| 17 | 1.13267e-09 | 1 |
| 18 | 0.035334 | 0.964666 |
| 19 | 0.0898649 | 0.910135 |
| 20 | 0.138946 | 0.861054 |
| 21 | 1.74845e-12 | 1 |
| 22 | 2.43836e-12 | 1 |
| 23 | 8.06619e-13 | 1 |
| 24 | 1.47618e-12 | 1 |
| 25 | 2.85276e-12 | 1 |
| 26 | 1.13422e-12 | 1 |
| 27 | 4.57797e-11 | 1 |
| 28 | 1.49179e-11 | 1 |
| 29 | 3.06207e-11 | 1 |
| 30 | 1.58755e-09 | 1 |
| 31 | 2.68392e-09 | 1 |
| 32 | 1.27164e-12 | 1 |
| 33 | 1.62557e-12 | 1 |
| 34 | 8.96541e-12 | 1 |
| 35 | 6.52234e-13 | 1 |
| 36 | 9.84119e-13 | 1 |
| 37 | 1.52061e-12 | 1 |
| 38 | 4.77175e-11 | 1 |
| 39 | 3.78073e-12 | 1 |
| 40 | 9.6486e-11 | 1 |
| 41 | 7.73031e-12 | 1 |
| 42 | 6.21584e-11 | 1 |
| 43 | 5.02819e-12 | 1 |
| 44 | 1.21918e-12 | 1 |
| 45 | 1.54228e-12 | 1 |
| 46 | 2.01655e-12 | 1 |
| 47 | 2.16249e-12 | 1 |
| 48 | 1.47618e-11 | 1 |
| 49 | 2.04233e-11 | 1 |
| 50 | 2.85276e-11 | 1 |
| 51 | 4.01224e-11 | 1 |
| 52 | 5.67109e-11 | 1 |
| 53 | 1.60903e-11 | 1 |
| 54 | 9.15571e-11 | 1 |
| 55 | 9.75344e-11 | 1 |
| 56 | 2.34742e-11 | 1 |
| 57 | 2.88848e-11 | 1 |
| 58 | 1.80917e-10 | 1 |
| 59 | 4.59695e-11 | 1 |
| 60 | 5.90471e-11 | 1 |
| 61 | 7.64912e-11 | 1 |
| 62 | 1.99474e-11 | 1 |
| 63 | 5.22792e-10 | 1 |
| 64 | 1.37545e-10 | 1 |
| 65 | 3.62952e-11 | 1 |
| 66 | 2.39996e-10 | 1 |
| 67 | 9.46784e-11 | 1 |
| 68 | 2.24135e-09 | 1 |
| 69 | 5.37746e-10 | 1 |
| 70 | 1.30448e-10 | 1 |
| 71 | 1.59651e-10 | 1 |
| 72 | 4.92059e-11 | 1 |
| 73 | 1.22042e-09 | 1 |
| 74 | 3.04122e-10 | 1 |
| 75 | 3.80389e-10 | 1 |
| 76 | 1.19293e-10 | 1 |
| 77 | 2.40034e-11 | 1 |
| 78 | 1.89036e-10 | 1 |
| 79 | 1.52693e-10 | 1 |
| 80 | 8.51421e-11 | 1 |
| 81 | 5.00062e-10 | 1 |
| 82 | 2.95891e-11 | 1 |
| 83 | 3.52348e-09 | 1 |
| 84 | 1.68706e-10 | 1 |
| 85 | 1.01408e-09 | 1 |
| 86 | 1.22349e-09 | 1 |
| 87 | 1.18441e-09 | 1 |
| 88 | 8.97985e-11 | 1 |
| 89 | 8.73175e-11 | 1 |
| 90 | 1.32893e-10 | 1 |
| 91 | 3.24062e-09 | 1 |
| 92 | 6.3294e-10 | 1 |
| 93 | 4.83388e-11 | 1 |
| 94 | 2.36477e-10 | 1 |
| 95 | 2.89432e-10 | 1 |
| 96 | 4.43108e-10 | 1 |
| 97 | 8.68895e-11 | 1 |
| 98 | 5.32775e-10 | 1 |
| 99 | 6.5375e-10 | 1 |
| 100 | 4.01354e-11 | 1 |
| 101 | 7.89016e-10 | 1 |
| 102 | 1.2128e-09 | 1 |
| 103 | 2.9852e-10 | 1 |
| 104 | 3.67767e-10 | 1 |
| 105 | 1.8145e-09 | 1 |
| 106 | 5.60653e-10 | 1 |
| 107 | 5.54707e-10 | 1 |
| 108 | 6.87777e-10 | 1 |
| 109 | 2.67359e-10 | 1 |
| 110 | 1.06754e-09 | 1 |
| 111 | 1.33823e-09 | 1 |
| 112 | 1.68812e-09 | 1 |
| 113 | 4.28244e-10 | 1 |
| 114 | 6.86965e-10 | 1 |
| 115 | 3.56606e-09 | 1 |
| 116 | 3.77817e-09 | 1 |
| 117 | 6.41445e-09 | 1 |
| 118 | 7.23969e-09 | 1 |
| 119 | 1.36448e-08 | 1 |
| 120 | 1.87508e-08 | 1 |
| 121 | 4.51507e-08 | 1 |
| 122 | 0.00232907 | 0.997671 |
| 123 | 0.0176067 | 0.982393 |
| 124 | 0.0325891 | 0.967411 |
| 125 | 0.0473326 | 0.952667 |
| 126 | 0.061831 | 0.938169 |
| 127 | 0.0761047 | 0.923895 |
| 128 | 0.0901541 | 0.909846 |
| 129 | 0.103985 | 0.896015 |
| 130 | 0.117604 | 0.882396 |
| 131 | 0.13101 | 0.86899 |
| 132 | 0.144212 | 0.855788 |
| 133 | 0.157215 | 0.842785 |
| 134 | 0.170021 | 0.829979 |
| 135 | 0.182635 | 0.817365 |
| 136 | 0.19506 | 0.80494 |
| 137 | 0.2073 | 0.7927 |
| 138 | 0.219358 | 0.780642 |
| 139 | 0.231237 | 0.768763 |
| 140 | 0.24294 | 0.75706 |
| 141 | 0.254469 | 0.745531 |
| 142 | 0.265827 | 0.734173 |
| 143 | 0.277014 | 0.722986 |
| 144 | 0.288033 | 0.711967 |
| 145 | 0.298885 | 0.701115 |
| 146 | 0.309568 | 0.690432 |
| 147 | 0.320085 | 0.679915 |
| 148 | 0.330432 | 0.669568 |
| 149 | 0.34061 | 0.65939 |
| 150 | 0.350614 | 0.649386 |
| 151 | 0.360442 | 0.639558 |
| 152 | 0.37009 | 0.62991 |
| 153 | 0.37955 | 0.62045 |
| 154 | 0.388818 | 0.611182 |
| 155 | 0.397884 | 0.602116 |
| 156 | 0.406739 | 0.593261 |
| 157 | 0.415372 | 0.584628 |
| 158 | 0.423772 | 0.576228 |
| 159 | 0.431926 | 0.568074 |
| 160 | 0.43982 | 0.56018 |
| 161 | 0.447442 | 0.552558 |
| 162 | 0.454779 | 0.545221 |
| 163 | 0.461819 | 0.538181 |
| 164 | 0.46855 | 0.53145 |
| 165 | 0.474966 | 0.525034 |
| 166 | 0.48106 | 0.51894 |
| 167 | 0.48683 | 0.51317 |
| 168 | 0.492276 | 0.507724 |
| 169 | 0.497404 | 0.502596 |
| 170 | 0.502219 | 0.497781 |
| 171 | 0.506732 | 0.493268 |
| 172 | 0.510955 | 0.489045 |
| 173 | 0.514901 | 0.485099 |
| 174 | 0.518586 | 0.481414 |
| 175 | 0.522025 | 0.477975 |
| 176 | 0.525235 | 0.474765 |
| 177 | 0.528232 | 0.471768 |
| 178 | 0.531031 | 0.468969 |
| 179 | 0.533647 | 0.466353 |
| 180 | 0.536096 | 0.463904 |
| 181 | 0.53839 | 0.46161 |
| 182 | 0.540541 | 0.459459 |
| 183 | 0.542563 | 0.457437 |
| 184 | 0.544465 | 0.455535 |
| 185 | 0.546257 | 0.453743 |
| 186 | 0.547949 | 0.452051 |
| 187 | 0.549548 | 0.450452 |
| 188 | 0.551063 | 0.448937 |
| 189 | 0.5525 | 0.4475 |
| 190 | 0.553865 | 0.446135 |
| 191 | 0.555165 | 0.444835 |
| 192 | 0.556404 | 0.443596 |
| 193 | 0.557587 | 0.442413 |
| 194 | 0.558719 | 0.441281 |
| 195 | 0.559803 | 0.440197 |
| 196 | 0.560843 | 0.439157 |
| 197 | 0.561842 | 0.438158 |
| 198 | 0.562803 | 0.437197 |
| 199 | 0.563729 | 0.436271 |
| 200 | 0.564622 | 0.435378 |

V1 T-I-biased

|  | Male | Female |
| --- | --- | --- |
| 1 | 1 | 0 |
| 2 | 1 | 0 |
| 3 | 1 | 0 |
| 4 | 1 | 0 |
| 5 | 1 | 0 |
| 6 | 0 | 1 |
| 7 | 0 | 1 |
| 8 | 0 | 1 |
| 9 | 0 | 1 |
| 10 | 0 | 1 |
| 11 | 0 | 1 |
| 12 | 0 | 1 |
| 13 | 0 | 1 |
| 14 | 0 | 1 |
| 15 | 0 | 1 |
| 16 | 0 | 1 |
| 17 | 0 | 1 |
| 18 | 1 | 1 |
| 19 | 1 | 1 |
| 20 | 1 | 1 |
| 21 | 0 | 2 |
| 22 | 0 | 2 |
| 23 | 0 | 2 |
| 24 | 0 | 2 |
| 25 | 0 | 2 |
| 26 | 0 | 2 |
| 27 | 0 | 2 |
| 28 | 0 | 2 |
| 29 | 0 | 2 |
| 30 | 0 | 2 |
| 31 | 0 | 2 |
| 32 | 0 | 3 |
| 33 | 0 | 3 |
| 34 | 0 | 3 |
| 35 | 0 | 3 |
| 36 | 0 | 3 |
| 37 | 0 | 3 |
| 38 | 0 | 3 |
| 39 | 0 | 3 |
| 40 | 0 | 3 |
| 41 | 0 | 3 |
| 42 | 0 | 3 |
| 43 | 0 | 4 |
| 44 | 0 | 4 |
| 45 | 0 | 4 |
| 46 | 0 | 4 |
| 47 | 0 | 4 |
| 48 | 0 | 4 |
| 49 | 0 | 4 |
| 50 | 0 | 4 |
| 51 | 0 | 4 |
| 52 | 0 | 4 |
| 53 | 0 | 4 |
| 54 | 0 | 4 |
| 55 | 0 | 5 |
| 56 | 0 | 5 |
| 57 | 0 | 5 |
| 58 | 0 | 5 |
| 59 | 0 | 5 |
| 60 | 0 | 5 |
| 61 | 0 | 5 |
| 62 | 0 | 5 |
| 63 | 0 | 5 |
| 64 | 0 | 5 |
| 65 | 0 | 5 |
| 66 | 0 | 5 |
| 67 | 0 | 6 |
| 68 | 0 | 6 |
| 69 | 0 | 6 |
| 70 | 0 | 6 |
| 71 | 0 | 6 |
| 72 | 0 | 6 |
| 73 | 0 | 6 |
| 74 | 0 | 6 |
| 75 | 0 | 6 |
| 76 | 0 | 6 |
| 77 | 0 | 6 |
| 78 | 0 | 6 |
| 79 | 0 | 6 |
| 80 | 0 | 7 |
| 81 | 0 | 7 |
| 82 | 0 | 7 |
| 83 | 0 | 7 |
| 84 | 0 | 7 |
| 85 | 0 | 7 |
| 86 | 0 | 7 |
| 87 | 0 | 7 |
| 88 | 0 | 7 |
| 89 | 0 | 7 |
| 90 | 0 | 7 |
| 91 | 0 | 7 |
| 92 | 0 | 7 |
| 93 | 0 | 7 |
| 94 | 0 | 7 |
| 95 | 0 | 7 |
| 96 | 0 | 7 |
| 97 | 0 | 7 |
| 98 | 0 | 7 |
| 99 | 0 | 7 |
| 100 | 0 | 7 |
| 101 | 0 | 7 |
| 102 | 0 | 7 |
| 103 | 0 | 7 |
| 104 | 0 | 7 |
| 105 | 0 | 7 |
| 106 | 0 | 7 |
| 107 | 0 | 7 |
| 108 | 0 | 7 |
| 109 | 0 | 7 |
| 110 | 0 | 7 |
| 111 | 0 | 7 |
| 112 | 0 | 7 |
| 113 | 0 | 7 |
| 114 | 0 | 7 |
| 115 | 0 | 7 |
| 116 | 0 | 7 |
| 117 | 0 | 7 |
| 118 | 0 | 7 |
| 119 | 0 | 7 |
| 120 | 0 | 7 |
| 121 | 0 | 7 |
| 122 | 1 | 7 |
| 123 | 1 | 7 |
| 124 | 1 | 7 |
| 125 | 1 | 7 |
| 126 | 1 | 7 |
| 127 | 1 | 7 |
| 128 | 1 | 7 |
| 129 | 1 | 7 |
| 130 | 1 | 7 |
| 131 | 1 | 7 |
| 132 | 1 | 7 |
| 133 | 1 | 7 |
| 134 | 1 | 7 |
| 135 | 1 | 7 |
| 136 | 1 | 7 |
| 137 | 1 | 7 |
| 138 | 1 | 7 |
| 139 | 1 | 7 |
| 140 | 1 | 7 |
| 141 | 1 | 7 |
| 142 | 1 | 7 |
| 143 | 1 | 7 |
| 144 | 1 | 7 |
| 145 | 1 | 7 |
| 146 | 1 | 7 |
| 147 | 1 | 7 |
| 148 | 1 | 7 |
| 149 | 1 | 7 |
| 150 | 1 | 7 |
| 151 | 1 | 7 |
| 152 | 1 | 7 |
| 153 | 1 | 7 |
| 154 | 1 | 7 |
| 155 | 1 | 7 |
| 156 | 1 | 7 |
| 157 | 1 | 7 |
| 158 | 1 | 7 |
| 159 | 1 | 7 |
| 160 | 1 | 7 |
| 161 | 1 | 7 |
| 162 | 1 | 7 |
| 163 | 1 | 7 |
| 164 | 1 | 7 |
| 165 | 1 | 7 |
| 166 | 1 | 7 |
| 167 | 1 | 7 |
| 168 | 1 | 7 |
| 169 | 1 | 7 |
| 170 | 1 | 7 |
| 171 | 1 | 7 |
| 172 | 1 | 7 |
| 173 | 1 | 7 |
| 174 | 1 | 7 |
| 175 | 1 | 7 |
| 176 | 1 | 7 |
| 177 | 1 | 7 |
| 178 | 1 | 7 |
| 179 | 1 | 7 |
| 180 | 1 | 7 |
| 181 | 1 | 7 |
| 182 | 1 | 7 |
| 183 | 1 | 7 |
| 184 | 1 | 7 |
| 185 | 1 | 7 |
| 186 | 1 | 7 |
| 187 | 1 | 7 |
| 188 | 1 | 7 |
| 189 | 1 | 7 |
| 190 | 1 | 7 |
| 191 | 1 | 7 |
| 192 | 1 | 7 |
| 193 | 1 | 7 |
| 194 | 1 | 7 |
| 195 | 1 | 7 |
| 196 | 1 | 7 |
| 197 | 1 | 7 |
| 198 | 1 | 7 |
| 199 | 1 | 7 |
| 200 | 1 | 7 |

Figure S4 | Calculation results of model version 1.

(a) Fitness functions of type I (red) and type II (blue) offspring

(b) Optimized total offspring fitness of type-I-biased (yellow), equal (black), type-II-biased (pruple) brood.

(c) The ratio of total care given to type I offspring (red) and type II offspring (blue) in a type-II-biased brood.

(d) The number of cared type I offspring (red) and type II offspring (blue) in a type-II-biased brood.

(e, f) same as (c, d) for the equal brood.

(g, h) same as (c, d) for the type-I-biased brood.

**Model Version 2**

V2 Functions

|  |  |  |
| --- | --- | --- |
| 0 | -1.86711e-05 | 1.82578e-17 |
| 0.01 | 1.84368e-05 | 4.62312e-08 |
| 0.02 | 5.55816e-05 | 9.31611e-08 |
| 0.03 | 9.27634e-05 | 1.408e-07 |
| 0.04 | 0.000129982 | 1.89159e-07 |
| 0.05 | 0.000167238 | 2.38249e-07 |
| 0.06 | 0.000204531 | 2.88081e-07 |
| 0.07 | 0.000241861 | 3.38666e-07 |
| 0.08 | 0.000279228 | 3.90016e-07 |
| 0.09 | 0.000316633 | 4.42141e-07 |
| 0.1 | 0.000354074 | 4.95055e-07 |
| 0.11 | 0.000391553 | 5.48768e-07 |
| 0.12 | 0.000429069 | 6.03292e-07 |
| 0.13 | 0.000466623 | 6.58641e-07 |
| 0.14 | 0.000504214 | 7.14826e-07 |
| 0.15 | 0.000541842 | 7.71861e-07 |
| 0.16 | 0.000579508 | 8.29757e-07 |
| 0.17 | 0.000617212 | 8.88528e-07 |
| 0.18 | 0.000654952 | 9.48188e-07 |
| 0.19 | 0.000692731 | 1.00875e-06 |
| 0.2 | 0.000730547 | 1.07023e-06 |
| 0.21 | 0.000768401 | 1.13263e-06 |
| 0.22 | 0.000806292 | 1.19598e-06 |
| 0.23 | 0.000844221 | 1.26029e-06 |
| 0.24 | 0.000882188 | 1.32556e-06 |
| 0.25 | 0.000920193 | 1.39183e-06 |
| 0.26 | 0.000958235 | 1.45909e-06 |
| 0.27 | 0.000996315 | 1.52738e-06 |
| 0.28 | 0.00103443 | 1.59669e-06 |
| 0.29 | 0.00107259 | 1.66705e-06 |
| 0.3 | 0.00111078 | 1.73848e-06 |
| 0.31 | 0.00114902 | 1.81098e-06 |
| 0.32 | 0.00118729 | 1.88458e-06 |
| 0.33 | 0.00122559 | 1.9593e-06 |
| 0.34 | 0.00126394 | 2.03514e-06 |
| 0.35 | 0.00130233 | 2.11213e-06 |
| 0.36 | 0.00134075 | 2.19028e-06 |
| 0.37 | 0.00137921 | 2.26961e-06 |
| 0.38 | 0.00141771 | 2.35015e-06 |
| 0.39 | 0.00145625 | 2.43189e-06 |
| 0.4 | 0.00149482 | 2.51488e-06 |
| 0.41 | 0.00153344 | 2.59912e-06 |
| 0.42 | 0.00157209 | 2.68463e-06 |
| 0.43 | 0.00161078 | 2.77143e-06 |
| 0.44 | 0.00164951 | 2.85955e-06 |
| 0.45 | 0.00168828 | 2.949e-06 |
| 0.46 | 0.00172709 | 3.0398e-06 |
| 0.47 | 0.00176593 | 3.13197e-06 |
| 0.48 | 0.00180482 | 3.22553e-06 |
| 0.49 | 0.00184374 | 3.32051e-06 |
| 0.5 | 0.0018827 | 3.41693e-06 |
| 0.51 | 0.0019217 | 3.5148e-06 |
| 0.52 | 0.00196074 | 3.61415e-06 |
| 0.53 | 0.00199982 | 3.715e-06 |
| 0.54 | 0.00203894 | 3.81738e-06 |
| 0.55 | 0.00207809 | 3.9213e-06 |
| 0.56 | 0.00211729 | 4.02679e-06 |
| 0.57 | 0.00215652 | 4.13388e-06 |
| 0.58 | 0.00219579 | 4.24259e-06 |
| 0.59 | 0.00223511 | 4.35294e-06 |
| 0.6 | 0.00227446 | 4.46496e-06 |
| 0.61 | 0.00231385 | 4.57867e-06 |
| 0.62 | 0.00235328 | 4.6941e-06 |
| 0.63 | 0.00239275 | 4.81127e-06 |
| 0.64 | 0.00243226 | 4.93021e-06 |
| 0.65 | 0.0024718 | 5.05095e-06 |
| 0.66 | 0.00251139 | 5.17352e-06 |
| 0.67 | 0.00255102 | 5.29794e-06 |
| 0.68 | 0.00259068 | 5.42424e-06 |
| 0.69 | 0.00263039 | 5.55245e-06 |
| 0.7 | 0.00267013 | 5.68259e-06 |
| 0.71 | 0.00270992 | 5.81471e-06 |
| 0.72 | 0.00274974 | 5.94881e-06 |
| 0.73 | 0.0027896 | 6.08495e-06 |
| 0.74 | 0.00282951 | 6.22314e-06 |
| 0.75 | 0.00286945 | 6.36343e-06 |
| 0.76 | 0.00290943 | 6.50583e-06 |
| 0.77 | 0.00294945 | 6.65038e-06 |
| 0.78 | 0.00298952 | 6.79712e-06 |
| 0.79 | 0.00302962 | 6.94608e-06 |
| 0.8 | 0.00306976 | 7.09728e-06 |
| 0.81 | 0.00310994 | 7.25078e-06 |
| 0.82 | 0.00315016 | 7.40659e-06 |
| 0.83 | 0.00319043 | 7.56476e-06 |
| 0.84 | 0.00323073 | 7.72531e-06 |
| 0.85 | 0.00327107 | 7.8883e-06 |
| 0.86 | 0.00331145 | 8.05375e-06 |
| 0.87 | 0.00335187 | 8.22169e-06 |
| 0.88 | 0.00339234 | 8.39218e-06 |
| 0.89 | 0.00343284 | 8.56524e-06 |
| 0.9 | 0.00347338 | 8.74092e-06 |
| 0.91 | 0.00351397 | 8.91925e-06 |
| 0.92 | 0.00355459 | 9.10028e-06 |
| 0.93 | 0.00359525 | 9.28405e-06 |
| 0.94 | 0.00363596 | 9.47059e-06 |
| 0.95 | 0.0036767 | 9.65995e-06 |
| 0.96 | 0.00371749 | 9.85217e-06 |
| 0.97 | 0.00375832 | 1.00473e-05 |
| 0.98 | 0.00379918 | 1.02454e-05 |
| 0.99 | 0.00384009 | 1.04464e-05 |
| 1 | 0.00388104 | 1.06506e-05 |
| 1.01 | 0.00392203 | 1.08577e-05 |
| 1.02 | 0.00396306 | 1.10681e-05 |
| 1.03 | 0.00400413 | 1.12816e-05 |
| 1.04 | 0.00404524 | 1.14983e-05 |
| 1.05 | 0.00408639 | 1.17183e-05 |
| 1.06 | 0.00412759 | 1.19416e-05 |
| 1.07 | 0.00416882 | 1.21683e-05 |
| 1.08 | 0.0042101 | 1.23985e-05 |
| 1.09 | 0.00425141 | 1.26321e-05 |
| 1.1 | 0.00429277 | 1.28692e-05 |
| 1.11 | 0.00433417 | 1.311e-05 |
| 1.12 | 0.00437561 | 1.33543e-05 |
| 1.13 | 0.00441709 | 1.36024e-05 |
| 1.14 | 0.00445862 | 1.38542e-05 |
| 1.15 | 0.00450018 | 1.41098e-05 |
| 1.16 | 0.00454178 | 1.43693e-05 |
| 1.17 | 0.00458343 | 1.46327e-05 |
| 1.18 | 0.00462512 | 1.49e-05 |
| 1.19 | 0.00466685 | 1.51714e-05 |
| 1.2 | 0.00470862 | 1.5447e-05 |
| 1.21 | 0.00475043 | 1.57266e-05 |
| 1.22 | 0.00479229 | 1.60106e-05 |
| 1.23 | 0.00483418 | 1.62987e-05 |
| 1.24 | 0.00487612 | 1.65913e-05 |
| 1.25 | 0.0049181 | 1.68883e-05 |
| 1.26 | 0.00496012 | 1.71897e-05 |
| 1.27 | 0.00500218 | 1.74958e-05 |
| 1.28 | 0.00504429 | 1.78064e-05 |
| 1.29 | 0.00508643 | 1.81217e-05 |
| 1.3 | 0.00512862 | 1.84419e-05 |
| 1.31 | 0.00517085 | 1.87668e-05 |
| 1.32 | 0.00521313 | 1.90967e-05 |
| 1.33 | 0.00525544 | 1.94315e-05 |
| 1.34 | 0.0052978 | 1.97714e-05 |
| 1.35 | 0.0053402 | 2.01164e-05 |
| 1.36 | 0.00538264 | 2.04667e-05 |
| 1.37 | 0.00542512 | 2.08222e-05 |
| 1.38 | 0.00546764 | 2.11831e-05 |
| 1.39 | 0.00551021 | 2.15495e-05 |
| 1.4 | 0.00555282 | 2.19214e-05 |
| 1.41 | 0.00559548 | 2.2299e-05 |
| 1.42 | 0.00563817 | 2.26822e-05 |
| 1.43 | 0.00568091 | 2.30712e-05 |
| 1.44 | 0.00572369 | 2.34661e-05 |
| 1.45 | 0.00576651 | 2.3867e-05 |
| 1.46 | 0.00580937 | 2.42739e-05 |
| 1.47 | 0.00585228 | 2.4687e-05 |
| 1.48 | 0.00589523 | 2.51063e-05 |
| 1.49 | 0.00593823 | 2.5532e-05 |
| 1.5 | 0.00598126 | 2.59641e-05 |
| 1.51 | 0.00602434 | 2.64027e-05 |
| 1.52 | 0.00606746 | 2.6848e-05 |
| 1.53 | 0.00611063 | 2.73e-05 |
| 1.54 | 0.00615383 | 2.77588e-05 |
| 1.55 | 0.00619708 | 2.82245e-05 |
| 1.56 | 0.00624038 | 2.86973e-05 |
| 1.57 | 0.00628371 | 2.91773e-05 |
| 1.58 | 0.00632709 | 2.96645e-05 |
| 1.59 | 0.00637052 | 3.0159e-05 |
| 1.6 | 0.00641398 | 3.0661e-05 |
| 1.61 | 0.00645749 | 3.11706e-05 |
| 1.62 | 0.00650104 | 3.1688e-05 |
| 1.63 | 0.00654464 | 3.22131e-05 |
| 1.64 | 0.00658828 | 3.27461e-05 |
| 1.65 | 0.00663196 | 3.32873e-05 |
| 1.66 | 0.00667569 | 3.38366e-05 |
| 1.67 | 0.00671946 | 3.43942e-05 |
| 1.68 | 0.00676327 | 3.49602e-05 |
| 1.69 | 0.00680713 | 3.55348e-05 |
| 1.7 | 0.00685103 | 3.61181e-05 |
| 1.71 | 0.00689497 | 3.67101e-05 |
| 1.72 | 0.00693896 | 3.73112e-05 |
| 1.73 | 0.00698299 | 3.79213e-05 |
| 1.74 | 0.00702706 | 3.85406e-05 |
| 1.75 | 0.00707118 | 3.91693e-05 |
| 1.76 | 0.00711535 | 3.98075e-05 |
| 1.77 | 0.00715955 | 4.04554e-05 |
| 1.78 | 0.0072038 | 4.1113e-05 |
| 1.79 | 0.0072481 | 4.17806e-05 |
| 1.8 | 0.00729244 | 4.24582e-05 |
| 1.81 | 0.00733682 | 4.31461e-05 |
| 1.82 | 0.00738125 | 4.38444e-05 |
| 1.83 | 0.00742572 | 4.45533e-05 |
| 1.84 | 0.00747023 | 4.52728e-05 |
| 1.85 | 0.00751479 | 4.60033e-05 |
| 1.86 | 0.0075594 | 4.67448e-05 |
| 1.87 | 0.00760404 | 4.74974e-05 |
| 1.88 | 0.00764874 | 4.82615e-05 |
| 1.89 | 0.00769347 | 4.90371e-05 |
| 1.9 | 0.00773826 | 4.98244e-05 |
| 1.91 | 0.00778308 | 5.06237e-05 |
| 1.92 | 0.00782795 | 5.1435e-05 |
| 1.93 | 0.00787287 | 5.22585e-05 |
| 1.94 | 0.00791783 | 5.30946e-05 |
| 1.95 | 0.00796283 | 5.39432e-05 |
| 1.96 | 0.00800788 | 5.48047e-05 |
| 1.97 | 0.00805298 | 5.56792e-05 |
| 1.98 | 0.00809812 | 5.65669e-05 |
| 1.99 | 0.0081433 | 5.7468e-05 |

V2 Total fitness

|  |  |  |  |
| --- | --- | --- | --- |
| 1 | 0.0038 | 0.00380636 | 0.0038437 |
| 2 | 0.0081 | 0.00811385 | 0.00815119 |
| 3 | 0.0128 | 0.0128715 | 0.0129088 |
| 4 | 0.0181 | 0.018126 | 0.0181633 |
| 5 | 0.0239 | 0.0239288 | 0.0239661 |
| 6 | 0.0303 | 0.0303367 | 0.030374 |
| 7 | 0.0374 | 0.0374121 | 0.0374494 |
| 8 | 0.0452 | 0.0452239 | 0.474199 |
| 9 | 1.8241 | 1.82416 | 1.8242 |
| 10 | 4.9999 | 4.99991 | 4.99994 |
| 11 | 8.1756 | 8.17565 | 8.17569 |
| 12 | 9.5256 | 9.52565 | 9.52569 |
| 13 | 9.89 | 9.89004 | 9.89007 |
| 14 | 9.9751 | 9.97518 | 9.97522 |
| 15 | 9.9943 | 9.99438 | 9.99442 |
| 16 | 9.9992 | 9.9992 | 9.99923 |
| 17 | 10.0033 | 10.0033 | 10.0033 |
| 18 | 10.0078 | 10.0079 | 10.0079 |
| 19 | 10.0129 | 10.013 | 10.013 |
| 20 | 10.0186 | 10.0186 | 10.0187 |
| 21 | 13.5834 | 13.5835 | 13.5835 |
| 22 | 16.3514 | 16.3514 | 16.3514 |
| 23 | 18.0929 | 18.0929 | 18.093 |
| 24 | 19.0514 | 19.0514 | 19.0514 |
| 25 | 19.5403 | 19.5404 | 19.5404 |
| 26 | 19.7801 | 19.7802 | 19.7802 |
| 27 | 19.8955 | 19.8955 | 19.8955 |
| 28 | 19.9504 | 19.9505 | 19.9505 |
| 29 | 19.9765 | 19.9765 | 19.9766 |
| 30 | 19.9888 | 19.9889 | 19.9889 |
| 31 | 19.9946 | 19.9947 | 19.9947 |
| 32 | 21.9316 | 21.9317 | 21.9317 |
| 33 | 24.5271 | 24.5271 | 24.5272 |
| 34 | 26.4238 | 26.4238 | 26.4239 |
| 35 | 27.7241 | 27.7242 | 27.7242 |
| 36 | 28.5771 | 28.5771 | 28.5772 |
| 37 | 29.1205 | 29.1205 | 29.1206 |
| 38 | 29.4603 | 29.4603 | 29.4604 |
| 39 | 29.6703 | 29.6703 | 29.6703 |
| 40 | 29.7991 | 29.7991 | 29.7992 |
| 41 | 29.8778 | 29.8778 | 29.8778 |
| 42 | 29.9257 | 29.9257 | 29.9258 |
| 43 | 29.9548 | 30.1965 | 30.1965 |
| 44 | 29.9725 | 32.7029 | 32.7029 |
| 45 | 29.9833 | 34.6813 | 34.6814 |
| 46 | 29.9898 | 36.1859 | 36.186 |
| 47 | 29.9939 | 37.298 | 37.2981 |
| 48 | 29.9981 | 38.1029 | 38.1029 |
| 49 | 30.0028 | 38.6765 | 38.6765 |
| 50 | 30.0081 | 39.0808 | 39.0808 |
| 51 | 30.0141 | 39.3637 | 39.3637 |
| 52 | 30.0209 | 39.5604 | 39.5605 |
| 53 | 30.0286 | 39.6968 | 39.6969 |
| 54 | 30.0373 | 39.7911 | 39.7911 |
| 55 | 30.0472 | 40.8786 | 40.8787 |
| 56 | 30.0583 | 42.9074 | 42.9074 |
| 57 | 30.0709 | 44.5451 | 44.5451 |
| 58 | 30.0851 | 45.8413 | 45.8413 |
| 59 | 30.1012 | 46.8512 | 46.8513 |
| 60 | 30.1193 | 47.6286 | 47.6286 |
| 61 | 30.1398 | 48.2213 | 48.2214 |
| 62 | 30.1629 | 48.6701 | 48.6701 |
| 63 | 30.189 | 49.0079 | 49.0079 |
| 64 | 30.2183 | 49.2612 | 49.2612 |
| 65 | 30.2514 | 49.4506 | 49.4506 |
| 66 | 30.2886 | 49.5918 | 49.5918 |
| 67 | 30.3304 | 49.6969 | 49.697 |
| 68 | 30.3773 | 49.7751 | 52.8478 |
| 69 | 30.4299 | 49.8332 | 54.279 |
| 70 | 30.4888 | 49.8763 | 55.4485 |
| 71 | 30.5547 | 49.9083 | 56.3947 |
| 72 | 30.6283 | 49.932 | 57.1544 |
| 73 | 30.7104 | 49.9496 | 57.7603 |
| 74 | 30.8017 | 49.9626 | 58.2412 |
| 75 | 30.903 | 49.9723 | 58.6213 |
| 76 | 31.0152 | 49.9794 | 58.9208 |
| 77 | 31.1392 | 49.9847 | 59.1561 |
| 78 | 31.2758 | 49.9887 | 59.3407 |
| 79 | 31.4258 | 49.9929 | 59.4853 |
| 80 | 31.5899 | 49.9978 | 59.5984 |
| 81 | 31.769 | 50.0035 | 59.6867 |
| 82 | 31.9635 | 50.01 | 59.7557 |
| 83 | 32.1741 | 50.0176 | 65.9329 |
| 84 | 32.4009 | 50.0264 | 66.6801 |
| 85 | 32.6442 | 50.0366 | 67.2957 |
| 86 | 32.9039 | 50.0484 | 67.8009 |
| 87 | 33.1797 | 50.062 | 68.2143 |
| 88 | 33.4712 | 50.0779 | 68.5516 |
| 89 | 33.7776 | 50.0961 | 68.8262 |
| 90 | 34.0979 | 50.1172 | 69.0496 |
| 91 | 34.431 | 50.1416 | 69.2309 |
| 92 | 34.7754 | 50.1697 | 69.3779 |
| 93 | 35.1296 | 50.2021 | 69.497 |
| 94 | 35.4919 | 50.2393 | 69.5934 |
| 95 | 35.8605 | 50.282 | 69.6715 |
| 96 | 36.2333 | 50.331 | 69.7346 |
| 97 | 36.6086 | 50.3871 | 69.7856 |
| 98 | 36.9844 | 50.4511 | 69.8269 |
| 99 | 37.3589 | 50.524 | 69.8602 |
| 100 | 37.7302 | 50.6067 | 69.8871 |
| 101 | 38.0967 | 50.7005 | 69.9088 |
| 102 | 38.4569 | 50.8062 | 69.9264 |
| 103 | 38.8094 | 50.925 | 69.9406 |
| 104 | 39.153 | 51.058 | 69.952 |
| 105 | 39.4866 | 51.2062 | 69.9612 |
| 106 | 39.8093 | 51.3706 | 69.9687 |
| 107 | 40.1204 | 51.5518 | 69.9747 |
| 108 | 40.4194 | 51.7507 | 69.9796 |
| 109 | 40.7059 | 51.9675 | 69.9836 |
| 110 | 40.9796 | 52.2026 | 69.9878 |
| 111 | 41.2404 | 52.4558 | 69.993 |
| 112 | 41.4882 | 52.7268 | 69.9991 |
| 113 | 41.7232 | 53.0149 | 70.0066 |
| 114 | 41.9455 | 53.3193 | 70.0155 |
| 115 | 42.1555 | 53.6386 | 70.0262 |
| 116 | 42.3533 | 53.9715 | 70.0391 |
| 117 | 42.5393 | 54.3163 | 70.0545 |
| 118 | 42.7141 | 54.6711 | 70.073 |
| 119 | 42.8779 | 55.0342 | 70.0951 |
| 120 | 43.0314 | 55.4034 | 70.1214 |
| 121 | 43.1748 | 55.7767 | 70.1528 |
| 122 | 43.3088 | 56.1522 | 70.19 |
| 123 | 43.4338 | 56.5279 | 70.2342 |
| 124 | 43.5503 | 56.902 | 70.2864 |
| 125 | 43.6587 | 57.2728 | 70.3477 |
| 126 | 43.7595 | 57.6386 | 70.4196 |
| 127 | 43.8532 | 57.9982 | 70.5034 |
| 128 | 43.9403 | 58.3502 | 70.6006 |
| 129 | 44.021 | 58.6935 | 70.7126 |
| 130 | 44.0959 | 59.0272 | 70.8409 |
| 131 | 44.1653 | 59.3505 | 70.9868 |
| 132 | 44.2296 | 59.6629 | 71.1513 |
| 133 | 44.2891 | 59.9639 | 71.3355 |
| 134 | 44.3442 | 60.253 | 71.5398 |
| 135 | 44.3951 | 60.5302 | 71.7647 |
| 136 | 44.4422 | 60.7953 | 72.01 |
| 137 | 44.4857 | 61.0483 | 72.2751 |
| 138 | 44.5258 | 61.2892 | 72.5592 |
| 139 | 44.563 | 61.5183 | 72.8612 |
| 140 | 44.5972 | 61.7357 | 73.1793 |
| 141 | 44.6289 | 61.9417 | 73.512 |
| 142 | 44.658 | 62.1366 | 73.8571 |
| 143 | 44.685 | 62.3208 | 74.2127 |
| 144 | 44.7098 | 62.4945 | 74.5765 |
| 145 | 44.7327 | 62.6583 | 74.9463 |
| 146 | 44.7538 | 62.8125 | 75.3201 |
| 147 | 44.7733 | 62.9575 | 75.6957 |
| 148 | 44.7912 | 63.0938 | 76.0713 |
| 149 | 44.8078 | 63.2217 | 76.4451 |
| 150 | 44.823 | 63.3416 | 76.8153 |
| 151 | 44.837 | 63.454 | 77.1805 |
| 152 | 44.85 | 63.5593 | 77.5394 |
| 153 | 44.8619 | 63.6579 | 77.8909 |
| 154 | 44.8728 | 63.75 | 78.234 |
| 155 | 44.8829 | 63.8362 | 78.5678 |
| 156 | 44.8922 | 63.9167 | 78.8917 |
| 157 | 44.9008 | 63.9918 | 79.2052 |
| 158 | 44.9087 | 64.062 | 79.5079 |
| 159 | 44.916 | 64.1274 | 79.7995 |
| 160 | 45.2162 | 64.1884 | 80.0797 |
| 161 | 45.5897 | 64.2453 | 80.3486 |
| 162 | 45.9615 | 64.2984 | 80.6061 |
| 163 | 46.3311 | 64.3478 | 80.8522 |
| 164 | 46.6982 | 64.3938 | 81.0872 |
| 165 | 47.0623 | 64.4366 | 81.3112 |
| 166 | 47.4231 | 64.4765 | 81.5244 |
| 167 | 47.7801 | 64.5136 | 81.727 |
| 168 | 48.133 | 64.5481 | 81.9195 |
| 169 | 48.4816 | 64.5802 | 82.102 |
| 170 | 48.8254 | 64.6101 | 82.275 |
| 171 | 49.1641 | 64.6379 | 82.4387 |
| 172 | 49.4976 | 64.6637 | 82.5935 |
| 173 | 49.8256 | 64.6877 | 82.7399 |
| 174 | 50.1478 | 64.71 | 82.878 |
| 175 | 50.4641 | 64.7307 | 83.0084 |
| 176 | 50.7742 | 64.75 | 83.1314 |
| 177 | 51.078 | 64.7679 | 83.2472 |
| 178 | 51.3755 | 64.7845 | 83.3564 |
| 179 | 51.6663 | 64.7999 | 83.4591 |
| 180 | 51.9506 | 64.8143 | 83.5557 |
| 181 | 52.2281 | 64.8276 | 83.6466 |
| 182 | 52.4989 | 64.84 | 83.732 |
| 183 | 52.7628 | 64.8515 | 83.8123 |
| 184 | 53.02 | 64.8621 | 83.8877 |
| 185 | 53.2703 | 64.8721 | 83.9585 |
| 186 | 53.5138 | 65.1328 | 84.0249 |
| 187 | 53.7505 | 65.5043 | 84.0872 |
| 188 | 53.9804 | 65.8736 | 84.1457 |
| 189 | 54.2036 | 66.2404 | 84.2005 |
| 190 | 54.4202 | 66.6042 | 84.2519 |
| 191 | 54.6303 | 66.9646 | 84.3001 |
| 192 | 54.8338 | 67.3213 | 84.3452 |
| 193 | 55.031 | 67.6739 | 84.3875 |
| 194 | 55.2218 | 68.0222 | 84.4271 |
| 195 | 55.4065 | 68.3659 | 84.4642 |
| 196 | 55.585 | 68.7046 | 84.4989 |
| 197 | 55.7577 | 69.0382 | 84.5314 |
| 198 | 55.9244 | 69.3664 | 84.5619 |
| 199 | 56.0855 | 69.6889 | 84.5904 |
| 200 | 56.241 | 70.0057 | 84.617 |

V2 T-II-biased

|  | Male | Female |
| --- | --- | --- |
| 1 | 1 | 0 |
| 2 | 1 | 0 |
| 3 | 1 | 0 |
| 4 | 1 | 0 |
| 5 | 1 | 0 |
| 6 | 1 | 0 |
| 7 | 1 | 0 |
| 8 | 1 | 0 |
| 9 | 0 | 1 |
| 10 | 0 | 1 |
| 11 | 0 | 1 |
| 12 | 0 | 1 |
| 13 | 0 | 1 |
| 14 | 0 | 1 |
| 15 | 0 | 1 |
| 16 | 0.0310375 | 0.968962 |
| 17 | 0.0922118 | 0.907788 |
| 18 | 0.146589 | 0.853411 |
| 19 | 0.195242 | 0.804758 |
| 20 | 0.239025 | 0.760975 |
| 21 | 0 | 1 |
| 22 | 0 | 1 |
| 23 | 0 | 1 |
| 24 | 0 | 1 |
| 25 | 0 | 1 |
| 26 | 0 | 1 |
| 27 | 0 | 1 |
| 28 | 0 | 1 |
| 29 | 0 | 1 |
| 30 | 0 | 1 |
| 31 | 0 | 1 |
| 32 | 0 | 1 |
| 33 | 0 | 1 |
| 34 | 0 | 0.999997 |
| 35 | 0 | 1 |
| 36 | 0 | 1 |
| 37 | 0 | 0.999997 |
| 38 | 0 | 1 |
| 39 | 0 | 1 |
| 40 | 0 | 0.999997 |
| 41 | 0 | 1 |
| 42 | 0 | 1 |
| 43 | 0 | 0.999998 |
| 44 | 0 | 1 |
| 45 | 0 | 1 |
| 46 | 0 | 0.999998 |
| 47 | 0.010383 | 0.989617 |
| 48 | 0.036175 | 0.963825 |
| 49 | 0.0609122 | 0.939086 |
| 50 | 0.084654 | 0.915348 |
| 51 | 0.107461 | 0.892541 |
| 52 | 0.129383 | 0.870617 |
| 53 | 0.150472 | 0.849526 |
| 54 | 0.170772 | 0.829228 |
| 55 | 0.190327 | 0.809673 |
| 56 | 0.209175 | 0.790825 |
| 57 | 0.227353 | 0.772647 |
| 58 | 0.244891 | 0.75511 |
| 59 | 0.261825 | 0.738173 |
| 60 | 0.27818 | 0.72182 |
| 61 | 0.293985 | 0.706013 |
| 62 | 0.309263 | 0.690735 |
| 63 | 0.324038 | 0.675962 |
| 64 | 0.338331 | 0.661669 |
| 65 | 0.352163 | 0.647838 |
| 66 | 0.365552 | 0.63445 |
| 67 | 0.378513 | 0.621488 |
| 68 | 0.391065 | 0.608934 |
| 69 | 0.403219 | 0.596783 |
| 70 | 0.414989 | 0.585013 |
| 71 | 0.426389 | 0.573613 |
| 72 | 0.437428 | 0.562571 |
| 73 | 0.448116 | 0.551885 |
| 74 | 0.458465 | 0.541536 |
| 75 | 0.46848 | 0.53152 |
| 76 | 0.47817 | 0.52183 |
| 77 | 0.487542 | 0.512458 |
| 78 | 0.4966 | 0.5034 |
| 79 | 0.505352 | 0.494647 |
| 80 | 0.513802 | 0.486199 |
| 81 | 0.521954 | 0.478044 |
| 82 | 0.529813 | 0.470188 |
| 83 | 0.537383 | 0.462618 |
| 84 | 0.544669 | 0.455332 |
| 85 | 0.551673 | 0.448327 |
| 86 | 0.558401 | 0.4416 |
| 87 | 0.564856 | 0.435145 |
| 88 | 0.571045 | 0.428956 |
| 89 | 0.576973 | 0.423027 |
| 90 | 0.582644 | 0.417357 |
| 91 | 0.588067 | 0.411933 |
| 92 | 0.593247 | 0.406754 |
| 93 | 0.598191 | 0.40181 |
| 94 | 0.602909 | 0.397091 |
| 95 | 0.607407 | 0.392593 |
| 96 | 0.611696 | 0.388303 |
| 97 | 0.615785 | 0.384216 |
| 98 | 0.61968 | 0.38032 |
| 99 | 0.623393 | 0.376606 |
| 100 | 0.626933 | 0.373068 |
| 101 | 0.630309 | 0.369692 |
| 102 | 0.63353 | 0.366471 |
| 103 | 0.636605 | 0.363396 |
| 104 | 0.639541 | 0.360459 |
| 105 | 0.64235 | 0.357651 |
| 106 | 0.645035 | 0.354965 |
| 107 | 0.647607 | 0.352393 |
| 108 | 0.650072 | 0.349928 |
| 109 | 0.652438 | 0.347562 |
| 110 | 0.654709 | 0.345292 |
| 111 | 0.656892 | 0.343108 |
| 112 | 0.658994 | 0.341006 |
| 113 | 0.661018 | 0.338981 |
| 114 | 0.66297 | 0.337029 |
| 115 | 0.664854 | 0.335147 |
| 116 | 0.666675 | 0.333326 |
| 117 | 0.668437 | 0.331564 |
| 118 | 0.670142 | 0.329858 |
| 119 | 0.671795 | 0.328205 |
| 120 | 0.673398 | 0.326603 |
| 121 | 0.674955 | 0.325044 |
| 122 | 0.676469 | 0.32353 |
| 123 | 0.677941 | 0.322059 |
| 124 | 0.679373 | 0.320627 |
| 125 | 0.680769 | 0.319231 |
| 126 | 0.682129 | 0.317871 |
| 127 | 0.683457 | 0.316543 |
| 128 | 0.684752 | 0.315248 |
| 129 | 0.686019 | 0.313981 |
| 130 | 0.687256 | 0.312743 |
| 131 | 0.688466 | 0.311533 |
| 132 | 0.689651 | 0.31035 |
| 133 | 0.690811 | 0.309189 |
| 134 | 0.691947 | 0.308053 |
| 135 | 0.693061 | 0.30694 |
| 136 | 0.694153 | 0.305848 |
| 137 | 0.695224 | 0.304776 |
| 138 | 0.696275 | 0.303724 |
| 139 | 0.697308 | 0.302691 |
| 140 | 0.698321 | 0.301678 |
| 141 | 0.699318 | 0.300683 |
| 142 | 0.700296 | 0.299704 |
| 143 | 0.701259 | 0.298741 |
| 144 | 0.702206 | 0.297794 |
| 145 | 0.703137 | 0.296863 |
| 146 | 0.704053 | 0.295946 |
| 147 | 0.704955 | 0.295045 |
| 148 | 0.705843 | 0.294156 |
| 149 | 0.706717 | 0.293283 |
| 150 | 0.707579 | 0.29242 |
| 151 | 0.708428 | 0.291572 |
| 152 | 0.709264 | 0.290736 |
| 153 | 0.710088 | 0.289912 |
| 154 | 0.7109 | 0.289101 |
| 155 | 0.711701 | 0.2883 |
| 156 | 0.71249 | 0.28751 |
| 157 | 0.713269 | 0.286731 |
| 158 | 0.714037 | 0.285963 |
| 159 | 0.714796 | 0.285204 |
| 160 | 0.76701 | 0.232989 |
| 161 | 0.768405 | 0.231594 |
| 162 | 0.769768 | 0.230231 |
| 163 | 0.771098 | 0.228902 |
| 164 | 0.772396 | 0.227603 |
| 165 | 0.773665 | 0.226335 |
| 166 | 0.774904 | 0.225096 |
| 167 | 0.776114 | 0.223886 |
| 168 | 0.777296 | 0.222704 |
| 169 | 0.778451 | 0.221549 |
| 170 | 0.779579 | 0.220421 |
| 171 | 0.780682 | 0.219319 |
| 172 | 0.781759 | 0.218241 |
| 173 | 0.782812 | 0.217188 |
| 174 | 0.783841 | 0.216159 |
| 175 | 0.784848 | 0.215151 |
| 176 | 0.785833 | 0.214168 |
| 177 | 0.786797 | 0.213203 |
| 178 | 0.787738 | 0.212262 |
| 179 | 0.788661 | 0.211339 |
| 180 | 0.789563 | 0.210437 |
| 181 | 0.790448 | 0.209552 |
| 182 | 0.791313 | 0.208688 |
| 183 | 0.792161 | 0.207839 |
| 184 | 0.792991 | 0.20701 |
| 185 | 0.793804 | 0.206196 |
| 186 | 0.794602 | 0.205398 |
| 187 | 0.795384 | 0.204616 |
| 188 | 0.796151 | 0.203848 |
| 189 | 0.796904 | 0.203097 |
| 190 | 0.797642 | 0.202358 |
| 191 | 0.798366 | 0.201635 |
| 192 | 0.799077 | 0.200923 |
| 193 | 0.799775 | 0.200224 |
| 194 | 0.800461 | 0.199539 |
| 195 | 0.801135 | 0.198865 |
| 196 | 0.801797 | 0.198204 |
| 197 | 0.802448 | 0.197552 |
| 198 | 0.803087 | 0.196912 |
| 199 | 0.803717 | 0.196283 |
| 200 | 0.804335 | 0.195664 |

V2 T-II-biased

|  | Male | Female |
| --- | --- | --- |
| 1 | 1 | 0 |
| 2 | 1 | 0 |
| 3 | 1 | 0 |
| 4 | 1 | 0 |
| 5 | 1 | 0 |
| 6 | 1 | 0 |
| 7 | 1 | 0 |
| 8 | 1 | 0 |
| 9 | 0 | 1 |
| 10 | 0 | 1 |
| 11 | 0 | 1 |
| 12 | 0 | 1 |
| 13 | 0 | 1 |
| 14 | 0 | 1 |
| 15 | 0 | 1 |
| 16 | 1 | 1 |
| 17 | 1 | 1 |
| 18 | 1 | 1 |
| 19 | 1 | 1 |
| 20 | 1 | 1 |
| 21 | 0 | 2 |
| 22 | 0 | 2 |
| 23 | 0 | 2 |
| 24 | 0 | 2 |
| 25 | 0 | 2 |
| 26 | 0 | 2 |
| 27 | 0 | 2 |
| 28 | 0 | 2 |
| 29 | 0 | 2 |
| 30 | 0 | 2 |
| 31 | 0 | 2 |
| 32 | 0 | 3 |
| 33 | 0 | 3 |
| 34 | 0 | 3 |
| 35 | 0 | 3 |
| 36 | 0 | 3 |
| 37 | 0 | 3 |
| 38 | 0 | 3 |
| 39 | 0 | 3 |
| 40 | 0 | 3 |
| 41 | 0 | 3 |
| 42 | 0 | 3 |
| 43 | 0 | 3 |
| 44 | 0 | 3 |
| 45 | 0 | 3 |
| 46 | 0 | 3 |
| 47 | 1 | 3 |
| 48 | 1 | 3 |
| 49 | 1 | 3 |
| 50 | 1 | 3 |
| 51 | 1 | 3 |
| 52 | 1 | 3 |
| 53 | 1 | 3 |
| 54 | 1 | 3 |
| 55 | 1 | 3 |
| 56 | 1 | 3 |
| 57 | 1 | 3 |
| 58 | 1 | 3 |
| 59 | 1 | 3 |
| 60 | 1 | 3 |
| 61 | 1 | 3 |
| 62 | 1 | 3 |
| 63 | 1 | 3 |
| 64 | 1 | 3 |
| 65 | 1 | 3 |
| 66 | 1 | 3 |
| 67 | 1 | 3 |
| 68 | 1 | 3 |
| 69 | 1 | 3 |
| 70 | 1 | 3 |
| 71 | 1 | 3 |
| 72 | 1 | 3 |
| 73 | 1 | 3 |
| 74 | 1 | 3 |
| 75 | 1 | 3 |
| 76 | 1 | 3 |
| 77 | 1 | 3 |
| 78 | 1 | 3 |
| 79 | 1 | 3 |
| 80 | 1 | 3 |
| 81 | 1 | 3 |
| 82 | 1 | 3 |
| 83 | 1 | 3 |
| 84 | 1 | 3 |
| 85 | 1 | 3 |
| 86 | 1 | 3 |
| 87 | 1 | 3 |
| 88 | 1 | 3 |
| 89 | 1 | 3 |
| 90 | 1 | 3 |
| 91 | 1 | 3 |
| 92 | 1 | 3 |
| 93 | 1 | 3 |
| 94 | 1 | 3 |
| 95 | 1 | 3 |
| 96 | 1 | 3 |
| 97 | 1 | 3 |
| 98 | 1 | 3 |
| 99 | 1 | 3 |
| 100 | 1 | 3 |
| 101 | 1 | 3 |
| 102 | 1 | 3 |
| 103 | 1 | 3 |
| 104 | 1 | 3 |
| 105 | 1 | 3 |
| 106 | 1 | 3 |
| 107 | 1 | 3 |
| 108 | 1 | 3 |
| 109 | 1 | 3 |
| 110 | 1 | 3 |
| 111 | 1 | 3 |
| 112 | 1 | 3 |
| 113 | 1 | 3 |
| 114 | 1 | 3 |
| 115 | 1 | 3 |
| 116 | 1 | 3 |
| 117 | 1 | 3 |
| 118 | 1 | 3 |
| 119 | 1 | 3 |
| 120 | 1 | 3 |
| 121 | 1 | 3 |
| 122 | 1 | 3 |
| 123 | 1 | 3 |
| 124 | 1 | 3 |
| 125 | 1 | 3 |
| 126 | 1 | 3 |
| 127 | 1 | 3 |
| 128 | 1 | 3 |
| 129 | 1 | 3 |
| 130 | 1 | 3 |
| 131 | 1 | 3 |
| 132 | 1 | 3 |
| 133 | 1 | 3 |
| 134 | 1 | 3 |
| 135 | 1 | 3 |
| 136 | 1 | 3 |
| 137 | 1 | 3 |
| 138 | 1 | 3 |
| 139 | 1 | 3 |
| 140 | 1 | 3 |
| 141 | 1 | 3 |
| 142 | 1 | 3 |
| 143 | 1 | 3 |
| 144 | 1 | 3 |
| 145 | 1 | 3 |
| 146 | 1 | 3 |
| 147 | 1 | 3 |
| 148 | 1 | 3 |
| 149 | 1 | 3 |
| 150 | 1 | 3 |
| 151 | 1 | 3 |
| 152 | 1 | 3 |
| 153 | 1 | 3 |
| 154 | 1 | 3 |
| 155 | 1 | 3 |
| 156 | 1 | 3 |
| 157 | 1 | 3 |
| 158 | 1 | 3 |
| 159 | 1 | 3 |
| 160 | 2 | 3 |
| 161 | 2 | 3 |
| 162 | 2 | 3 |
| 163 | 2 | 3 |
| 164 | 2 | 3 |
| 165 | 2 | 3 |
| 166 | 2 | 3 |
| 167 | 2 | 3 |
| 168 | 2 | 3 |
| 169 | 2 | 3 |
| 170 | 2 | 3 |
| 171 | 2 | 3 |
| 172 | 2 | 3 |
| 173 | 2 | 3 |
| 174 | 2 | 3 |
| 175 | 2 | 3 |
| 176 | 2 | 3 |
| 177 | 2 | 3 |
| 178 | 2 | 3 |
| 179 | 2 | 3 |
| 180 | 2 | 3 |
| 181 | 2 | 3 |
| 182 | 2 | 3 |
| 183 | 2 | 3 |
| 184 | 2 | 3 |
| 185 | 2 | 3 |
| 186 | 2 | 3 |
| 187 | 2 | 3 |
| 188 | 2 | 3 |
| 189 | 2 | 3 |
| 190 | 2 | 3 |
| 191 | 2 | 3 |
| 192 | 2 | 3 |
| 193 | 2 | 3 |
| 194 | 2 | 3 |
| 195 | 2 | 3 |
| 196 | 2 | 3 |
| 197 | 2 | 3 |
| 198 | 2 | 3 |
| 199 | 2 | 3 |
| 200 | 2 | 3 |

V2 Equal

|  | Male | Female |
| --- | --- | --- |
| 1 | 1 | 7.95949e-09 |
| 2 | 1 | 9.1343e-10 |
| 3 | 1 | 2.23014e-09 |
| 4 | 1 | 3.82055e-10 |
| 5 | 1 | 1.3941e-09 |
| 6 | 1 | 8.46819e-10 |
| 7 | 1 | 8.25926e-10 |
| 8 | 1 | 1.3148e-10 |
| 9 | 2.21119e-10 | 1 |
| 10 | 1.18642e-12 | 1 |
| 11 | 1.80916e-10 | 1 |
| 12 | 1.09942e-10 | 1 |
| 13 | 8.59513e-11 | 1 |
| 14 | 9.58585e-11 | 1 |
| 15 | 8.50885e-11 | 1 |
| 16 | 0.0310361 | 0.968964 |
| 17 | 0.0922134 | 0.907787 |
| 18 | 0.146591 | 0.853409 |
| 19 | 0.195242 | 0.804758 |
| 20 | 0.239025 | 0.760975 |
| 21 | 5.83439e-13 | 1 |
| 22 | 4.07028e-12 | 1 |
| 23 | 6.73998e-12 | 1 |
| 24 | 2.47302e-12 | 1 |
| 25 | 3.84348e-12 | 1 |
| 26 | 9.65874e-12 | 1 |
| 27 | 1.59763e-11 | 1 |
| 28 | 6.86632e-12 | 1 |
| 29 | 3.19532e-10 | 1 |
| 30 | 9.31987e-10 | 1 |
| 31 | 1.5882e-08 | 1 |
| 32 | 2.1219e-12 | 1 |
| 33 | 2.71352e-12 | 1 |
| 34 | 1.87194e-13 | 1 |
| 35 | 5.45337e-12 | 1 |
| 36 | 8.2434e-12 | 1 |
| 37 | 2.55522e-12 | 1 |
| 38 | 1.6118e-11 | 1 |
| 39 | 6.43906e-12 | 1 |
| 40 | 8.33229e-12 | 1 |
| 41 | 1.367e-11 | 1 |
| 42 | 9.15507e-11 | 1 |
| 43 | 1.67817e-12 | 1 |
| 44 | 2.03514e-12 | 1 |
| 45 | 2.57565e-12 | 1 |
| 46 | 3.36999e-12 | 1 |
| 47 | 1.80875e-11 | 1 |
| 48 | 1.23651e-12 | 1 |
| 49 | 3.4288e-11 | 1 |
| 50 | 1.20109e-11 | 1 |
| 51 | 1.35758e-11 | 1 |
| 52 | 3.86367e-12 | 1 |
| 53 | 2.76735e-11 | 1 |
| 54 | 3.99409e-11 | 1 |
| 55 | 1.62811e-10 | 1 |
| 56 | 3.91986e-11 | 1 |
| 57 | 4.82567e-11 | 1 |
| 58 | 6.04899e-11 | 1 |
| 59 | 7.69186e-11 | 1 |
| 60 | 9.89206e-11 | 1 |
| 61 | 2.5671e-11 | 1 |
| 62 | 1.34187e-10 | 1 |
| 63 | 2.20465e-10 | 1 |
| 64 | 2.91202e-10 | 1 |
| 65 | 3.09076e-10 | 1 |
| 66 | 8.2366e-11 | 1 |
| 67 | 5.51277e-10 | 1 |
| 68 | 7.41878e-10 | 1 |
| 69 | 2.51258e-10 | 1 |
| 70 | 1.37326e-09 | 1 |
| 71 | 1.89892e-09 | 1 |
| 72 | 2.67073e-09 | 1 |
| 73 | 7.70566e-10 | 1 |
| 74 | 1.15777e-09 | 1 |
| 75 | 7.45552e-09 | 1 |
| 76 | 2.77521e-09 | 1 |
| 77 | 3.95092e-08 | 1 |
| 78 | 0.00610305 | 0.993897 |
| 79 | 0.0249647 | 0.975035 |
| 80 | 0.0433494 | 0.956651 |
| 81 | 0.0612697 | 0.93873 |
| 82 | 0.0787438 | 0.921256 |
| 83 | 0.0957848 | 0.904215 |
| 84 | 0.112408 | 0.887592 |
| 85 | 0.128624 | 0.871376 |
| 86 | 0.144447 | 0.855553 |
| 87 | 0.159887 | 0.840113 |
| 88 | 0.174953 | 0.825047 |
| 89 | 0.189656 | 0.810344 |
| 90 | 0.204004 | 0.795996 |
| 91 | 0.218004 | 0.781996 |
| 92 | 0.231662 | 0.768338 |
| 93 | 0.244985 | 0.755015 |
| 94 | 0.257977 | 0.742023 |
| 95 | 0.270642 | 0.729358 |
| 96 | 0.282983 | 0.717017 |
| 97 | 0.295003 | 0.704997 |
| 98 | 0.306701 | 0.693299 |
| 99 | 0.318079 | 0.681921 |
| 100 | 0.329137 | 0.670863 |
| 101 | 0.339872 | 0.660128 |
| 102 | 0.350284 | 0.649716 |
| 103 | 0.36037 | 0.63963 |
| 104 | 0.370128 | 0.629872 |
| 105 | 0.379555 | 0.620445 |
| 106 | 0.38865 | 0.61135 |
| 107 | 0.397411 | 0.602589 |
| 108 | 0.405836 | 0.594164 |
| 109 | 0.413925 | 0.586075 |
| 110 | 0.421681 | 0.578319 |
| 111 | 0.429104 | 0.570896 |
| 112 | 0.436198 | 0.563802 |
| 113 | 0.44297 | 0.55703 |
| 114 | 0.449425 | 0.550575 |
| 115 | 0.455571 | 0.544429 |
| 116 | 0.461419 | 0.538581 |
| 117 | 0.466978 | 0.533022 |
| 118 | 0.47226 | 0.52774 |
| 119 | 0.477277 | 0.522723 |
| 120 | 0.482042 | 0.517958 |
| 121 | 0.486567 | 0.513433 |
| 122 | 0.490866 | 0.509134 |
| 123 | 0.494951 | 0.505049 |
| 124 | 0.498835 | 0.501165 |
| 125 | 0.50253 | 0.49747 |
| 126 | 0.506049 | 0.493951 |
| 127 | 0.509403 | 0.490597 |
| 128 | 0.512602 | 0.487398 |
| 129 | 0.515657 | 0.484343 |
| 130 | 0.518577 | 0.481423 |
| 131 | 0.521372 | 0.478628 |
| 132 | 0.52405 | 0.47595 |
| 133 | 0.526618 | 0.473382 |
| 134 | 0.529085 | 0.470915 |
| 135 | 0.531457 | 0.468543 |
| 136 | 0.53374 | 0.46626 |
| 137 | 0.53594 | 0.46406 |
| 138 | 0.538062 | 0.461938 |
| 139 | 0.540113 | 0.459887 |
| 140 | 0.542095 | 0.457905 |
| 141 | 0.544014 | 0.455986 |
| 142 | 0.545873 | 0.454127 |
| 143 | 0.547677 | 0.452323 |
| 144 | 0.549428 | 0.450572 |
| 145 | 0.551129 | 0.448871 |
| 146 | 0.552784 | 0.447216 |
| 147 | 0.554395 | 0.445605 |
| 148 | 0.555965 | 0.444035 |
| 149 | 0.557496 | 0.442504 |
| 150 | 0.558989 | 0.441011 |
| 151 | 0.560448 | 0.439552 |
| 152 | 0.561873 | 0.438127 |
| 153 | 0.563266 | 0.436734 |
| 154 | 0.56463 | 0.43537 |
| 155 | 0.565965 | 0.434035 |
| 156 | 0.567273 | 0.432727 |
| 157 | 0.568555 | 0.431445 |
| 158 | 0.569812 | 0.430188 |
| 159 | 0.571045 | 0.428955 |
| 160 | 0.572256 | 0.427744 |
| 161 | 0.573445 | 0.426555 |
| 162 | 0.574613 | 0.425387 |
| 163 | 0.575761 | 0.424239 |
| 164 | 0.57689 | 0.42311 |
| 165 | 0.578001 | 0.421999 |
| 166 | 0.579094 | 0.420906 |
| 167 | 0.580169 | 0.419831 |
| 168 | 0.581228 | 0.418772 |
| 169 | 0.582271 | 0.417729 |
| 170 | 0.583298 | 0.416702 |
| 171 | 0.584311 | 0.415689 |
| 172 | 0.585309 | 0.414691 |
| 173 | 0.586293 | 0.413707 |
| 174 | 0.587263 | 0.412737 |
| 175 | 0.588221 | 0.411779 |
| 176 | 0.589165 | 0.410835 |
| 177 | 0.590097 | 0.409903 |
| 178 | 0.591016 | 0.408984 |
| 179 | 0.591924 | 0.408076 |
| 180 | 0.592821 | 0.407179 |
| 181 | 0.593706 | 0.406294 |
| 182 | 0.59458 | 0.40542 |
| 183 | 0.595444 | 0.404556 |
| 184 | 0.596297 | 0.403703 |
| 185 | 0.59714 | 0.40286 |
| 186 | 0.665877 | 0.334123 |
| 187 | 0.667564 | 0.332436 |
| 188 | 0.669212 | 0.330788 |
| 189 | 0.670821 | 0.329179 |
| 190 | 0.672393 | 0.327607 |
| 191 | 0.673928 | 0.326072 |
| 192 | 0.675428 | 0.324572 |
| 193 | 0.676893 | 0.323107 |
| 194 | 0.678325 | 0.321675 |
| 195 | 0.679725 | 0.320275 |
| 196 | 0.681093 | 0.318907 |
| 197 | 0.682431 | 0.317569 |
| 198 | 0.683739 | 0.316261 |
| 199 | 0.685019 | 0.314981 |
| 200 | 0.686271 | 0.313729 |

V2 Equal

|  | Male | Female |
| --- | --- | --- |
| 1 | 1 | 0 |
| 2 | 1 | 0 |
| 3 | 1 | 0 |
| 4 | 1 | 0 |
| 5 | 1 | 0 |
| 6 | 1 | 0 |
| 7 | 1 | 0 |
| 8 | 1 | 0 |
| 9 | 0 | 1 |
| 10 | 0 | 1 |
| 11 | 0 | 1 |
| 12 | 0 | 1 |
| 13 | 0 | 1 |
| 14 | 0 | 1 |
| 15 | 0 | 1 |
| 16 | 1 | 1 |
| 17 | 1 | 1 |
| 18 | 1 | 1 |
| 19 | 1 | 1 |
| 20 | 1 | 1 |
| 21 | 0 | 2 |
| 22 | 0 | 2 |
| 23 | 0 | 2 |
| 24 | 0 | 2 |
| 25 | 0 | 2 |
| 26 | 0 | 2 |
| 27 | 0 | 2 |
| 28 | 0 | 2 |
| 29 | 0 | 2 |
| 30 | 0 | 2 |
| 31 | 0 | 2 |
| 32 | 0 | 3 |
| 33 | 0 | 3 |
| 34 | 0 | 3 |
| 35 | 0 | 3 |
| 36 | 0 | 3 |
| 37 | 0 | 3 |
| 38 | 0 | 3 |
| 39 | 0 | 3 |
| 40 | 0 | 3 |
| 41 | 0 | 3 |
| 42 | 0 | 3 |
| 43 | 0 | 4 |
| 44 | 0 | 4 |
| 45 | 0 | 4 |
| 46 | 0 | 4 |
| 47 | 0 | 4 |
| 48 | 0 | 4 |
| 49 | 0 | 4 |
| 50 | 0 | 4 |
| 51 | 0 | 4 |
| 52 | 0 | 4 |
| 53 | 0 | 4 |
| 54 | 0 | 4 |
| 55 | 0 | 5 |
| 56 | 0 | 5 |
| 57 | 0 | 5 |
| 58 | 0 | 5 |
| 59 | 0 | 5 |
| 60 | 0 | 5 |
| 61 | 0 | 5 |
| 62 | 0 | 5 |
| 63 | 0 | 5 |
| 64 | 0 | 5 |
| 65 | 0 | 5 |
| 66 | 0 | 5 |
| 67 | 0 | 5 |
| 68 | 0 | 5 |
| 69 | 0 | 5 |
| 70 | 0 | 5 |
| 71 | 0 | 5 |
| 72 | 0 | 5 |
| 73 | 0 | 5 |
| 74 | 0 | 5 |
| 75 | 0 | 5 |
| 76 | 0 | 5 |
| 77 | 0 | 5 |
| 78 | 1 | 5 |
| 79 | 1 | 5 |
| 80 | 1 | 5 |
| 81 | 1 | 5 |
| 82 | 1 | 5 |
| 83 | 1 | 5 |
| 84 | 1 | 5 |
| 85 | 1 | 5 |
| 86 | 1 | 5 |
| 87 | 1 | 5 |
| 88 | 1 | 5 |
| 89 | 1 | 5 |
| 90 | 1 | 5 |
| 91 | 1 | 5 |
| 92 | 1 | 5 |
| 93 | 1 | 5 |
| 94 | 1 | 5 |
| 95 | 1 | 5 |
| 96 | 1 | 5 |
| 97 | 1 | 5 |
| 98 | 1 | 5 |
| 99 | 1 | 5 |
| 100 | 1 | 5 |
| 101 | 1 | 5 |
| 102 | 1 | 5 |
| 103 | 1 | 5 |
| 104 | 1 | 5 |
| 105 | 1 | 5 |
| 106 | 1 | 5 |
| 107 | 1 | 5 |
| 108 | 1 | 5 |
| 109 | 1 | 5 |
| 110 | 1 | 5 |
| 111 | 1 | 5 |
| 112 | 1 | 5 |
| 113 | 1 | 5 |
| 114 | 1 | 5 |
| 115 | 1 | 5 |
| 116 | 1 | 5 |
| 117 | 1 | 5 |
| 118 | 1 | 5 |
| 119 | 1 | 5 |
| 120 | 1 | 5 |
| 121 | 1 | 5 |
| 122 | 1 | 5 |
| 123 | 1 | 5 |
| 124 | 1 | 5 |
| 125 | 1 | 5 |
| 126 | 1 | 5 |
| 127 | 1 | 5 |
| 128 | 1 | 5 |
| 129 | 1 | 5 |
| 130 | 1 | 5 |
| 131 | 1 | 5 |
| 132 | 1 | 5 |
| 133 | 1 | 5 |
| 134 | 1 | 5 |
| 135 | 1 | 5 |
| 136 | 1 | 5 |
| 137 | 1 | 5 |
| 138 | 1 | 5 |
| 139 | 1 | 5 |
| 140 | 1 | 5 |
| 141 | 1 | 5 |
| 142 | 1 | 5 |
| 143 | 1 | 5 |
| 144 | 1 | 5 |
| 145 | 1 | 5 |
| 146 | 1 | 5 |
| 147 | 1 | 5 |
| 148 | 1 | 5 |
| 149 | 1 | 5 |
| 150 | 1 | 5 |
| 151 | 1 | 5 |
| 152 | 1 | 5 |
| 153 | 1 | 5 |
| 154 | 1 | 5 |
| 155 | 1 | 5 |
| 156 | 1 | 5 |
| 157 | 1 | 5 |
| 158 | 1 | 5 |
| 159 | 1 | 5 |
| 160 | 1 | 5 |
| 161 | 1 | 5 |
| 162 | 1 | 5 |
| 163 | 1 | 5 |
| 164 | 1 | 5 |
| 165 | 1 | 5 |
| 166 | 1 | 5 |
| 167 | 1 | 5 |
| 168 | 1 | 5 |
| 169 | 1 | 5 |
| 170 | 1 | 5 |
| 171 | 1 | 5 |
| 172 | 1 | 5 |
| 173 | 1 | 5 |
| 174 | 1 | 5 |
| 175 | 1 | 5 |
| 176 | 1 | 5 |
| 177 | 1 | 5 |
| 178 | 1 | 5 |
| 179 | 1 | 5 |
| 180 | 1 | 5 |
| 181 | 1 | 5 |
| 182 | 1 | 5 |
| 183 | 1 | 5 |
| 184 | 1 | 5 |
| 185 | 1 | 5 |
| 186 | 2 | 5 |
| 187 | 2 | 5 |
| 188 | 2 | 5 |
| 189 | 2 | 5 |
| 190 | 2 | 5 |
| 191 | 2 | 5 |
| 192 | 2 | 5 |
| 193 | 2 | 5 |
| 194 | 2 | 5 |
| 195 | 2 | 5 |
| 196 | 2 | 5 |
| 197 | 2 | 5 |
| 198 | 2 | 5 |
| 199 | 2 | 5 |
| 200 | 2 | 5 |

V2 T-I-biased

|  | Male | Female |
| --- | --- | --- |
| 1 | 1 | 2.25068e-09 |
| 2 | 1 | 1.03797e-09 |
| 3 | 1 | 7.93272e-10 |
| 4 | 1 | 5.4365e-10 |
| 5 | 1 | 1.58591e-09 |
| 6 | 1 | 3.00728e-10 |
| 7 | 1 | 2.34373e-10 |
| 8 | 4.94803e-08 | 1 |
| 9 | 1.32677e-10 | 1 |
| 10 | 7.11865e-13 | 1 |
| 11 | 2.71385e-11 | 1 |
| 12 | 3.29869e-10 | 1 |
| 13 | 5.16005e-11 | 1 |
| 14 | 5.76759e-11 | 1 |
| 15 | 5.20825e-11 | 1 |
| 16 | 0.031036 | 0.968964 |
| 17 | 0.0922134 | 0.907787 |
| 18 | 0.146591 | 0.853409 |
| 19 | 0.195242 | 0.804758 |
| 20 | 0.239025 | 0.760975 |
| 21 | 1.75032e-12 | 1 |
| 22 | 1.22108e-13 | 1 |
| 23 | 8.08797e-13 | 1 |
| 24 | 1.48381e-12 | 1 |
| 25 | 2.88261e-12 | 1 |
| 26 | 1.15903e-12 | 1 |
| 27 | 9.58556e-12 | 1 |
| 28 | 4.11982e-12 | 1 |
| 29 | 3.83452e-11 | 1 |
| 30 | 1.11837e-10 | 1 |
| 31 | 3.81027e-08 | 1 |
| 32 | 1.27314e-12 | 1 |
| 33 | 8.14055e-14 | 1 |
| 34 | 2.24632e-12 | 1 |
| 35 | 6.54403e-13 | 1 |
| 36 | 4.94604e-12 | 1 |
| 37 | 1.53314e-12 | 1 |
| 38 | 2.4177e-12 | 1 |
| 39 | 1.54537e-11 | 1 |
| 40 | 2.49969e-11 | 1 |
| 41 | 8.20209e-12 | 1 |
| 42 | 2.74653e-10 | 1 |
| 43 | 1.0069e-12 | 1 |
| 44 | 1.22108e-12 | 1 |
| 45 | 3.09078e-13 | 1 |
| 46 | 2.02199e-12 | 1 |
| 47 | 2.71312e-12 | 1 |
| 48 | 1.48381e-11 | 1 |
| 49 | 1.02864e-12 | 1 |
| 50 | 5.76521e-12 | 1 |
| 51 | 8.14548e-12 | 1 |
| 52 | 1.15903e-11 | 1 |
| 53 | 1.66042e-11 | 1 |
| 54 | 4.7929e-12 | 1 |
| 55 | 9.76866e-13 | 1 |
| 56 | 1.17595e-12 | 1 |
| 57 | 5.7908e-12 | 1 |
| 58 | 3.62939e-11 | 1 |
| 59 | 4.61512e-11 | 1 |
| 60 | 1.18705e-11 | 1 |
| 61 | 3.85064e-12 | 1 |
| 62 | 4.0256e-12 | 1 |
| 63 | 2.6456e-11 | 1 |
| 64 | 6.98886e-12 | 1 |
| 65 | 4.63611e-11 | 1 |
| 66 | 1.23549e-11 | 1 |
| 67 | 3.30766e-10 | 1 |
| 68 | 1.12316e-10 | 1 |
| 69 | 2.69599e-11 | 1 |
| 70 | 1.63601e-10 | 1 |
| 71 | 4.00776e-11 | 1 |
| 72 | 4.94606e-11 | 1 |
| 73 | 6.14179e-11 | 1 |
| 74 | 7.66586e-11 | 1 |
| 75 | 3.84347e-10 | 1 |
| 76 | 9.67078e-11 | 1 |
| 77 | 3.0516e-11 | 1 |
| 78 | 7.72685e-10 | 1 |
| 79 | 1.96236e-10 | 1 |
| 80 | 2.49969e-10 | 1 |
| 81 | 3.99446e-10 | 1 |
| 82 | 4.10101e-10 | 1 |
| 83 | 1.4154e-10 | 1 |
| 84 | 4.23945e-09 | 1 |
| 85 | 1.02047e-09 | 1 |
| 86 | 6.16451e-11 | 1 |
| 87 | 1.49448e-09 | 1 |
| 88 | 9.08376e-11 | 1 |
| 89 | 4.42832e-10 | 1 |
| 90 | 2.16384e-09 | 1 |
| 91 | 6.62302e-10 | 1 |
| 92 | 8.12643e-10 | 1 |
| 93 | 1.99877e-10 | 1 |
| 94 | 6.16041e-11 | 1 |
| 95 | 1.52315e-09 | 1 |
| 96 | 7.55649e-11 | 1 |
| 97 | 4.7045e-10 | 1 |
| 98 | 1.47166e-10 | 1 |
| 99 | 7.40761e-10 | 1 |
| 100 | 7.51672e-10 | 1 |
| 101 | 2.40783e-10 | 1 |
| 102 | 1.56384e-09 | 1 |
| 103 | 1.03512e-10 | 1 |
| 104 | 5.63268e-10 | 1 |
| 105 | 7.98747e-10 | 1 |
| 106 | 1.21872e-09 | 1 |
| 107 | 2.12134e-09 | 1 |
| 108 | 2.18418e-08 | 1 |
| 109 | 0.00419955 | 0.9958 |
| 110 | 0.0211347 | 0.978865 |
| 111 | 0.0377453 | 0.962255 |
| 112 | 0.0540416 | 0.945958 |
| 113 | 0.0700272 | 0.929973 |
| 114 | 0.085707 | 0.914293 |
| 115 | 0.101083 | 0.898917 |
| 116 | 0.116158 | 0.883842 |
| 117 | 0.130933 | 0.869067 |
| 118 | 0.145407 | 0.854593 |
| 119 | 0.159579 | 0.840421 |
| 120 | 0.173445 | 0.826555 |
| 121 | 0.187001 | 0.812999 |
| 122 | 0.200241 | 0.799759 |
| 123 | 0.213157 | 0.786843 |
| 124 | 0.22574 | 0.77426 |
| 125 | 0.23798 | 0.76202 |
| 126 | 0.249864 | 0.750136 |
| 127 | 0.261381 | 0.738619 |
| 128 | 0.272516 | 0.727484 |
| 129 | 0.283259 | 0.716741 |
| 130 | 0.293596 | 0.706404 |
| 131 | 0.303518 | 0.696482 |
| 132 | 0.313017 | 0.686983 |
| 133 | 0.322087 | 0.677913 |
| 134 | 0.330726 | 0.669274 |
| 135 | 0.338935 | 0.661065 |
| 136 | 0.346721 | 0.653279 |
| 137 | 0.354091 | 0.645909 |
| 138 | 0.361058 | 0.638942 |
| 139 | 0.367635 | 0.632365 |
| 140 | 0.37384 | 0.62616 |
| 141 | 0.37969 | 0.62031 |
| 142 | 0.385204 | 0.614796 |
| 143 | 0.390403 | 0.609597 |
| 144 | 0.395307 | 0.604693 |
| 145 | 0.399934 | 0.600066 |
| 146 | 0.404304 | 0.595696 |
| 147 | 0.408435 | 0.591565 |
| 148 | 0.412345 | 0.587655 |
| 149 | 0.416049 | 0.583951 |
| 150 | 0.419564 | 0.580436 |
| 151 | 0.422902 | 0.577098 |
| 152 | 0.426078 | 0.573922 |
| 153 | 0.429104 | 0.570896 |
| 154 | 0.431991 | 0.568009 |
| 155 | 0.434749 | 0.565251 |
| 156 | 0.437388 | 0.562612 |
| 157 | 0.439916 | 0.560084 |
| 158 | 0.442342 | 0.557658 |
| 159 | 0.444672 | 0.555328 |
| 160 | 0.446914 | 0.553086 |
| 161 | 0.449074 | 0.550926 |
| 162 | 0.451157 | 0.548843 |
| 163 | 0.453169 | 0.546831 |
| 164 | 0.455114 | 0.544886 |
| 165 | 0.456996 | 0.543004 |
| 166 | 0.45882 | 0.54118 |
| 167 | 0.46059 | 0.53941 |
| 168 | 0.462308 | 0.537692 |
| 169 | 0.463978 | 0.536022 |
| 170 | 0.465602 | 0.534398 |
| 171 | 0.467184 | 0.532816 |
| 172 | 0.468725 | 0.531275 |
| 173 | 0.470229 | 0.529771 |
| 174 | 0.471697 | 0.528303 |
| 175 | 0.473131 | 0.526869 |
| 176 | 0.474532 | 0.525468 |
| 177 | 0.475903 | 0.524097 |
| 178 | 0.477245 | 0.522755 |
| 179 | 0.47856 | 0.52144 |
| 180 | 0.479848 | 0.520152 |
| 181 | 0.481112 | 0.518888 |
| 182 | 0.482351 | 0.517649 |
| 183 | 0.483568 | 0.516432 |
| 184 | 0.484763 | 0.515237 |
| 185 | 0.485938 | 0.514062 |
| 186 | 0.487092 | 0.512908 |
| 187 | 0.488227 | 0.511773 |
| 188 | 0.489344 | 0.510656 |
| 189 | 0.490444 | 0.509556 |
| 190 | 0.491526 | 0.508474 |
| 191 | 0.492592 | 0.507408 |
| 192 | 0.493642 | 0.506358 |
| 193 | 0.494677 | 0.505323 |
| 194 | 0.495698 | 0.504302 |
| 195 | 0.496704 | 0.503296 |
| 196 | 0.497697 | 0.502303 |
| 197 | 0.498676 | 0.501324 |
| 198 | 0.499642 | 0.500358 |
| 199 | 0.500596 | 0.499404 |
| 200 | 0.501538 | 0.498462 |

V2 T-I-biased

|  | Male | Female |
| --- | --- | --- |
| 1 | 1 | 0 |
| 2 | 1 | 0 |
| 3 | 1 | 0 |
| 4 | 1 | 0 |
| 5 | 1 | 0 |
| 6 | 1 | 0 |
| 7 | 1 | 0 |
| 8 | 0 | 1 |
| 9 | 0 | 1 |
| 10 | 0 | 1 |
| 11 | 0 | 1 |
| 12 | 0 | 1 |
| 13 | 0 | 1 |
| 14 | 0 | 1 |
| 15 | 0 | 1 |
| 16 | 1 | 1 |
| 17 | 1 | 1 |
| 18 | 1 | 1 |
| 19 | 1 | 1 |
| 20 | 1 | 1 |
| 21 | 0 | 2 |
| 22 | 0 | 2 |
| 23 | 0 | 2 |
| 24 | 0 | 2 |
| 25 | 0 | 2 |
| 26 | 0 | 2 |
| 27 | 0 | 2 |
| 28 | 0 | 2 |
| 29 | 0 | 2 |
| 30 | 0 | 2 |
| 31 | 0 | 2 |
| 32 | 0 | 3 |
| 33 | 0 | 3 |
| 34 | 0 | 3 |
| 35 | 0 | 3 |
| 36 | 0 | 3 |
| 37 | 0 | 3 |
| 38 | 0 | 3 |
| 39 | 0 | 3 |
| 40 | 0 | 3 |
| 41 | 0 | 3 |
| 42 | 0 | 3 |
| 43 | 0 | 4 |
| 44 | 0 | 4 |
| 45 | 0 | 4 |
| 46 | 0 | 4 |
| 47 | 0 | 4 |
| 48 | 0 | 4 |
| 49 | 0 | 4 |
| 50 | 0 | 4 |
| 51 | 0 | 4 |
| 52 | 0 | 4 |
| 53 | 0 | 4 |
| 54 | 0 | 4 |
| 55 | 0 | 5 |
| 56 | 0 | 5 |
| 57 | 0 | 5 |
| 58 | 0 | 5 |
| 59 | 0 | 5 |
| 60 | 0 | 5 |
| 61 | 0 | 5 |
| 62 | 0 | 5 |
| 63 | 0 | 5 |
| 64 | 0 | 5 |
| 65 | 0 | 5 |
| 66 | 0 | 5 |
| 67 | 0 | 5 |
| 68 | 0 | 6 |
| 69 | 0 | 6 |
| 70 | 0 | 6 |
| 71 | 0 | 6 |
| 72 | 0 | 6 |
| 73 | 0 | 6 |
| 74 | 0 | 6 |
| 75 | 0 | 6 |
| 76 | 0 | 6 |
| 77 | 0 | 6 |
| 78 | 0 | 6 |
| 79 | 0 | 6 |
| 80 | 0 | 6 |
| 81 | 0 | 6 |
| 82 | 0 | 6 |
| 83 | 0 | 7 |
| 84 | 0 | 7 |
| 85 | 0 | 7 |
| 86 | 0 | 7 |
| 87 | 0 | 7 |
| 88 | 0 | 7 |
| 89 | 0 | 7 |
| 90 | 0 | 7 |
| 91 | 0 | 7 |
| 92 | 0 | 7 |
| 93 | 0 | 7 |
| 94 | 0 | 7 |
| 95 | 0 | 7 |
| 96 | 0 | 7 |
| 97 | 0 | 7 |
| 98 | 0 | 7 |
| 99 | 0 | 7 |
| 100 | 0 | 7 |
| 101 | 0 | 7 |
| 102 | 0 | 7 |
| 103 | 0 | 7 |
| 104 | 0 | 7 |
| 105 | 0 | 7 |
| 106 | 0 | 7 |
| 107 | 0 | 7 |
| 108 | 0 | 7 |
| 109 | 1 | 7 |
| 110 | 1 | 7 |
| 111 | 1 | 7 |
| 112 | 1 | 7 |
| 113 | 1 | 7 |
| 114 | 1 | 7 |
| 115 | 1 | 7 |
| 116 | 1 | 7 |
| 117 | 1 | 7 |
| 118 | 1 | 7 |
| 119 | 1 | 7 |
| 120 | 1 | 7 |
| 121 | 1 | 7 |
| 122 | 1 | 7 |
| 123 | 1 | 7 |
| 124 | 1 | 7 |
| 125 | 1 | 7 |
| 126 | 1 | 7 |
| 127 | 1 | 7 |
| 128 | 1 | 7 |
| 129 | 1 | 7 |
| 130 | 1 | 7 |
| 131 | 1 | 7 |
| 132 | 1 | 7 |
| 133 | 1 | 7 |
| 134 | 1 | 7 |
| 135 | 1 | 7 |
| 136 | 1 | 7 |
| 137 | 1 | 7 |
| 138 | 1 | 7 |
| 139 | 1 | 7 |
| 140 | 1 | 7 |
| 141 | 1 | 7 |
| 142 | 1 | 7 |
| 143 | 1 | 7 |
| 144 | 1 | 7 |
| 145 | 1 | 7 |
| 146 | 1 | 7 |
| 147 | 1 | 7 |
| 148 | 1 | 7 |
| 149 | 1 | 7 |
| 150 | 1 | 7 |
| 151 | 1 | 7 |
| 152 | 1 | 7 |
| 153 | 1 | 7 |
| 154 | 1 | 7 |
| 155 | 1 | 7 |
| 156 | 1 | 7 |
| 157 | 1 | 7 |
| 158 | 1 | 7 |
| 159 | 1 | 7 |
| 160 | 1 | 7 |
| 161 | 1 | 7 |
| 162 | 1 | 7 |
| 163 | 1 | 7 |
| 164 | 1 | 7 |
| 165 | 1 | 7 |
| 166 | 1 | 7 |
| 167 | 1 | 7 |
| 168 | 1 | 7 |
| 169 | 1 | 7 |
| 170 | 1 | 7 |
| 171 | 1 | 7 |
| 172 | 1 | 7 |
| 173 | 1 | 7 |
| 174 | 1 | 7 |
| 175 | 1 | 7 |
| 176 | 1 | 7 |
| 177 | 1 | 7 |
| 178 | 1 | 7 |
| 179 | 1 | 7 |
| 180 | 1 | 7 |
| 181 | 1 | 7 |
| 182 | 1 | 7 |
| 183 | 1 | 7 |
| 184 | 1 | 7 |
| 185 | 1 | 7 |
| 186 | 1 | 7 |
| 187 | 1 | 7 |
| 188 | 1 | 7 |
| 189 | 1 | 7 |
| 190 | 1 | 7 |
| 191 | 1 | 7 |
| 192 | 1 | 7 |
| 193 | 1 | 7 |
| 194 | 1 | 7 |
| 195 | 1 | 7 |
| 196 | 1 | 7 |
| 197 | 1 | 7 |
| 198 | 1 | 7 |
| 199 | 1 | 7 |
| 200 | 1 | 7 |

Figure S5 | Same as Figure S4 for model version 2.

**Model Version 3**

V3 Functions

|  |  |  |
| --- | --- | --- |
| 0 | -0.000354096 | -3.88578e-15 |
| 0.01 | -0.000205365 | 0.00150382 |
| 0.02 | -5.58915e-05 | 0.00301095 |
| 0.03 | 9.43273e-05 | 0.0045214 |
| 0.04 | 0.000245295 | 0.00603516 |
| 0.05 | 0.000397016 | 0.00755226 |
| 0.06 | 0.000549493 | 0.0090727 |
| 0.07 | 0.000702731 | 0.0105965 |
| 0.08 | 0.000856733 | 0.0121236 |
| 0.09 | 0.0010115 | 0.0136541 |
| 0.1 | 0.00116705 | 0.0151879 |
| 0.11 | 0.00132336 | 0.0167251 |
| 0.12 | 0.00148046 | 0.0182657 |
| 0.13 | 0.00163834 | 0.0198096 |
| 0.14 | 0.00179701 | 0.021357 |
| 0.15 | 0.00195646 | 0.0229077 |
| 0.16 | 0.00211672 | 0.0244618 |
| 0.17 | 0.00227777 | 0.0260194 |
| 0.18 | 0.00243963 | 0.0275803 |
| 0.19 | 0.00260229 | 0.0291447 |
| 0.2 | 0.00276576 | 0.0307125 |
| 0.21 | 0.00293005 | 0.0322837 |
| 0.22 | 0.00309516 | 0.0338583 |
| 0.23 | 0.00326109 | 0.0354364 |
| 0.24 | 0.00342784 | 0.037018 |
| 0.25 | 0.00359543 | 0.038603 |
| 0.26 | 0.00376386 | 0.0401914 |
| 0.27 | 0.00393312 | 0.0417833 |
| 0.28 | 0.00410322 | 0.0433787 |
| 0.29 | 0.00427418 | 0.0449776 |
| 0.3 | 0.00444599 | 0.04658 |
| 0.31 | 0.00461865 | 0.0481859 |
| 0.32 | 0.00479217 | 0.0497952 |
| 0.33 | 0.00496656 | 0.0514081 |
| 0.34 | 0.00514182 | 0.0530245 |
| 0.35 | 0.00531795 | 0.0546444 |
| 0.36 | 0.00549495 | 0.0562678 |
| 0.37 | 0.00567284 | 0.0578947 |
| 0.38 | 0.00585162 | 0.0595252 |
| 0.39 | 0.00603129 | 0.0611593 |
| 0.4 | 0.00621185 | 0.0627968 |
| 0.41 | 0.00639331 | 0.064438 |
| 0.42 | 0.00657568 | 0.0660827 |
| 0.43 | 0.00675895 | 0.067731 |
| 0.44 | 0.00694314 | 0.0693828 |
| 0.45 | 0.00712824 | 0.0710383 |
| 0.46 | 0.00731427 | 0.0726973 |
| 0.47 | 0.00750123 | 0.0743599 |
| 0.48 | 0.00768911 | 0.0760262 |
| 0.49 | 0.00787793 | 0.077696 |
| 0.5 | 0.00806769 | 0.0793695 |
| 0.51 | 0.0082584 | 0.0810465 |
| 0.52 | 0.00845005 | 0.0827272 |
| 0.53 | 0.00864266 | 0.0844116 |
| 0.54 | 0.00883623 | 0.0860996 |
| 0.55 | 0.00903077 | 0.0877912 |
| 0.56 | 0.00922627 | 0.0894865 |
| 0.57 | 0.00942275 | 0.0911854 |
| 0.58 | 0.0096202 | 0.092888 |
| 0.59 | 0.00981864 | 0.0945943 |
| 0.6 | 0.0100181 | 0.0963043 |
| 0.61 | 0.0102185 | 0.098018 |
| 0.62 | 0.0104199 | 0.0997353 |
| 0.63 | 0.0106223 | 0.101456 |
| 0.64 | 0.0108257 | 0.103181 |
| 0.65 | 0.0110302 | 0.10491 |
| 0.66 | 0.0112356 | 0.106642 |
| 0.67 | 0.0114421 | 0.108378 |
| 0.68 | 0.0116496 | 0.110117 |
| 0.69 | 0.0118582 | 0.111861 |
| 0.7 | 0.0120678 | 0.113608 |
| 0.71 | 0.0122784 | 0.115359 |
| 0.72 | 0.01249 | 0.117113 |
| 0.73 | 0.0127028 | 0.118872 |
| 0.74 | 0.0129166 | 0.120634 |
| 0.75 | 0.0131314 | 0.1224 |
| 0.76 | 0.0133473 | 0.12417 |
| 0.77 | 0.0135643 | 0.125943 |
| 0.78 | 0.0137824 | 0.127721 |
| 0.79 | 0.0140015 | 0.129502 |
| 0.8 | 0.0142218 | 0.131287 |
| 0.81 | 0.0144431 | 0.133076 |
| 0.82 | 0.0146656 | 0.134868 |
| 0.83 | 0.0148891 | 0.136665 |
| 0.84 | 0.0151138 | 0.138465 |
| 0.85 | 0.0153395 | 0.140269 |
| 0.86 | 0.0155664 | 0.142077 |
| 0.87 | 0.0157945 | 0.143889 |
| 0.88 | 0.0160236 | 0.145705 |
| 0.89 | 0.016254 | 0.147524 |
| 0.9 | 0.0164854 | 0.149348 |
| 0.91 | 0.016718 | 0.151175 |
| 0.92 | 0.0169518 | 0.153007 |
| 0.93 | 0.0171867 | 0.154842 |
| 0.94 | 0.0174228 | 0.156681 |
| 0.95 | 0.01766 | 0.158524 |
| 0.96 | 0.0178985 | 0.160371 |
| 0.97 | 0.0181381 | 0.162222 |
| 0.98 | 0.0183789 | 0.164076 |
| 0.99 | 0.0186209 | 0.165935 |
| 1 | 0.0188642 | 0.167798 |
| 1.01 | 0.0191086 | 0.169664 |
| 1.02 | 0.0193542 | 0.171535 |
| 1.03 | 0.0196011 | 0.173409 |
| 1.04 | 0.0198492 | 0.175288 |
| 1.05 | 0.0200985 | 0.17717 |
| 1.06 | 0.0203491 | 0.179057 |
| 1.07 | 0.0206009 | 0.180947 |
| 1.08 | 0.020854 | 0.182842 |
| 1.09 | 0.0211083 | 0.18474 |
| 1.1 | 0.0213639 | 0.186643 |
| 1.11 | 0.0216207 | 0.188549 |
| 1.12 | 0.0218789 | 0.19046 |
| 1.13 | 0.0221383 | 0.192375 |
| 1.14 | 0.022399 | 0.194293 |
| 1.15 | 0.022661 | 0.196216 |
| 1.16 | 0.0229243 | 0.198143 |
| 1.17 | 0.0231889 | 0.200073 |
| 1.18 | 0.0234548 | 0.202008 |
| 1.19 | 0.023722 | 0.203947 |
| 1.2 | 0.0239906 | 0.20589 |
| 1.21 | 0.0242605 | 0.207837 |
| 1.22 | 0.0245318 | 0.209789 |
| 1.23 | 0.0248044 | 0.211744 |
| 1.24 | 0.0250783 | 0.213703 |
| 1.25 | 0.0253536 | 0.215667 |
| 1.26 | 0.0256303 | 0.217635 |
| 1.27 | 0.0259083 | 0.219606 |
| 1.28 | 0.0261878 | 0.221582 |
| 1.29 | 0.0264686 | 0.223562 |
| 1.3 | 0.0267508 | 0.225547 |
| 1.31 | 0.0270344 | 0.227535 |
| 1.32 | 0.0273194 | 0.229528 |
| 1.33 | 0.0276059 | 0.231524 |
| 1.34 | 0.0278937 | 0.233525 |
| 1.35 | 0.028183 | 0.23553 |
| 1.36 | 0.0284737 | 0.23754 |
| 1.37 | 0.0287659 | 0.239553 |
| 1.38 | 0.0290595 | 0.241571 |
| 1.39 | 0.0293546 | 0.243593 |
| 1.4 | 0.0296511 | 0.245619 |
| 1.41 | 0.0299491 | 0.247649 |
| 1.42 | 0.0302486 | 0.249684 |
| 1.43 | 0.0305496 | 0.251723 |
| 1.44 | 0.0308521 | 0.253766 |
| 1.45 | 0.031156 | 0.255813 |
| 1.46 | 0.0314615 | 0.257865 |
| 1.47 | 0.0317685 | 0.259921 |
| 1.48 | 0.032077 | 0.261981 |
| 1.49 | 0.0323871 | 0.264045 |
| 1.5 | 0.0326986 | 0.266114 |
| 1.51 | 0.0330118 | 0.268187 |
| 1.52 | 0.0333264 | 0.270264 |
| 1.53 | 0.0336427 | 0.272346 |
| 1.54 | 0.0339605 | 0.274432 |
| 1.55 | 0.0342799 | 0.276522 |
| 1.56 | 0.0346009 | 0.278617 |
| 1.57 | 0.0349234 | 0.280716 |
| 1.58 | 0.0352476 | 0.282819 |
| 1.59 | 0.0355733 | 0.284926 |
| 1.6 | 0.0359007 | 0.287038 |
| 1.61 | 0.0362297 | 0.289155 |
| 1.62 | 0.0365604 | 0.291275 |
| 1.63 | 0.0368926 | 0.293401 |
| 1.64 | 0.0372266 | 0.29553 |
| 1.65 | 0.0375621 | 0.297664 |
| 1.66 | 0.0378994 | 0.299802 |
| 1.67 | 0.0382383 | 0.301945 |
| 1.68 | 0.0385789 | 0.304092 |
| 1.69 | 0.0389212 | 0.306244 |
| 1.7 | 0.0392651 | 0.308399 |
| 1.71 | 0.0396108 | 0.31056 |
| 1.72 | 0.0399582 | 0.312725 |
| 1.73 | 0.0403073 | 0.314894 |
| 1.74 | 0.0406582 | 0.317068 |
| 1.75 | 0.0410107 | 0.319246 |
| 1.76 | 0.0413651 | 0.321429 |
| 1.77 | 0.0417212 | 0.323616 |
| 1.78 | 0.042079 | 0.325807 |
| 1.79 | 0.0424386 | 0.328003 |
| 1.8 | 0.0428 | 0.330204 |
| 1.81 | 0.0431632 | 0.332409 |
| 1.82 | 0.0435282 | 0.334619 |
| 1.83 | 0.043895 | 0.336833 |
| 1.84 | 0.0442636 | 0.339051 |
| 1.85 | 0.044634 | 0.341275 |
| 1.86 | 0.0450063 | 0.343502 |
| 1.87 | 0.0453804 | 0.345735 |
| 1.88 | 0.0457564 | 0.347971 |
| 1.89 | 0.0461342 | 0.350213 |
| 1.9 | 0.0465139 | 0.352459 |
| 1.91 | 0.0468954 | 0.354709 |
| 1.92 | 0.0472789 | 0.356964 |
| 1.93 | 0.0476643 | 0.359224 |
| 1.94 | 0.0480515 | 0.361488 |
| 1.95 | 0.0484407 | 0.363757 |
| 1.96 | 0.0488318 | 0.366031 |
| 1.97 | 0.0492248 | 0.368309 |
| 1.98 | 0.0496198 | 0.370592 |
| 1.99 | 0.0500168 | 0.372879 |

V3 Total fitness

|  | Male biased | Equal | Female biased |
| --- | --- | --- | --- |
| 1 | 0.165319 | 0.166 | 0.1667 |
| 2 | 0.372692 | 0.3734 | 0.3741 |
| 3 | 0.626476 | 0.6272 | 0.6279 |
| 4 | 0.933358 | 0.9341 | 0.9348 |
| 5 | 1.29912 | 1.2998 | 1.3005 |
| 6 | 1.72761 | 1.7283 | 1.729 |
| 7 | 2.21959 | 2.2203 | 2.221 |
| 8 | 2.77155 | 2.7723 | 2.773 |
| 9 | 3.375 | 3.3757 | 3.3764 |
| 10 | 4.01637 | 4.0171 | 4.0178 |
| 11 | 4.67788 | 4.6786 | 4.6793 |
| 12 | 5.98283 | 5.9835 | 5.9842 |
| 13 | 7.45602 | 7.4567 | 7.4574 |
| 14 | 8.76248 | 8.7632 | 8.7639 |
| 15 | 9.80328 | 9.804 | 9.8047 |
| 16 | 10.5639 | 10.5646 | 10.5653 |
| 17 | 11.0853 | 11.086 | 11.0867 |
| 18 | 11.4273 | 11.428 | 11.4288 |
| 19 | 11.6452 | 11.646 | 11.6467 |
| 20 | 11.8157 | 11.8164 | 11.8171 |
| 21 | 12.0567 | 12.0574 | 12.0581 |
| 22 | 12.3897 | 12.3904 | 12.3911 |
| 23 | 12.8216 | 12.8223 | 12.823 |
| 24 | 13.3453 | 13.346 | 13.3467 |
| 25 | 13.9414 | 13.9421 | 13.9429 |
| 26 | 14.9145 | 14.9152 | 14.9159 |
| 27 | 16.2792 | 16.2799 | 16.2806 |
| 28 | 17.5274 | 17.5282 | 17.5289 |
| 29 | 18.64 | 18.6407 | 18.6414 |
| 30 | 19.609 | 19.6097 | 19.6105 |
| 31 | 20.4362 | 20.4369 | 20.4376 |
| 32 | 21.1302 | 21.1309 | 21.1316 |
| 33 | 21.7041 | 21.7048 | 21.7055 |
| 34 | 22.1731 | 22.1738 | 22.1745 |
| 35 | 22.5525 | 22.5532 | 22.5539 |
| 36 | 22.8572 | 22.8579 | 22.8586 |
| 37 | 23.1001 | 23.1008 | 23.1016 |
| 38 | 23.293 | 23.2937 | 23.2944 |
| 39 | 23.489 | 23.4897 | 23.4904 |
| 40 | 23.9264 | 23.9271 | 23.9278 |
| 41 | 25.0654 | 25.0661 | 25.0668 |
| 42 | 26.2924 | 26.2931 | 26.2938 |
| 43 | 27.4287 | 27.4294 | 27.4301 |
| 44 | 28.4698 | 28.4705 | 28.4713 |
| 45 | 29.4148 | 29.4155 | 29.4162 |
| 46 | 30.2649 | 30.2657 | 30.2664 |
| 47 | 31.0238 | 31.0245 | 31.0253 |
| 48 | 31.6965 | 31.6972 | 31.6979 |
| 49 | 32.2891 | 32.2898 | 32.2905 |
| 50 | 32.8082 | 32.8089 | 32.8096 |
| 51 | 33.2608 | 33.2615 | 33.2623 |
| 52 | 33.6538 | 33.6545 | 33.6552 |
| 53 | 33.9938 | 33.9945 | 33.9952 |
| 54 | 34.287 | 34.2877 | 34.2884 |
| 55 | 34.5391 | 34.5398 | 34.5405 |
| 56 | 35.0574 | 35.0581 | 34.7569 |
| 57 | 36.2053 | 36.2061 | 35.0759 |
| 58 | 37.2825 | 37.2832 | 35.721 |
| 59 | 38.2876 | 38.2883 | 36.385 |
| 60 | 39.2205 | 39.2213 | 37.0424 |
| 61 | 40.0823 | 40.083 | 37.6813 |
| 62 | 40.8748 | 40.8756 | 38.295 |
| 63 | 41.6006 | 41.6014 | 38.8796 |
| 64 | 42.2628 | 42.2636 | 39.4329 |
| 65 | 42.8649 | 42.8657 | 39.954 |
| 66 | 43.4107 | 43.4114 | 40.4425 |
| 67 | 43.9039 | 43.9046 | 40.8989 |
| 68 | 44.3486 | 44.3493 | 41.3239 |
| 69 | 44.7486 | 44.7493 | 41.7186 |
| 70 | 45.1075 | 45.1083 | 42.0841 |
| 71 | 45.4291 | 45.4299 | 42.422 |
| 72 | 46.0759 | 46.0766 | 42.7336 |
| 73 | 47.1172 | 47.1179 | 43.031 |
| 74 | 48.1006 | 48.1013 | 43.6646 |
| 75 | 49.0263 | 49.027 | 44.283 |
| 76 | 49.8951 | 49.8958 | 44.8842 |
| 77 | 50.7081 | 50.7088 | 45.467 |
| 78 | 51.4669 | 51.4676 | 46.0301 |
| 79 | 52.1732 | 52.1739 | 46.5727 |
| 80 | 52.8292 | 52.8299 | 47.0944 |
| 81 | 53.4371 | 53.4378 | 47.5949 |
| 82 | 53.9993 | 54 | 48.0739 |
| 83 | 54.5182 | 54.5189 | 48.5316 |
| 84 | 54.9964 | 54.9971 | 48.9682 |
| 85 | 55.4364 | 55.4371 | 49.3839 |
| 86 | 55.8406 | 55.8413 | 49.7792 |
| 87 | 56.2114 | 56.2121 | 50.1546 |
| 88 | 56.9422 | 56.5519 | 50.5105 |
| 89 | 57.9111 | 56.8629 | 50.8766 |
| 90 | 58.8321 | 57.1473 | 51.4721 |
| 91 | 59.7056 | 57.5208 | 52.0539 |
| 92 | 60.5324 | 58.1755 | 52.6215 |
| 93 | 61.3135 | 58.8134 | 53.1743 |
| 94 | 62.0502 | 59.4304 | 53.7119 |
| 95 | 62.7437 | 60.0246 | 54.2339 |
| 96 | 63.3955 | 60.5947 | 54.7402 |
| 97 | 64.0073 | 61.14 | 55.2305 |
| 98 | 64.5806 | 61.6606 | 55.7049 |
| 99 | 65.1173 | 62.1564 | 56.1633 |
| 100 | 65.6189 | 62.6278 | 56.6058 |
| 101 | 66.0873 | 63.0752 | 57.0325 |
| 102 | 66.5242 | 63.4994 | 57.4435 |
| 103 | 66.9312 | 63.9009 | 57.8392 |
| 104 | 67.3101 | 64.2805 | 58.2196 |
| 105 | 68.6378 | 64.6391 | 58.6408 |
| 106 | 69.5147 | 64.9774 | 59.211 |
| 107 | 70.3514 | 65.4166 | 59.7696 |
| 108 | 71.1487 | 66.0218 | 60.3163 |
| 109 | 71.9074 | 66.6113 | 60.8509 |
| 110 | 72.6285 | 67.1844 | 61.3731 |
| 111 | 73.3129 | 67.7405 | 61.8828 |
| 112 | 73.9618 | 68.2793 | 62.3798 |
| 113 | 74.5764 | 68.8005 | 62.8642 |
| 114 | 75.1578 | 69.3041 | 63.3358 |
| 115 | 75.7074 | 69.79 | 63.7946 |
| 116 | 76.2263 | 70.2584 | 64.2406 |
| 117 | 76.7158 | 70.7094 | 64.674 |
| 118 | 77.1773 | 71.1431 | 65.0947 |
| 119 | 77.6119 | 71.56 | 65.5029 |
| 120 | 78.021 | 71.9602 | 65.8988 |
| 121 | 78.4058 | 72.3442 | 66.3618 |
| 122 | 78.7675 | 72.7124 | 66.9144 |
| 123 | 79.1073 | 73.1981 | 67.4571 |
| 124 | 79.4262 | 73.7727 | 67.9897 |
| 125 | 79.9454 | 74.3345 | 68.5121 |
| 126 | 80.5647 | 74.8831 | 69.024 |
| 127 | 81.1651 | 75.4183 | 69.5255 |
| 128 | 81.7457 | 75.9399 | 70.0165 |
| 129 | 82.3063 | 76.4478 | 70.4968 |
| 130 | 82.8467 | 76.942 | 70.9665 |
| 131 | 83.3669 | 77.4224 | 71.4256 |
| 132 | 83.8671 | 77.889 | 71.8741 |
| 133 | 84.3476 | 78.342 | 72.3119 |
| 134 | 84.8088 | 78.7814 | 72.7392 |
| 135 | 85.2509 | 79.2074 | 73.1559 |
| 136 | 85.6746 | 79.62 | 73.5622 |
| 137 | 86.0802 | 80.0195 | 74.0575 |
| 138 | 86.4682 | 80.4061 | 74.5973 |
| 139 | 86.8392 | 80.9256 | 75.1285 |
| 140 | 87.1938 | 81.4803 | 75.651 |
| 141 | 87.7526 | 82.0242 | 76.1645 |
| 142 | 88.333 | 82.5571 | 76.6692 |
| 143 | 88.8988 | 83.079 | 77.1648 |
| 144 | 89.4497 | 83.5897 | 77.6514 |
| 145 | 89.9857 | 84.0891 | 78.1289 |
| 146 | 90.5066 | 84.5773 | 78.5973 |
| 147 | 91.0124 | 85.0541 | 79.0565 |
| 148 | 91.5031 | 85.5197 | 79.5067 |
| 149 | 91.9789 | 85.974 | 79.9477 |
| 150 | 92.4399 | 86.417 | 80.3797 |
| 151 | 92.8862 | 86.8489 | 80.8026 |
| 152 | 93.318 | 87.2698 | 81.2165 |
| 153 | 93.7357 | 87.6797 | 81.7372 |
| 154 | 94.1393 | 88.0787 | 82.2673 |
| 155 | 94.5292 | 88.6239 | 82.7898 |
| 156 | 94.92 | 89.1645 | 83.3046 |
| 157 | 95.4887 | 89.6959 | 83.8116 |
| 158 | 96.0459 | 90.218 | 84.3107 |
| 159 | 96.5912 | 90.7306 | 84.8019 |
| 160 | 97.1245 | 91.2336 | 85.2852 |
| 161 | 97.6458 | 91.7272 | 85.7606 |
| 162 | 98.155 | 92.2112 | 86.228 |
| 163 | 98.6521 | 92.6855 | 86.6874 |
| 164 | 99.1371 | 93.1503 | 87.1388 |
| 165 | 99.6099 | 93.6055 | 87.5823 |
| 166 | 100.071 | 94.0512 | 88.0178 |
| 167 | 100.52 | 94.4873 | 88.4453 |
| 168 | 100.957 | 94.914 | 88.865 |
| 169 | 101.382 | 95.3313 | 89.2768 |
| 170 | 101.796 | 95.7392 | 89.6807 |
| 171 | 102.198 | 96.1379 | 90.0769 |
| 172 | 102.59 | 96.5274 | 90.4653 |
| 173 | 102.97 | 96.9079 | 90.846 |
| 174 | 103.339 | 97.2793 | 91.2192 |
| 175 | 103.697 | 97.6419 | 91.5847 |
| 176 | 104.045 | 97.9958 | 91.9428 |
| 177 | 104.382 | 98.341 | 92.2935 |
| 178 | 104.71 | 98.6777 | 92.6369 |
| 179 | 105.027 | 99.006 | 92.9729 |
| 180 | 105.335 | 99.326 | 93.3019 |
| 181 | 105.633 | 99.6379 | 93.6237 |
| 182 | 105.922 | 99.9418 | 93.9386 |
| 183 | 106.202 | 100.238 | 94.2466 |
| 184 | 106.473 | 100.526 | 94.5477 |
| 185 | 106.736 | 100.807 | 94.8422 |
| 186 | 106.99 | 101.08 | 95.13 |
| 187 | 107.236 | 101.346 | 95.4113 |
| 188 | 107.474 | 101.605 | 95.6862 |
| 189 | 107.704 | 101.857 | 95.9548 |
| 190 | 107.927 | 102.102 | 96.2172 |
| 191 | 108.142 | 102.34 | 96.4735 |
| 192 | 108.35 | 102.572 | 96.7238 |
| 193 | 108.551 | 102.797 | 96.9681 |
| 194 | 108.746 | 103.016 | 97.2067 |
| 195 | 108.934 | 103.229 | 97.4396 |
| 196 | 109.115 | 103.436 | 97.6668 |
| 197 | 109.29 | 103.637 | 97.8886 |
| 198 | 109.46 | 103.832 | 98.105 |
| 199 | 109.623 | 104.022 | 98.3162 |
| 200 | 109.781 | 104.206 | 98.5221 |

V3 Type-II-biased

|  | Male | Female |
| --- | --- | --- |
| 1 | 3.43631e-10 | 1 |
| 2 | 7.01732e-10 | 1 |
| 3 | 1.53453e-09 | 1 |
| 4 | 7.6379e-10 | 1 |
| 5 | 6.43688e-10 | 1 |
| 6 | 4.60793e-10 | 1 |
| 7 | 3.46919e-10 | 1 |
| 8 | 2.73416e-10 | 1 |
| 9 | 2.25067e-10 | 1 |
| 10 | 4.8316e-11 | 1 |
| 11 | 8.64662e-10 | 1 |
| 12 | 1 | 1.65359e-10 |
| 13 | 1 | 1.63619e-10 |
| 14 | 1 | 1.8622e-10 |
| 15 | 1 | 2.41058e-10 |
| 16 | 1 | 3.52937e-10 |
| 17 | 1 | 1.1927e-10 |
| 18 | 1 | 5.17338e-11 |
| 19 | 1 | 3.07634e-10 |
| 20 | 0.933364 | 0.0666363 |
| 21 | 0.852172 | 0.147828 |
| 22 | 0.783116 | 0.216884 |
| 23 | 0.725634 | 0.274366 |
| 24 | 0.678813 | 0.321187 |
| 25 | 0.641288 | 0.358712 |
| 26 | 1 | 3.65331e-12 |
| 27 | 1 | 7.65998e-13 |
| 28 | 1 | 2.07558e-13 |
| 29 | 1 | 9.28807e-13 |
| 30 | 1 | 1.07108e-12 |
| 31 | 1 | 1.27239e-12 |
| 32 | 1 | 1.5582e-12 |
| 33 | 1 | 1.97291e-12 |
| 34 | 1 | 2.60031e-12 |
| 35 | 1 | 3.61872e-12 |
| 36 | 1 | 2.19437e-11 |
| 37 | 1 | 3.9357e-11 |
| 38 | 1 | 2.41402e-11 |
| 39 | 0.90495 | 0.0950502 |
| 40 | 0.822746 | 0.177254 |
| 41 | 1 | 2.61331e-12 |
| 42 | 1 | 2.76744e-12 |
| 43 | 1 | 5.94437e-13 |
| 44 | 1 | 2.58873e-12 |
| 45 | 1 | 3.57028e-12 |
| 46 | 1 | 3.99144e-12 |
| 47 | 1 | 3.61757e-12 |
| 48 | 1 | 5.19399e-12 |
| 49 | 1 | 1.2109e-12 |
| 50 | 1 | 1.43495e-12 |
| 51 | 1 | 1.73354e-12 |
| 52 | 1 | 2.14483e-12 |
| 53 | 1 | 2.73855e-12 |
| 54 | 1 | 3.65728e-12 |
| 55 | 1 | 5.24522e-12 |
| 56 | 1 | 2.07558e-12 |
| 57 | 1 | 1.74951e-12 |
| 58 | 1 | 2.32203e-12 |
| 59 | 1 | 4.96851e-13 |
| 60 | 1 | 2.14217e-12 |
| 61 | 1 | 2.90768e-12 |
| 62 | 1 | 3.18097e-12 |
| 63 | 1 | 2.8051e-12 |
| 64 | 1 | 3.8955e-12 |
| 65 | 1 | 1.74551e-11 |
| 66 | 1 | 9.86455e-13 |
| 67 | 1 | 5.63046e-12 |
| 68 | 1 | 5.20063e-12 |
| 69 | 1 | 3.04262e-11 |
| 70 | 1 | 7.23746e-12 |
| 71 | 1 | 8.78835e-12 |
| 72 | 1 | 3.62387e-11 |
| 73 | 1 | 3.81353e-11 |
| 74 | 1 | 8.06493e-12 |
| 75 | 1 | 8.56868e-12 |
| 76 | 1 | 9.14708e-12 |
| 77 | 1 | 9.81101e-12 |
| 78 | 1 | 5.28688e-11 |
| 79 | 1 | 2.29036e-12 |
| 80 | 1 | 6.2328e-11 |
| 81 | 1 | 1.36412e-11 |
| 82 | 1 | 1.50118e-11 |
| 83 | 1 | 1.66207e-11 |
| 84 | 1 | 3.70504e-12 |
| 85 | 1 | 1.04012e-10 |
| 86 | 1 | 2.35599e-11 |
| 87 | 1 | 5.3901e-12 |
| 88 | 1 | 3.23591e-11 |
| 89 | 1 | 3.39337e-11 |
| 90 | 1 | 3.57029e-11 |
| 91 | 1 | 3.76879e-11 |
| 92 | 1 | 3.99146e-11 |
| 93 | 1 | 4.24129e-11 |
| 94 | 1 | 4.52199e-11 |
| 95 | 1 | 4.8378e-11 |
| 96 | 1 | 5.194e-11 |
| 97 | 1 | 5.59699e-11 |
| 98 | 1 | 1.2109e-09 |
| 99 | 1 | 2.63055e-10 |
| 100 | 1 | 1.434e-11 |
| 101 | 1 | 7.87129e-11 |
| 102 | 1 | 3.47059e-10 |
| 103 | 1 | 1.92233e-11 |
| 104 | 1 | 1.07308e-10 |
| 105 | 1 | 6.11429e-10 |
| 106 | 1 | 3.19811e-09 |
| 107 | 1 | 3.36446e-09 |
| 108 | 1 | 1.76944e-08 |
| 109 | 1 | 3.73506e-11 |
| 110 | 1 | 7.90909e-10 |
| 111 | 1 | 8.37611e-12 |
| 112 | 1 | 8.904e-10 |
| 113 | 1 | 1.89823e-10 |
| 114 | 1 | 2.53615e-08 |
| 115 | 1 | 2.17565e-10 |
| 116 | 1 | 1.16959e-09 |
| 117 | 1 | 1.26203e-09 |
| 118 | 1 | 1.3667e-09 |
| 119 | 1 | 2.97179e-10 |
| 120 | 1 | 3.2452e-10 |
| 121 | 1 | 3.56063e-10 |
| 122 | 1 | 1.96417e-09 |
| 123 | 1 | 4.36127e-10 |
| 124 | 1 | 4.87723e-08 |
| 125 | 0.896721 | 0.103279 |
| 126 | 0.893918 | 0.106082 |
| 127 | 0.891479 | 0.108521 |
| 128 | 0.889309 | 0.110691 |
| 129 | 0.887347 | 0.112653 |
| 130 | 0.885554 | 0.114446 |
| 131 | 0.883898 | 0.116102 |
| 132 | 0.882359 | 0.117641 |
| 133 | 0.880918 | 0.119082 |
| 134 | 0.879564 | 0.120436 |
| 135 | 0.878284 | 0.121716 |
| 136 | 0.877071 | 0.122929 |
| 137 | 0.875916 | 0.124084 |
| 138 | 0.874815 | 0.125185 |
| 139 | 0.873761 | 0.126239 |
| 140 | 0.87275 | 0.12725 |
| 141 | 0.80363 | 0.19637 |
| 142 | 0.801092 | 0.198908 |
| 143 | 0.798713 | 0.201287 |
| 144 | 0.796475 | 0.203525 |
| 145 | 0.794361 | 0.205639 |
| 146 | 0.792358 | 0.207642 |
| 147 | 0.790453 | 0.209547 |
| 148 | 0.788638 | 0.211362 |
| 149 | 0.786903 | 0.213097 |
| 150 | 0.785243 | 0.214757 |
| 151 | 0.78365 | 0.21635 |
| 152 | 0.78212 | 0.21788 |
| 153 | 0.780646 | 0.219354 |
| 154 | 0.779226 | 0.220774 |
| 155 | 0.777855 | 0.222145 |
| 156 | 0.72892 | 0.27108 |
| 157 | 0.726571 | 0.273429 |
| 158 | 0.724326 | 0.275674 |
| 159 | 0.722177 | 0.277823 |
| 160 | 0.720116 | 0.279884 |
| 161 | 0.718137 | 0.281863 |
| 162 | 0.716233 | 0.283767 |
| 163 | 0.7144 | 0.2856 |
| 164 | 0.712632 | 0.287368 |
| 165 | 0.710924 | 0.289076 |
| 166 | 0.709274 | 0.290726 |
| 167 | 0.707677 | 0.292323 |
| 168 | 0.70613 | 0.29387 |
| 169 | 0.70463 | 0.29537 |
| 170 | 0.703174 | 0.296826 |
| 171 | 0.70176 | 0.29824 |
| 172 | 0.700386 | 0.299614 |
| 173 | 0.699048 | 0.300952 |
| 174 | 0.697747 | 0.302253 |
| 175 | 0.696478 | 0.303522 |
| 176 | 0.695242 | 0.304758 |
| 177 | 0.694036 | 0.305964 |
| 178 | 0.692858 | 0.307142 |
| 179 | 0.691709 | 0.308291 |
| 180 | 0.690585 | 0.309415 |
| 181 | 0.689487 | 0.310513 |
| 182 | 0.688412 | 0.311588 |
| 183 | 0.687361 | 0.312639 |
| 184 | 0.686331 | 0.313669 |
| 185 | 0.685322 | 0.314678 |
| 186 | 0.684334 | 0.315666 |
| 187 | 0.683365 | 0.316635 |
| 188 | 0.682415 | 0.317585 |
| 189 | 0.681482 | 0.318518 |
| 190 | 0.680567 | 0.319433 |
| 191 | 0.679669 | 0.320331 |
| 192 | 0.678786 | 0.321214 |
| 193 | 0.677919 | 0.322081 |
| 194 | 0.677067 | 0.322933 |
| 195 | 0.676229 | 0.323771 |
| 196 | 0.675406 | 0.324594 |
| 197 | 0.674595 | 0.325405 |
| 198 | 0.673798 | 0.326202 |
| 199 | 0.673013 | 0.326987 |
| 200 | 0.67224 | 0.32776 |

V3 Type-II-biased

|  | Male | Female |
| --- | --- | --- |
| 1 | 0 | 1 |
| 2 | 0 | 1 |
| 3 | 0 | 1 |
| 4 | 0 | 1 |
| 5 | 0 | 1 |
| 6 | 0 | 1 |
| 7 | 0 | 1 |
| 8 | 0 | 1 |
| 9 | 0 | 1 |
| 10 | 0 | 1 |
| 11 | 0 | 1 |
| 12 | 1 | 0 |
| 13 | 1 | 0 |
| 14 | 1 | 0 |
| 15 | 1 | 0 |
| 16 | 1 | 0 |
| 17 | 1 | 0 |
| 18 | 1 | 0 |
| 19 | 1 | 0 |
| 20 | 1 | 1 |
| 21 | 1 | 1 |
| 22 | 1 | 1 |
| 23 | 1 | 1 |
| 24 | 1 | 1 |
| 25 | 1 | 1 |
| 26 | 2 | 0 |
| 27 | 2 | 0 |
| 28 | 2 | 0 |
| 29 | 2 | 0 |
| 30 | 2 | 0 |
| 31 | 2 | 0 |
| 32 | 2 | 0 |
| 33 | 2 | 0 |
| 34 | 2 | 0 |
| 35 | 2 | 0 |
| 36 | 2 | 0 |
| 37 | 2 | 0 |
| 38 | 2 | 0 |
| 39 | 2 | 1 |
| 40 | 2 | 1 |
| 41 | 3 | 0 |
| 42 | 3 | 0 |
| 43 | 3 | 0 |
| 44 | 3 | 0 |
| 45 | 3 | 0 |
| 46 | 3 | 0 |
| 47 | 3 | 0 |
| 48 | 3 | 0 |
| 49 | 3 | 0 |
| 50 | 3 | 0 |
| 51 | 3 | 0 |
| 52 | 3 | 0 |
| 53 | 3 | 0 |
| 54 | 3 | 0 |
| 55 | 3 | 0 |
| 56 | 4 | 0 |
| 57 | 4 | 0 |
| 58 | 4 | 0 |
| 59 | 4 | 0 |
| 60 | 4 | 0 |
| 61 | 4 | 0 |
| 62 | 4 | 0 |
| 63 | 4 | 0 |
| 64 | 4 | 0 |
| 65 | 4 | 0 |
| 66 | 4 | 0 |
| 67 | 4 | 0 |
| 68 | 4 | 0 |
| 69 | 4 | 0 |
| 70 | 4 | 0 |
| 71 | 4 | 0 |
| 72 | 5 | 0 |
| 73 | 5 | 0 |
| 74 | 5 | 0 |
| 75 | 5 | 0 |
| 76 | 5 | 0 |
| 77 | 5 | 0 |
| 78 | 5 | 0 |
| 79 | 5 | 0 |
| 80 | 5 | 0 |
| 81 | 5 | 0 |
| 82 | 5 | 0 |
| 83 | 5 | 0 |
| 84 | 5 | 0 |
| 85 | 5 | 0 |
| 86 | 5 | 0 |
| 87 | 5 | 0 |
| 88 | 6 | 0 |
| 89 | 6 | 0 |
| 90 | 6 | 0 |
| 91 | 6 | 0 |
| 92 | 6 | 0 |
| 93 | 6 | 0 |
| 94 | 6 | 0 |
| 95 | 6 | 0 |
| 96 | 6 | 0 |
| 97 | 6 | 0 |
| 98 | 6 | 0 |
| 99 | 6 | 0 |
| 100 | 6 | 0 |
| 101 | 6 | 0 |
| 102 | 6 | 0 |
| 103 | 6 | 0 |
| 104 | 6 | 0 |
| 105 | 7 | 0 |
| 106 | 7 | 0 |
| 107 | 7 | 0 |
| 108 | 7 | 0 |
| 109 | 7 | 0 |
| 110 | 7 | 0 |
| 111 | 7 | 0 |
| 112 | 7 | 0 |
| 113 | 7 | 0 |
| 114 | 7 | 0 |
| 115 | 7 | 0 |
| 116 | 7 | 0 |
| 117 | 7 | 0 |
| 118 | 7 | 0 |
| 119 | 7 | 0 |
| 120 | 7 | 0 |
| 121 | 7 | 0 |
| 122 | 7 | 0 |
| 123 | 7 | 0 |
| 124 | 7 | 0 |
| 125 | 7 | 1 |
| 126 | 7 | 1 |
| 127 | 7 | 1 |
| 128 | 7 | 1 |
| 129 | 7 | 1 |
| 130 | 7 | 1 |
| 131 | 7 | 1 |
| 132 | 7 | 1 |
| 133 | 7 | 1 |
| 134 | 7 | 1 |
| 135 | 7 | 1 |
| 136 | 7 | 1 |
| 137 | 7 | 1 |
| 138 | 7 | 1 |
| 139 | 7 | 1 |
| 140 | 7 | 1 |
| 141 | 7 | 2 |
| 142 | 7 | 2 |
| 143 | 7 | 2 |
| 144 | 7 | 2 |
| 145 | 7 | 2 |
| 146 | 7 | 2 |
| 147 | 7 | 2 |
| 148 | 7 | 2 |
| 149 | 7 | 2 |
| 150 | 7 | 2 |
| 151 | 7 | 2 |
| 152 | 7 | 2 |
| 153 | 7 | 2 |
| 154 | 7 | 2 |
| 155 | 7 | 2 |
| 156 | 7 | 3 |
| 157 | 7 | 3 |
| 158 | 7 | 3 |
| 159 | 7 | 3 |
| 160 | 7 | 3 |
| 161 | 7 | 3 |
| 162 | 7 | 3 |
| 163 | 7 | 3 |
| 164 | 7 | 3 |
| 165 | 7 | 3 |
| 166 | 7 | 3 |
| 167 | 7 | 3 |
| 168 | 7 | 3 |
| 169 | 7 | 3 |
| 170 | 7 | 3 |
| 171 | 7 | 3 |
| 172 | 7 | 3 |
| 173 | 7 | 3 |
| 174 | 7 | 3 |
| 175 | 7 | 3 |
| 176 | 7 | 3 |
| 177 | 7 | 3 |
| 178 | 7 | 3 |
| 179 | 7 | 3 |
| 180 | 7 | 3 |
| 181 | 7 | 3 |
| 182 | 7 | 3 |
| 183 | 7 | 3 |
| 184 | 7 | 3 |
| 185 | 7 | 3 |
| 186 | 7 | 3 |
| 187 | 7 | 3 |
| 188 | 7 | 3 |
| 189 | 7 | 3 |
| 190 | 7 | 3 |
| 191 | 7 | 3 |
| 192 | 7 | 3 |
| 193 | 7 | 3 |
| 194 | 7 | 3 |
| 195 | 7 | 3 |
| 196 | 7 | 3 |
| 197 | 7 | 3 |
| 198 | 7 | 3 |
| 199 | 7 | 3 |
| 200 | 7 | 3 |

V3 Equal

|  | Male | Female |
| --- | --- | --- |
| 1 | 2.415e-10 | 1 |
| 2 | 1.5775e-09 | 1 |
| 3 | 2.7e-10 | 1 |
| 4 | 6.7375e-10 | 1 |
| 5 | 4.55e-10 | 1 |
| 6 | 3.26667e-10 | 1 |
| 7 | 2.45714e-10 | 1 |
| 8 | 1.94375e-10 | 1 |
| 9 | 1.6e-10 | 1 |
| 10 | 1.375e-10 | 1 |
| 11 | 6.13636e-10 | 1 |
| 12 | 1 | 6.875e-11 |
| 13 | 1 | 2.71923e-10 |
| 14 | 1 | 3.09286e-10 |
| 15 | 1 | 4e-10 |
| 16 | 1 | 5.84375e-10 |
| 17 | 1 | 1.96176e-10 |
| 18 | 1 | 8.38889e-11 |
| 19 | 1 | 1.90526e-09 |
| 20 | 0.933365 | 0.066635 |
| 21 | 0.852171 | 0.147829 |
| 22 | 0.783118 | 0.216882 |
| 23 | 0.725635 | 0.274365 |
| 24 | 0.678813 | 0.321188 |
| 25 | 0.641288 | 0.358712 |
| 26 | 1 | 1.21731e-12 |
| 27 | 1 | 6.38889e-12 |
| 28 | 1 | 6.91071e-12 |
| 29 | 1 | 7.74138e-12 |
| 30 | 1 | 1.78333e-12 |
| 31 | 1 | 2.1129e-12 |
| 32 | 1 | 1.29844e-11 |
| 33 | 1 | 3.28788e-12 |
| 34 | 1 | 3.47059e-12 |
| 35 | 1 | 4.82857e-12 |
| 36 | 1 | 7.31944e-12 |
| 37 | 1 | 5.24324e-11 |
| 38 | 1 | 2.01316e-10 |
| 39 | 0.904949 | 0.0950513 |
| 40 | 0.822745 | 0.177255 |
| 41 | 1 | 4.35366e-12 |
| 42 | 1 | 4.60714e-12 |
| 43 | 0.999998 | 4.95349e-12 |
| 44 | 1 | 1.07841e-12 |
| 45 | 1 | 5.95556e-12 |
| 46 | 0.999998 | 1.32609e-12 |
| 47 | 1 | 1.51064e-12 |
| 48 | 1 | 1.72917e-12 |
| 49 | 0.999998 | 2.02041e-12 |
| 50 | 1 | 2.39e-12 |
| 51 | 1 | 1.15686e-11 |
| 52 | 0.999998 | 3.57692e-12 |
| 53 | 1 | 9.13208e-11 |
| 54 | 1 | 6.09259e-12 |
| 55 | 0.999998 | 1.74545e-10 |
| 56 | 1 | 1.38393e-11 |
| 57 | 1 | 1.45614e-11 |
| 58 | 1 | 7.74138e-11 |
| 59 | 1 | 8.27966e-11 |
| 60 | 1 | 8.91667e-13 |
| 61 | 1 | 9.67213e-11 |
| 62 | 1 | 8.46774e-11 |
| 63 | 1 | 2.34127e-11 |
| 64 | 1 | 5.19531e-12 |
| 65 | 1 | 2.90769e-11 |
| 66 | 1 | 3.28788e-11 |
| 67 | 1 | 7.53731e-12 |
| 68 | 1 | 4.33088e-11 |
| 69 | 1 | 1.01449e-11 |
| 70 | 1 | 6.02857e-11 |
| 71 | 1 | 5.85915e-11 |
| 72 | 1 | 6.04167e-11 |
| 73 | 1 | 1.58904e-11 |
| 74 | 1 | 6.72297e-11 |
| 75 | 1 | 1.42667e-11 |
| 76 | 1 | 7.63158e-11 |
| 77 | 1 | 8.18182e-11 |
| 78 | 1 | 8.78205e-11 |
| 79 | 1 | 2.38608e-11 |
| 80 | 1 | 1.0375e-10 |
| 81 | 1 | 1.1358e-10 |
| 82 | 1 | 1.25e-10 |
| 83 | 1 | 1.72892e-10 |
| 84 | 1 | 1.54167e-10 |
| 85 | 1 | 6.94118e-10 |
| 86 | 1 | 1.96512e-10 |
| 87 | 1 | 4.49425e-11 |
| 88 | 1 | 8.29545e-10 |
| 89 | 1 | 3.05618e-10 |
| 90 | 1 | 1.17222e-09 |
| 91 | 0.872456 | 0.127546 |
| 92 | 0.866071 | 0.13393 |
| 93 | 0.861027 | 0.138972 |
| 94 | 0.856819 | 0.143183 |
| 95 | 0.853179 | 0.14682 |
| 96 | 0.849969 | 0.150032 |
| 97 | 0.847082 | 0.152916 |
| 98 | 0.844464 | 0.155538 |
| 99 | 0.842056 | 0.157944 |
| 100 | 0.83983 | 0.160171 |
| 101 | 0.837752 | 0.162246 |
| 102 | 0.835814 | 0.164187 |
| 103 | 0.833985 | 0.166015 |
| 104 | 0.83226 | 0.167741 |
| 105 | 0.830624 | 0.169378 |
| 106 | 0.829066 | 0.170935 |
| 107 | 0.75135 | 0.248652 |
| 108 | 0.747412 | 0.252589 |
| 109 | 0.743771 | 0.256229 |
| 110 | 0.740382 | 0.259618 |
| 111 | 0.737216 | 0.262782 |
| 112 | 0.734246 | 0.265752 |
| 113 | 0.731451 | 0.26855 |
| 114 | 0.728807 | 0.271193 |
| 115 | 0.726304 | 0.273697 |
| 116 | 0.723922 | 0.276078 |
| 117 | 0.721658 | 0.278344 |
| 118 | 0.719492 | 0.280508 |
| 119 | 0.71742 | 0.282578 |
| 120 | 0.715437 | 0.284562 |
| 121 | 0.713533 | 0.286468 |
| 122 | 0.711701 | 0.2883 |
| 123 | 0.659191 | 0.340807 |
| 124 | 0.656141 | 0.343858 |
| 125 | 0.653236 | 0.346764 |
| 126 | 0.650468 | 0.349533 |
| 127 | 0.647823 | 0.352179 |
| 128 | 0.645289 | 0.35471 |
| 129 | 0.64286 | 0.357137 |
| 130 | 0.640535 | 0.359467 |
| 131 | 0.638294 | 0.361706 |
| 132 | 0.63614 | 0.363859 |
| 133 | 0.634064 | 0.365935 |
| 134 | 0.632063 | 0.367937 |
| 135 | 0.63013 | 0.369869 |
| 136 | 0.628261 | 0.371738 |
| 137 | 0.626456 | 0.373546 |
| 138 | 0.624703 | 0.375296 |
| 139 | 0.586669 | 0.413332 |
| 140 | 0.584211 | 0.415789 |
| 141 | 0.58184 | 0.418159 |
| 142 | 0.579556 | 0.420445 |
| 143 | 0.57735 | 0.422649 |
| 144 | 0.575219 | 0.424783 |
| 145 | 0.573155 | 0.426844 |
| 146 | 0.571158 | 0.428841 |
| 147 | 0.569224 | 0.430776 |
| 148 | 0.567348 | 0.432651 |
| 149 | 0.56553 | 0.43447 |
| 150 | 0.563763 | 0.436235 |
| 151 | 0.56205 | 0.437952 |
| 152 | 0.560382 | 0.439618 |
| 153 | 0.558758 | 0.441242 |
| 154 | 0.557179 | 0.442821 |
| 155 | 0.528277 | 0.471723 |
| 156 | 0.526253 | 0.473747 |
| 157 | 0.52429 | 0.47571 |
| 158 | 0.522383 | 0.477617 |
| 159 | 0.520528 | 0.479472 |
| 160 | 0.518725 | 0.481275 |
| 161 | 0.516972 | 0.483028 |
| 162 | 0.515265 | 0.484735 |
| 163 | 0.513604 | 0.486396 |
| 164 | 0.511985 | 0.488015 |
| 165 | 0.510409 | 0.489591 |
| 166 | 0.50887 | 0.49113 |
| 167 | 0.507368 | 0.492632 |
| 168 | 0.505905 | 0.494095 |
| 169 | 0.504473 | 0.495527 |
| 170 | 0.503076 | 0.496924 |
| 171 | 0.501711 | 0.498289 |
| 172 | 0.500375 | 0.499625 |
| 173 | 0.499069 | 0.500931 |
| 174 | 0.497793 | 0.502207 |
| 175 | 0.496543 | 0.503457 |
| 176 | 0.495318 | 0.504682 |
| 177 | 0.494121 | 0.505879 |
| 178 | 0.492947 | 0.507053 |
| 179 | 0.491796 | 0.508204 |
| 180 | 0.490667 | 0.509333 |
| 181 | 0.489561 | 0.510439 |
| 182 | 0.488475 | 0.511525 |
| 183 | 0.48741 | 0.51259 |
| 184 | 0.486364 | 0.513636 |
| 185 | 0.485338 | 0.514662 |
| 186 | 0.484331 | 0.515669 |
| 187 | 0.48334 | 0.51666 |
| 188 | 0.482367 | 0.517633 |
| 189 | 0.48141 | 0.51859 |
| 190 | 0.480471 | 0.519529 |
| 191 | 0.479547 | 0.520453 |
| 192 | 0.478638 | 0.521362 |
| 193 | 0.477744 | 0.522256 |
| 194 | 0.476863 | 0.523137 |
| 195 | 0.475995 | 0.524005 |
| 196 | 0.475143 | 0.524857 |
| 197 | 0.474305 | 0.525695 |
| 198 | 0.473477 | 0.526523 |
| 199 | 0.472663 | 0.527337 |
| 200 | 0.47186 | 0.52814 |

V3 Equal

|  | Male | Female |
| --- | --- | --- |
| 1 | 0 | 1 |
| 2 | 0 | 1 |
| 3 | 0 | 1 |
| 4 | 0 | 1 |
| 5 | 0 | 1 |
| 6 | 0 | 1 |
| 7 | 0 | 1 |
| 8 | 0 | 1 |
| 9 | 0 | 1 |
| 10 | 0 | 1 |
| 11 | 0 | 1 |
| 12 | 1 | 0 |
| 13 | 1 | 0 |
| 14 | 1 | 0 |
| 15 | 1 | 0 |
| 16 | 1 | 0 |
| 17 | 1 | 0 |
| 18 | 1 | 0 |
| 19 | 1 | 0 |
| 20 | 1 | 1 |
| 21 | 1 | 1 |
| 22 | 1 | 1 |
| 23 | 1 | 1 |
| 24 | 1 | 1 |
| 25 | 1 | 1 |
| 26 | 2 | 0 |
| 27 | 2 | 0 |
| 28 | 2 | 0 |
| 29 | 2 | 0 |
| 30 | 2 | 0 |
| 31 | 2 | 0 |
| 32 | 2 | 0 |
| 33 | 2 | 0 |
| 34 | 2 | 0 |
| 35 | 2 | 0 |
| 36 | 2 | 0 |
| 37 | 2 | 0 |
| 38 | 2 | 0 |
| 39 | 2 | 1 |
| 40 | 2 | 1 |
| 41 | 3 | 0 |
| 42 | 3 | 0 |
| 43 | 3 | 0 |
| 44 | 3 | 0 |
| 45 | 3 | 0 |
| 46 | 3 | 0 |
| 47 | 3 | 0 |
| 48 | 3 | 0 |
| 49 | 3 | 0 |
| 50 | 3 | 0 |
| 51 | 3 | 0 |
| 52 | 3 | 0 |
| 53 | 3 | 0 |
| 54 | 3 | 0 |
| 55 | 3 | 0 |
| 56 | 4 | 0 |
| 57 | 4 | 0 |
| 58 | 4 | 0 |
| 59 | 4 | 0 |
| 60 | 4 | 0 |
| 61 | 4 | 0 |
| 62 | 4 | 0 |
| 63 | 4 | 0 |
| 64 | 4 | 0 |
| 65 | 4 | 0 |
| 66 | 4 | 0 |
| 67 | 4 | 0 |
| 68 | 4 | 0 |
| 69 | 4 | 0 |
| 70 | 4 | 0 |
| 71 | 4 | 0 |
| 72 | 5 | 0 |
| 73 | 5 | 0 |
| 74 | 5 | 0 |
| 75 | 5 | 0 |
| 76 | 5 | 0 |
| 77 | 5 | 0 |
| 78 | 5 | 0 |
| 79 | 5 | 0 |
| 80 | 5 | 0 |
| 81 | 5 | 0 |
| 82 | 5 | 0 |
| 83 | 5 | 0 |
| 84 | 5 | 0 |
| 85 | 5 | 0 |
| 86 | 5 | 0 |
| 87 | 5 | 0 |
| 88 | 5 | 0 |
| 89 | 5 | 0 |
| 90 | 5 | 0 |
| 91 | 5 | 1 |
| 92 | 5 | 1 |
| 93 | 5 | 1 |
| 94 | 5 | 1 |
| 95 | 5 | 1 |
| 96 | 5 | 1 |
| 97 | 5 | 1 |
| 98 | 5 | 1 |
| 99 | 5 | 1 |
| 100 | 5 | 1 |
| 101 | 5 | 1 |
| 102 | 5 | 1 |
| 103 | 5 | 1 |
| 104 | 5 | 1 |
| 105 | 5 | 1 |
| 106 | 5 | 1 |
| 107 | 5 | 2 |
| 108 | 5 | 2 |
| 109 | 5 | 2 |
| 110 | 5 | 2 |
| 111 | 5 | 2 |
| 112 | 5 | 2 |
| 113 | 5 | 2 |
| 114 | 5 | 2 |
| 115 | 5 | 2 |
| 116 | 5 | 2 |
| 117 | 5 | 2 |
| 118 | 5 | 2 |
| 119 | 5 | 2 |
| 120 | 5 | 2 |
| 121 | 5 | 2 |
| 122 | 5 | 2 |
| 123 | 5 | 3 |
| 124 | 5 | 3 |
| 125 | 5 | 3 |
| 126 | 5 | 3 |
| 127 | 5 | 3 |
| 128 | 5 | 3 |
| 129 | 5 | 3 |
| 130 | 5 | 3 |
| 131 | 5 | 3 |
| 132 | 5 | 3 |
| 133 | 5 | 3 |
| 134 | 5 | 3 |
| 135 | 5 | 3 |
| 136 | 5 | 3 |
| 137 | 5 | 3 |
| 138 | 5 | 3 |
| 139 | 5 | 4 |
| 140 | 5 | 4 |
| 141 | 5 | 4 |
| 142 | 5 | 4 |
| 143 | 5 | 4 |
| 144 | 5 | 4 |
| 145 | 5 | 4 |
| 146 | 5 | 4 |
| 147 | 5 | 4 |
| 148 | 5 | 4 |
| 149 | 5 | 4 |
| 150 | 5 | 4 |
| 151 | 5 | 4 |
| 152 | 5 | 4 |
| 153 | 5 | 4 |
| 154 | 5 | 4 |
| 155 | 5 | 5 |
| 156 | 5 | 5 |
| 157 | 5 | 5 |
| 158 | 5 | 5 |
| 159 | 5 | 5 |
| 160 | 5 | 5 |
| 161 | 5 | 5 |
| 162 | 5 | 5 |
| 163 | 5 | 5 |
| 164 | 5 | 5 |
| 165 | 5 | 5 |
| 166 | 5 | 5 |
| 167 | 5 | 5 |
| 168 | 5 | 5 |
| 169 | 5 | 5 |
| 170 | 5 | 5 |
| 171 | 5 | 5 |
| 172 | 5 | 5 |
| 173 | 5 | 5 |
| 174 | 5 | 5 |
| 175 | 5 | 5 |
| 176 | 5 | 5 |
| 177 | 5 | 5 |
| 178 | 5 | 5 |
| 179 | 5 | 5 |
| 180 | 5 | 5 |
| 181 | 5 | 5 |
| 182 | 5 | 5 |
| 183 | 5 | 5 |
| 184 | 5 | 5 |
| 185 | 5 | 5 |
| 186 | 5 | 5 |
| 187 | 5 | 5 |
| 188 | 5 | 5 |
| 189 | 5 | 5 |
| 190 | 5 | 5 |
| 191 | 5 | 5 |
| 192 | 5 | 5 |
| 193 | 5 | 5 |
| 194 | 5 | 5 |
| 195 | 5 | 5 |
| 196 | 5 | 5 |
| 197 | 5 | 5 |
| 198 | 5 | 5 |
| 199 | 5 | 5 |
| 200 | 5 | 5 |

V3 Type-I-biased

|  | Male | Female |
| --- | --- | --- |
| 1 | 1.44e-10 | 1 |
| 2 | 1.173e-09 | 1 |
| 3 | 5.14e-10 | 1 |
| 4 | 1.005e-10 | 1 |
| 5 | 2.712e-10 | 1 |
| 6 | 1.945e-10 | 1 |
| 7 | 1.46571e-10 | 1 |
| 8 | 1.15875e-10 | 1 |
| 9 | 9.53333e-11 | 1 |
| 10 | 8.19e-11 | 1 |
| 11 | 3.65455e-10 | 1 |
| 12 | 1 | 1.91917e-11 |
| 13 | 1 | 3.79615e-10 |
| 14 | 1 | 4.32e-10 |
| 15 | 1 | 5.55333e-10 |
| 16 | 1 | 8.1375e-10 |
| 17 | 1 | 2.72176e-10 |
| 18 | 1 | 1.155e-10 |
| 19 | 1 | 1.02421e-10 |
| 20 | 0.933365 | 0.066635 |
| 21 | 0.852171 | 0.147829 |
| 22 | 0.783118 | 0.216882 |
| 23 | 0.725635 | 0.274365 |
| 24 | 0.678813 | 0.321188 |
| 25 | 0.641288 | 0.358712 |
| 26 | 1 | 1.70423e-12 |
| 27 | 1 | 1.7863e-12 |
| 28 | 1 | 1.9375e-12 |
| 29 | 1 | 2.16759e-12 |
| 30 | 1 | 2.49667e-12 |
| 31 | 1 | 1.48355e-11 |
| 32 | 1 | 3.63125e-12 |
| 33 | 1 | 2.31212e-11 |
| 34 | 1 | 6.07353e-12 |
| 35 | 1 | 3.38e-11 |
| 36 | 1 | 1.02472e-11 |
| 37 | 1 | 7.34054e-11 |
| 38 | 1 | 9.00789e-10 |
| 39 | 0.904949 | 0.0950513 |
| 40 | 0.822745 | 0.177255 |
| 41 | 1 | 6.09512e-12 |
| 42 | 1 | 6.45e-12 |
| 43 | 0.999998 | 6.93488e-12 |
| 44 | 1 | 3.77045e-11 |
| 45 | 1 | 1.66444e-12 |
| 46 | 0.999998 | 3.72826e-11 |
| 47 | 1 | 1.05447e-11 |
| 48 | 1 | 4.84167e-11 |
| 49 | 0.999998 | 2.82857e-12 |
| 50 | 1 | 3.346e-12 |
| 51 | 1 | 8.08431e-11 |
| 52 | 0.999998 | 2.00577e-11 |
| 53 | 1 | 2.56226e-11 |
| 54 | 1 | 1.71111e-10 |
| 55 | 0.999998 | 9.78727e-10 |
| 56 | 1 | 1.2875e-09 |
| 57 | 0.844937 | 0.155061 |
| 58 | 0.821405 | 0.178597 |
| 59 | 0.806934 | 0.193066 |
| 60 | 0.79627 | 0.203728 |
| 61 | 0.787785 | 0.212213 |
| 62 | 0.780726 | 0.219274 |
| 63 | 0.774671 | 0.225327 |
| 64 | 0.769373 | 0.230625 |
| 65 | 0.764663 | 0.235337 |
| 66 | 0.760418 | 0.239583 |
| 67 | 0.756555 | 0.243446 |
| 68 | 0.753004 | 0.246994 |
| 69 | 0.74973 | 0.250271 |
| 70 | 0.746679 | 0.25332 |
| 71 | 0.743831 | 0.25617 |
| 72 | 0.741154 | 0.258846 |
| 73 | 0.656022 | 0.343981 |
| 74 | 0.649532 | 0.350468 |
| 75 | 0.643608 | 0.356392 |
| 76 | 0.638171 | 0.361829 |
| 77 | 0.633156 | 0.366847 |
| 78 | 0.628508 | 0.371495 |
| 79 | 0.624178 | 0.37582 |
| 80 | 0.620141 | 0.37986 |
| 81 | 0.616356 | 0.383647 |
| 82 | 0.612794 | 0.387205 |
| 83 | 0.609437 | 0.390561 |
| 84 | 0.606264 | 0.393736 |
| 85 | 0.603254 | 0.396746 |
| 86 | 0.600398 | 0.399602 |
| 87 | 0.597676 | 0.402324 |
| 88 | 0.595077 | 0.404923 |
| 89 | 0.543519 | 0.456482 |
| 90 | 0.539393 | 0.460607 |
| 91 | 0.535487 | 0.464512 |
| 92 | 0.531789 | 0.468212 |
| 93 | 0.528277 | 0.471723 |
| 94 | 0.524936 | 0.475063 |
| 95 | 0.521757 | 0.478241 |
| 96 | 0.518725 | 0.481275 |
| 97 | 0.51583 | 0.484169 |
| 98 | 0.513061 | 0.48694 |
| 99 | 0.510409 | 0.489591 |
| 100 | 0.507864 | 0.492135 |
| 101 | 0.505423 | 0.494578 |
| 102 | 0.503076 | 0.496924 |
| 103 | 0.500817 | 0.499183 |
| 104 | 0.49864 | 0.501358 |
| 105 | 0.46396 | 0.536038 |
| 106 | 0.461021 | 0.538977 |
| 107 | 0.458198 | 0.541802 |
| 108 | 0.455489 | 0.544511 |
| 109 | 0.452882 | 0.547119 |
| 110 | 0.450374 | 0.549625 |
| 111 | 0.447957 | 0.552043 |
| 112 | 0.445629 | 0.554371 |
| 113 | 0.443381 | 0.556616 |
| 114 | 0.441213 | 0.558786 |
| 115 | 0.439119 | 0.56088 |
| 116 | 0.437095 | 0.562907 |
| 117 | 0.435133 | 0.564865 |
| 118 | 0.433236 | 0.566763 |
| 119 | 0.431397 | 0.568602 |
| 120 | 0.429615 | 0.570383 |
| 121 | 0.40464 | 0.59536 |
| 122 | 0.402425 | 0.597574 |
| 123 | 0.40028 | 0.59972 |
| 124 | 0.398204 | 0.601798 |
| 125 | 0.396192 | 0.603808 |
| 126 | 0.39424 | 0.605758 |
| 127 | 0.39235 | 0.60765 |
| 128 | 0.390516 | 0.609484 |
| 129 | 0.388735 | 0.611264 |
| 130 | 0.387007 | 0.612992 |
| 131 | 0.385326 | 0.614676 |
| 132 | 0.383691 | 0.616307 |
| 133 | 0.382103 | 0.617898 |
| 134 | 0.380557 | 0.619444 |
| 135 | 0.379051 | 0.620948 |
| 136 | 0.377585 | 0.622415 |
| 137 | 0.358721 | 0.641277 |
| 138 | 0.356987 | 0.643013 |
| 139 | 0.355299 | 0.644702 |
| 140 | 0.353655 | 0.646346 |
| 141 | 0.352057 | 0.647945 |
| 142 | 0.350499 | 0.6495 |
| 143 | 0.348982 | 0.651017 |
| 144 | 0.347502 | 0.652496 |
| 145 | 0.346061 | 0.653938 |
| 146 | 0.344655 | 0.655344 |
| 147 | 0.343282 | 0.656718 |
| 148 | 0.341943 | 0.658058 |
| 149 | 0.340637 | 0.659364 |
| 150 | 0.33936 | 0.66064 |
| 151 | 0.338112 | 0.661887 |
| 152 | 0.336894 | 0.663107 |
| 153 | 0.322133 | 0.677865 |
| 154 | 0.320737 | 0.679264 |
| 155 | 0.319372 | 0.680626 |
| 156 | 0.31804 | 0.681957 |
| 157 | 0.316739 | 0.683262 |
| 158 | 0.315467 | 0.684534 |
| 159 | 0.314225 | 0.685775 |
| 160 | 0.313009 | 0.686993 |
| 161 | 0.311819 | 0.688183 |
| 162 | 0.310656 | 0.689344 |
| 163 | 0.309517 | 0.690483 |
| 164 | 0.308402 | 0.691596 |
| 165 | 0.307311 | 0.69269 |
| 166 | 0.306242 | 0.693759 |
| 167 | 0.305197 | 0.694802 |
| 168 | 0.304171 | 0.695829 |
| 169 | 0.303165 | 0.696836 |
| 170 | 0.302181 | 0.697818 |
| 171 | 0.301214 | 0.698784 |
| 172 | 0.300267 | 0.699731 |
| 173 | 0.299338 | 0.700664 |
| 174 | 0.298426 | 0.701573 |
| 175 | 0.297531 | 0.702468 |
| 176 | 0.296652 | 0.703349 |
| 177 | 0.29579 | 0.704208 |
| 178 | 0.294944 | 0.705057 |
| 179 | 0.294112 | 0.705889 |
| 180 | 0.293295 | 0.706704 |
| 181 | 0.292492 | 0.70751 |
| 182 | 0.291702 | 0.708296 |
| 183 | 0.290926 | 0.709073 |
| 184 | 0.290164 | 0.709838 |
| 185 | 0.289412 | 0.710587 |
| 186 | 0.288674 | 0.711324 |
| 187 | 0.287949 | 0.71205 |
| 188 | 0.287234 | 0.712764 |
| 189 | 0.286532 | 0.713467 |
| 190 | 0.28584 | 0.714162 |
| 191 | 0.285159 | 0.714843 |
| 192 | 0.284488 | 0.715513 |
| 193 | 0.283826 | 0.716173 |
| 194 | 0.283177 | 0.716825 |
| 195 | 0.282535 | 0.717464 |
| 196 | 0.281904 | 0.718096 |
| 197 | 0.281281 | 0.718719 |
| 198 | 0.280668 | 0.719331 |
| 199 | 0.280063 | 0.719938 |
| 200 | 0.279467 | 0.720534 |

V3 Type-I-biased

|  | Male | Female |
| --- | --- | --- |
| 1 | 0 | 1 |
| 2 | 0 | 1 |
| 3 | 0 | 1 |
| 4 | 0 | 1 |
| 5 | 0 | 1 |
| 6 | 0 | 1 |
| 7 | 0 | 1 |
| 8 | 0 | 1 |
| 9 | 0 | 1 |
| 10 | 0 | 1 |
| 11 | 0 | 1 |
| 12 | 1 | 0 |
| 13 | 1 | 0 |
| 14 | 1 | 0 |
| 15 | 1 | 0 |
| 16 | 1 | 0 |
| 17 | 1 | 0 |
| 18 | 1 | 0 |
| 19 | 1 | 0 |
| 20 | 1 | 1 |
| 21 | 1 | 1 |
| 22 | 1 | 1 |
| 23 | 1 | 1 |
| 24 | 1 | 1 |
| 25 | 1 | 1 |
| 26 | 2 | 0 |
| 27 | 2 | 0 |
| 28 | 2 | 0 |
| 29 | 2 | 0 |
| 30 | 2 | 0 |
| 31 | 2 | 0 |
| 32 | 2 | 0 |
| 33 | 2 | 0 |
| 34 | 2 | 0 |
| 35 | 2 | 0 |
| 36 | 2 | 0 |
| 37 | 2 | 0 |
| 38 | 2 | 0 |
| 39 | 2 | 1 |
| 40 | 2 | 1 |
| 41 | 3 | 0 |
| 42 | 3 | 0 |
| 43 | 3 | 0 |
| 44 | 3 | 0 |
| 45 | 3 | 0 |
| 46 | 3 | 0 |
| 47 | 3 | 0 |
| 48 | 3 | 0 |
| 49 | 3 | 0 |
| 50 | 3 | 0 |
| 51 | 3 | 0 |
| 52 | 3 | 0 |
| 53 | 3 | 0 |
| 54 | 3 | 0 |
| 55 | 3 | 0 |
| 56 | 3 | 0 |
| 57 | 3 | 1 |
| 58 | 3 | 1 |
| 59 | 3 | 1 |
| 60 | 3 | 1 |
| 61 | 3 | 1 |
| 62 | 3 | 1 |
| 63 | 3 | 1 |
| 64 | 3 | 1 |
| 65 | 3 | 1 |
| 66 | 3 | 1 |
| 67 | 3 | 1 |
| 68 | 3 | 1 |
| 69 | 3 | 1 |
| 70 | 3 | 1 |
| 71 | 3 | 1 |
| 72 | 3 | 1 |
| 73 | 3 | 2 |
| 74 | 3 | 2 |
| 75 | 3 | 2 |
| 76 | 3 | 2 |
| 77 | 3 | 2 |
| 78 | 3 | 2 |
| 79 | 3 | 2 |
| 80 | 3 | 2 |
| 81 | 3 | 2 |
| 82 | 3 | 2 |
| 83 | 3 | 2 |
| 84 | 3 | 2 |
| 85 | 3 | 2 |
| 86 | 3 | 2 |
| 87 | 3 | 2 |
| 88 | 3 | 2 |
| 89 | 3 | 3 |
| 90 | 3 | 3 |
| 91 | 3 | 3 |
| 92 | 3 | 3 |
| 93 | 3 | 3 |
| 94 | 3 | 3 |
| 95 | 3 | 3 |
| 96 | 3 | 3 |
| 97 | 3 | 3 |
| 98 | 3 | 3 |
| 99 | 3 | 3 |
| 100 | 3 | 3 |
| 101 | 3 | 3 |
| 102 | 3 | 3 |
| 103 | 3 | 3 |
| 104 | 3 | 3 |
| 105 | 3 | 4 |
| 106 | 3 | 4 |
| 107 | 3 | 4 |
| 108 | 3 | 4 |
| 109 | 3 | 4 |
| 110 | 3 | 4 |
| 111 | 3 | 4 |
| 112 | 3 | 4 |
| 113 | 3 | 4 |
| 114 | 3 | 4 |
| 115 | 3 | 4 |
| 116 | 3 | 4 |
| 117 | 3 | 4 |
| 118 | 3 | 4 |
| 119 | 3 | 4 |
| 120 | 3 | 4 |
| 121 | 3 | 5 |
| 122 | 3 | 5 |
| 123 | 3 | 5 |
| 124 | 3 | 5 |
| 125 | 3 | 5 |
| 126 | 3 | 5 |
| 127 | 3 | 5 |
| 128 | 3 | 5 |
| 129 | 3 | 5 |
| 130 | 3 | 5 |
| 131 | 3 | 5 |
| 132 | 3 | 5 |
| 133 | 3 | 5 |
| 134 | 3 | 5 |
| 135 | 3 | 5 |
| 136 | 3 | 5 |
| 137 | 3 | 6 |
| 138 | 3 | 6 |
| 139 | 3 | 6 |
| 140 | 3 | 6 |
| 141 | 3 | 6 |
| 142 | 3 | 6 |
| 143 | 3 | 6 |
| 144 | 3 | 6 |
| 145 | 3 | 6 |
| 146 | 3 | 6 |
| 147 | 3 | 6 |
| 148 | 3 | 6 |
| 149 | 3 | 6 |
| 150 | 3 | 6 |
| 151 | 3 | 6 |
| 152 | 3 | 6 |
| 153 | 3 | 7 |
| 154 | 3 | 7 |
| 155 | 3 | 7 |
| 156 | 3 | 7 |
| 157 | 3 | 7 |
| 158 | 3 | 7 |
| 159 | 3 | 7 |
| 160 | 3 | 7 |
| 161 | 3 | 7 |
| 162 | 3 | 7 |
| 163 | 3 | 7 |
| 164 | 3 | 7 |
| 165 | 3 | 7 |
| 166 | 3 | 7 |
| 167 | 3 | 7 |
| 168 | 3 | 7 |
| 169 | 3 | 7 |
| 170 | 3 | 7 |
| 171 | 3 | 7 |
| 172 | 3 | 7 |
| 173 | 3 | 7 |
| 174 | 3 | 7 |
| 175 | 3 | 7 |
| 176 | 3 | 7 |
| 177 | 3 | 7 |
| 178 | 3 | 7 |
| 179 | 3 | 7 |
| 180 | 3 | 7 |
| 181 | 3 | 7 |
| 182 | 3 | 7 |
| 183 | 3 | 7 |
| 184 | 3 | 7 |
| 185 | 3 | 7 |
| 186 | 3 | 7 |
| 187 | 3 | 7 |
| 188 | 3 | 7 |
| 189 | 3 | 7 |
| 190 | 3 | 7 |
| 191 | 3 | 7 |
| 192 | 3 | 7 |
| 193 | 3 | 7 |
| 194 | 3 | 7 |
| 195 | 3 | 7 |
| 196 | 3 | 7 |
| 197 | 3 | 7 |
| 198 | 3 | 7 |
| 199 | 3 | 7 |
| 200 | 3 | 7 |

Figure S6 | Same as Figure S4 for model version 3.

**Model Version 4**

V4 Functions

|  |  |  |
| --- | --- | --- |
| 0 | 0 | -4.28267e-05 |
| 0.01 | 2.07149e-10 | 0.000317246 |
| 0.02 | 4.16381e-10 | 0.000678014 |
| 0.03 | 6.27715e-10 | 0.00103948 |
| 0.04 | 8.41173e-10 | 0.00140164 |
| 0.05 | 1.05678e-09 | 0.0017645 |
| 0.06 | 1.27455e-09 | 0.00212806 |
| 0.07 | 1.49451e-09 | 0.00249232 |
| 0.08 | 1.71667e-09 | 0.00285728 |
| 0.09 | 1.94108e-09 | 0.00322295 |
| 0.1 | 2.16773e-09 | 0.00358932 |
| 0.11 | 2.39667e-09 | 0.0039564 |
| 0.12 | 2.62791e-09 | 0.00432419 |
| 0.13 | 2.86147e-09 | 0.00469269 |
| 0.14 | 3.09737e-09 | 0.0050619 |
| 0.15 | 3.33565e-09 | 0.00543181 |
| 0.16 | 3.57633e-09 | 0.00580245 |
| 0.17 | 3.81942e-09 | 0.00617379 |
| 0.18 | 4.06495e-09 | 0.00654586 |
| 0.19 | 4.31296e-09 | 0.00691863 |
| 0.2 | 4.56345e-09 | 0.00729213 |
| 0.21 | 4.81646e-09 | 0.00766635 |
| 0.22 | 5.07202e-09 | 0.00804129 |
| 0.23 | 5.33014e-09 | 0.00841695 |
| 0.24 | 5.59086e-09 | 0.00879333 |
| 0.25 | 5.8542e-09 | 0.00917044 |
| 0.26 | 6.12019e-09 | 0.00954828 |
| 0.27 | 6.38884e-09 | 0.00992684 |
| 0.28 | 6.6602e-09 | 0.0103061 |
| 0.29 | 6.93429e-09 | 0.0106862 |
| 0.3 | 7.21113e-09 | 0.0110669 |
| 0.31 | 7.49075e-09 | 0.0114484 |
| 0.32 | 7.77318e-09 | 0.0118306 |
| 0.33 | 8.05845e-09 | 0.0122136 |
| 0.34 | 8.34659e-09 | 0.0125973 |
| 0.35 | 8.63763e-09 | 0.0129817 |
| 0.36 | 8.93158e-09 | 0.0133669 |
| 0.37 | 9.2285e-09 | 0.0137528 |
| 0.38 | 9.5284e-09 | 0.0141395 |
| 0.39 | 9.83131e-09 | 0.0145269 |
| 0.4 | 1.01373e-08 | 0.014915 |
| 0.41 | 1.04463e-08 | 0.0153039 |
| 0.42 | 1.07584e-08 | 0.0156936 |
| 0.43 | 1.10737e-08 | 0.0160839 |
| 0.44 | 1.13921e-08 | 0.0164751 |
| 0.45 | 1.17138e-08 | 0.016867 |
| 0.46 | 1.20387e-08 | 0.0172596 |
| 0.47 | 1.23668e-08 | 0.017653 |
| 0.48 | 1.26982e-08 | 0.0180472 |
| 0.49 | 1.3033e-08 | 0.0184421 |
| 0.5 | 1.33711e-08 | 0.0188378 |
| 0.51 | 1.37127e-08 | 0.0192342 |
| 0.52 | 1.40576e-08 | 0.0196314 |
| 0.53 | 1.44061e-08 | 0.0200294 |
| 0.54 | 1.4758e-08 | 0.0204281 |
| 0.55 | 1.51135e-08 | 0.0208276 |
| 0.56 | 1.54725e-08 | 0.0212279 |
| 0.57 | 1.58352e-08 | 0.0216289 |
| 0.58 | 1.62015e-08 | 0.0220307 |
| 0.59 | 1.65714e-08 | 0.0224333 |
| 0.6 | 1.69451e-08 | 0.0228367 |
| 0.61 | 1.73226e-08 | 0.0232408 |
| 0.62 | 1.77038e-08 | 0.0236457 |
| 0.63 | 1.80889e-08 | 0.0240514 |
| 0.64 | 1.84778e-08 | 0.0244578 |
| 0.65 | 1.88707e-08 | 0.024865 |
| 0.66 | 1.92675e-08 | 0.0252731 |
| 0.67 | 1.96683e-08 | 0.0256819 |
| 0.68 | 2.00731e-08 | 0.0260915 |
| 0.69 | 2.0482e-08 | 0.0265018 |
| 0.7 | 2.0895e-08 | 0.026913 |
| 0.71 | 2.13121e-08 | 0.0273249 |
| 0.72 | 2.17335e-08 | 0.0277377 |
| 0.73 | 2.21591e-08 | 0.0281512 |
| 0.74 | 2.25889e-08 | 0.0285655 |
| 0.75 | 2.30231e-08 | 0.0289807 |
| 0.76 | 2.34616e-08 | 0.0293966 |
| 0.77 | 2.39046e-08 | 0.0298133 |
| 0.78 | 2.4352e-08 | 0.0302308 |
| 0.79 | 2.48038e-08 | 0.0306491 |
| 0.8 | 2.52603e-08 | 0.0310682 |
| 0.81 | 2.57213e-08 | 0.0314882 |
| 0.82 | 2.6187e-08 | 0.0319089 |
| 0.83 | 2.66573e-08 | 0.0323304 |
| 0.84 | 2.71323e-08 | 0.0327528 |
| 0.85 | 2.76122e-08 | 0.0331759 |
| 0.86 | 2.80968e-08 | 0.0335999 |
| 0.87 | 2.85864e-08 | 0.0340247 |
| 0.88 | 2.90808e-08 | 0.0344502 |
| 0.89 | 2.95802e-08 | 0.0348767 |
| 0.9 | 3.00847e-08 | 0.0353039 |
| 0.91 | 3.05942e-08 | 0.0357319 |
| 0.92 | 3.11088e-08 | 0.0361608 |
| 0.93 | 3.16286e-08 | 0.0365905 |
| 0.94 | 3.21536e-08 | 0.037021 |
| 0.95 | 3.26839e-08 | 0.0374523 |
| 0.96 | 3.32195e-08 | 0.0378844 |
| 0.97 | 3.37606e-08 | 0.0383174 |
| 0.98 | 3.4307e-08 | 0.0387512 |
| 0.99 | 3.48589e-08 | 0.0391859 |
| 1 | 3.54164e-08 | 0.0396214 |
| 1.01 | 3.59795e-08 | 0.0400577 |
| 1.02 | 3.65483e-08 | 0.0404948 |
| 1.03 | 3.71227e-08 | 0.0409328 |
| 1.04 | 3.7703e-08 | 0.0413716 |
| 1.05 | 3.8289e-08 | 0.0418113 |
| 1.06 | 3.8881e-08 | 0.0422518 |
| 1.07 | 3.94789e-08 | 0.0426931 |
| 1.08 | 4.00828e-08 | 0.0431353 |
| 1.09 | 4.06928e-08 | 0.0435783 |
| 1.1 | 4.13089e-08 | 0.0440222 |
| 1.11 | 4.19313e-08 | 0.0444669 |
| 1.12 | 4.25598e-08 | 0.0449125 |
| 1.13 | 4.31947e-08 | 0.0453589 |
| 1.14 | 4.3836e-08 | 0.0458062 |
| 1.15 | 4.44837e-08 | 0.0462544 |
| 1.16 | 4.51379e-08 | 0.0467034 |
| 1.17 | 4.57987e-08 | 0.0471532 |
| 1.18 | 4.64661e-08 | 0.0476039 |
| 1.19 | 4.71403e-08 | 0.0480555 |
| 1.2 | 4.78212e-08 | 0.048508 |
| 1.21 | 4.85089e-08 | 0.0489613 |
| 1.22 | 4.92036e-08 | 0.0494154 |
| 1.23 | 4.99053e-08 | 0.0498705 |
| 1.24 | 5.0614e-08 | 0.0503264 |
| 1.25 | 5.13298e-08 | 0.0507832 |
| 1.26 | 5.20528e-08 | 0.0512408 |
| 1.27 | 5.27831e-08 | 0.0516993 |
| 1.28 | 5.35207e-08 | 0.0521587 |
| 1.29 | 5.42658e-08 | 0.052619 |
| 1.3 | 5.50183e-08 | 0.0530802 |
| 1.31 | 5.57784e-08 | 0.0535422 |
| 1.32 | 5.65461e-08 | 0.0540051 |
| 1.33 | 5.73216e-08 | 0.0544689 |
| 1.34 | 5.81048e-08 | 0.0549336 |
| 1.35 | 5.88959e-08 | 0.0553992 |
| 1.36 | 5.9695e-08 | 0.0558656 |
| 1.37 | 6.05021e-08 | 0.056333 |
| 1.38 | 6.13173e-08 | 0.0568012 |
| 1.39 | 6.21407e-08 | 0.0572704 |
| 1.4 | 6.29724e-08 | 0.0577404 |
| 1.41 | 6.38124e-08 | 0.0582113 |
| 1.42 | 6.46609e-08 | 0.0586832 |
| 1.43 | 6.55179e-08 | 0.0591559 |
| 1.44 | 6.63835e-08 | 0.0596295 |
| 1.45 | 6.72578e-08 | 0.060104 |
| 1.46 | 6.81409e-08 | 0.0605795 |
| 1.47 | 6.90329e-08 | 0.0610558 |
| 1.48 | 6.99338e-08 | 0.0615331 |
| 1.49 | 7.08438e-08 | 0.0620112 |
| 1.5 | 7.1763e-08 | 0.0624903 |
| 1.51 | 7.26913e-08 | 0.0629703 |
| 1.52 | 7.3629e-08 | 0.0634512 |
| 1.53 | 7.45762e-08 | 0.063933 |
| 1.54 | 7.55328e-08 | 0.0644157 |
| 1.55 | 7.64991e-08 | 0.0648994 |
| 1.56 | 7.74751e-08 | 0.0653839 |
| 1.57 | 7.84609e-08 | 0.0658694 |
| 1.58 | 7.94566e-08 | 0.0663558 |
| 1.59 | 8.04623e-08 | 0.0668432 |
| 1.6 | 8.14781e-08 | 0.0673314 |
| 1.61 | 8.25041e-08 | 0.0678206 |
| 1.62 | 8.35404e-08 | 0.0683108 |
| 1.63 | 8.45872e-08 | 0.0688018 |
| 1.64 | 8.56444e-08 | 0.0692938 |
| 1.65 | 8.67123e-08 | 0.0697868 |
| 1.66 | 8.77909e-08 | 0.0702807 |
| 1.67 | 8.88804e-08 | 0.0707755 |
| 1.68 | 8.99808e-08 | 0.0712712 |
| 1.69 | 9.10923e-08 | 0.0717679 |
| 1.7 | 9.22149e-08 | 0.0722656 |
| 1.71 | 9.33489e-08 | 0.0727641 |
| 1.72 | 9.44942e-08 | 0.0732637 |
| 1.73 | 9.5651e-08 | 0.0737642 |
| 1.74 | 9.68195e-08 | 0.0742656 |
| 1.75 | 9.79997e-08 | 0.074768 |
| 1.76 | 9.91917e-08 | 0.0752713 |
| 1.77 | 1.00396e-07 | 0.0757756 |
| 1.78 | 1.01612e-07 | 0.0762809 |
| 1.79 | 1.0284e-07 | 0.0767871 |
| 1.8 | 1.04081e-07 | 0.0772943 |
| 1.81 | 1.05334e-07 | 0.0778024 |
| 1.82 | 1.066e-07 | 0.0783115 |
| 1.83 | 1.07878e-07 | 0.0788216 |
| 1.84 | 1.0917e-07 | 0.0793327 |
| 1.85 | 1.10474e-07 | 0.0798447 |
| 1.86 | 1.11792e-07 | 0.0803577 |
| 1.87 | 1.13122e-07 | 0.0808716 |
| 1.88 | 1.14466e-07 | 0.0813865 |
| 1.89 | 1.15824e-07 | 0.0819025 |
| 1.9 | 1.17195e-07 | 0.0824194 |
| 1.91 | 1.1858e-07 | 0.0829372 |
| 1.92 | 1.19979e-07 | 0.0834561 |
| 1.93 | 1.21392e-07 | 0.0839759 |
| 1.94 | 1.22819e-07 | 0.0844967 |
| 1.95 | 1.24261e-07 | 0.0850186 |
| 1.96 | 1.25716e-07 | 0.0855414 |
| 1.97 | 1.27187e-07 | 0.0860652 |
| 1.98 | 1.28673e-07 | 0.0865899 |
| 1.99 | 1.30173e-07 | 0.0871157 |

V4 Total fitness

|  | Fast-biased | Equal | Slow-biased |
| --- | --- | --- | --- |
| 1 | 0.0395357 | 0.0394501 | 0.0393644 |
| 2 | 0.0875569 | 0.0874712 | 0.0873855 |
| 3 | 0.145586 | 0.1455 | 0.145414 |
| 4 | 0.215547 | 0.215461 | 0.215375 |
| 5 | 0.299661 | 0.299576 | 0.29949 |
| 6 | 0.400458 | 0.400372 | 0.400287 |
| 7 | 0.520765 | 0.520679 | 0.520593 |
| 8 | 0.663679 | 0.663593 | 0.663507 |
| 9 | 0.832494 | 0.832408 | 0.832322 |
| 10 | 1.03058 | 1.0305 | 1.03041 |
| 11 | 1.26121 | 1.26113 | 1.26104 |
| 12 | 1.5273 | 1.52722 | 1.52713 |
| 13 | 1.83112 | 1.83103 | 1.83094 |
| 14 | 2.17387 | 2.17379 | 2.1737 |
| 15 | 2.5554 | 2.55531 | 2.55523 |
| 16 | 2.97377 | 2.97368 | 2.97359 |
| 17 | 3.42507 | 3.42498 | 3.4249 |
| 18 | 3.90336 | 3.90327 | 3.90319 |
| 19 | 4.40084 | 4.40076 | 4.40067 |
| 20 | 4.99987 | 4.99979 | 4.9997 |
| 21 | 7.31046 | 7.31037 | 7.31029 |
| 22 | 8.80784 | 8.80776 | 8.80767 |
| 23 | 9.52561 | 9.52553 | 9.52544 |
| 24 | 9.82001 | 9.81992 | 9.81984 |
| 25 | 9.93294 | 9.93286 | 9.93277 |
| 26 | 9.97806 | 9.97798 | 9.97789 |
| 27 | 10.0225 | 10.0224 | 10.0223 |
| 28 | 10.0787 | 10.0786 | 10.0786 |
| 29 | 10.1496 | 10.1495 | 10.1494 |
| 30 | 10.2383 | 10.2382 | 10.2381 |
| 31 | 10.3489 | 10.3488 | 10.3487 |
| 32 | 10.4855 | 10.4854 | 10.4853 |
| 33 | 10.6528 | 10.6527 | 10.6526 |
| 34 | 10.8553 | 10.8552 | 10.8551 |
| 35 | 11.0973 | 11.0972 | 11.0971 |
| 36 | 11.3821 | 11.382 | 11.3819 |
| 37 | 11.7113 | 11.7112 | 11.7112 |
| 38 | 12.0848 | 12.0848 | 12.0847 |
| 39 | 12.4999 | 12.4998 | 12.4997 |
| 40 | 12.9511 | 12.951 | 12.9509 |
| 41 | 13.4309 | 13.4308 | 13.4307 |
| 42 | 14.621 | 14.621 | 14.6209 |
| 43 | 16.3514 | 16.3513 | 16.3512 |
| 44 | 17.6158 | 17.6157 | 17.6156 |
| 45 | 18.4827 | 18.4826 | 18.4825 |
| 46 | 19.0514 | 19.0513 | 19.0512 |
| 47 | 19.4136 | 19.4135 | 19.4135 |
| 48 | 19.6401 | 19.6401 | 19.64 |
| 49 | 19.7801 | 19.78 | 19.78 |
| 50 | 19.866 | 19.8659 | 19.8658 |
| 51 | 19.9185 | 19.9184 | 19.9183 |
| 52 | 19.9585 | 19.9584 | 19.9583 |
| 53 | 20.0119 | 20.0118 | 20.0118 |
| 54 | 20.0842 | 20.0841 | 20.084 |
| 55 | 20.1807 | 20.1807 | 20.1806 |
| 56 | 20.3081 | 20.308 | 20.3079 |
| 57 | 20.4731 | 20.473 | 20.4729 |
| 58 | 20.6823 | 20.6822 | 20.6822 |
| 59 | 20.9411 | 20.941 | 20.9409 |
| 60 | 21.2522 | 21.2521 | 21.252 |
| 61 | 21.6152 | 21.6151 | 21.615 |
| 62 | 22.026 | 22.0259 | 22.0258 |
| 63 | 22.4771 | 22.477 | 22.4769 |
| 64 | 23.7416 | 23.7415 | 23.7414 |
| 65 | 25.2338 | 25.2337 | 25.2336 |
| 66 | 26.4238 | 26.4237 | 26.4236 |
| 67 | 27.3479 | 27.3478 | 27.3477 |
| 68 | 28.0508 | 28.0507 | 28.0506 |
| 69 | 28.5771 | 28.577 | 28.5769 |
| 70 | 28.9665 | 28.9664 | 28.9663 |
| 71 | 29.2521 | 29.2521 | 29.252 |
| 72 | 29.4603 | 29.4602 | 29.4601 |
| 73 | 29.6113 | 29.6112 | 29.6111 |
| 74 | 29.7204 | 29.7203 | 29.7202 |
| 75 | 29.7991 | 29.799 | 29.7989 |
| 76 | 29.8557 | 29.8556 | 29.8556 |
| 77 | 29.8967 | 29.8966 | 29.8966 |
| 78 | 29.9452 | 29.9451 | 29.9451 |
| 79 | 30.0202 | 30.0201 | 30.02 |
| 80 | 30.1324 | 30.1323 | 30.1322 |
| 81 | 30.2936 | 30.2935 | 30.2934 |
| 82 | 30.5138 | 30.5137 | 30.5136 |
| 83 | 30.7985 | 30.7984 | 30.7983 |
| 84 | 31.1469 | 31.1469 | 31.1468 |
| 85 | 31.5523 | 31.5522 | 31.5521 |
| 86 | 32.7029 | 32.7028 | 32.003 |
| 87 | 34.078 | 34.0779 | 32.4862 |
| 88 | 35.2318 | 35.2317 | 32.9882 |
| 89 | 36.1859 | 36.1858 | 33.4969 |
| 90 | 36.9655 | 36.9655 | 34.0019 |
| 91 | 37.5964 | 37.5963 | 34.4948 |
| 92 | 38.1028 | 38.1028 | 34.9692 |
| 93 | 38.5068 | 38.5067 | 35.4207 |
| 94 | 38.8274 | 38.8273 | 35.8461 |
| 95 | 39.0808 | 39.0807 | 36.2435 |
| 96 | 39.2804 | 39.2803 | 36.6123 |
| 97 | 39.4373 | 39.4372 | 36.9523 |
| 98 | 39.5604 | 39.5603 | 37.2641 |
| 99 | 39.6568 | 39.6567 | 37.5488 |
| 100 | 39.7322 | 39.7321 | 37.8076 |
| 101 | 39.7911 | 39.791 | 38.0421 |
| 102 | 39.8371 | 39.837 | 38.2539 |
| 103 | 39.8789 | 39.8788 | 38.4446 |
| 104 | 39.9611 | 39.961 | 38.616 |
| 105 | 40.1144 | 40.1143 | 38.7696 |
| 106 | 40.3548 | 40.3547 | 38.9072 |
| 107 | 40.6809 | 40.6808 | 39.03 |
| 108 | 41.6008 | 41.0786 | 39.1396 |
| 109 | 42.9073 | 41.5293 | 39.2373 |
| 110 | 44.0397 | 42.0143 | 39.3242 |
| 111 | 45.0123 | 45.0123 | 39.4181 |
| 112 | 45.8412 | 43.0265 | 39.9171 |
| 113 | 46.543 | 46.5429 | 40.4084 |
| 114 | 47.1337 | 44.0218 | 40.8903 |
| 115 | 47.6286 | 44.4952 | 41.361 |
| 116 | 48.0416 | 48.0415 | 41.8192 |
| 117 | 48.3851 | 48.385 | 42.2638 |
| 118 | 48.67 | 45.775 | 42.6937 |
| 119 | 48.9058 | 48.9057 | 43.1084 |
| 120 | 49.1006 | 46.4978 | 43.5072 |
| 121 | 49.2612 | 49.2611 | 43.8896 |
| 122 | 49.3934 | 49.3934 | 44.2556 |
| 123 | 49.5023 | 47.3895 | 44.6051 |
| 124 | 49.5917 | 47.6393 | 44.938 |
| 125 | 49.6652 | 49.6651 | 45.2545 |
| 126 | 49.7256 | 48.0756 | 45.555 |
| 127 | 49.7751 | 48.2649 | 45.8396 |
| 128 | 49.8157 | 49.8156 | 46.1089 |
| 129 | 49.9365 | 48.5926 | 46.3632 |
| 130 | 50.219 | 50.219 | 46.6031 |
| 131 | 50.6053 | 48.8614 | 46.829 |
| 132 | 51.0556 | 48.9766 | 47.0416 |
| 133 | 51.5427 | 51.5426 | 47.2414 |
| 134 | 52.0474 | 52.0473 | 47.4289 |
| 135 | 52.556 | 52.5559 | 47.6048 |
| 136 | 53.0588 | 53.0587 | 47.7696 |
| 137 | 53.5488 | 53.5487 | 47.9239 |
| 138 | 54.0212 | 50.887 | 48.0682 |
| 139 | 54.4727 | 54.4726 | 48.2134 |
| 140 | 57.9332 | 54.9012 | 48.6649 |
| 141 | 55.3056 | 55.3056 | 49.1068 |
| 142 | 58.5044 | 52.6367 | 49.5385 |
| 143 | 58.7291 | 56.0404 | 49.9598 |
| 144 | 56.3713 | 56.3712 | 50.3701 |
| 145 | 59.0838 | 56.6785 | 50.7694 |
| 146 | 56.9633 | 56.9632 | 51.1573 |
| 147 | 59.3407 | 57.2262 | 51.5338 |
| 148 | 59.4409 | 57.4687 | 51.8987 |
| 149 | 59.526 | 57.6919 | 52.252 |
| 150 | 57.897 | 57.8969 | 52.5936 |
| 151 | 59.6596 | 55.6835 | 52.9237 |
| 152 | 59.7116 | 58.2574 | 53.2421 |
| 153 | 58.4151 | 58.415 | 53.5492 |
| 154 | 60.1322 | 58.5591 | 53.845 |
| 155 | 60.5819 | 58.6907 | 54.1296 |
| 156 | 61.0711 | 58.8108 | 54.4033 |
| 157 | 58.9203 | 57.0624 | 54.6663 |
| 158 | 62.0853 | 59.0199 | 54.9187 |
| 159 | 62.5869 | 59.4628 | 55.1609 |
| 160 | 63.0756 | 59.1932 | 55.3931 |
| 161 | 63.5471 | 59.2682 | 55.6155 |
| 162 | 63.9986 | 59.3365 | 55.8284 |
| 163 | 64.4285 | 61.3158 | 56.0322 |
| 164 | 64.8356 | 58.6329 | 56.227 |
| 165 | 65.2198 | 62.1647 | 56.4132 |
| 166 | 65.5809 | 62.5675 | 56.5911 |
| 167 | 65.9193 | 62.9554 | 56.8071 |
| 168 | 66.2356 | 63.3284 | 57.2236 |
| 169 | 66.5306 | 63.6863 | 57.6317 |
| 170 | 66.8051 | 64.0291 | 58.0312 |
| 171 | 67.0601 | 64.357 | 58.4221 |
| 172 | 67.2966 | 64.6702 | 58.8042 |
| 173 | 67.5156 | 64.969 | 59.1774 |
| 174 | 67.7182 | 65.2536 | 59.5417 |
| 175 | 67.9053 | 65.5245 | 59.8971 |
| 176 | 68.078 | 65.7819 | 60.2434 |
| 177 | 68.2372 | 66.0264 | 60.5808 |
| 178 | 68.3838 | 66.2583 | 60.9093 |
| 179 | 68.5188 | 66.4781 | 61.2289 |
| 180 | 68.6429 | 66.6862 | 61.5396 |
| 181 | 68.757 | 66.8832 | 61.8415 |
| 182 | 68.99 | 67.0694 | 62.1348 |
| 183 | 69.4697 | 67.2454 | 62.4194 |
| 184 | 69.9389 | 67.4116 | 62.6955 |
| 185 | 70.3966 | 67.5684 | 62.9633 |
| 186 | 70.8418 | 67.7165 | 63.2229 |
| 187 | 74.3652 | 68.1589 | 63.4743 |
| 188 | 71.6926 | 68.5916 | 63.7177 |
| 189 | 75.1195 | 69.0144 | 63.9534 |
| 190 | 72.4875 | 69.427 | 64.1814 |
| 191 | 72.8635 | 69.8291 | 64.4018 |
| 192 | 73.2251 | 70.2206 | 64.615 |
| 193 | 76.3774 | 70.6014 | 64.8209 |
| 194 | 73.9051 | 70.9714 | 65.0199 |
| 195 | 74.2238 | 71.3305 | 65.2929 |
| 196 | 74.5287 | 71.6787 | 65.6856 |
| 197 | 74.82 | 72.0161 | 66.0711 |
| 198 | 75.0981 | 72.3427 | 66.4495 |
| 199 | 77.7255 | 72.6587 | 66.8207 |
| 200 | 75.6157 | 72.964 | 67.1847 |

V4 Type-II-biased

|  |  |  |
| --- | --- | --- |
| 1 | 1.07527e-09 | 1 |
| 2 | 5.67914e-10 | 1 |
| 3 | 3.17408e-10 | 1 |
| 4 | 9.95648e-10 | 1 |
| 5 | 5.33293e-10 | 1 |
| 6 | 3.72739e-10 | 1 |
| 7 | 2.68881e-10 | 1 |
| 8 | 2.4863e-10 | 1 |
| 9 | 7.51642e-10 | 1 |
| 10 | 5.79155e-10 | 1 |
| 11 | 4.54464e-10 | 1 |
| 12 | 3.63051e-10 | 1 |
| 13 | 2.953e-10 | 1 |
| 14 | 2.44686e-10 | 1 |
| 15 | 2.06689e-10 | 1 |
| 16 | 1.78139e-10 | 1 |
| 17 | 1.56784e-10 | 1 |
| 18 | 1.41015e-10 | 1 |
| 19 | 1.29685e-10 | 1 |
| 20 | 1 | 2.70852e-12 |
| 21 | 1 | 1.647e-11 |
| 22 | 1 | 5.9927e-12 |
| 23 | 1 | 5.61193e-11 |
| 24 | 1 | 1.6077e-10 |
| 25 | 1 | 2.97457e-11 |
| 26 | 0.981885 | 0.0181149 |
| 27 | 0.936673 | 0.0633269 |
| 28 | 0.8948 | 0.1052 |
| 29 | 0.855948 | 0.144052 |
| 30 | 0.819848 | 0.180152 |
| 31 | 0.786272 | 0.213728 |
| 32 | 0.755029 | 0.244971 |
| 33 | 0.725958 | 0.274042 |
| 34 | 0.698926 | 0.301074 |
| 35 | 0.673819 | 0.326181 |
| 36 | 0.650542 | 0.349458 |
| 37 | 0.62901 | 0.37099 |
| 38 | 0.609144 | 0.390856 |
| 39 | 0.590864 | 0.409136 |
| 40 | 0.574086 | 0.425914 |
| 41 | 0.558717 | 0.441283 |
| 42 | 1 | 2.96054e-13 |
| 43 | 1 | 9.58681e-14 |
| 44 | 1 | 2.68972e-12 |
| 45 | 1 | 8.01927e-13 |
| 46 | 1 | 2.5096e-13 |
| 47 | 1 | 2.05441e-12 |
| 48 | 1 | 7.10962e-13 |
| 49 | 1 | 5.39057e-12 |
| 50 | 1 | 2.51738e-12 |
| 51 | 1 | 1.65006e-11 |
| 52 | 0.976185 | 0.0238153 |
| 53 | 0.946077 | 0.0539234 |
| 54 | 0.917398 | 0.0826025 |
| 55 | 0.890166 | 0.109834 |
| 56 | 0.864419 | 0.135581 |
| 57 | 0.840205 | 0.159795 |
| 58 | 0.81758 | 0.18242 |
| 59 | 0.796589 | 0.203411 |
| 60 | 0.777257 | 0.222743 |
| 61 | 0.759574 | 0.240426 |
| 62 | 0.74349 | 0.25651 |
| 63 | 0.728914 | 0.271086 |
| 64 | 1 | 1.16104e-12 |
| 65 | 1 | 1.41977e-12 |
| 66 | 1 | 1.79315e-12 |
| 67 | 1 | 4.65278e-13 |
| 68 | 1 | 3.08781e-12 |
| 69 | 1 | 8.36535e-13 |
| 70 | 1 | 2.3118e-13 |
| 71 | 1 | 3.26504e-13 |
| 72 | 1 | 4.73975e-13 |
| 73 | 1 | 2.86237e-12 |
| 74 | 1 | 4.60695e-12 |
| 75 | 1 | 8.39132e-12 |
| 76 | 1 | 4.26588e-12 |
| 77 | 0.995683 | 0.00431738 |
| 78 | 0.965477 | 0.034523 |
| 79 | 0.936975 | 0.0630249 |
| 80 | 0.910465 | 0.0895353 |
| 81 | 0.886226 | 0.113774 |
| 82 | 0.864456 | 0.135544 |
| 83 | 0.845201 | 0.154799 |
| 84 | 0.828348 | 0.171652 |
| 85 | 0.813662 | 0.186338 |
| 86 | 1 | 4.79341e-12 |
| 87 | 1 | 1.12567e-12 |
| 88 | 1 | 1.34486e-12 |
| 89 | 1 | 1.63114e-12 |
| 90 | 1 | 4.00963e-13 |
| 91 | 1 | 2.49377e-12 |
| 92 | 1 | 3.13701e-12 |
| 93 | 1 | 1.59613e-11 |
| 94 | 1 | 4.10882e-12 |
| 95 | 1 | 1.07205e-12 |
| 96 | 1 | 7.10966e-12 |
| 97 | 1 | 4.81902e-13 |
| 98 | 1 | 2.69527e-12 |
| 99 | 1 | 3.95258e-12 |
| 100 | 1 | 6.29345e-12 |
| 101 | 1 | 1.1914e-11 |
| 102 | 1 | 1.65075e-10 |
| 103 | 0.978036 | 0.0219639 |
| 104 | 0.940498 | 0.0595021 |
| 105 | 0.910237 | 0.0897627 |
| 106 | 0.886754 | 0.113246 |
| 107 | 0.86854 | 0.13146 |
| 108 | 1 | 4.07997e-12 |
| 109 | 1 | 1.86388e-11 |
| 110 | 1 | 1.07589e-12 |
| 111 | 1 | 6.27057e-12 |
| 112 | 1 | 2.94924e-11 |
| 113 | 1 | 6.99098e-12 |
| 114 | 1 | 4.17374e-11 |
| 115 | 1 | 1.00384e-11 |
| 116 | 1 | 1.2158e-13 |
| 117 | 1 | 1.48347e-11 |
| 118 | 1 | 1.82476e-11 |
| 119 | 1 | 9.06164e-13 |
| 120 | 1 | 2.84385e-11 |
| 121 | 1 | 7.23994e-12 |
| 122 | 1 | 3.75418e-11 |
| 123 | 1 | 2.49537e-10 |
| 124 | 1 | 6.88384e-11 |
| 125 | 1 | 5.03474e-10 |
| 126 | 1 | 1.63944e-10 |
| 127 | 1 | 1.37626e-09 |
| 128 | 1 | 2.04338e-08 |
| 129 | 0.922658 | 0.0773415 |
| 130 | 0.896247 | 0.103753 |
| 131 | 0.880562 | 0.119438 |
| 132 | 0.869332 | 0.130668 |
| 133 | 0.860544 | 0.139456 |
| 134 | 0.853293 | 0.146707 |
| 135 | 0.847095 | 0.152905 |
| 136 | 0.841661 | 0.158339 |
| 137 | 0.836805 | 0.163195 |
| 138 | 0.832401 | 0.167599 |
| 139 | 0.828358 | 0.171642 |
| 140 | 1 | 1.1559e-09 |
| 141 | 0.821112 | 0.178888 |
| 142 | 1 | 6.53025e-11 |
| 143 | 1 | 1.95913e-09 |
| 144 | 0.811745 | 0.188255 |
| 145 | 1 | 2.31557e-09 |
| 146 | 0.806211 | 0.193789 |
| 147 | 1 | 1.43748e-10 |
| 148 | 1 | 4.60697e-09 |
| 149 | 1 | 6.08191e-09 |
| 150 | 0.796327 | 0.203673 |
| 151 | 1 | 1.2455e-08 |
| 152 | 1 | 2.13298e-10 |
| 153 | 0.789677 | 0.210323 |
| 154 | 0.899416 | 0.100584 |
| 155 | 0.888227 | 0.111773 |
| 156 | 0.880244 | 0.119756 |
| 157 | 0.781553 | 0.218447 |
| 158 | 0.86861 | 0.13139 |
| 159 | 0.86401 | 0.13599 |
| 160 | 0.859924 | 0.140076 |
| 161 | 0.85623 | 0.14377 |
| 162 | 0.852845 | 0.147155 |
| 163 | 0.849711 | 0.150289 |
| 164 | 0.846784 | 0.153216 |
| 165 | 0.84403 | 0.15597 |
| 166 | 0.841425 | 0.158575 |
| 167 | 0.838947 | 0.161053 |
| 168 | 0.83658 | 0.16342 |
| 169 | 0.834311 | 0.165689 |
| 170 | 0.832129 | 0.167871 |
| 171 | 0.830024 | 0.169976 |
| 172 | 0.827988 | 0.172012 |
| 173 | 0.826016 | 0.173984 |
| 174 | 0.8241 | 0.1759 |
| 175 | 0.822237 | 0.177763 |
| 176 | 0.820422 | 0.179578 |
| 177 | 0.818651 | 0.181349 |
| 178 | 0.81692 | 0.18308 |
| 179 | 0.815228 | 0.184772 |
| 180 | 0.813571 | 0.186429 |
| 181 | 0.811948 | 0.188052 |
| 182 | 0.755677 | 0.244323 |
| 183 | 0.752303 | 0.247697 |
| 184 | 0.749067 | 0.250933 |
| 185 | 0.74596 | 0.25404 |
| 186 | 0.74297 | 0.25703 |
| 187 | 0.866082 | 0.133918 |
| 188 | 0.737306 | 0.262694 |
| 189 | 0.861527 | 0.138473 |
| 190 | 0.732012 | 0.267988 |
| 191 | 0.729487 | 0.270513 |
| 192 | 0.727036 | 0.272964 |
| 193 | 0.853617 | 0.146383 |
| 194 | 0.722335 | 0.277665 |
| 195 | 0.720077 | 0.279923 |
| 196 | 0.717875 | 0.282125 |
| 197 | 0.715727 | 0.284273 |
| 198 | 0.713627 | 0.286373 |
| 199 | 0.843634 | 0.156366 |
| 200 | 0.709567 | 0.290433 |

V4 Type-II-biased

|  |  |  |
| --- | --- | --- |
| 1 | 0 | 1 |
| 2 | 0 | 1 |
| 3 | 0 | 1 |
| 4 | 0 | 1 |
| 5 | 0 | 1 |
| 6 | 0 | 1 |
| 7 | 0 | 1 |
| 8 | 0 | 1 |
| 9 | 0 | 1 |
| 10 | 0 | 1 |
| 11 | 0 | 1 |
| 12 | 0 | 1 |
| 13 | 0 | 1 |
| 14 | 0 | 1 |
| 15 | 0 | 1 |
| 16 | 0 | 1 |
| 17 | 0 | 1 |
| 18 | 0 | 1 |
| 19 | 0 | 1 |
| 20 | 1 | 0 |
| 21 | 1 | 0 |
| 22 | 1 | 0 |
| 23 | 1 | 0 |
| 24 | 1 | 0 |
| 25 | 1 | 0 |
| 26 | 1 | 1 |
| 27 | 1 | 1 |
| 28 | 1 | 1 |
| 29 | 1 | 1 |
| 30 | 1 | 1 |
| 31 | 1 | 1 |
| 32 | 1 | 1 |
| 33 | 1 | 1 |
| 34 | 1 | 1 |
| 35 | 1 | 1 |
| 36 | 1 | 1 |
| 37 | 1 | 1 |
| 38 | 1 | 1 |
| 39 | 1 | 1 |
| 40 | 1 | 1 |
| 41 | 1 | 1 |
| 42 | 2 | 0 |
| 43 | 2 | 0 |
| 44 | 2 | 0 |
| 45 | 2 | 0 |
| 46 | 2 | 0 |
| 47 | 2 | 0 |
| 48 | 2 | 0 |
| 49 | 2 | 0 |
| 50 | 2 | 0 |
| 51 | 2 | 0 |
| 52 | 2 | 1 |
| 53 | 2 | 1 |
| 54 | 2 | 1 |
| 55 | 2 | 1 |
| 56 | 2 | 1 |
| 57 | 2 | 1 |
| 58 | 2 | 1 |
| 59 | 2 | 1 |
| 60 | 2 | 1 |
| 61 | 2 | 1 |
| 62 | 2 | 1 |
| 63 | 2 | 1 |
| 64 | 3 | 0 |
| 65 | 3 | 0 |
| 66 | 3 | 0 |
| 67 | 3 | 0 |
| 68 | 3 | 0 |
| 69 | 3 | 0 |
| 70 | 3 | 0 |
| 71 | 3 | 0 |
| 72 | 3 | 0 |
| 73 | 3 | 0 |
| 74 | 3 | 0 |
| 75 | 3 | 0 |
| 76 | 3 | 0 |
| 77 | 3 | 1 |
| 78 | 3 | 1 |
| 79 | 3 | 1 |
| 80 | 3 | 1 |
| 81 | 3 | 1 |
| 82 | 3 | 1 |
| 83 | 3 | 1 |
| 84 | 3 | 1 |
| 85 | 3 | 1 |
| 86 | 4 | 0 |
| 87 | 4 | 0 |
| 88 | 4 | 0 |
| 89 | 4 | 0 |
| 90 | 4 | 0 |
| 91 | 4 | 0 |
| 92 | 4 | 0 |
| 93 | 4 | 0 |
| 94 | 4 | 0 |
| 95 | 4 | 0 |
| 96 | 4 | 0 |
| 97 | 4 | 0 |
| 98 | 4 | 0 |
| 99 | 4 | 0 |
| 100 | 4 | 0 |
| 101 | 4 | 0 |
| 102 | 4 | 0 |
| 103 | 4 | 1 |
| 104 | 4 | 1 |
| 105 | 4 | 1 |
| 106 | 4 | 1 |
| 107 | 4 | 1 |
| 108 | 5 | 0 |
| 109 | 5 | 0 |
| 110 | 5 | 0 |
| 111 | 5 | 0 |
| 112 | 5 | 0 |
| 113 | 5 | 0 |
| 114 | 5 | 0 |
| 115 | 5 | 0 |
| 116 | 5 | 0 |
| 117 | 5 | 0 |
| 118 | 5 | 0 |
| 119 | 5 | 0 |
| 120 | 5 | 0 |
| 121 | 5 | 0 |
| 122 | 5 | 0 |
| 123 | 5 | 0 |
| 124 | 5 | 0 |
| 125 | 5 | 0 |
| 126 | 5 | 0 |
| 127 | 5 | 0 |
| 128 | 5 | 0 |
| 129 | 5 | 1 |
| 130 | 5 | 1 |
| 131 | 5 | 1 |
| 132 | 5 | 1 |
| 133 | 5 | 1 |
| 134 | 5 | 1 |
| 135 | 5 | 1 |
| 136 | 5 | 1 |
| 137 | 5 | 1 |
| 138 | 5 | 1 |
| 139 | 5 | 1 |
| 140 | 6 | 0 |
| 141 | 5 | 1 |
| 142 | 6 | 0 |
| 143 | 6 | 0 |
| 144 | 5 | 1 |
| 145 | 6 | 0 |
| 146 | 5 | 1 |
| 147 | 6 | 0 |
| 148 | 6 | 0 |
| 149 | 6 | 0 |
| 150 | 5 | 1 |
| 151 | 6 | 0 |
| 152 | 6 | 0 |
| 153 | 5 | 1 |
| 154 | 6 | 1 |
| 155 | 6 | 1 |
| 156 | 6 | 1 |
| 157 | 5 | 1 |
| 158 | 6 | 1 |
| 159 | 6 | 1 |
| 160 | 6 | 1 |
| 161 | 6 | 1 |
| 162 | 6 | 1 |
| 163 | 6 | 1 |
| 164 | 6 | 1 |
| 165 | 6 | 1 |
| 166 | 6 | 1 |
| 167 | 6 | 1 |
| 168 | 6 | 1 |
| 169 | 6 | 1 |
| 170 | 6 | 1 |
| 171 | 6 | 1 |
| 172 | 6 | 1 |
| 173 | 6 | 1 |
| 174 | 6 | 1 |
| 175 | 6 | 1 |
| 176 | 6 | 1 |
| 177 | 6 | 1 |
| 178 | 6 | 1 |
| 179 | 6 | 1 |
| 180 | 6 | 1 |
| 181 | 6 | 1 |
| 182 | 6 | 2 |
| 183 | 6 | 2 |
| 184 | 6 | 2 |
| 185 | 6 | 2 |
| 186 | 6 | 2 |
| 187 | 7 | 1 |
| 188 | 6 | 2 |
| 189 | 7 | 1 |
| 190 | 6 | 2 |
| 191 | 6 | 2 |
| 192 | 6 | 2 |
| 193 | 7 | 1 |
| 194 | 6 | 2 |
| 195 | 6 | 2 |
| 196 | 6 | 2 |
| 197 | 6 | 2 |
| 198 | 6 | 2 |
| 199 | 7 | 1 |
| 200 | 6 | 2 |

V4 Equal

|  |  |  |
| --- | --- | --- |
| 1 | 7.5716e-10 | 1 |
| 2 | 7.98179e-11 | 1 |
| 3 | 2.23382e-10 | 1 |
| 4 | 5.61792e-10 | 1 |
| 5 | 3.7692e-10 | 1 |
| 6 | 2.63914e-10 | 1 |
| 7 | 2.3832e-10 | 1 |
| 8 | 1.7651e-10 | 1 |
| 9 | 5.34127e-10 | 1 |
| 10 | 1.02969e-10 | 1 |
| 11 | 3.23404e-10 | 1 |
| 12 | 2.58482e-10 | 1 |
| 13 | 2.1033e-10 | 1 |
| 14 | 1.74334e-10 | 1 |
| 15 | 1.47299e-10 | 1 |
| 16 | 1.26975e-10 | 1 |
| 17 | 5.58848e-10 | 1 |
| 18 | 4.02153e-10 | 1 |
| 19 | 2.31163e-11 | 1 |
| 20 | 1 | 4.51255e-12 |
| 21 | 1 | 1.0975e-10 |
| 22 | 1 | 1.9958e-10 |
| 23 | 1 | 9.33267e-11 |
| 24 | 1 | 5.32438e-11 |
| 25 | 1 | 4.82198e-11 |
| 26 | 0.981885 | 0.0181149 |
| 27 | 0.936673 | 0.0633269 |
| 28 | 0.8948 | 0.1052 |
| 29 | 0.855948 | 0.144052 |
| 30 | 0.819848 | 0.180152 |
| 31 | 0.786272 | 0.213728 |
| 32 | 0.755029 | 0.244971 |
| 33 | 0.725958 | 0.274042 |
| 34 | 0.698926 | 0.301074 |
| 35 | 0.673819 | 0.326181 |
| 36 | 0.650542 | 0.349458 |
| 37 | 0.62901 | 0.37099 |
| 38 | 0.609144 | 0.390856 |
| 39 | 0.590864 | 0.409136 |
| 40 | 0.574086 | 0.425914 |
| 41 | 0.558717 | 0.441283 |
| 42 | 1 | 2.46712e-12 |
| 43 | 1 | 6.39119e-13 |
| 44 | 1 | 4.48286e-12 |
| 45 | 1 | 1.33654e-12 |
| 46 | 1 | 2.09134e-12 |
| 47 | 1 | 3.42401e-12 |
| 48 | 1 | 5.92469e-12 |
| 49 | 1 | 4.4921e-11 |
| 50 | 1 | 2.09783e-11 |
| 51 | 1 | 1.10004e-10 |
| 52 | 0.976185 | 0.0238151 |
| 53 | 0.946077 | 0.0539234 |
| 54 | 0.917398 | 0.0826025 |
| 55 | 0.890166 | 0.109834 |
| 56 | 0.864419 | 0.135581 |
| 57 | 0.840205 | 0.159795 |
| 58 | 0.81758 | 0.18242 |
| 59 | 0.796589 | 0.203411 |
| 60 | 0.777257 | 0.222743 |
| 61 | 0.759574 | 0.240426 |
| 62 | 0.74349 | 0.25651 |
| 63 | 0.728914 | 0.271086 |
| 64 | 1 | 1.93506e-12 |
| 65 | 1 | 1.18314e-11 |
| 66 | 1 | 2.98858e-12 |
| 67 | 1 | 7.75463e-13 |
| 68 | 1 | 1.02926e-12 |
| 69 | 1 | 1.39422e-12 |
| 70 | 1 | 3.85301e-11 |
| 71 | 1 | 5.44173e-13 |
| 72 | 1 | 1.57992e-11 |
| 73 | 1 | 2.3853e-11 |
| 74 | 1 | 3.83913e-11 |
| 75 | 1 | 1.39855e-11 |
| 76 | 1 | 1.42199e-10 |
| 77 | 0.995683 | 0.00431738 |
| 78 | 0.965477 | 0.034523 |
| 79 | 0.936975 | 0.0630249 |
| 80 | 0.910465 | 0.0895353 |
| 81 | 0.886226 | 0.113774 |
| 82 | 0.864456 | 0.135544 |
| 83 | 0.845201 | 0.154799 |
| 84 | 0.828348 | 0.171652 |
| 85 | 0.813662 | 0.186338 |
| 86 | 1 | 3.19559e-13 |
| 87 | 1 | 3.75225e-11 |
| 88 | 1 | 1.12072e-11 |
| 89 | 1 | 5.43715e-11 |
| 90 | 1 | 6.68272e-11 |
| 91 | 1 | 8.31258e-11 |
| 92 | 1 | 1.04567e-12 |
| 93 | 1 | 2.66023e-11 |
| 94 | 1 | 3.42401e-11 |
| 95 | 1 | 4.46693e-11 |
| 96 | 1 | 5.92479e-11 |
| 97 | 1 | 8.03199e-11 |
| 98 | 1 | 4.49211e-10 |
| 99 | 1 | 3.29387e-11 |
| 100 | 1 | 1.0489e-09 |
| 101 | 1 | 1.58851e-09 |
| 102 | 1 | 3.43762e-10 |
| 103 | 0.978036 | 0.0219638 |
| 104 | 0.940498 | 0.0595021 |
| 105 | 0.910237 | 0.0897627 |
| 106 | 0.886754 | 0.113246 |
| 107 | 0.86854 | 0.13146 |
| 108 | 0.85411 | 0.14589 |
| 109 | 0.84236 | 0.15764 |
| 110 | 0.832536 | 0.167464 |
| 111 | 1 | 2.09027e-10 |
| 112 | 0.816785 | 0.183215 |
| 113 | 1 | 2.9129e-10 |
| 114 | 0.804401 | 0.195599 |
| 115 | 0.799058 | 0.200942 |
| 116 | 1 | 2.02634e-09 |
| 117 | 1 | 1.23622e-10 |
| 118 | 0.785317 | 0.214683 |
| 119 | 1 | 7.55139e-10 |
| 120 | 0.777518 | 0.222482 |
| 121 | 1 | 6.03331e-09 |
| 122 | 1 | 3.12871e-10 |
| 123 | 0.767195 | 0.232805 |
| 124 | 0.764037 | 0.235963 |
| 125 | 1 | 8.39152e-10 |
| 126 | 0.758061 | 0.241939 |
| 127 | 0.755222 | 0.244778 |
| 128 | 0.999999 | 5.44539e-07 |
| 129 | 0.749796 | 0.250204 |
| 130 | 0.896247 | 0.103753 |
| 131 | 0.744664 | 0.255336 |
| 132 | 0.742195 | 0.257805 |
| 133 | 0.860544 | 0.139456 |
| 134 | 0.853293 | 0.146707 |
| 135 | 0.847095 | 0.152905 |
| 136 | 0.841661 | 0.158339 |
| 137 | 0.836805 | 0.163195 |
| 138 | 0.665789 | 0.334211 |
| 139 | 0.828358 | 0.171642 |
| 140 | 0.824612 | 0.175388 |
| 141 | 0.821112 | 0.178888 |
| 142 | 0.651001 | 0.348999 |
| 143 | 0.814705 | 0.185295 |
| 144 | 0.811745 | 0.188255 |
| 145 | 0.808919 | 0.191081 |
| 146 | 0.806211 | 0.193789 |
| 147 | 0.803609 | 0.196391 |
| 148 | 0.8011 | 0.1989 |
| 149 | 0.798675 | 0.201325 |
| 150 | 0.796327 | 0.203673 |
| 151 | 0.623981 | 0.376019 |
| 152 | 0.791834 | 0.208166 |
| 153 | 0.789677 | 0.210323 |
| 154 | 0.787574 | 0.212426 |
| 155 | 0.785522 | 0.214478 |
| 156 | 0.783516 | 0.216484 |
| 157 | 0.60919 | 0.39081 |
| 158 | 0.779631 | 0.220369 |
| 159 | 0.720771 | 0.279229 |
| 160 | 0.775899 | 0.224101 |
| 161 | 0.774085 | 0.225915 |
| 162 | 0.772304 | 0.227696 |
| 163 | 0.706539 | 0.293461 |
| 164 | 0.561855 | 0.438145 |
| 165 | 0.700189 | 0.299811 |
| 166 | 0.697173 | 0.302827 |
| 167 | 0.694254 | 0.305746 |
| 168 | 0.691425 | 0.308575 |
| 169 | 0.68868 | 0.31132 |
| 170 | 0.686012 | 0.313988 |
| 171 | 0.683418 | 0.316582 |
| 172 | 0.680893 | 0.319107 |
| 173 | 0.678432 | 0.321568 |
| 174 | 0.676031 | 0.323969 |
| 175 | 0.673688 | 0.326312 |
| 176 | 0.671398 | 0.328602 |
| 177 | 0.669159 | 0.330841 |
| 178 | 0.666968 | 0.333032 |
| 179 | 0.664823 | 0.335177 |
| 180 | 0.662721 | 0.337279 |
| 181 | 0.660661 | 0.339339 |
| 182 | 0.658639 | 0.341361 |
| 183 | 0.656655 | 0.343345 |
| 184 | 0.654706 | 0.345294 |
| 185 | 0.652792 | 0.347208 |
| 186 | 0.618593 | 0.381407 |
| 187 | 0.615914 | 0.384086 |
| 188 | 0.613299 | 0.386701 |
| 189 | 0.610746 | 0.389254 |
| 190 | 0.608253 | 0.391747 |
| 191 | 0.605817 | 0.394183 |
| 192 | 0.603436 | 0.396564 |
| 193 | 0.601106 | 0.398894 |
| 194 | 0.598826 | 0.401174 |
| 195 | 0.596594 | 0.403406 |
| 196 | 0.594407 | 0.405593 |
| 197 | 0.592265 | 0.407735 |
| 198 | 0.590164 | 0.409836 |
| 199 | 0.588103 | 0.411897 |
| 200 | 0.586081 | 0.413919 |

V4 Equal

|  |  |  |
| --- | --- | --- |
| 1 | 0 | 1 |
| 2 | 0 | 1 |
| 3 | 0 | 1 |
| 4 | 0 | 1 |
| 5 | 0 | 1 |
| 6 | 0 | 1 |
| 7 | 0 | 1 |
| 8 | 0 | 1 |
| 9 | 0 | 1 |
| 10 | 0 | 1 |
| 11 | 0 | 1 |
| 12 | 0 | 1 |
| 13 | 0 | 1 |
| 14 | 0 | 1 |
| 15 | 0 | 1 |
| 16 | 0 | 1 |
| 17 | 0 | 1 |
| 18 | 0 | 1 |
| 19 | 0 | 1 |
| 20 | 1 | 0 |
| 21 | 1 | 0 |
| 22 | 1 | 0 |
| 23 | 1 | 0 |
| 24 | 1 | 0 |
| 25 | 1 | 0 |
| 26 | 1 | 1 |
| 27 | 1 | 1 |
| 28 | 1 | 1 |
| 29 | 1 | 1 |
| 30 | 1 | 1 |
| 31 | 1 | 1 |
| 32 | 1 | 1 |
| 33 | 1 | 1 |
| 34 | 1 | 1 |
| 35 | 1 | 1 |
| 36 | 1 | 1 |
| 37 | 1 | 1 |
| 38 | 1 | 1 |
| 39 | 1 | 1 |
| 40 | 1 | 1 |
| 41 | 1 | 1 |
| 42 | 2 | 0 |
| 43 | 2 | 0 |
| 44 | 2 | 0 |
| 45 | 2 | 0 |
| 46 | 2 | 0 |
| 47 | 2 | 0 |
| 48 | 2 | 0 |
| 49 | 2 | 0 |
| 50 | 2 | 0 |
| 51 | 2 | 0 |
| 52 | 2 | 1 |
| 53 | 2 | 1 |
| 54 | 2 | 1 |
| 55 | 2 | 1 |
| 56 | 2 | 1 |
| 57 | 2 | 1 |
| 58 | 2 | 1 |
| 59 | 2 | 1 |
| 60 | 2 | 1 |
| 61 | 2 | 1 |
| 62 | 2 | 1 |
| 63 | 2 | 1 |
| 64 | 3 | 0 |
| 65 | 3 | 0 |
| 66 | 3 | 0 |
| 67 | 3 | 0 |
| 68 | 3 | 0 |
| 69 | 3 | 0 |
| 70 | 3 | 0 |
| 71 | 3 | 0 |
| 72 | 3 | 0 |
| 73 | 3 | 0 |
| 74 | 3 | 0 |
| 75 | 3 | 0 |
| 76 | 3 | 0 |
| 77 | 3 | 1 |
| 78 | 3 | 1 |
| 79 | 3 | 1 |
| 80 | 3 | 1 |
| 81 | 3 | 1 |
| 82 | 3 | 1 |
| 83 | 3 | 1 |
| 84 | 3 | 1 |
| 85 | 3 | 1 |
| 86 | 4 | 0 |
| 87 | 4 | 0 |
| 88 | 4 | 0 |
| 89 | 4 | 0 |
| 90 | 4 | 0 |
| 91 | 4 | 0 |
| 92 | 4 | 0 |
| 93 | 4 | 0 |
| 94 | 4 | 0 |
| 95 | 4 | 0 |
| 96 | 4 | 0 |
| 97 | 4 | 0 |
| 98 | 4 | 0 |
| 99 | 4 | 0 |
| 100 | 4 | 0 |
| 101 | 4 | 0 |
| 102 | 4 | 0 |
| 103 | 4 | 1 |
| 104 | 4 | 1 |
| 105 | 4 | 1 |
| 106 | 4 | 1 |
| 107 | 4 | 1 |
| 108 | 4 | 1 |
| 109 | 4 | 1 |
| 110 | 4 | 1 |
| 111 | 5 | 0 |
| 112 | 4 | 1 |
| 113 | 5 | 0 |
| 114 | 4 | 1 |
| 115 | 4 | 1 |
| 116 | 5 | 0 |
| 117 | 5 | 0 |
| 118 | 4 | 1 |
| 119 | 5 | 0 |
| 120 | 4 | 1 |
| 121 | 5 | 0 |
| 122 | 5 | 0 |
| 123 | 4 | 1 |
| 124 | 4 | 1 |
| 125 | 5 | 0 |
| 126 | 4 | 1 |
| 127 | 4 | 1 |
| 128 | 5 | 0 |
| 129 | 4 | 1 |
| 130 | 5 | 1 |
| 131 | 4 | 1 |
| 132 | 4 | 1 |
| 133 | 5 | 1 |
| 134 | 5 | 1 |
| 135 | 5 | 1 |
| 136 | 5 | 1 |
| 137 | 5 | 1 |
| 138 | 4 | 2 |
| 139 | 5 | 1 |
| 140 | 5 | 1 |
| 141 | 5 | 1 |
| 142 | 4 | 2 |
| 143 | 5 | 1 |
| 144 | 5 | 1 |
| 145 | 5 | 1 |
| 146 | 5 | 1 |
| 147 | 5 | 1 |
| 148 | 5 | 1 |
| 149 | 5 | 1 |
| 150 | 5 | 1 |
| 151 | 4 | 2 |
| 152 | 5 | 1 |
| 153 | 5 | 1 |
| 154 | 5 | 1 |
| 155 | 5 | 1 |
| 156 | 5 | 1 |
| 157 | 4 | 2 |
| 158 | 5 | 1 |
| 159 | 5 | 2 |
| 160 | 5 | 1 |
| 161 | 5 | 1 |
| 162 | 5 | 1 |
| 163 | 5 | 2 |
| 164 | 4 | 3 |
| 165 | 5 | 2 |
| 166 | 5 | 2 |
| 167 | 5 | 2 |
| 168 | 5 | 2 |
| 169 | 5 | 2 |
| 170 | 5 | 2 |
| 171 | 5 | 2 |
| 172 | 5 | 2 |
| 173 | 5 | 2 |
| 174 | 5 | 2 |
| 175 | 5 | 2 |
| 176 | 5 | 2 |
| 177 | 5 | 2 |
| 178 | 5 | 2 |
| 179 | 5 | 2 |
| 180 | 5 | 2 |
| 181 | 5 | 2 |
| 182 | 5 | 2 |
| 183 | 5 | 2 |
| 184 | 5 | 2 |
| 185 | 5 | 2 |
| 186 | 5 | 3 |
| 187 | 5 | 3 |
| 188 | 5 | 3 |
| 189 | 5 | 3 |
| 190 | 5 | 3 |
| 191 | 5 | 3 |
| 192 | 5 | 3 |
| 193 | 5 | 3 |
| 194 | 5 | 3 |
| 195 | 5 | 3 |
| 196 | 5 | 3 |
| 197 | 5 | 3 |
| 198 | 5 | 3 |
| 199 | 5 | 3 |
| 200 | 5 | 3 |

V4 Type-I-biased

|  |  |  |
| --- | --- | --- |
| 1 | 2.25517e-09 | 1 |
| 2 | 9.48654e-10 | 1 |
| 3 | 5.30866e-10 | 1 |
| 4 | 4.17498e-10 | 1 |
| 5 | 2.80385e-10 | 1 |
| 6 | 1.96523e-10 | 1 |
| 7 | 1.42111e-10 | 1 |
| 8 | 1.05343e-10 | 1 |
| 9 | 3.19007e-10 | 1 |
| 10 | 6.15362e-11 | 1 |
| 11 | 4.83431e-11 | 1 |
| 12 | 1.5462e-10 | 1 |
| 13 | 1.2586e-10 | 1 |
| 14 | 1.0435e-10 | 1 |
| 15 | 8.81869e-11 | 1 |
| 16 | 7.60331e-11 | 1 |
| 17 | 6.69362e-11 | 1 |
| 18 | 6.02153e-11 | 1 |
| 19 | 5.53833e-11 | 1 |
| 20 | 1 | 6.31533e-12 |
| 21 | 1 | 7.67897e-10 |
| 22 | 1 | 2.79171e-10 |
| 23 | 1 | 6.51934e-10 |
| 24 | 1 | 3.70562e-10 |
| 25 | 1 | 6.62152e-11 |
| 26 | 0.981885 | 0.0181149 |
| 27 | 0.936673 | 0.0633268 |
| 28 | 0.8948 | 0.1052 |
| 29 | 0.855948 | 0.144052 |
| 30 | 0.819848 | 0.180152 |
| 31 | 0.786272 | 0.213728 |
| 32 | 0.755029 | 0.244971 |
| 33 | 0.725958 | 0.274042 |
| 34 | 0.698926 | 0.301074 |
| 35 | 0.673819 | 0.326181 |
| 36 | 0.650542 | 0.349458 |
| 37 | 0.62901 | 0.37099 |
| 38 | 0.609144 | 0.390856 |
| 39 | 0.590864 | 0.409136 |
| 40 | 0.574086 | 0.425914 |
| 41 | 0.558717 | 0.441283 |
| 42 | 1 | 3.45397e-12 |
| 43 | 1 | 4.47383e-12 |
| 44 | 1 | 6.27601e-12 |
| 45 | 1 | 9.35581e-12 |
| 46 | 1 | 2.92787e-12 |
| 47 | 1 | 4.79361e-12 |
| 48 | 1 | 1.65893e-12 |
| 49 | 1 | 6.28897e-11 |
| 50 | 1 | 2.93699e-11 |
| 51 | 1 | 6.16021e-10 |
| 52 | 0.976185 | 0.0238153 |
| 53 | 0.946077 | 0.0539234 |
| 54 | 0.917398 | 0.0826025 |
| 55 | 0.890166 | 0.109834 |
| 56 | 0.864419 | 0.135581 |
| 57 | 0.840205 | 0.159795 |
| 58 | 0.81758 | 0.18242 |
| 59 | 0.796589 | 0.203411 |
| 60 | 0.777257 | 0.222743 |
| 61 | 0.759574 | 0.240426 |
| 62 | 0.74349 | 0.25651 |
| 63 | 0.728914 | 0.271086 |
| 64 | 1 | 1.35454e-11 |
| 65 | 1 | 1.65639e-11 |
| 66 | 1 | 1.6736e-11 |
| 67 | 1 | 2.71413e-11 |
| 68 | 1 | 1.44097e-10 |
| 69 | 1 | 1.95191e-12 |
| 70 | 1 | 5.39427e-11 |
| 71 | 1 | 7.61842e-11 |
| 72 | 1 | 1.10594e-10 |
| 73 | 1 | 8.3486e-12 |
| 74 | 1 | 2.14991e-10 |
| 75 | 1 | 3.91592e-10 |
| 76 | 1 | 9.95391e-10 |
| 77 | 0.995683 | 0.00431731 |
| 78 | 0.965477 | 0.034523 |
| 79 | 0.936975 | 0.0630249 |
| 80 | 0.910465 | 0.0895353 |
| 81 | 0.886226 | 0.113774 |
| 82 | 0.864456 | 0.135544 |
| 83 | 0.845201 | 0.154799 |
| 84 | 0.828348 | 0.171652 |
| 85 | 0.813662 | 0.186338 |
| 86 | 0.800852 | 0.199148 |
| 87 | 0.789621 | 0.210379 |
| 88 | 0.779699 | 0.220301 |
| 89 | 0.770855 | 0.229145 |
| 90 | 0.762899 | 0.237101 |
| 91 | 0.755677 | 0.244323 |
| 92 | 0.749067 | 0.250933 |
| 93 | 0.74297 | 0.25703 |
| 94 | 0.737306 | 0.262694 |
| 95 | 0.732012 | 0.267988 |
| 96 | 0.727036 | 0.272964 |
| 97 | 0.722335 | 0.277665 |
| 98 | 0.717875 | 0.282125 |
| 99 | 0.713627 | 0.286373 |
| 100 | 0.709567 | 0.290433 |
| 101 | 0.705675 | 0.294325 |
| 102 | 0.701933 | 0.298067 |
| 103 | 0.698327 | 0.301673 |
| 104 | 0.694844 | 0.305156 |
| 105 | 0.691474 | 0.308526 |
| 106 | 0.688208 | 0.311792 |
| 107 | 0.685037 | 0.314963 |
| 108 | 0.681955 | 0.318045 |
| 109 | 0.678955 | 0.321045 |
| 110 | 0.676031 | 0.323969 |
| 111 | 0.618443 | 0.381557 |
| 112 | 0.613324 | 0.386676 |
| 113 | 0.608416 | 0.391584 |
| 114 | 0.603708 | 0.396292 |
| 115 | 0.599187 | 0.400813 |
| 116 | 0.59484 | 0.40516 |
| 117 | 0.590656 | 0.409344 |
| 118 | 0.586625 | 0.413375 |
| 119 | 0.582737 | 0.417263 |
| 120 | 0.578983 | 0.421017 |
| 121 | 0.575353 | 0.424647 |
| 122 | 0.571841 | 0.428159 |
| 123 | 0.568439 | 0.431561 |
| 124 | 0.56514 | 0.43486 |
| 125 | 0.561937 | 0.438063 |
| 126 | 0.558827 | 0.441173 |
| 127 | 0.555802 | 0.444198 |
| 128 | 0.552858 | 0.447142 |
| 129 | 0.54999 | 0.45001 |
| 130 | 0.547195 | 0.452805 |
| 131 | 0.544468 | 0.455532 |
| 132 | 0.541807 | 0.458193 |
| 133 | 0.539207 | 0.460793 |
| 134 | 0.536665 | 0.463335 |
| 135 | 0.53418 | 0.46582 |
| 136 | 0.531747 | 0.468253 |
| 137 | 0.529365 | 0.470635 |
| 138 | 0.527032 | 0.472968 |
| 139 | 0.496169 | 0.503831 |
| 140 | 0.493119 | 0.506881 |
| 141 | 0.490148 | 0.509852 |
| 142 | 0.487252 | 0.512748 |
| 143 | 0.484429 | 0.515571 |
| 144 | 0.481675 | 0.518325 |
| 145 | 0.478988 | 0.521012 |
| 146 | 0.476364 | 0.523636 |
| 147 | 0.473802 | 0.526198 |
| 148 | 0.471299 | 0.528701 |
| 149 | 0.468852 | 0.531148 |
| 150 | 0.466459 | 0.533541 |
| 151 | 0.464117 | 0.535883 |
| 152 | 0.461826 | 0.538174 |
| 153 | 0.459583 | 0.540417 |
| 154 | 0.457385 | 0.542615 |
| 155 | 0.455232 | 0.544768 |
| 156 | 0.453121 | 0.546879 |
| 157 | 0.451052 | 0.548948 |
| 158 | 0.449021 | 0.550979 |
| 159 | 0.447029 | 0.552971 |
| 160 | 0.445073 | 0.554927 |
| 161 | 0.443153 | 0.556847 |
| 162 | 0.441266 | 0.558734 |
| 163 | 0.439413 | 0.560587 |
| 164 | 0.437591 | 0.562409 |
| 165 | 0.4358 | 0.5642 |
| 166 | 0.434038 | 0.565962 |
| 167 | 0.414587 | 0.585413 |
| 168 | 0.412509 | 0.587491 |
| 169 | 0.410471 | 0.589529 |
| 170 | 0.408471 | 0.591529 |
| 171 | 0.406509 | 0.593491 |
| 172 | 0.404584 | 0.595416 |
| 173 | 0.402693 | 0.597307 |
| 174 | 0.400836 | 0.599164 |
| 175 | 0.399013 | 0.600987 |
| 176 | 0.397222 | 0.602778 |
| 177 | 0.395461 | 0.604539 |
| 178 | 0.393731 | 0.606269 |
| 179 | 0.39203 | 0.60797 |
| 180 | 0.390358 | 0.609642 |
| 181 | 0.388713 | 0.611287 |
| 182 | 0.387094 | 0.612906 |
| 183 | 0.385502 | 0.614498 |
| 184 | 0.383935 | 0.616065 |
| 185 | 0.382392 | 0.617608 |
| 186 | 0.380873 | 0.619127 |
| 187 | 0.379377 | 0.620623 |
| 188 | 0.377904 | 0.622096 |
| 189 | 0.376452 | 0.623548 |
| 190 | 0.375022 | 0.624978 |
| 191 | 0.373612 | 0.626388 |
| 192 | 0.372222 | 0.627778 |
| 193 | 0.370852 | 0.629148 |
| 194 | 0.369501 | 0.630499 |
| 195 | 0.356061 | 0.643939 |
| 196 | 0.354545 | 0.645455 |
| 197 | 0.353052 | 0.646948 |
| 198 | 0.351582 | 0.648418 |
| 199 | 0.350133 | 0.649867 |
| 200 | 0.348707 | 0.651293 |

V4 Type-I-biased

|  |  |  |
| --- | --- | --- |
| 1 | 0 | 1 |
| 2 | 0 | 1 |
| 3 | 0 | 1 |
| 4 | 0 | 1 |
| 5 | 0 | 1 |
| 6 | 0 | 1 |
| 7 | 0 | 1 |
| 8 | 0 | 1 |
| 9 | 0 | 1 |
| 10 | 0 | 1 |
| 11 | 0 | 1 |
| 12 | 0 | 1 |
| 13 | 0 | 1 |
| 14 | 0 | 1 |
| 15 | 0 | 1 |
| 16 | 0 | 1 |
| 17 | 0 | 1 |
| 18 | 0 | 1 |
| 19 | 0 | 1 |
| 20 | 1 | 0 |
| 21 | 1 | 0 |
| 22 | 1 | 0 |
| 23 | 1 | 0 |
| 24 | 1 | 0 |
| 25 | 1 | 0 |
| 26 | 1 | 1 |
| 27 | 1 | 1 |
| 28 | 1 | 1 |
| 29 | 1 | 1 |
| 30 | 1 | 1 |
| 31 | 1 | 1 |
| 32 | 1 | 1 |
| 33 | 1 | 1 |
| 34 | 1 | 1 |
| 35 | 1 | 1 |
| 36 | 1 | 1 |
| 37 | 1 | 1 |
| 38 | 1 | 1 |
| 39 | 1 | 1 |
| 40 | 1 | 1 |
| 41 | 1 | 1 |
| 42 | 2 | 0 |
| 43 | 2 | 0 |
| 44 | 2 | 0 |
| 45 | 2 | 0 |
| 46 | 2 | 0 |
| 47 | 2 | 0 |
| 48 | 2 | 0 |
| 49 | 2 | 0 |
| 50 | 2 | 0 |
| 51 | 2 | 0 |
| 52 | 2 | 1 |
| 53 | 2 | 1 |
| 54 | 2 | 1 |
| 55 | 2 | 1 |
| 56 | 2 | 1 |
| 57 | 2 | 1 |
| 58 | 2 | 1 |
| 59 | 2 | 1 |
| 60 | 2 | 1 |
| 61 | 2 | 1 |
| 62 | 2 | 1 |
| 63 | 2 | 1 |
| 64 | 3 | 0 |
| 65 | 3 | 0 |
| 66 | 3 | 0 |
| 67 | 3 | 0 |
| 68 | 3 | 0 |
| 69 | 3 | 0 |
| 70 | 3 | 0 |
| 71 | 3 | 0 |
| 72 | 3 | 0 |
| 73 | 3 | 0 |
| 74 | 3 | 0 |
| 75 | 3 | 0 |
| 76 | 3 | 0 |
| 77 | 3 | 1 |
| 78 | 3 | 1 |
| 79 | 3 | 1 |
| 80 | 3 | 1 |
| 81 | 3 | 1 |
| 82 | 3 | 1 |
| 83 | 3 | 1 |
| 84 | 3 | 1 |
| 85 | 3 | 1 |
| 86 | 3 | 1 |
| 87 | 3 | 1 |
| 88 | 3 | 1 |
| 89 | 3 | 1 |
| 90 | 3 | 1 |
| 91 | 3 | 1 |
| 92 | 3 | 1 |
| 93 | 3 | 1 |
| 94 | 3 | 1 |
| 95 | 3 | 1 |
| 96 | 3 | 1 |
| 97 | 3 | 1 |
| 98 | 3 | 1 |
| 99 | 3 | 1 |
| 100 | 3 | 1 |
| 101 | 3 | 1 |
| 102 | 3 | 1 |
| 103 | 3 | 1 |
| 104 | 3 | 1 |
| 105 | 3 | 1 |
| 106 | 3 | 1 |
| 107 | 3 | 1 |
| 108 | 3 | 1 |
| 109 | 3 | 1 |
| 110 | 3 | 1 |
| 111 | 3 | 2 |
| 112 | 3 | 2 |
| 113 | 3 | 2 |
| 114 | 3 | 2 |
| 115 | 3 | 2 |
| 116 | 3 | 2 |
| 117 | 3 | 2 |
| 118 | 3 | 2 |
| 119 | 3 | 2 |
| 120 | 3 | 2 |
| 121 | 3 | 2 |
| 122 | 3 | 2 |
| 123 | 3 | 2 |
| 124 | 3 | 2 |
| 125 | 3 | 2 |
| 126 | 3 | 2 |
| 127 | 3 | 2 |
| 128 | 3 | 2 |
| 129 | 3 | 2 |
| 130 | 3 | 2 |
| 131 | 3 | 2 |
| 132 | 3 | 2 |
| 133 | 3 | 2 |
| 134 | 3 | 2 |
| 135 | 3 | 2 |
| 136 | 3 | 2 |
| 137 | 3 | 2 |
| 138 | 3 | 2 |
| 139 | 3 | 3 |
| 140 | 3 | 3 |
| 141 | 3 | 3 |
| 142 | 3 | 3 |
| 143 | 3 | 3 |
| 144 | 3 | 3 |
| 145 | 3 | 3 |
| 146 | 3 | 3 |
| 147 | 3 | 3 |
| 148 | 3 | 3 |
| 149 | 3 | 3 |
| 150 | 3 | 3 |
| 151 | 3 | 3 |
| 152 | 3 | 3 |
| 153 | 3 | 3 |
| 154 | 3 | 3 |
| 155 | 3 | 3 |
| 156 | 3 | 3 |
| 157 | 3 | 3 |
| 158 | 3 | 3 |
| 159 | 3 | 3 |
| 160 | 3 | 3 |
| 161 | 3 | 3 |
| 162 | 3 | 3 |
| 163 | 3 | 3 |
| 164 | 3 | 3 |
| 165 | 3 | 3 |
| 166 | 3 | 3 |
| 167 | 3 | 4 |
| 168 | 3 | 4 |
| 169 | 3 | 4 |
| 170 | 3 | 4 |
| 171 | 3 | 4 |
| 172 | 3 | 4 |
| 173 | 3 | 4 |
| 174 | 3 | 4 |
| 175 | 3 | 4 |
| 176 | 3 | 4 |
| 177 | 3 | 4 |
| 178 | 3 | 4 |
| 179 | 3 | 4 |
| 180 | 3 | 4 |
| 181 | 3 | 4 |
| 182 | 3 | 4 |
| 183 | 3 | 4 |
| 184 | 3 | 4 |
| 185 | 3 | 4 |
| 186 | 3 | 4 |
| 187 | 3 | 4 |
| 188 | 3 | 4 |
| 189 | 3 | 4 |
| 190 | 3 | 4 |
| 191 | 3 | 4 |
| 192 | 3 | 4 |
| 193 | 3 | 4 |
| 194 | 3 | 4 |
| 195 | 3 | 5 |
| 196 | 3 | 5 |
| 197 | 3 | 5 |
| 198 | 3 | 5 |
| 199 | 3 | 5 |
| 200 | 3 | 5 |

Figure S7 | Same as Figure S4 for model version 4.

**Model Version 5-1**

V5-1 Functions

|  |  |  |
| --- | --- | --- |
| 0 | 0.000793255 | 5.37764e-16 |
| 1 | 0.00209324 | 0.00564835 |
| 2 | 0.0035496 | 0.0123587 |
| 3 | 0.00518107 | 0.02033 |
| 4 | 0.00700865 | 0.0297984 |
| 5 | 0.00905583 | 0.0410439 |
| 6 | 0.0113489 | 0.0543982 |
| 7 | 0.0139172 | 0.0702543 |
| 8 | 0.0167937 | 0.0890775 |
| 9 | 0.020015 | 0.111418 |
| 10 | 0.0236224 | 0.137927 |
| 11 | 0.0276617 | 0.169371 |
| 12 | 0.0321842 | 0.206657 |
| 13 | 0.0372472 | 0.25085 |
| 14 | 0.0429146 | 0.303204 |
| 15 | 0.0492578 | 0.365187 |
| 16 | 0.0563564 | 0.438518 |
| 17 | 0.0642989 | 0.525202 |
| 18 | 0.0731842 | 0.627567 |
| 19 | 0.0831221 | 0.748307 |
| 20 | 0.0942348 | 0.890523 |
| 21 | 0.106658 | 1.05776 |
| 22 | 0.120542 | 1.25404 |
| 23 | 0.136055 | 1.48388 |
| 24 | 0.15338 | 1.75231 |
| 25 | 0.172723 | 2.06484 |
| 26 | 0.194307 | 2.42741 |
| 27 | 0.218383 | 2.84627 |
| 28 | 0.245221 | 3.32783 |
| 29 | 0.275122 | 3.87841 |
| 30 | 0.308411 | 4.50392 |
| 31 | 0.345445 | 5.20945 |
| 32 | 0.38661 | 5.99882 |
| 33 | 0.432324 | 6.874 |
| 34 | 0.483037 | 7.83459 |
| 35 | 0.53923 | 8.87735 |
| 36 | 0.601416 | 9.99583 |
| 37 | 0.670138 | 11.1802 |
| 38 | 0.745962 | 12.4175 |
| 39 | 0.829478 | 13.6917 |
| 40 | 0.921293 | 14.985 |
| 41 | 1.02202 | 16.2783 |
| 42 | 1.13227 | 17.5525 |
| 43 | 1.25264 | 18.7898 |
| 44 | 1.3837 | 19.9742 |
| 45 | 1.52598 | 21.0927 |
| 46 | 1.67994 | 22.1354 |
| 47 | 1.84594 | 23.096 |
| 48 | 2.02427 | 23.9712 |
| 49 | 2.21505 | 24.7605 |
| 50 | 2.41828 | 25.4661 |
| 51 | 2.63377 | 26.0916 |
| 52 | 2.86114 | 26.6422 |
| 53 | 3.09981 | 27.1237 |
| 54 | 3.34898 | 27.5426 |
| 55 | 3.60765 | 27.9052 |
| 56 | 3.8746 | 28.2177 |
| 57 | 4.14844 | 28.4861 |
| 58 | 4.4276 | 28.716 |
| 59 | 4.71038 | 28.9122 |
| 60 | 4.995 | 29.0795 |
| 61 | 5.27962 | 29.2217 |
| 62 | 5.5624 | 29.3424 |
| 63 | 5.84156 | 29.4448 |
| 64 | 6.1154 | 29.5315 |
| 65 | 6.38235 | 29.6048 |
| 66 | 6.64102 | 29.6668 |
| 67 | 6.89019 | 29.7191 |
| 68 | 7.12886 | 29.7633 |
| 69 | 7.35623 | 29.8006 |
| 70 | 7.57172 | 29.8321 |
| 71 | 7.77495 | 29.8586 |
| 72 | 7.96573 | 29.8809 |
| 73 | 8.14406 | 29.8997 |
| 74 | 8.31006 | 29.9156 |
| 75 | 8.46402 | 29.929 |
| 76 | 8.6063 | 29.9402 |
| 77 | 8.73736 | 29.9497 |
| 78 | 8.85773 | 29.9576 |
| 79 | 8.96798 | 29.9644 |
| 80 | 9.06871 | 29.97 |
| 81 | 9.16052 | 29.9748 |
| 82 | 9.24404 | 29.9788 |
| 83 | 9.31986 | 29.9821 |
| 84 | 9.38858 | 29.985 |
| 85 | 9.45077 | 29.9873 |
| 86 | 9.50696 | 29.9893 |
| 87 | 9.55768 | 29.991 |
| 88 | 9.60339 | 29.9925 |
| 89 | 9.64455 | 29.9937 |
| 90 | 9.68159 | 29.9947 |
| 91 | 9.71488 | 29.9955 |
| 92 | 9.74478 | 29.9962 |
| 93 | 9.77162 | 29.9968 |
| 94 | 9.79569 | 29.9973 |
| 95 | 9.81728 | 29.9977 |
| 96 | 9.83662 | 29.9981 |
| 97 | 9.85395 | 29.9984 |
| 98 | 9.86946 | 29.9987 |
| 99 | 9.88334 | 29.9989 |
| 100 | 9.89577 | 29.9991 |
| 101 | 9.90688 | 29.9992 |
| 102 | 9.91682 | 29.9993 |
| 103 | 9.9257 | 29.9994 |
| 104 | 9.93364 | 29.9995 |
| 105 | 9.94074 | 29.9996 |
| 106 | 9.94709 | 29.9997 |
| 107 | 9.95275 | 29.9997 |
| 108 | 9.95782 | 29.9998 |
| 109 | 9.96234 | 29.9998 |
| 110 | 9.96638 | 29.9998 |
| 111 | 9.96998 | 29.9999 |
| 112 | 9.97321 | 29.9999 |
| 113 | 9.97608 | 29.9999 |
| 114 | 9.97865 | 29.9999 |
| 115 | 9.98094 | 29.9999 |
| 116 | 9.98299 | 29.9999 |
| 117 | 9.98482 | 29.9999 |
| 118 | 9.98645 | 30 |
| 119 | 9.98791 | 30 |
| 120 | 9.98921 | 30 |
| 121 | 9.99037 | 30 |
| 122 | 9.9914 | 30 |
| 123 | 9.99233 | 30 |
| 124 | 9.99315 | 30 |
| 125 | 9.99389 | 30 |
| 126 | 9.99455 | 30 |
| 127 | 9.99513 | 30 |
| 128 | 9.99566 | 30 |
| 129 | 9.99612 | 30 |
| 130 | 9.99654 | 30 |
| 131 | 9.99691 | 30 |
| 132 | 9.99724 | 30 |
| 133 | 9.99754 | 30 |
| 134 | 9.99781 | 30 |
| 135 | 9.99804 | 30 |
| 136 | 9.99825 | 30 |
| 137 | 9.99844 | 30 |
| 138 | 9.99861 | 30 |
| 139 | 9.99876 | 30 |
| 140 | 9.99889 | 30 |
| 141 | 9.99901 | 30 |
| 142 | 9.99912 | 30 |
| 143 | 9.99921 | 30 |
| 144 | 9.9993 | 30 |
| 145 | 9.99937 | 30 |
| 146 | 9.99944 | 30 |
| 147 | 9.9995 | 30 |
| 148 | 9.99955 | 30 |
| 149 | 9.9996 | 30 |
| 150 | 9.99965 | 30 |
| 151 | 9.99968 | 30 |
| 152 | 9.99972 | 30 |
| 153 | 9.99975 | 30 |
| 154 | 9.99977 | 30 |
| 155 | 9.9998 | 30 |
| 156 | 9.99982 | 30 |
| 157 | 9.99984 | 30 |
| 158 | 9.99986 | 30 |
| 159 | 9.99987 | 30 |
| 160 | 9.99989 | 30 |
| 161 | 9.9999 | 30 |
| 162 | 9.99991 | 30 |
| 163 | 9.99992 | 30 |
| 164 | 9.99993 | 30 |
| 165 | 9.99994 | 30 |
| 166 | 9.99994 | 30 |
| 167 | 9.99995 | 30 |
| 168 | 9.99995 | 30 |
| 169 | 9.99996 | 30 |
| 170 | 9.99996 | 30 |
| 171 | 9.99997 | 30 |
| 172 | 9.99997 | 30 |
| 173 | 9.99997 | 30 |
| 174 | 9.99998 | 30 |
| 175 | 9.99998 | 30 |
| 176 | 9.99998 | 30 |
| 177 | 9.99998 | 30 |
| 178 | 9.99999 | 30 |
| 179 | 9.99999 | 30 |
| 180 | 9.99999 | 30 |
| 181 | 9.99999 | 30 |
| 182 | 9.99999 | 30 |
| 183 | 9.99999 | 30 |
| 184 | 9.99999 | 30 |
| 185 | 9.99999 | 30 |
| 186 | 9.99999 | 30 |
| 187 | 9.99999 | 30 |
| 188 | 10 | 30 |
| 189 | 10 | 30 |
| 190 | 10 | 30 |
| 191 | 10 | 30 |
| 192 | 10 | 30 |
| 193 | 10 | 30 |
| 194 | 10 | 30 |
| 195 | 10 | 30 |
| 196 | 10 | 30 |
| 197 | 10 | 30 |
| 198 | 10 | 30 |
| 199 | 10 | 30 |

V5-1 Equal

|  |  |  |
| --- | --- | --- |
| 2 | 0 | 1 |
| 4 | 0 | 1 |
| 6 | 0 | 1 |
| 8 | 0 | 1 |
| 10 | 0 | 1 |
| 12 | 0 | 1 |
| 14 | 0 | 1 |
| 16 | 0 | 1 |
| 18 | 0 | 1 |
| 20 | 0 | 1 |
| 22 | 0 | 1 |
| 24 | 0 | 1 |
| 26 | 0 | 1 |
| 28 | 0 | 1 |
| 30 | 0 | 1 |
| 32 | 0 | 1 |
| 34 | 0 | 1 |
| 36 | 0 | 1 |
| 38 | 0 | 1 |
| 40 | 0 | 1 |
| 42 | 0 | 1 |
| 44 | 0 | 1 |
| 46 | 0 | 1 |
| 48 | 0 | 1 |
| 50 | 0 | 1 |
| 52 | 0 | 1 |
| 54 | 0 | 1 |
| 56 | 0 | 1 |
| 58 | 0 | 1 |
| 60 | 0 | 1 |
| 62 | 0 | 1 |
| 64 | 0 | 1 |
| 66 | 0 | 1 |
| 68 | 0 | 1 |
| 70 | 0 | 1 |
| 72 | 0 | 1 |
| 74 | 0 | 1 |
| 76 | 0 | 1 |
| 78 | 0 | 1 |
| 80 | 0 | 1 |
| 82 | 0 | 2 |
| 84 | 0 | 2 |
| 86 | 0 | 2 |
| 88 | 0 | 2 |
| 90 | 0 | 2 |
| 92 | 0 | 2 |
| 94 | 0 | 2 |
| 96 | 0 | 2 |
| 98 | 0 | 2 |
| 100 | 0 | 2 |
| 102 | 0 | 2 |
| 104 | 0 | 2 |
| 106 | 0 | 2 |
| 108 | 0 | 2 |
| 110 | 0 | 2 |
| 112 | 0 | 2 |
| 114 | 0 | 2 |
| 116 | 0 | 2 |
| 118 | 0 | 2 |
| 120 | 0 | 2 |
| 122 | 0 | 2 |
| 124 | 0 | 2 |
| 126 | 0 | 2 |
| 128 | 0 | 2 |
| 130 | 0 | 2 |
| 132 | 0 | 3 |
| 134 | 0 | 3 |
| 136 | 0 | 3 |
| 138 | 0 | 3 |
| 140 | 0 | 3 |
| 142 | 0 | 3 |
| 144 | 0 | 3 |
| 146 | 0 | 3 |
| 148 | 0 | 3 |
| 150 | 0 | 3 |
| 152 | 0 | 3 |
| 154 | 0 | 3 |
| 156 | 0 | 3 |
| 158 | 0 | 3 |
| 160 | 0 | 3 |
| 162 | 0 | 3 |
| 164 | 0 | 3 |
| 166 | 0 | 3 |
| 168 | 0 | 3 |
| 170 | 0 | 3 |
| 172 | 0 | 3 |
| 174 | 0 | 3 |
| 176 | 0 | 3 |
| 178 | 0 | 3 |
| 180 | 0 | 3 |
| 182 | 0 | 3 |
| 184 | 0 | 4 |
| 186 | 0 | 4 |
| 188 | 0 | 4 |
| 190 | 0 | 4 |
| 192 | 0 | 4 |
| 194 | 0 | 4 |
| 196 | 0 | 4 |
| 198 | 0 | 4 |
| 200 | 0 | 4 |
| 202 | 0 | 4 |
| 204 | 0 | 4 |
| 206 | 0 | 4 |
| 208 | 0 | 4 |
| 210 | 0 | 4 |
| 212 | 0 | 4 |
| 214 | 0 | 4 |
| 216 | 0 | 4 |
| 218 | 0 | 4 |
| 220 | 0 | 4 |
| 222 | 0 | 4 |
| 224 | 0 | 4 |
| 226 | 0 | 4 |
| 228 | 0 | 4 |
| 230 | 0 | 4 |
| 232 | 0 | 4 |
| 234 | 0 | 4 |
| 236 | 0 | 5 |
| 238 | 0 | 5 |
| 240 | 0 | 5 |
| 242 | 0 | 5 |
| 244 | 0 | 5 |
| 246 | 0 | 5 |
| 248 | 0 | 5 |
| 250 | 0 | 5 |
| 252 | 0 | 5 |
| 254 | 0 | 5 |
| 256 | 0 | 5 |
| 258 | 0 | 5 |
| 260 | 0 | 5 |
| 262 | 0 | 5 |
| 264 | 0 | 5 |
| 266 | 0 | 5 |
| 268 | 0 | 5 |
| 270 | 0 | 5 |
| 272 | 0 | 5 |
| 274 | 0 | 5 |
| 276 | 0 | 5 |
| 278 | 0 | 5 |
| 280 | 0 | 5 |
| 282 | 0 | 5 |
| 284 | 0 | 5 |
| 286 | 0 | 5 |
| 288 | 0 | 5 |
| 290 | 0 | 5 |
| 292 | 0 | 5 |
| 294 | 0 | 5 |
| 296 | 0 | 5 |
| 298 | 0 | 5 |
| 300 | 0 | 5 |
| 302 | 0 | 5 |
| 304 | 0 | 5 |
| 306 | 0 | 5 |
| 308 | 0 | 5 |
| 310 | 0 | 5 |
| 312 | 0 | 5 |
| 314 | 0 | 5 |
| 316 | 0 | 5 |
| 318 | 0 | 5 |
| 320 | 0 | 5 |
| 322 | 0 | 5 |
| 324 | 0 | 5 |
| 326 | 0 | 5 |
| 328 | 0 | 5 |
| 330 | 0 | 5 |
| 332 | 0 | 5 |
| 334 | 0 | 5 |
| 336 | 0 | 5 |
| 338 | 0 | 5 |
| 340 | 0 | 5 |
| 342 | 0 | 5 |
| 344 | 0 | 5 |
| 346 | 0 | 5 |
| 348 | 0 | 5 |
| 350 | 0 | 5 |
| 352 | 1 | 5 |
| 354 | 1 | 5 |
| 356 | 1 | 5 |
| 358 | 1 | 5 |
| 360 | 1 | 5 |
| 362 | 1 | 5 |
| 364 | 1 | 5 |
| 366 | 1 | 5 |
| 368 | 1 | 5 |
| 370 | 1 | 5 |
| 372 | 1 | 5 |
| 374 | 1 | 5 |
| 376 | 1 | 5 |
| 378 | 1 | 5 |
| 380 | 1 | 5 |
| 382 | 1 | 5 |
| 384 | 1 | 5 |
| 386 | 1 | 5 |
| 388 | 1 | 5 |
| 390 | 1 | 5 |
| 392 | 1 | 5 |
| 394 | 1 | 5 |
| 396 | 1 | 5 |
| 398 | 1 | 5 |
| 400 | 1 | 5 |

V5-1 Equal

|  |  |  |
| --- | --- | --- |
| 2 | 2.781e-09 | 1 |
| 4 | 1.18932e-09 | 1 |
| 6 | 5.46905e-10 | 1 |
| 8 | 2.85225e-10 | 1 |
| 10 | 6.3895e-10 | 1 |
| 12 | 3.74799e-10 | 1 |
| 14 | 2.27243e-10 | 1 |
| 16 | 1.41325e-10 | 1 |
| 18 | 1.12189e-10 | 1 |
| 20 | 2.9036e-10 | 1 |
| 22 | 4.78182e-11 | 1 |
| 24 | 1.28379e-10 | 1 |
| 26 | 8.79731e-11 | 1 |
| 28 | 3.08804e-10 | 1 |
| 30 | 2.232e-10 | 1 |
| 32 | 3.34234e-11 | 1 |
| 34 | 1.3053e-10 | 1 |
| 36 | 2.14278e-11 | 1 |
| 38 | 1.86013e-11 | 1 |
| 40 | 4.2885e-12 | 1 |
| 42 | 2.10369e-11 | 1 |
| 44 | 2.19148e-11 | 1 |
| 46 | 9.64793e-11 | 1 |
| 48 | 2.78531e-11 | 1 |
| 50 | 6.6961e-12 | 1 |
| 52 | 8.31413e-12 | 1 |
| 54 | 1.05889e-11 | 1 |
| 56 | 1.37554e-11 | 1 |
| 58 | 1.81379e-11 | 1 |
| 60 | 2.41967e-11 | 1 |
| 62 | 2.60573e-11 | 1 |
| 64 | 3.53313e-11 | 1 |
| 66 | 4.8203e-11 | 1 |
| 68 | 6.61412e-11 | 1 |
| 70 | 2.28193e-11 | 1 |
| 72 | 3.16917e-11 | 1 |
| 74 | 3.5475e-11 | 1 |
| 76 | 1.00158e-11 | 1 |
| 78 | 1.42622e-11 | 1 |
| 80 | 1.12197e-11 | 1 |
| 82 | 7.58841e-14 | 1 |
| 84 | 7.5744e-14 | 1 |
| 86 | 7.675e-14 | 1 |
| 88 | 7.89034e-14 | 1 |
| 90 | 8.225e-14 | 1 |
| 92 | 4.34239e-13 | 1 |
| 94 | 9.28138e-14 | 1 |
| 96 | 1.00292e-13 | 1 |
| 98 | 5.47347e-13 | 1 |
| 100 | 6.0285e-13 | 1 |
| 102 | 6.69363e-13 | 1 |
| 104 | 1.49712e-13 | 1 |
| 106 | 8.425e-13 | 1 |
| 108 | 1.90704e-13 | 1 |
| 110 | 1.0845e-12 | 1 |
| 112 | 1.23879e-12 | 1 |
| 114 | 1.42039e-12 | 1 |
| 116 | 3.26793e-13 | 1 |
| 118 | 3.7703e-13 | 1 |
| 120 | 4.36125e-13 | 1 |
| 122 | 5.05574e-13 | 1 |
| 124 | 5.87339e-13 | 1 |
| 126 | 6.83611e-13 | 1 |
| 128 | 7.96992e-13 | 1 |
| 130 | 9.30692e-13 | 1 |
| 132 | 2.63019e-13 | 1 |
| 134 | 2.69993e-13 | 1.00001 |
| 136 | 1.39393e-12 | 0.999993 |
| 138 | 1.44746e-12 | 1 |
| 140 | 3.02221e-13 | 1.00001 |
| 142 | 3.17109e-13 | 0.999993 |
| 144 | 3.34312e-13 | 1 |
| 146 | 3.54041e-13 | 1.00001 |
| 148 | 1.88233e-12 | 0.999993 |
| 150 | 4.019e-13 | 1 |
| 152 | 4.30559e-13 | 1.00001 |
| 154 | 4.62825e-13 | 0.999994 |
| 156 | 3.99231e-13 | 1 |
| 158 | 4.31677e-13 | 1.00001 |
| 160 | 4.67969e-13 | 0.999994 |
| 162 | 6.35679e-13 | 1 |
| 164 | 6.92287e-13 | 1.00001 |
| 166 | 7.55452e-13 | 0.999994 |
| 168 | 6.60685e-13 | 1 |
| 170 | 9.04353e-13 | 1.00001 |
| 172 | 9.91831e-13 | 0.999994 |
| 174 | 2.17862e-13 | 1 |
| 176 | 1.1979e-12 | 1.00001 |
| 178 | 2.63781e-13 | 0.999994 |
| 180 | 2.9075e-13 | 1 |
| 182 | 3.20769e-13 | 1.00001 |
| 184 | 1.73696e-11 | 1 |
| 186 | 4.48253e-12 | 1 |
| 188 | 4.64069e-12 | 1 |
| 190 | 4.81763e-12 | 1 |
| 192 | 1.25367e-12 | 1 |
| 194 | 5.23273e-12 | 1 |
| 196 | 5.47347e-12 | 1 |
| 198 | 5.73788e-12 | 1 |
| 200 | 1.50708e-12 | 1 |
| 202 | 6.34604e-12 | 1 |
| 204 | 1.67336e-12 | 1 |
| 206 | 3.53617e-13 | 1 |
| 208 | 7.48558e-12 | 1 |
| 210 | 1.98386e-12 | 1 |
| 212 | 2.1062e-12 | 1 |
| 214 | 8.95701e-12 | 1 |
| 216 | 9.53519e-12 | 1 |
| 218 | 2.03264e-12 | 1 |
| 220 | 5.4225e-13 | 1 |
| 222 | 5.79257e-13 | 1 |
| 224 | 2.47768e-12 | 1 |
| 226 | 1.32591e-11 | 1 |
| 228 | 2.84079e-12 | 1 |
| 230 | 7.61435e-13 | 1 |
| 232 | 3.26875e-12 | 1 |
| 234 | 3.50897e-12 | 1 |
| 236 | 3.76737e-10 | 1 |
| 238 | 1.94221e-11 | 1 |
| 240 | 8.02354e-11 | 1 |
| 242 | 4.15e-12 | 1 |
| 244 | 4.29939e-12 | 1 |
| 246 | 1.78443e-11 | 1 |
| 248 | 2.31754e-11 | 1 |
| 250 | 9.6454e-13 | 1 |
| 252 | 1.00484e-12 | 1 |
| 254 | 5.24075e-12 | 1 |
| 256 | 5.47285e-12 | 1 |
| 258 | 2.86085e-11 | 1 |
| 260 | 2.39538e-11 | 1 |
| 262 | 3.13702e-11 | 1 |
| 264 | 6.57936e-12 | 1 |
| 266 | 6.90583e-12 | 1 |
| 268 | 2.90205e-11 | 1 |
| 270 | 7.62815e-12 | 1 |
| 272 | 8.02684e-12 | 1 |
| 274 | 1.69049e-12 | 1 |
| 276 | 8.90707e-12 | 1 |
| 278 | 4.69622e-13 | 1 |
| 280 | 9.91036e-12 | 1 |
| 282 | 2.61578e-12 | 1 |
| 284 | 5.52658e-13 | 1 |
| 286 | 1.1683e-11 | 1 |
| 288 | 1.23549e-11 | 1 |
| 290 | 1.30719e-11 | 1 |
| 292 | 2.76729e-12 | 1 |
| 294 | 3.66327e-12 | 1 |
| 296 | 3.10473e-12 | 1 |
| 298 | 1.64529e-11 | 1 |
| 300 | 3.48883e-12 | 1 |
| 302 | 3.7005e-12 | 1 |
| 304 | 3.92615e-12 | 1 |
| 306 | 2.08366e-11 | 1 |
| 308 | 4.42435e-12 | 1 |
| 310 | 1.17463e-12 | 1 |
| 312 | 4.99167e-12 | 1 |
| 314 | 5.30462e-12 | 1 |
| 316 | 5.63845e-12 | 1 |
| 318 | 2.39796e-11 | 1 |
| 320 | 6.37594e-12 | 1 |
| 322 | 1.69565e-12 | 1 |
| 324 | 5.77392e-12 | 1 |
| 326 | 1.53635e-12 | 1 |
| 328 | 1.63567e-12 | 1 |
| 330 | 8.70818e-12 | 1 |
| 332 | 9.27575e-12 | 1 |
| 334 | 7.90569e-12 | 1 |
| 336 | 1.0531e-11 | 1 |
| 338 | 2.24497e-12 | 1 |
| 340 | 9.58559e-12 | 1 |
| 342 | 1.27637e-11 | 1 |
| 344 | 1.36158e-11 | 1 |
| 346 | 2.90578e-12 | 1 |
| 348 | 3.10158e-12 | 1 |
| 350 | 4.13914e-12 | 1 |
| 352 | 0.189685 | 0.810313 |
| 354 | 0.190949 | 0.809054 |
| 356 | 0.192081 | 0.807921 |
| 358 | 0.193101 | 0.806899 |
| 360 | 0.194025 | 0.805972 |
| 362 | 0.19487 | 0.805124 |
| 364 | 0.195648 | 0.804354 |
| 366 | 0.196369 | 0.803634 |
| 368 | 0.197033 | 0.802962 |
| 370 | 0.197654 | 0.802351 |
| 372 | 0.198234 | 0.801761 |
| 374 | 0.198775 | 0.80123 |
| 376 | 0.199287 | 0.800718 |
| 378 | 0.199767 | 0.800238 |
| 380 | 0.200221 | 0.799776 |
| 382 | 0.200649 | 0.799346 |
| 384 | 0.201055 | 0.798945 |
| 386 | 0.20144 | 0.798562 |
| 388 | 0.201809 | 0.798196 |
| 390 | 0.202159 | 0.797846 |
| 392 | 0.202492 | 0.797513 |
| 394 | 0.202812 | 0.797183 |
| 396 | 0.203119 | 0.796881 |
| 398 | 0.20341 | 0.796595 |
| 400 | 0.203693 | 0.796312 |

V5-1 Equal

|  |  |  |
| --- | --- | --- |
| 2 | 0 | 2 |
| 4 | 0 | 4 |
| 6 | 0 | 6 |
| 8 | 0 | 8 |
| 10 | 0 | 10 |
| 12 | 0 | 12 |
| 14 | 0 | 14 |
| 16 | 0 | 16 |
| 18 | 0 | 18 |
| 20 | 0 | 20 |
| 22 | 0 | 22 |
| 24 | 0 | 24 |
| 26 | 0 | 26 |
| 28 | 0 | 28 |
| 30 | 0 | 30 |
| 32 | 0 | 32 |
| 34 | 0 | 34 |
| 36 | 0 | 36 |
| 38 | 0 | 38 |
| 40 | 0 | 40 |
| 42 | 0 | 42 |
| 44 | 0 | 44 |
| 46 | 0 | 46 |
| 48 | 0 | 48 |
| 50 | 0 | 50 |
| 52 | 0 | 52 |
| 54 | 0 | 54 |
| 56 | 0 | 56 |
| 58 | 0 | 58 |
| 60 | 0 | 60 |
| 62 | 0 | 62 |
| 64 | 0 | 64 |
| 66 | 0 | 66 |
| 68 | 0 | 68 |
| 70 | 0 | 70 |
| 72 | 0 | 72 |
| 74 | 0 | 74 |
| 76 | 0 | 76 |
| 78 | 0 | 78 |
| 80 | 0 | 80 |
| 82 | 0 | 41 |
| 84 | 0 | 42 |
| 86 | 0 | 43 |
| 88 | 0 | 44 |
| 90 | 0 | 45 |
| 92 | 0 | 46 |
| 94 | 0 | 47 |
| 96 | 0 | 48 |
| 98 | 0 | 49 |
| 100 | 0 | 50 |
| 102 | 0 | 51 |
| 104 | 0 | 52 |
| 106 | 0 | 53 |
| 108 | 0 | 54 |
| 110 | 0 | 55 |
| 112 | 0 | 56 |
| 114 | 0 | 57 |
| 116 | 0 | 58 |
| 118 | 0 | 59 |
| 120 | 0 | 60 |
| 122 | 0 | 61 |
| 124 | 0 | 62 |
| 126 | 0 | 63 |
| 128 | 0 | 64 |
| 130 | 0 | 65 |
| 132 | 0 | 44 |
| 134 | 0 | 44.667 |
| 136 | 0 | 45.333 |
| 138 | 0 | 46 |
| 140 | 0 | 46.667 |
| 142 | 0 | 47.333 |
| 144 | 0 | 48 |
| 146 | 0 | 48.667 |
| 148 | 0 | 49.333 |
| 150 | 0 | 50 |
| 152 | 0 | 50.667 |
| 154 | 0 | 51.333 |
| 156 | 0 | 52 |
| 158 | 0 | 52.667 |
| 160 | 0 | 53.333 |
| 162 | 0 | 54 |
| 164 | 0 | 54.667 |
| 166 | 0 | 55.333 |
| 168 | 0 | 56 |
| 170 | 0 | 56.667 |
| 172 | 0 | 57.333 |
| 174 | 0 | 58 |
| 176 | 0 | 58.667 |
| 178 | 0 | 59.333 |
| 180 | 0 | 60 |
| 182 | 0 | 60.667 |
| 184 | 0 | 46 |
| 186 | 0 | 46.5 |
| 188 | 0 | 47 |
| 190 | 0 | 47.5 |
| 192 | 0 | 48 |
| 194 | 0 | 48.5 |
| 196 | 0 | 49 |
| 198 | 0 | 49.5 |
| 200 | 0 | 50 |
| 202 | 0 | 50.5 |
| 204 | 0 | 51 |
| 206 | 0 | 51.5 |
| 208 | 0 | 52 |
| 210 | 0 | 52.5 |
| 212 | 0 | 53 |
| 214 | 0 | 53.5 |
| 216 | 0 | 54 |
| 218 | 0 | 54.5 |
| 220 | 0 | 55 |
| 222 | 0 | 55.5 |
| 224 | 0 | 56 |
| 226 | 0 | 56.5 |
| 228 | 0 | 57 |
| 230 | 0 | 57.5 |
| 232 | 0 | 58 |
| 234 | 0 | 58.5 |
| 236 | 0 | 47.2 |
| 238 | 0 | 47.6 |
| 240 | 0 | 48 |
| 242 | 0 | 48.4 |
| 244 | 0 | 48.8 |
| 246 | 0 | 49.2 |
| 248 | 0 | 49.6 |
| 250 | 0 | 50 |
| 252 | 0 | 50.4 |
| 254 | 0 | 50.8 |
| 256 | 0 | 51.2 |
| 258 | 0 | 51.6 |
| 260 | 0 | 52 |
| 262 | 0 | 52.4 |
| 264 | 0 | 52.8 |
| 266 | 0 | 53.2 |
| 268 | 0 | 53.6 |
| 270 | 0 | 54 |
| 272 | 0 | 54.4 |
| 274 | 0 | 54.8 |
| 276 | 0 | 55.2 |
| 278 | 0 | 55.6 |
| 280 | 0 | 56 |
| 282 | 0 | 56.4 |
| 284 | 0 | 56.8 |
| 286 | 0 | 57.2 |
| 288 | 0 | 57.6 |
| 290 | 0 | 58 |
| 292 | 0 | 58.4 |
| 294 | 0 | 58.8 |
| 296 | 0 | 59.2 |
| 298 | 0 | 59.6 |
| 300 | 0 | 60 |
| 302 | 0 | 60.4 |
| 304 | 0 | 60.8 |
| 306 | 0 | 61.2 |
| 308 | 0 | 61.6 |
| 310 | 0 | 62 |
| 312 | 0 | 62.4 |
| 314 | 0 | 62.8 |
| 316 | 0 | 63.2 |
| 318 | 0 | 63.6 |
| 320 | 0 | 64 |
| 322 | 0 | 64.4 |
| 324 | 0 | 64.8 |
| 326 | 0 | 65.2 |
| 328 | 0 | 65.6 |
| 330 | 0 | 66 |
| 332 | 0 | 66.4 |
| 334 | 0 | 66.8 |
| 336 | 0 | 67.2 |
| 338 | 0 | 67.6 |
| 340 | 0 | 68 |
| 342 | 0 | 68.4 |
| 344 | 0 | 68.8 |
| 346 | 0 | 69.2 |
| 348 | 0 | 69.6 |
| 350 | 0 | 70 |
| 352 | 66.769 | 57.046 |
| 354 | 67.596 | 57.281 |
| 356 | 68.381 | 57.524 |
| 358 | 69.13 | 57.774 |
| 360 | 69.849 | 58.03 |
| 362 | 70.543 | 58.291 |
| 364 | 71.216 | 58.557 |
| 366 | 71.871 | 58.826 |
| 368 | 72.508 | 59.098 |
| 370 | 73.132 | 59.374 |
| 372 | 73.743 | 59.651 |
| 374 | 74.342 | 59.932 |
| 376 | 74.932 | 60.214 |
| 378 | 75.512 | 60.498 |
| 380 | 76.084 | 60.783 |
| 382 | 76.648 | 61.07 |
| 384 | 77.205 | 61.359 |
| 386 | 77.756 | 61.649 |
| 388 | 78.302 | 61.94 |
| 390 | 78.842 | 62.232 |
| 392 | 79.377 | 62.525 |
| 394 | 79.908 | 62.818 |
| 396 | 80.435 | 63.113 |
| 398 | 80.957 | 63.409 |
| 400 | 81.477 | 63.705 |

Figure S8 | Calculation results of model version 5-1.

(a) Fitness functions of offspring. Upper bound of red line is 30 and that of blue line is 10. For sake of brevity, computational results from the brood which is composed of equal ratio is shown: That is, 5 offspring with upper bound of 10, and the other 5 offspring with upper bound of 30. For all of the figures of model version 5, the red line stays the same and blue line brood is composed of 5 offspring with variable number of upper bounds in the subsequent models of version 5.

all graphs of model version 5 is based on equal brood

(b) The Optimized number of cared offspring of each brood.

(c) The optimized ratio of total care given to each brood.

(d) The optimized per capita investment to cared offspring.

**Model Version 5-2**

V5-2 Functions

|  |  |  |
| --- | --- | --- |
| 0 | 3.40006e-16 | 5.37764e-16 |
| 1 | 0.00243763 | 0.00564835 |
| 2 | 0.005172 | 0.0123587 |
| 3 | 0.00823911 | 0.02033 |
| 4 | 0.0116794 | 0.0297984 |
| 5 | 0.015538 | 0.0410439 |
| 6 | 0.0198656 | 0.0543982 |
| 7 | 0.0247191 | 0.0702543 |
| 8 | 0.030162 | 0.0890775 |
| 9 | 0.0362654 | 0.111418 |
| 10 | 0.0431092 | 0.137927 |
| 11 | 0.0507825 | 0.169371 |
| 12 | 0.059385 | 0.206657 |
| 13 | 0.0690284 | 0.25085 |
| 14 | 0.0798373 | 0.303204 |
| 15 | 0.0919512 | 0.365187 |
| 16 | 0.105526 | 0.438518 |
| 17 | 0.120734 | 0.525202 |
| 18 | 0.137771 | 0.627567 |
| 19 | 0.156852 | 0.748307 |
| 20 | 0.178218 | 0.890523 |
| 21 | 0.202136 | 1.05776 |
| 22 | 0.228903 | 1.25404 |
| 23 | 0.258851 | 1.48388 |
| 24 | 0.292345 | 1.75231 |
| 25 | 0.329791 | 2.06484 |
| 26 | 0.371637 | 2.42741 |
| 27 | 0.418378 | 2.84627 |
| 28 | 0.470557 | 3.32783 |
| 29 | 0.528774 | 3.87841 |
| 30 | 0.593682 | 4.50392 |
| 31 | 0.665996 | 5.20945 |
| 32 | 0.746496 | 5.99882 |
| 33 | 0.836023 | 6.874 |
| 34 | 0.935489 | 7.83459 |
| 35 | 1.04587 | 8.87735 |
| 36 | 1.16821 | 9.99583 |
| 37 | 1.3036 | 11.1802 |
| 38 | 1.45322 | 12.4175 |
| 39 | 1.61827 | 13.6917 |
| 40 | 1.8 | 14.985 |
| 41 | 1.99967 | 16.2783 |
| 42 | 2.21855 | 17.5525 |
| 43 | 2.45789 | 18.7898 |
| 44 | 2.71887 | 19.9742 |
| 45 | 3.00261 | 21.0927 |
| 46 | 3.31008 | 22.1354 |
| 47 | 3.64208 | 23.096 |
| 48 | 3.99922 | 23.9712 |
| 49 | 4.38181 | 24.7605 |
| 50 | 4.78987 | 25.4661 |
| 51 | 5.22306 | 26.0916 |
| 52 | 5.68064 | 26.6422 |
| 53 | 6.16145 | 27.1237 |
| 54 | 6.66389 | 27.5426 |
| 55 | 7.1859 | 27.9052 |
| 56 | 7.725 | 28.2177 |
| 57 | 8.27832 | 28.4861 |
| 58 | 8.84262 | 28.716 |
| 59 | 9.41441 | 28.9122 |
| 60 | 9.99 | 29.0795 |
| 61 | 10.5656 | 29.2217 |
| 62 | 11.1374 | 29.3424 |
| 63 | 11.7017 | 29.4448 |
| 64 | 12.255 | 29.5315 |
| 65 | 12.7941 | 29.6048 |
| 66 | 13.3161 | 29.6668 |
| 67 | 13.8185 | 29.7191 |
| 68 | 14.2994 | 29.7633 |
| 69 | 14.7569 | 29.8006 |
| 70 | 15.1901 | 29.8321 |
| 71 | 15.5982 | 29.8586 |
| 72 | 15.9808 | 29.8809 |
| 73 | 16.3379 | 29.8997 |
| 74 | 16.6699 | 29.9156 |
| 75 | 16.9774 | 29.929 |
| 76 | 17.2611 | 29.9402 |
| 77 | 17.5221 | 29.9497 |
| 78 | 17.7614 | 29.9576 |
| 79 | 17.9803 | 29.9644 |
| 80 | 18.18 | 29.97 |
| 81 | 18.3617 | 29.9748 |
| 82 | 18.5268 | 29.9788 |
| 83 | 18.6764 | 29.9821 |
| 84 | 18.8118 | 29.985 |
| 85 | 18.9341 | 29.9873 |
| 86 | 19.0445 | 29.9893 |
| 87 | 19.144 | 29.991 |
| 88 | 19.2335 | 29.9925 |
| 89 | 19.314 | 29.9937 |
| 90 | 19.3863 | 29.9947 |
| 91 | 19.4512 | 29.9955 |
| 92 | 19.5094 | 29.9962 |
| 93 | 19.5616 | 29.9968 |
| 94 | 19.6084 | 29.9973 |
| 95 | 19.6502 | 29.9977 |
| 96 | 19.6877 | 29.9981 |
| 97 | 19.7211 | 29.9984 |
| 98 | 19.7511 | 29.9987 |
| 99 | 19.7779 | 29.9989 |
| 100 | 19.8018 | 29.9991 |
| 101 | 19.8231 | 29.9992 |
| 102 | 19.8422 | 29.9993 |
| 103 | 19.8593 | 29.9994 |
| 104 | 19.8745 | 29.9995 |
| 105 | 19.888 | 29.9996 |
| 106 | 19.9002 | 29.9997 |
| 107 | 19.911 | 29.9997 |
| 108 | 19.9206 | 29.9998 |
| 109 | 19.9292 | 29.9998 |
| 110 | 19.9369 | 29.9998 |
| 111 | 19.9437 | 29.9999 |
| 112 | 19.9498 | 29.9999 |
| 113 | 19.9553 | 29.9999 |
| 114 | 19.9601 | 29.9999 |
| 115 | 19.9645 | 29.9999 |
| 116 | 19.9683 | 29.9999 |
| 117 | 19.9718 | 29.9999 |
| 118 | 19.9748 | 30 |
| 119 | 19.9776 | 30 |
| 120 | 19.98 | 30 |
| 121 | 19.9822 | 30 |
| 122 | 19.9841 | 30 |
| 123 | 19.9858 | 30 |
| 124 | 19.9874 | 30 |
| 125 | 19.9887 | 30 |
| 126 | 19.99 | 30 |
| 127 | 19.9911 | 30 |
| 128 | 19.992 | 30 |
| 129 | 19.9929 | 30 |
| 130 | 19.9937 | 30 |
| 131 | 19.9944 | 30 |
| 132 | 19.995 | 30 |
| 133 | 19.9955 | 30 |
| 134 | 19.996 | 30 |
| 135 | 19.9964 | 30 |
| 136 | 19.9968 | 30 |
| 137 | 19.9972 | 30 |
| 138 | 19.9975 | 30 |
| 139 | 19.9978 | 30 |
| 140 | 19.998 | 30 |
| 141 | 19.9982 | 30 |
| 142 | 19.9984 | 30 |
| 143 | 19.9986 | 30 |
| 144 | 19.9987 | 30 |
| 145 | 19.9989 | 30 |
| 146 | 19.999 | 30 |
| 147 | 19.9991 | 30 |
| 148 | 19.9992 | 30 |
| 149 | 19.9993 | 30 |
| 150 | 19.9994 | 30 |
| 151 | 19.9994 | 30 |
| 152 | 19.9995 | 30 |
| 153 | 19.9996 | 30 |
| 154 | 19.9996 | 30 |
| 155 | 19.9996 | 30 |
| 156 | 19.9997 | 30 |
| 157 | 19.9997 | 30 |
| 158 | 19.9997 | 30 |
| 159 | 19.9998 | 30 |
| 160 | 19.9998 | 30 |
| 161 | 19.9998 | 30 |
| 162 | 19.9998 | 30 |
| 163 | 19.9999 | 30 |
| 164 | 19.9999 | 30 |
| 165 | 19.9999 | 30 |
| 166 | 19.9999 | 30 |
| 167 | 19.9999 | 30 |
| 168 | 19.9999 | 30 |
| 169 | 19.9999 | 30 |
| 170 | 19.9999 | 30 |
| 171 | 19.9999 | 30 |
| 172 | 19.9999 | 30 |
| 173 | 20 | 30 |
| 174 | 20 | 30 |
| 175 | 20 | 30 |
| 176 | 20 | 30 |
| 177 | 20 | 30 |
| 178 | 20 | 30 |
| 179 | 20 | 30 |
| 180 | 20 | 30 |
| 181 | 20 | 30 |
| 182 | 20 | 30 |
| 183 | 20 | 30 |
| 184 | 20 | 30 |
| 185 | 20 | 30 |
| 186 | 20 | 30 |
| 187 | 20 | 30 |
| 188 | 20 | 30 |
| 189 | 20 | 30 |
| 190 | 20 | 30 |
| 191 | 20 | 30 |
| 192 | 20 | 30 |
| 193 | 20 | 30 |
| 194 | 20 | 30 |
| 195 | 20 | 30 |
| 196 | 20 | 30 |
| 197 | 20 | 30 |
| 198 | 20 | 30 |
| 199 | 20 | 30 |

V5-2 Equal

|  |  |  |
| --- | --- | --- |
| 2 | 0 | 1 |
| 4 | 0 | 1 |
| 6 | 0 | 1 |
| 8 | 0 | 1 |
| 10 | 0 | 1 |
| 12 | 0 | 1 |
| 14 | 0 | 1 |
| 16 | 0 | 1 |
| 18 | 0 | 1 |
| 20 | 0 | 1 |
| 22 | 0 | 1 |
| 24 | 0 | 1 |
| 26 | 0 | 1 |
| 28 | 0 | 1 |
| 30 | 0 | 1 |
| 32 | 0 | 1 |
| 34 | 0 | 1 |
| 36 | 0 | 1 |
| 38 | 0 | 1 |
| 40 | 0 | 1 |
| 42 | 0 | 1 |
| 44 | 0 | 1 |
| 46 | 0 | 1 |
| 48 | 0 | 1 |
| 50 | 0 | 1 |
| 52 | 0 | 1 |
| 54 | 0 | 1 |
| 56 | 0 | 1 |
| 58 | 0 | 1 |
| 60 | 0 | 1 |
| 62 | 0 | 1 |
| 64 | 0 | 1 |
| 66 | 0 | 1 |
| 68 | 0 | 1 |
| 70 | 0 | 1 |
| 72 | 0 | 1 |
| 74 | 0 | 1 |
| 76 | 0 | 1 |
| 78 | 0 | 1 |
| 80 | 0 | 1 |
| 82 | 0 | 2 |
| 84 | 0 | 2 |
| 86 | 0 | 2 |
| 88 | 0 | 2 |
| 90 | 0 | 2 |
| 92 | 0 | 2 |
| 94 | 0 | 2 |
| 96 | 0 | 2 |
| 98 | 0 | 2 |
| 100 | 0 | 2 |
| 102 | 0 | 2 |
| 104 | 0 | 2 |
| 106 | 0 | 2 |
| 108 | 0 | 2 |
| 110 | 0 | 2 |
| 112 | 0 | 2 |
| 114 | 0 | 2 |
| 116 | 0 | 2 |
| 118 | 0 | 2 |
| 120 | 0 | 2 |
| 122 | 0 | 2 |
| 124 | 0 | 2 |
| 126 | 0 | 2 |
| 128 | 0 | 2 |
| 130 | 0 | 2 |
| 132 | 0 | 3 |
| 134 | 0 | 3 |
| 136 | 0 | 3 |
| 138 | 0 | 3 |
| 140 | 0 | 3 |
| 142 | 0 | 3 |
| 144 | 0 | 3 |
| 146 | 0 | 3 |
| 148 | 0 | 3 |
| 150 | 0 | 3 |
| 152 | 0 | 3 |
| 154 | 0 | 3 |
| 156 | 0 | 3 |
| 158 | 0 | 3 |
| 160 | 0 | 3 |
| 162 | 0 | 3 |
| 164 | 0 | 3 |
| 166 | 0 | 3 |
| 168 | 0 | 3 |
| 170 | 0 | 3 |
| 172 | 0 | 3 |
| 174 | 0 | 3 |
| 176 | 0 | 3 |
| 178 | 0 | 3 |
| 180 | 0 | 3 |
| 182 | 0 | 3 |
| 184 | 0 | 4 |
| 186 | 0 | 4 |
| 188 | 0 | 4 |
| 190 | 0 | 4 |
| 192 | 0 | 4 |
| 194 | 0 | 4 |
| 196 | 0 | 4 |
| 198 | 0 | 4 |
| 200 | 0 | 4 |
| 202 | 0 | 4 |
| 204 | 0 | 4 |
| 206 | 0 | 4 |
| 208 | 0 | 4 |
| 210 | 0 | 4 |
| 212 | 0 | 4 |
| 214 | 0 | 4 |
| 216 | 0 | 4 |
| 218 | 0 | 4 |
| 220 | 0 | 4 |
| 222 | 0 | 4 |
| 224 | 0 | 4 |
| 226 | 0 | 4 |
| 228 | 0 | 4 |
| 230 | 0 | 4 |
| 232 | 0 | 4 |
| 234 | 0 | 4 |
| 236 | 0 | 5 |
| 238 | 0 | 5 |
| 240 | 0 | 5 |
| 242 | 0 | 5 |
| 244 | 0 | 5 |
| 246 | 0 | 5 |
| 248 | 0 | 5 |
| 250 | 0 | 5 |
| 252 | 0 | 5 |
| 254 | 0 | 5 |
| 256 | 0 | 5 |
| 258 | 0 | 5 |
| 260 | 0 | 5 |
| 262 | 0 | 5 |
| 264 | 0 | 5 |
| 266 | 0 | 5 |
| 268 | 0 | 5 |
| 270 | 0 | 5 |
| 272 | 0 | 5 |
| 274 | 0 | 5 |
| 276 | 0 | 5 |
| 278 | 0 | 5 |
| 280 | 0 | 5 |
| 282 | 0 | 5 |
| 284 | 0 | 5 |
| 286 | 0 | 5 |
| 288 | 0 | 5 |
| 290 | 0 | 5 |
| 292 | 0 | 5 |
| 294 | 0 | 5 |
| 296 | 0 | 5 |
| 298 | 0 | 5 |
| 300 | 0 | 5 |
| 302 | 0 | 5 |
| 304 | 0 | 5 |
| 306 | 0 | 5 |
| 308 | 0 | 5 |
| 310 | 0 | 5 |
| 312 | 0 | 5 |
| 314 | 0 | 5 |
| 316 | 0 | 5 |
| 318 | 0 | 5 |
| 320 | 0 | 5 |
| 322 | 0 | 5 |
| 324 | 0 | 5 |
| 326 | 0 | 5 |
| 328 | 0 | 5 |
| 330 | 1 | 5 |
| 332 | 1 | 5 |
| 334 | 1 | 5 |
| 336 | 1 | 5 |
| 338 | 1 | 5 |
| 340 | 1 | 5 |
| 342 | 1 | 5 |
| 344 | 1 | 5 |
| 346 | 1 | 5 |
| 348 | 1 | 5 |
| 350 | 1 | 5 |
| 352 | 1 | 5 |
| 354 | 1 | 5 |
| 356 | 1 | 5 |
| 358 | 1 | 5 |
| 360 | 1 | 5 |
| 362 | 1 | 5 |
| 364 | 1 | 5 |
| 366 | 1 | 5 |
| 368 | 1 | 5 |
| 370 | 1 | 5 |
| 372 | 1 | 5 |
| 374 | 1 | 5 |
| 376 | 1 | 5 |
| 378 | 1 | 5 |
| 380 | 1 | 5 |
| 382 | 1 | 5 |
| 384 | 1 | 5 |
| 386 | 1 | 5 |
| 388 | 1 | 5 |
| 390 | 1 | 5 |
| 392 | 1 | 5 |
| 394 | 1 | 5 |
| 396 | 1 | 5 |
| 398 | 1 | 5 |
| 400 | 1 | 5 |

v9-2 Equal

|  |  |  |
| --- | --- | --- |
| 2 | 3.406e-09 | 1 |
| 4 | 5.4285e-09 | 1 |
| 6 | 2.38952e-09 | 1 |
| 8 | 3.02931e-10 | 1 |
| 10 | 6.661e-10 | 1 |
| 12 | 3.85846e-10 | 1 |
| 14 | 2.31932e-10 | 1 |
| 16 | 1.79227e-10 | 1 |
| 18 | 1.1335e-10 | 1 |
| 20 | 7.31275e-11 | 1 |
| 22 | 3.84614e-11 | 1 |
| 24 | 1.28883e-10 | 1 |
| 26 | 2.20577e-11 | 1 |
| 28 | 3.09482e-10 | 1 |
| 30 | 1.11792e-11 | 1 |
| 32 | 1.67344e-10 | 1 |
| 34 | 1.30677e-10 | 1 |
| 36 | 2.14486e-11 | 1 |
| 38 | 2.32724e-11 | 1 |
| 40 | 4.29225e-12 | 1 |
| 42 | 2.1056e-11 | 1 |
| 44 | 2.19364e-11 | 1 |
| 46 | 2.41467e-11 | 1 |
| 48 | 2.78906e-11 | 1 |
| 50 | 6.7075e-12 | 1 |
| 52 | 8.3324e-12 | 1 |
| 54 | 1.06204e-11 | 1 |
| 56 | 5.52348e-13 | 1 |
| 58 | 1.82353e-11 | 1 |
| 60 | 2.43767e-11 | 1 |
| 62 | 2.63266e-11 | 1 |
| 64 | 3.58453e-11 | 1 |
| 66 | 4.91977e-11 | 1 |
| 68 | 1.70228e-11 | 1 |
| 70 | 1.90314e-11 | 1 |
| 72 | 2.69271e-11 | 1 |
| 74 | 9.68716e-12 | 1 |
| 76 | 1.14283e-11 | 1 |
| 78 | 1.7466e-11 | 1 |
| 80 | 1.32913e-11 | 1 |
| 82 | 7.59451e-14 | 1 |
| 84 | 7.58095e-14 | 1 |
| 86 | 7.68198e-14 | 1 |
| 88 | 7.89773e-14 | 1 |
| 90 | 8.23278e-14 | 1 |
| 92 | 8.69402e-14 | 1 |
| 94 | 1.16154e-13 | 1 |
| 96 | 1.00422e-13 | 1 |
| 98 | 1.09628e-13 | 1 |
| 100 | 1.5095e-13 | 1 |
| 102 | 6.70588e-13 | 1 |
| 104 | 1.50024e-13 | 1 |
| 106 | 1.68901e-13 | 1 |
| 108 | 1.91231e-13 | 1 |
| 110 | 2.17595e-13 | 1 |
| 112 | 2.48683e-13 | 1 |
| 114 | 2.85316e-13 | 1 |
| 116 | 3.28466e-13 | 1 |
| 118 | 3.79292e-13 | 1 |
| 120 | 4.39208e-13 | 1 |
| 122 | 5.09795e-13 | 1 |
| 124 | 5.93145e-13 | 1 |
| 126 | 5.53254e-13 | 1 |
| 128 | 8.08047e-13 | 1 |
| 130 | 9.46038e-13 | 1 |
| 132 | 2.63265e-13 | 1 |
| 134 | 2.70254e-13 | 1.00001 |
| 136 | 1.39537e-12 | 0.999993 |
| 138 | 2.89804e-13 | 1 |
| 140 | 3.02564e-13 | 1.00001 |
| 142 | 1.58746e-12 | 0.999993 |
| 144 | 3.34743e-13 | 1 |
| 146 | 3.54521e-13 | 1.00001 |
| 148 | 3.77027e-13 | 0.999993 |
| 150 | 4.02533e-13 | 1 |
| 152 | 4.31316e-13 | 1.00001 |
| 154 | 4.63701e-13 | 0.999994 |
| 156 | 4.00064e-13 | 1 |
| 158 | 4.32658e-13 | 1.00001 |
| 160 | 4.69156e-13 | 0.999994 |
| 162 | 5.09938e-13 | 1 |
| 164 | 5.55518e-13 | 1.00001 |
| 166 | 7.57982e-13 | 0.999994 |
| 168 | 8.28958e-13 | 1 |
| 170 | 9.08118e-13 | 1.00001 |
| 172 | 7.97122e-13 | 0.999994 |
| 174 | 1.09489e-12 | 1 |
| 176 | 1.20472e-12 | 1.00001 |
| 178 | 1.32725e-12 | 0.999994 |
| 180 | 1.46397e-12 | 1 |
| 182 | 1.61654e-12 | 1.00001 |
| 184 | 2.17353e-11 | 1 |
| 186 | 4.48763e-12 | 1 |
| 188 | 4.64628e-12 | 1 |
| 190 | 4.82368e-12 | 1 |
| 192 | 1.25529e-12 | 1 |
| 194 | 6.55e-12 | 1 |
| 196 | 5.48138e-12 | 1 |
| 198 | 5.74672e-12 | 1 |
| 200 | 1.50953e-12 | 1 |
| 202 | 1.27141e-12 | 1 |
| 204 | 3.35294e-13 | 1 |
| 206 | 1.77158e-12 | 1 |
| 208 | 1.87531e-12 | 1 |
| 210 | 1.98829e-12 | 1 |
| 212 | 8.44505e-12 | 1 |
| 214 | 4.48995e-13 | 1 |
| 216 | 4.78079e-13 | 1 |
| 218 | 5.09679e-13 | 1 |
| 220 | 5.44e-13 | 1 |
| 222 | 5.81261e-13 | 1 |
| 224 | 6.21719e-13 | 1 |
| 226 | 2.66261e-12 | 1 |
| 228 | 7.13289e-13 | 1 |
| 230 | 3.06e-12 | 1 |
| 232 | 8.21164e-13 | 1 |
| 234 | 8.82094e-13 | 1 |
| 236 | 3.77182e-12 | 1 |
| 238 | 1.94462e-11 | 1 |
| 240 | 4.01688e-12 | 1 |
| 242 | 4.15558e-12 | 1 |
| 244 | 4.30553e-12 | 1 |
| 246 | 8.93516e-13 | 1 |
| 248 | 4.64234e-12 | 1 |
| 250 | 1.20762e-12 | 1 |
| 252 | 2.51647e-11 | 1 |
| 254 | 1.3125e-12 | 1 |
| 256 | 1.09664e-12 | 1 |
| 258 | 5.73314e-12 | 1 |
| 260 | 6.00096e-12 | 1 |
| 262 | 6.28779e-12 | 1 |
| 264 | 6.5947e-12 | 1 |
| 266 | 6.92293e-12 | 1 |
| 268 | 1.45481e-12 | 1 |
| 270 | 1.52985e-12 | 1 |
| 272 | 8.05037e-12 | 1 |
| 274 | 8.47883e-12 | 1 |
| 276 | 4.4683e-13 | 1 |
| 278 | 1.88507e-12 | 1 |
| 280 | 1.98946e-12 | 1 |
| 282 | 2.62624e-12 | 1 |
| 284 | 1.11e-11 | 1 |
| 286 | 5.86783e-11 | 1 |
| 288 | 2.48281e-12 | 1 |
| 290 | 6.56931e-13 | 1 |
| 292 | 1.39121e-11 | 1 |
| 294 | 2.94762e-12 | 1 |
| 296 | 3.12399e-12 | 1 |
| 298 | 3.31242e-12 | 1 |
| 300 | 3.5135e-12 | 1 |
| 302 | 3.72831e-12 | 1 |
| 304 | 7.91579e-13 | 1 |
| 306 | 1.05077e-12 | 1 |
| 308 | 1.11627e-12 | 1 |
| 310 | 1.18627e-12 | 1 |
| 312 | 5.04439e-12 | 1 |
| 314 | 1.07288e-12 | 1 |
| 316 | 5.70665e-12 | 1 |
| 318 | 1.21454e-12 | 1 |
| 320 | 1.29286e-12 | 1 |
| 322 | 1.37669e-12 | 1 |
| 324 | 7.33225e-12 | 1 |
| 326 | 1.56258e-12 | 1 |
| 328 | 8.32774e-12 | 1 |
| 330 | 0.204509 | 0.795485 |
| 332 | 0.205452 | 0.794548 |
| 334 | 0.206296 | 0.793698 |
| 336 | 0.207054 | 0.792946 |
| 338 | 0.207743 | 0.792263 |
| 340 | 0.208371 | 0.791632 |
| 342 | 0.208947 | 0.791053 |
| 344 | 0.209477 | 0.790523 |
| 346 | 0.209968 | 0.790029 |
| 348 | 0.210422 | 0.789583 |
| 350 | 0.210843 | 0.789157 |
| 352 | 0.211239 | 0.788764 |
| 354 | 0.211607 | 0.78839 |
| 356 | 0.211952 | 0.788048 |
| 358 | 0.212277 | 0.787723 |
| 360 | 0.212581 | 0.787417 |
| 362 | 0.212867 | 0.787127 |
| 364 | 0.21314 | 0.786854 |
| 366 | 0.213396 | 0.786598 |
| 368 | 0.213641 | 0.786359 |
| 370 | 0.213873 | 0.786122 |
| 372 | 0.214094 | 0.785901 |
| 374 | 0.214302 | 0.785695 |
| 376 | 0.214503 | 0.785492 |
| 378 | 0.214693 | 0.785304 |
| 380 | 0.214876 | 0.785118 |
| 382 | 0.21505 | 0.784948 |
| 384 | 0.215216 | 0.784779 |
| 386 | 0.215378 | 0.784624 |
| 388 | 0.215531 | 0.784472 |
| 390 | 0.215679 | 0.784321 |
| 392 | 0.215821 | 0.784184 |
| 394 | 0.215959 | 0.784036 |
| 396 | 0.216091 | 0.783914 |
| 398 | 0.216219 | 0.783781 |
| 400 | 0.216343 | 0.783662 |

V5-2 Equal

|  |  |  |
| --- | --- | --- |
| 2 | 0 | 2 |
| 4 | 0 | 4 |
| 6 | 0 | 6 |
| 8 | 0 | 8 |
| 10 | 0 | 10 |
| 12 | 0 | 12 |
| 14 | 0 | 14 |
| 16 | 0 | 16 |
| 18 | 0 | 18 |
| 20 | 0 | 20 |
| 22 | 0 | 22 |
| 24 | 0 | 24 |
| 26 | 0 | 26 |
| 28 | 0 | 28 |
| 30 | 0 | 30 |
| 32 | 0 | 32 |
| 34 | 0 | 34 |
| 36 | 0 | 36 |
| 38 | 0 | 38 |
| 40 | 0 | 40 |
| 42 | 0 | 42 |
| 44 | 0 | 44 |
| 46 | 0 | 46 |
| 48 | 0 | 48 |
| 50 | 0 | 50 |
| 52 | 0 | 52 |
| 54 | 0 | 54 |
| 56 | 0 | 56 |
| 58 | 0 | 58 |
| 60 | 0 | 60 |
| 62 | 0 | 62 |
| 64 | 0 | 64 |
| 66 | 0 | 66 |
| 68 | 0 | 68 |
| 70 | 0 | 70 |
| 72 | 0 | 72 |
| 74 | 0 | 74 |
| 76 | 0 | 76 |
| 78 | 0 | 78 |
| 80 | 0 | 80 |
| 82 | 0 | 41 |
| 84 | 0 | 42 |
| 86 | 0 | 43 |
| 88 | 0 | 44 |
| 90 | 0 | 45 |
| 92 | 0 | 46 |
| 94 | 0 | 47 |
| 96 | 0 | 48 |
| 98 | 0 | 49 |
| 100 | 0 | 50 |
| 102 | 0 | 51 |
| 104 | 0 | 52 |
| 106 | 0 | 53 |
| 108 | 0 | 54 |
| 110 | 0 | 55 |
| 112 | 0 | 56 |
| 114 | 0 | 57 |
| 116 | 0 | 58 |
| 118 | 0 | 59 |
| 120 | 0 | 60 |
| 122 | 0 | 61 |
| 124 | 0 | 62 |
| 126 | 0 | 63 |
| 128 | 0 | 64 |
| 130 | 0 | 65 |
| 132 | 0 | 44 |
| 134 | 0 | 44.667 |
| 136 | 0 | 45.333 |
| 138 | 0 | 46 |
| 140 | 0 | 46.667 |
| 142 | 0 | 47.333 |
| 144 | 0 | 48 |
| 146 | 0 | 48.667 |
| 148 | 0 | 49.333 |
| 150 | 0 | 50 |
| 152 | 0 | 50.667 |
| 154 | 0 | 51.333 |
| 156 | 0 | 52 |
| 158 | 0 | 52.667 |
| 160 | 0 | 53.333 |
| 162 | 0 | 54 |
| 164 | 0 | 54.667 |
| 166 | 0 | 55.333 |
| 168 | 0 | 56 |
| 170 | 0 | 56.667 |
| 172 | 0 | 57.333 |
| 174 | 0 | 58 |
| 176 | 0 | 58.667 |
| 178 | 0 | 59.333 |
| 180 | 0 | 60 |
| 182 | 0 | 60.667 |
| 184 | 0 | 46 |
| 186 | 0 | 46.5 |
| 188 | 0 | 47 |
| 190 | 0 | 47.5 |
| 192 | 0 | 48 |
| 194 | 0 | 48.5 |
| 196 | 0 | 49 |
| 198 | 0 | 49.5 |
| 200 | 0 | 50 |
| 202 | 0 | 50.5 |
| 204 | 0 | 51 |
| 206 | 0 | 51.5 |
| 208 | 0 | 52 |
| 210 | 0 | 52.5 |
| 212 | 0 | 53 |
| 214 | 0 | 53.5 |
| 216 | 0 | 54 |
| 218 | 0 | 54.5 |
| 220 | 0 | 55 |
| 222 | 0 | 55.5 |
| 224 | 0 | 56 |
| 226 | 0 | 56.5 |
| 228 | 0 | 57 |
| 230 | 0 | 57.5 |
| 232 | 0 | 58 |
| 234 | 0 | 58.5 |
| 236 | 0 | 47.2 |
| 238 | 0 | 47.6 |
| 240 | 0 | 48 |
| 242 | 0 | 48.4 |
| 244 | 0 | 48.8 |
| 246 | 0 | 49.2 |
| 248 | 0 | 49.6 |
| 250 | 0 | 50 |
| 252 | 0 | 50.4 |
| 254 | 0 | 50.8 |
| 256 | 0 | 51.2 |
| 258 | 0 | 51.6 |
| 260 | 0 | 52 |
| 262 | 0 | 52.4 |
| 264 | 0 | 52.8 |
| 266 | 0 | 53.2 |
| 268 | 0 | 53.6 |
| 270 | 0 | 54 |
| 272 | 0 | 54.4 |
| 274 | 0 | 54.8 |
| 276 | 0 | 55.2 |
| 278 | 0 | 55.6 |
| 280 | 0 | 56 |
| 282 | 0 | 56.4 |
| 284 | 0 | 56.8 |
| 286 | 0 | 57.2 |
| 288 | 0 | 57.6 |
| 290 | 0 | 58 |
| 292 | 0 | 58.4 |
| 294 | 0 | 58.8 |
| 296 | 0 | 59.2 |
| 298 | 0 | 59.6 |
| 300 | 0 | 60 |
| 302 | 0 | 60.4 |
| 304 | 0 | 60.8 |
| 306 | 0 | 61.2 |
| 308 | 0 | 61.6 |
| 310 | 0 | 62 |
| 312 | 0 | 62.4 |
| 314 | 0 | 62.8 |
| 316 | 0 | 63.2 |
| 318 | 0 | 63.6 |
| 320 | 0 | 64 |
| 322 | 0 | 64.4 |
| 324 | 0 | 64.8 |
| 326 | 0 | 65.2 |
| 328 | 0 | 65.6 |
| 330 | 67.488 | 52.502 |
| 332 | 68.21 | 52.758 |
| 334 | 68.903 | 53.019 |
| 336 | 69.57 | 53.286 |
| 338 | 70.217 | 53.557 |
| 340 | 70.846 | 53.831 |
| 342 | 71.46 | 54.108 |
| 344 | 72.06 | 54.388 |
| 346 | 72.649 | 54.67 |
| 348 | 73.227 | 54.955 |
| 350 | 73.795 | 55.241 |
| 352 | 74.356 | 55.529 |
| 354 | 74.909 | 55.818 |
| 356 | 75.455 | 56.109 |
| 358 | 75.995 | 56.401 |
| 360 | 76.529 | 56.694 |
| 362 | 77.058 | 56.988 |
| 364 | 77.583 | 57.283 |
| 366 | 78.103 | 57.579 |
| 368 | 78.62 | 57.876 |
| 370 | 79.133 | 58.173 |
| 372 | 79.643 | 58.471 |
| 374 | 80.149 | 58.77 |
| 376 | 80.653 | 59.069 |
| 378 | 81.154 | 59.369 |
| 380 | 81.653 | 59.669 |
| 382 | 82.149 | 59.97 |
| 384 | 82.643 | 60.271 |
| 386 | 83.136 | 60.573 |
| 388 | 83.626 | 60.875 |
| 390 | 84.115 | 61.177 |
| 392 | 84.602 | 61.48 |
| 394 | 85.088 | 61.782 |
| 396 | 85.572 | 62.086 |
| 398 | 86.055 | 62.389 |
| 400 | 86.537 | 62.693 |

Figure S9 | Same as Figure S8 in which upper bound of blue fitness function is 20.

**Model Version 5-3**

V5-3 Functions

|  |  |  |
| --- | --- | --- |
| 0 | 5.13478e-16 | 5.37764e-16 |
| 1 | 0.00365645 | 0.00564835 |
| 2 | 0.007758 | 0.0123587 |
| 3 | 0.0123587 | 0.02033 |
| 4 | 0.017519 | 0.0297984 |
| 5 | 0.0233069 | 0.0410439 |
| 6 | 0.0297984 | 0.0543982 |
| 7 | 0.0370786 | 0.0702543 |
| 8 | 0.0452429 | 0.0890775 |
| 9 | 0.0543982 | 0.111418 |
| 10 | 0.0646638 | 0.137927 |
| 11 | 0.0761737 | 0.169371 |
| 12 | 0.0890775 | 0.206657 |
| 13 | 0.103543 | 0.25085 |
| 14 | 0.119756 | 0.303204 |
| 15 | 0.137927 | 0.365187 |
| 16 | 0.158288 | 0.438518 |
| 17 | 0.181102 | 0.525202 |
| 18 | 0.206657 | 0.627567 |
| 19 | 0.235278 | 0.748307 |
| 20 | 0.267327 | 0.890523 |
| 21 | 0.303204 | 1.05776 |
| 22 | 0.343355 | 1.25404 |
| 23 | 0.388277 | 1.48388 |
| 24 | 0.438518 | 1.75231 |
| 25 | 0.494687 | 2.06484 |
| 26 | 0.557456 | 2.42741 |
| 27 | 0.627567 | 2.84627 |
| 28 | 0.705836 | 3.32783 |
| 29 | 0.793161 | 3.87841 |
| 30 | 0.890523 | 4.50392 |
| 31 | 0.998995 | 5.20945 |
| 32 | 1.11974 | 5.99882 |
| 33 | 1.25404 | 6.874 |
| 34 | 1.40323 | 7.83459 |
| 35 | 1.5688 | 8.87735 |
| 36 | 1.75231 | 9.99583 |
| 37 | 1.95541 | 11.1802 |
| 38 | 2.17983 | 12.4175 |
| 39 | 2.42741 | 13.6917 |
| 40 | 2.7 | 14.985 |
| 41 | 2.99951 | 16.2783 |
| 42 | 3.32783 | 17.5525 |
| 43 | 3.68683 | 18.7898 |
| 44 | 4.07831 | 19.9742 |
| 45 | 4.50392 | 21.0927 |
| 46 | 4.96512 | 22.1354 |
| 47 | 5.46312 | 23.096 |
| 48 | 5.99882 | 23.9712 |
| 49 | 6.57271 | 24.7605 |
| 50 | 7.1848 | 25.4661 |
| 51 | 7.83459 | 26.0916 |
| 52 | 8.52096 | 26.6422 |
| 53 | 9.24218 | 27.1237 |
| 54 | 9.99583 | 27.5426 |
| 55 | 10.7788 | 27.9052 |
| 56 | 11.5875 | 28.2177 |
| 57 | 12.4175 | 28.4861 |
| 58 | 13.2639 | 28.716 |
| 59 | 14.1216 | 28.9122 |
| 60 | 14.985 | 29.0795 |
| 61 | 15.8484 | 29.2217 |
| 62 | 16.7061 | 29.3424 |
| 63 | 17.5525 | 29.4448 |
| 64 | 18.3825 | 29.5315 |
| 65 | 19.1912 | 29.6048 |
| 66 | 19.9742 | 29.6668 |
| 67 | 20.7278 | 29.7191 |
| 68 | 21.449 | 29.7633 |
| 69 | 22.1354 | 29.8006 |
| 70 | 22.7852 | 29.8321 |
| 71 | 23.3973 | 29.8586 |
| 72 | 23.9712 | 29.8809 |
| 73 | 24.5069 | 29.8997 |
| 74 | 25.0049 | 29.9156 |
| 75 | 25.4661 | 29.929 |
| 76 | 25.8917 | 29.9402 |
| 77 | 26.2832 | 29.9497 |
| 78 | 26.6422 | 29.9576 |
| 79 | 26.9705 | 29.9644 |
| 80 | 27.27 | 29.97 |
| 81 | 27.5426 | 29.9748 |
| 82 | 27.7902 | 29.9788 |
| 83 | 28.0146 | 29.9821 |
| 84 | 28.2177 | 29.985 |
| 85 | 28.4012 | 29.9873 |
| 86 | 28.5668 | 29.9893 |
| 87 | 28.716 | 29.991 |
| 88 | 28.8503 | 29.9925 |
| 89 | 28.971 | 29.9937 |
| 90 | 29.0795 | 29.9947 |
| 91 | 29.1768 | 29.9955 |
| 92 | 29.2642 | 29.9962 |
| 93 | 29.3424 | 29.9968 |
| 94 | 29.4125 | 29.9973 |
| 95 | 29.4753 | 29.9977 |
| 96 | 29.5315 | 29.9981 |
| 97 | 29.5817 | 29.9984 |
| 98 | 29.6266 | 29.9987 |
| 99 | 29.6668 | 29.9989 |
| 100 | 29.7027 | 29.9991 |
| 101 | 29.7347 | 29.9992 |
| 102 | 29.7633 | 29.9993 |
| 103 | 29.7889 | 29.9994 |
| 104 | 29.8117 | 29.9995 |
| 105 | 29.8321 | 29.9996 |
| 106 | 29.8502 | 29.9997 |
| 107 | 29.8665 | 29.9997 |
| 108 | 29.8809 | 29.9998 |
| 109 | 29.8938 | 29.9998 |
| 110 | 29.9053 | 29.9998 |
| 111 | 29.9156 | 29.9999 |
| 112 | 29.9248 | 29.9999 |
| 113 | 29.9329 | 29.9999 |
| 114 | 29.9402 | 29.9999 |
| 115 | 29.9467 | 29.9999 |
| 116 | 29.9525 | 29.9999 |
| 117 | 29.9576 | 29.9999 |
| 118 | 29.9622 | 30 |
| 119 | 29.9663 | 30 |
| 120 | 29.97 | 30 |
| 121 | 29.9733 | 30 |
| 122 | 29.9762 | 30 |
| 123 | 29.9788 | 30 |
| 124 | 29.9811 | 30 |
| 125 | 29.9831 | 30 |
| 126 | 29.985 | 30 |
| 127 | 29.9866 | 30 |
| 128 | 29.988 | 30 |
| 129 | 29.9893 | 30 |
| 130 | 29.9905 | 30 |
| 131 | 29.9915 | 30 |
| 132 | 29.9925 | 30 |
| 133 | 29.9933 | 30 |
| 134 | 29.994 | 30 |
| 135 | 29.9947 | 30 |
| 136 | 29.9952 | 30 |
| 137 | 29.9958 | 30 |
| 138 | 29.9962 | 30 |
| 139 | 29.9966 | 30 |
| 140 | 29.997 | 30 |
| 141 | 29.9973 | 30 |
| 142 | 29.9976 | 30 |
| 143 | 29.9979 | 30 |
| 144 | 29.9981 | 30 |
| 145 | 29.9983 | 30 |
| 146 | 29.9985 | 30 |
| 147 | 29.9987 | 30 |
| 148 | 29.9988 | 30 |
| 149 | 29.9989 | 30 |
| 150 | 29.9991 | 30 |
| 151 | 29.9992 | 30 |
| 152 | 29.9992 | 30 |
| 153 | 29.9993 | 30 |
| 154 | 29.9994 | 30 |
| 155 | 29.9995 | 30 |
| 156 | 29.9995 | 30 |
| 157 | 29.9996 | 30 |
| 158 | 29.9996 | 30 |
| 159 | 29.9997 | 30 |
| 160 | 29.9997 | 30 |
| 161 | 29.9997 | 30 |
| 162 | 29.9998 | 30 |
| 163 | 29.9998 | 30 |
| 164 | 29.9998 | 30 |
| 165 | 29.9998 | 30 |
| 166 | 29.9998 | 30 |
| 167 | 29.9999 | 30 |
| 168 | 29.9999 | 30 |
| 169 | 29.9999 | 30 |
| 170 | 29.9999 | 30 |
| 171 | 29.9999 | 30 |
| 172 | 29.9999 | 30 |
| 173 | 29.9999 | 30 |
| 174 | 29.9999 | 30 |
| 175 | 29.9999 | 30 |
| 176 | 30 | 30 |
| 177 | 30 | 30 |
| 178 | 30 | 30 |
| 179 | 30 | 30 |
| 180 | 30 | 30 |
| 181 | 30 | 30 |
| 182 | 30 | 30 |
| 183 | 30 | 30 |
| 184 | 30 | 30 |
| 185 | 30 | 30 |
| 186 | 30 | 30 |
| 187 | 30 | 30 |
| 188 | 30 | 30 |
| 189 | 30 | 30 |
| 190 | 30 | 30 |
| 191 | 30 | 30 |
| 192 | 30 | 30 |
| 193 | 30 | 30 |
| 194 | 30 | 30 |
| 195 | 30 | 30 |
| 196 | 30 | 30 |
| 197 | 30 | 30 |
| 198 | 30 | 30 |
| 199 | 30 | 30 |

V5-3 Equal

|  |  |  |
| --- | --- | --- |
| 2 | 0 | 1 |
| 4 | 0 | 1 |
| 6 | 0 | 1 |
| 8 | 0 | 1 |
| 10 | 0 | 1 |
| 12 | 0 | 1 |
| 14 | 0 | 1 |
| 16 | 0 | 1 |
| 18 | 0 | 1 |
| 20 | 0 | 1 |
| 22 | 0 | 1 |
| 24 | 0 | 1 |
| 26 | 0 | 1 |
| 28 | 0 | 1 |
| 30 | 0 | 1 |
| 32 | 0 | 1 |
| 34 | 0 | 1 |
| 36 | 0 | 1 |
| 38 | 0 | 1 |
| 40 | 0 | 1 |
| 42 | 0 | 1 |
| 44 | 0 | 1 |
| 46 | 0 | 1 |
| 48 | 0 | 1 |
| 50 | 0 | 1 |
| 52 | 0 | 1 |
| 54 | 0 | 1 |
| 56 | 0 | 1 |
| 58 | 0 | 1 |
| 60 | 0 | 1 |
| 62 | 0 | 1 |
| 64 | 0 | 1 |
| 66 | 0 | 1 |
| 68 | 0 | 1 |
| 70 | 0 | 1 |
| 72 | 0 | 1 |
| 74 | 0 | 1 |
| 76 | 0 | 1 |
| 78 | 0 | 1 |
| 80 | 0 | 1 |
| 82 | 0 | 2 |
| 84 | 0 | 2 |
| 86 | 0 | 2 |
| 88 | 0 | 2 |
| 90 | 0 | 2 |
| 92 | 0 | 2 |
| 94 | 0 | 2 |
| 96 | 0 | 2 |
| 98 | 0 | 2 |
| 100 | 0 | 2 |
| 102 | 0 | 2 |
| 104 | 0 | 2 |
| 106 | 0 | 2 |
| 108 | 0 | 2 |
| 110 | 0 | 2 |
| 112 | 0 | 2 |
| 114 | 0 | 2 |
| 116 | 0 | 2 |
| 118 | 0 | 2 |
| 120 | 0 | 2 |
| 122 | 0 | 2 |
| 124 | 0 | 2 |
| 126 | 0 | 2 |
| 128 | 0 | 2 |
| 130 | 0 | 2 |
| 132 | 0 | 3 |
| 134 | 0 | 3 |
| 136 | 0 | 3 |
| 138 | 0 | 3 |
| 140 | 0 | 3 |
| 142 | 0 | 3 |
| 144 | 0 | 3 |
| 146 | 0 | 3 |
| 148 | 0 | 3 |
| 150 | 0 | 3 |
| 152 | 0 | 3 |
| 154 | 0 | 3 |
| 156 | 0 | 3 |
| 158 | 0 | 3 |
| 160 | 0 | 3 |
| 162 | 0 | 3 |
| 164 | 0 | 3 |
| 166 | 0 | 3 |
| 168 | 0 | 3 |
| 170 | 0 | 3 |
| 172 | 0 | 3 |
| 174 | 0 | 3 |
| 176 | 0 | 3 |
| 178 | 0 | 3 |
| 180 | 0 | 3 |
| 182 | 0 | 3 |
| 184 | 0 | 4 |
| 186 | 0 | 4 |
| 188 | 0 | 4 |
| 190 | 0 | 4 |
| 192 | 0 | 4 |
| 194 | 0 | 4 |
| 196 | 0 | 4 |
| 198 | 0 | 4 |
| 200 | 0 | 4 |
| 202 | 0 | 4 |
| 204 | 0 | 4 |
| 206 | 0 | 4 |
| 208 | 0 | 4 |
| 210 | 0 | 4 |
| 212 | 0 | 4 |
| 214 | 0 | 4 |
| 216 | 0 | 4 |
| 218 | 0 | 4 |
| 220 | 0 | 4 |
| 222 | 0 | 4 |
| 224 | 0 | 4 |
| 226 | 0 | 4 |
| 228 | 0 | 4 |
| 230 | 0 | 4 |
| 232 | 0 | 4 |
| 234 | 0 | 4 |
| 236 | 0 | 5 |
| 238 | 0 | 5 |
| 240 | 0 | 5 |
| 242 | 0 | 5 |
| 244 | 0 | 5 |
| 246 | 0 | 5 |
| 248 | 0 | 5 |
| 250 | 0 | 5 |
| 252 | 0 | 5 |
| 254 | 0 | 5 |
| 256 | 0 | 5 |
| 258 | 0 | 5 |
| 260 | 0 | 5 |
| 262 | 0 | 5 |
| 264 | 0 | 5 |
| 266 | 0 | 5 |
| 268 | 0 | 5 |
| 270 | 0 | 5 |
| 272 | 0 | 5 |
| 274 | 0 | 5 |
| 276 | 0 | 5 |
| 278 | 0 | 5 |
| 280 | 0 | 5 |
| 282 | 0 | 5 |
| 284 | 0 | 5 |
| 286 | 0 | 5 |
| 288 | 0 | 5 |
| 290 | 0 | 5 |
| 292 | 0 | 5 |
| 294 | 0 | 5 |
| 296 | 0 | 5 |
| 298 | 0 | 5 |
| 300 | 0 | 5 |
| 302 | 0 | 5 |
| 304 | 0 | 5 |
| 306 | 0 | 5 |
| 308 | 0 | 5 |
| 310 | 0 | 5 |
| 312 | 0 | 5 |
| 314 | 0 | 5 |
| 316 | 1 | 5 |
| 318 | 1 | 5 |
| 320 | 1 | 5 |
| 322 | 1 | 5 |
| 324 | 1 | 5 |
| 326 | 1 | 5 |
| 328 | 1 | 5 |
| 330 | 1 | 5 |
| 332 | 1 | 5 |
| 334 | 1 | 5 |
| 336 | 1 | 5 |
| 338 | 1 | 5 |
| 340 | 1 | 5 |
| 342 | 1 | 5 |
| 344 | 1 | 5 |
| 346 | 1 | 5 |
| 348 | 1 | 5 |
| 350 | 1 | 5 |
| 352 | 1 | 5 |
| 354 | 1 | 5 |
| 356 | 1 | 5 |
| 358 | 1 | 5 |
| 360 | 1 | 5 |
| 362 | 1 | 5 |
| 364 | 1 | 5 |
| 366 | 1 | 5 |
| 368 | 1 | 5 |
| 370 | 1 | 5 |
| 372 | 1 | 5 |
| 374 | 1 | 5 |
| 376 | 1 | 5 |
| 378 | 1 | 5 |
| 380 | 1 | 5 |
| 382 | 1 | 5 |
| 384 | 1 | 5 |
| 386 | 1 | 5 |
| 388 | 1 | 5 |
| 390 | 1 | 5 |
| 392 | 1 | 5 |
| 394 | 2 | 5 |
| 396 | 2 | 5 |
| 398 | 2 | 5 |
| 400 | 2 | 5 |

Figure S10 | Same as Figure S8 in which upper bound of blue fitness function is 30.

**Model Version 5-4**

Figure S11 | Same as Figure S8 in which upper bound of blue fitness function is 40.

**Model Version 5-5**

Figure S12 | Same as Figure S8 in which upper bound of blue fitness function is 50.

**Model Version 5-6**

Figure S13 | Same as Figure S8 in which upper bound of blue fitness function is 60.

**Model Version 5-7**

Figure S14 | Same as Figure S8 in which upper bound of blue fitness function is 70.

**Model Version 5-8**

Figure S15 | Same as Figure S8 in which upper bound of blue fitness function is 80.

**Model Version 5-9**

Figure S16 | Same as Figure S8 in which upper bound of blue fitness function is 90.

**Model Version 5-10**

Figure S17 | Same as Figure S8 in which upper bound of blue fitness function is 100.

**Model Version 5-11**

Figure S18 | Same as Figure S8 in which upper bound of blue fitness function is 110.

**Model Version 5-12**

Figure S19 | Same as Figure S8 in which upper bound of blue fitness function is 120.

**Model Version 5-13**

Figure S20 | Same as Figure S8 in which upper bound of blue fitness function is 130.

**Model Version 5-14**

Figure S21 | Same as Figure S8 in which upper bound of blue fitness function is 140.

**Model Version 5-15**

Figure S22 | Same as Figure S8 in which upper bound of blue fitness function is 150.
